# Na_2_CO_3_-Responsive Photosynthetic and ROS Scavenging Mechanisms in Chloroplasts of Alkaligrass Revealed by Phosphoproteomics

**DOI:** 10.1101/871046

**Authors:** Jinwei Suo, Heng Zhang, Qi Zhao, Nan Zhang, Yongxue Zhang, Ying Li, Baohua Song, Juanjuan Yu, Jian’guo Cao, Tai Wang, Ji Luo, Lihai Guo, Jun Ma, Xumin Zhang, Yimin She, Lianwei Peng, Weimin Ma, Siyi Guo, Yuchen Miao, Sixue Chen, Zhi Qin, Shaojun Dai

## Abstract

Alkali-salinity exerts severe osmotic, ionic and high-pH stresses to plants. To understand the alkali-salinity responsive mechanisms underlying photosynthetic modulation and reactive oxygen species (ROS) homeostasis, physiological and diverse quantitative proteomics analyses of alkaligrass (*Puccinellia tenuiflora*) under Na_2_CO_3_ stress were conducted. In addition, Western blot, real-time PCR, and transgenic techniques were applied to validate the proteomic results and test the functions of the Na_2_CO_3_-responsive proteins. A total of 104 and 102 Na_2_CO_3_-responsive proteins were identified in leaves and chloroplasts, respectively. In addition, 84 Na_2_CO_3_-responsive phosphoproteins were identified, including 56 new phosphorylation sites in 56 phosphoproteins from chloroplasts, which are crucial for the regulation of photosynthesis, ion transport, signal transduction and energy homeostasis. A full-length *PtFBA* encoding an alkaligrass chloroplastic fructose-bisphosphate aldolase (FBA) was overexpressed in wild-type cells of cyanobacterium *Synechocystis* sp. Strain PCC 6803, leading to enhanced Na_2_CO_3_ tolerance. All these results indicate that thermal dissipation, state transition, cyclic electron transport, photorespiration, repair of photosystem (PS) II, PSI activity, and ROS homeostasis were altered in response to Na_2_CO_3_ stress, and they have improved our understanding of the Na_2_CO_3_-responsive mechanisms in halophytes.

## Introduction

Soil salinization and alkalization frequently occur simultaneously. In northeast China, more than 70% of the land area has become alkaline grassland [1]. Alkali-salinity is one of the most severe abiotic stresses, limiting the productivity and geographical distribution of plants. Saline-alkali stress exerts osmotic stress and ion damage, as well as high-pH stress to plants [2]. However, little attention has been given to the sophisticated tolerance mechanisms underlying plant response to saline-alkali (*e.g.* Na_2_CO_3_ and NaHCO_3_) stresses [3,4]. As the organelle for photosynthesis, chloroplasts are extremely susceptible to saline-alkali stress [5]. Excessive accumulation of Na^+^ reduces the CO_2_ diffusion through stomata and mesophyll, negatively affecting plant photosynthesis [6]. As a consequence, excessive excitation energy causes generation of reactive oxygen species (ROS), resulting in damage to the thylakoid membrane [6].

Current high-throughput proteomic approaches are powerful to untangle the complicated mechanisms of chloroplast development, metabolism and stress response [7-10]. More than 522 NaCl-responsive chloroplast proteins were found in different plant species, such as tomato (*Solanum lycopersicum*) [11], wheat (*Triticum aestivum*) [12], and other plant species [13-18]. The presence of these proteins indicate that the light harvesting, photosynthetic electron transfer, carbon assimilation, ROS homeostasis, energy metabolism, signaling, and membrane trafficking were modulated in chloroplasts in response to NaCl stress. However, only about 53 salinity-responsive genes encoding chloroplast proteins have been characterized [5], which are insufficient to address the sophisticated salinity-responsive networks in chloroplasts. Additionally, NaCl stress altered phosphorylation levels of several chloroplast proteins in Arabidopsis [19,20], *Brachypodium distachyon* [21] and sugar beet (*Beta vulgaris*) [22], implying that state transition, PSII damage repair, thermal dissipation, and thylakoid membrane organization were crucial for plant acclimation to salt stress [23]. However, the critical roles of reversible protein phosphorylation in salinity-/alkali-responsive metabolic networks are virtually unknown.

Alkaligrass (*Puccinellia tenuiflora*) is a monocotyledonous halophyte species belonging to the Gramineae, and is widely distributed in the Songnen Plain in Northeastern China. It has strong ability to survive in extreme saline-alkali soil (pH range of 9-10). Several salinity-/alkali-responsive genes and/or proteins in leaves and roots of alkaligrass have been reported [24-26]. A previous transcriptomic study also revealed that a number of Na_2_CO_3_ responsive genes were overrepresented in metabolism, signal transduction, transcription, and cell rescue [27]. Despite this progress, the precise alkali-responsive mechanisms in chloroplasts are still poorly understood. Analyses of the photosynthetic and ROS scavenging mechanisms in chloroplasts regulated by the reversible protein phosphorylation and the expression of nuclear and chloroplast genes are critical for understanding the Na_2_CO_3_-responsive mechanisms in alkaligrass. In this study, we investigated the alkali-responsive characteristics in chloroplasts and leaves of alkaligrass. By integrative analyses of protein phosphorylation, patterns of protein abundance, gene expression, photosynthesis parameters, antioxidant enzyme activities, and chloroplast ultrastructure, we revealed several important Na_2_CO_3_-responsive strategies in the halophyte alkaligrass. These results have yielded important insights into the alkali-responsive mechanisms in halophytes.

## Results

### Na_2_CO_3_ treatment decreased seedling growth and biomass

Na_2_CO_3_ treatment clearly affected the morphology and biomass of alkaligrass seedlings. The leaves withered with the increase of Na_2_CO_3_ concentration and treatment time (Figure S1). The shoot length and relative water content decreased significantly at 200 mM Na_2_CO_3_ of 24 h after treatment (24 HAT200) (**Figure 1**A). The fresh and dry weights of leaves also clearly decreased under 200 mM Na_2_CO_3_ (Figure 1B).

**Figure 1.**
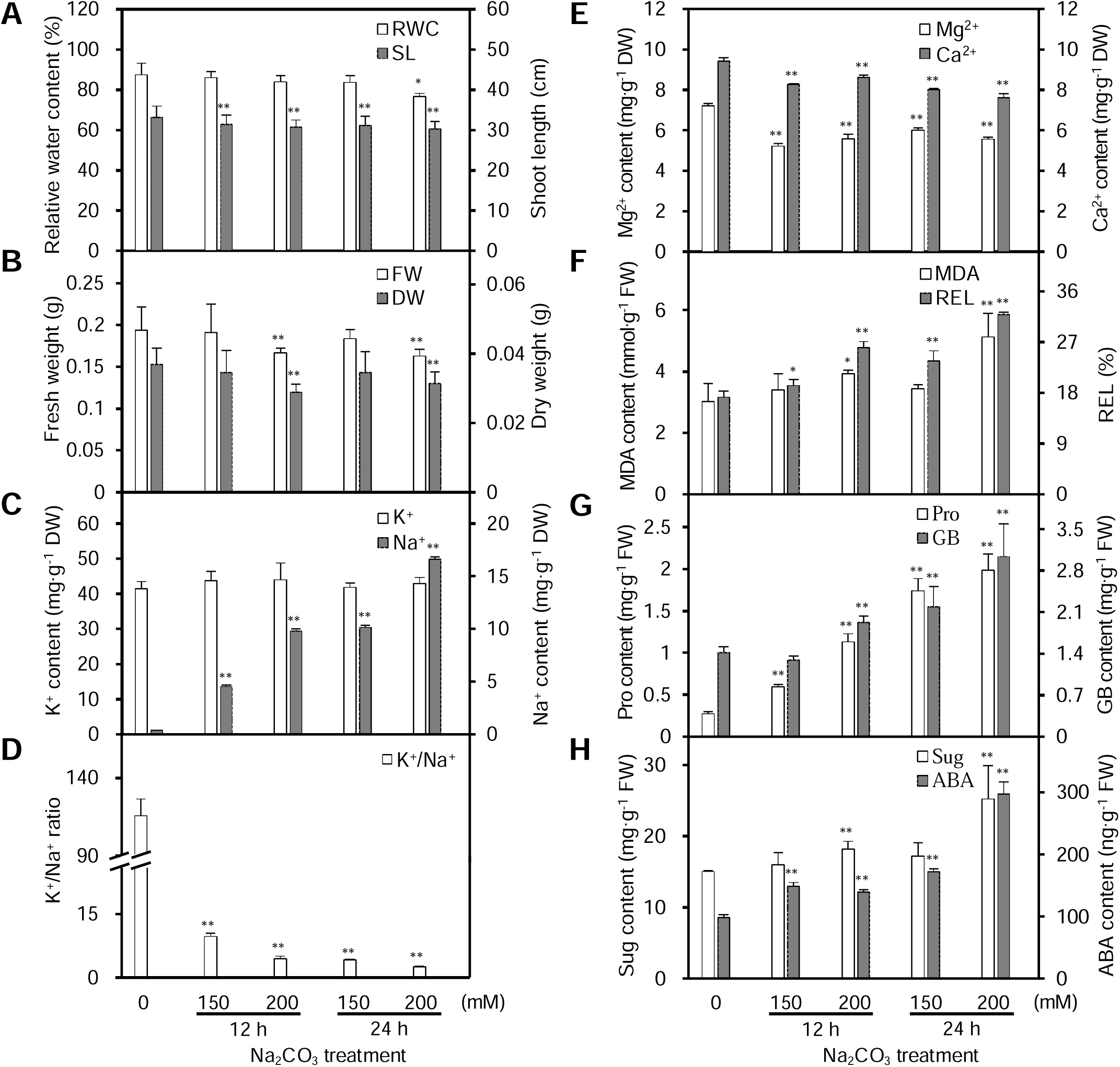
Leaf physiological characteristics in alkaligrass under Na_2_CO_3_ treatment. **A.** Relative water content (RWC) in leaves and shoot length (SL) of seedlings. **B.** Fresh weight (FW) and dry weight (DW) of leaves. **C.** K^+^ and Na^+^ contents. **D.** K^+^/Na^+^ ratio. **E.** Ca^2+^ and Mg^2+^ contents. **F.** Malondialdehyde (MDA) content and relative electrolyte leakage (REL). **G.** Proline (Pro) and glycine betaine (GB) contents. **H.** Soluble sugar (Sug) and abscisic acid (ABA) contents. The values were determined after plants were treated with 0, 150, and 200 mM Na_2_CO_3_ for 12 and 24 h, and were presented as means ± standard deviation (n ≥ 3), respectively. The asterisks indicate significant differences (*, P < 0.05; **, P < 0.01).

### Na_2_CO_3_ treatment changed ionic and osmotic homeostasis, cell membrane integrity, and abscisic acid (ABA) level

Na_2_CO_3_ treatment perturbed the ion and pH homeostasis in leaves. Na^+^ in leaves was gradually accumulated, but K^+^ content did not show obvious changes, resulting in the sharp decline of the K^+^/Na^+^ ratio (Figure 1C, D). In addition, the Mg^2+^ and Ca^2+^ contents gradually decreased under the Na_2_CO_3_ treatment (Figure 1E). Malondialdehyde content and relative electrolyte leakage significantly increased under different Na_2_CO_3_ treatments, indicating that the membrane integrity was affected by Na_2_CO_3_ treatment (Figure 1F). In addition, proline and glycine betaine gradually accumulated with the increase of Na_2_CO_3_ concentrations (Figure 1G), while the soluble sugar content only showed marked accumulation at 200 mM Na_2_CO_3_ (Figure 1H). The endogenous ABA content in leaves increased significantly (Figure 1H).

### Photosynthesis and chlorophyll (Chl) content decreased under Na_2_CO_3_

In seedlings, net photosynthetic rate, stomatal conductance, and transpiration rate (**Figure 2**A, B) gradually decreased, while the intercellular CO_2_ concentration did not exhibit obvious changes under the Na_2_CO_3_ (Figure 2B).

**Figure 2.**
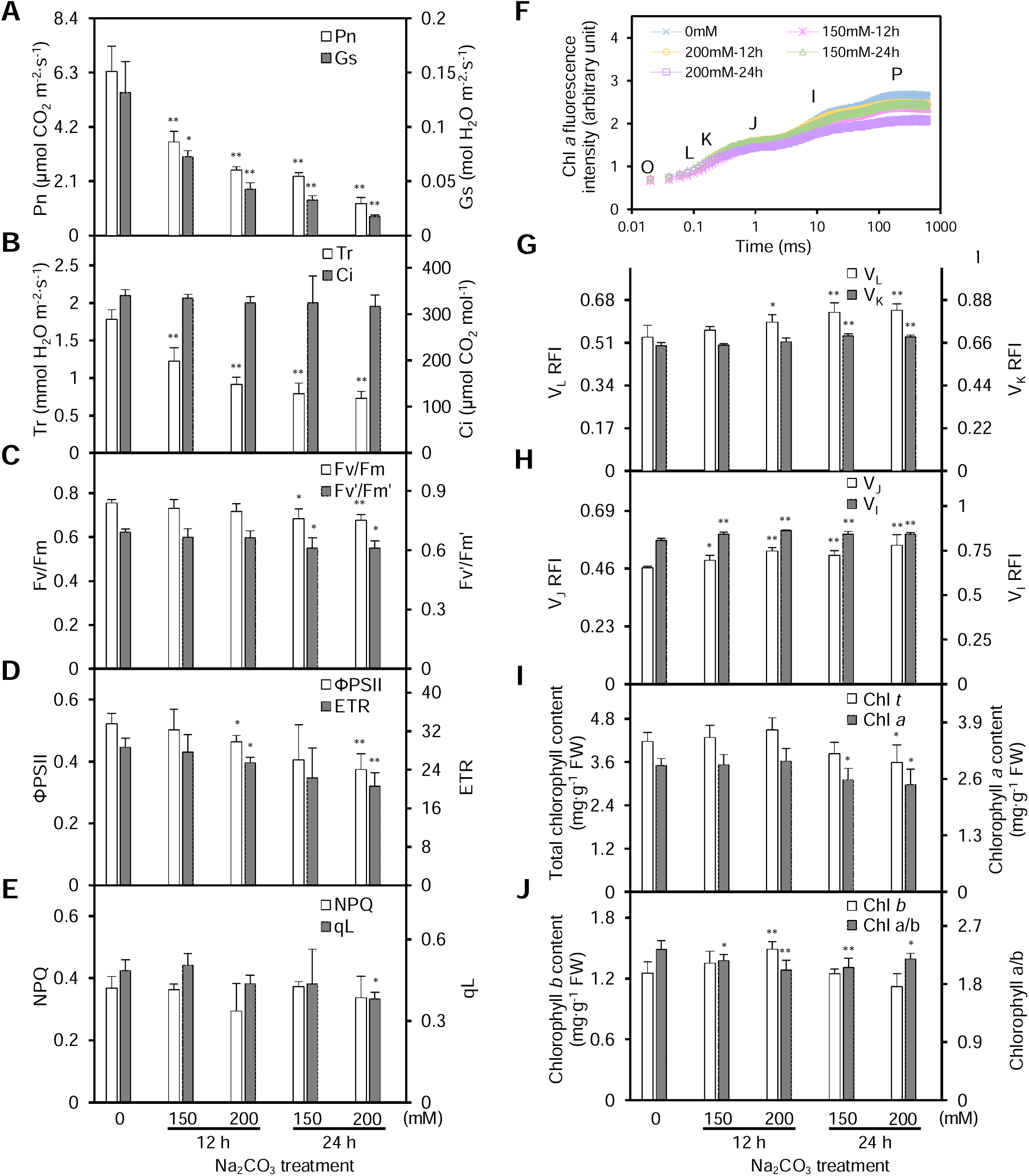
Photosynthetic characteristics of alkaligrass under Na_2_CO_3_ treatment. **A.** Net photosynthetic rate (Pn) and stomatal conductance (Gs). **B.** Transpiration rate (Tr) and intercellular CO_2_ concentration (Ci). **C.** Maximum quantum efficiency of PSII photochemistry (Fv/Fm) and PSII maximum efficiency (Fv’/Fm’). **D.** Actual PSII efficiency (*ϕ*PSII) and electron transport rate (ETR). **E.** Non-photochemical quenching (NPQ) and the fraction of open PSII centers (qL). **F.** Chlorophyll fluorescence OJIP transient, fluorescence intensity (F_t_) was recorded between 0.01 and 1000 ms time period. **G.** Relative fluorescence intensity (RFI) of band L (V_L_) and K (V_K_) after double normalization between the two fluorescence extreme F_O_ and F_K_, F_O_ and F_J_ phases: V_L_ = (F_L_ – F_O_)/(F_K_ – F_O_), V_K_ = (F_K_ – F_O_)/(F_J_ – F_O_). **H.** Relative fluorescence intensity (RFI) of step J (V_J_) and I (V_I_) after double normalization between the two fluorescence extreme F_O_ and F_P_ phases: V_J_ = (F_J_ – F_O_)/(F_P_ – F_O_), V_I_ = (F_I_ – F_O_)/(F_P_ – F_O_). **I.** Total chl and chl *a* contents. **J.** Chl *b* content and Chl *a*/*b* ratio. The values were determined after plants were treated with 0, 150 and 200 mM Na_2_CO_3_ for 12 and 24 h, and were presented as means ± standard deviation (n ≥ 3). The asterisks indicate significant differences (*, P < 0.05; **, P < 0.01).

To evaluate the photosynthetic performance, we investigated the changes of Chl fluorescence and the polyphasic fluorescence transients (OJIP). The maximum quantum efficiency of PSII photochemistry and PSII maximum efficiency significantly decreased at 24 HAT (Figure 2C), and the actual PSII efficiency and electron transport rate were declined remarkably at 200 mM Na_2_CO_3_ (Figure 2D). In addition, the non-photochemical quenching did not change and the fraction of open PSII centers significantly decreased at 24 HAT200 (Figure 2E). The fluorescence transient gradually decreased, reaching the lowest level at 24 HAT200 (Figure 2F). After normalization, the relative fluorescence intensities of V_L_ and V_K_, two specific indicators of thylakoid dissociation and oxygen-evolving complex (OEC) damage increased at 24 HAT (Figure 2G). V_J_ and V_I_, however, obviously increased. The relative variable fluorescence intensity of V_J_ and V_I_ can be considered as a measurement of the accumulation of Q_A_^-^ and the proportion of the Q_B_-non-reducing reaction center. This suggests that the accumulation of Q_A_^-^ and increased proportion of Q_B_-non-reducing reaction center in the Na_2_CO_3_-stressed leaves (Figure 2H). In addition, the contents of total Chl and Chl *a* decreased at 24 HAT (Figure 2I), and the ratio of Chl *a*/*b* also decreased under the Na_2_CO_3_ treatment (Figure 2J).

### Na_2_CO_3_ treatment affected chloroplast ultrastructure

Na_2_CO_3_ treatment changed the chloroplast ultrastructure in mesophyll cells and bundle sheath cells from lateral veins, minor veins and midveins (**Figure 3**). Under normal conditions, chloroplasts in mesophyll cells and bundle sheath cells exhibited long ellipsoidal or shuttle-shaped, double membrane compartment, with only a few osmophilia plastoglobules in the stroma (Figure 3A, F, K, and P). Thylakoids were dispersed in the chloroplasts, and the fully developed thylakoid membrane systems were well organized in grana and stromal lamellae (Figure 3A, F, K, and P). At 12 HAT, slight swelling of the chloroplast stroma occurred, and the membranes of the individual thylakoid fused, eliminating the intraspace (Figure 3C, H, and M). While at 24 HAT, chloroplast volume increased obviously to become round-shaped. The thylakoid membrane systems in various cells became distorted and incomplete, showing a dilated intraspace (Figure 3D, E, J, and O). The size and number of grana somewhat decreased, and some grana completely disappeared (Figure 3D, E, and J). At 24 HAT200, numerous plastoglobules were observed in chloroplasts, and the size and number of plastoglobules appeared to be Na_2_CO_3_ concentration-dependent (Figure 3). This implies that lipid peroxidation-mediated destruction of the thylakoid membranes takes place in chloroplasts. In addition, the aforementioned changes in thylakoids appeared more drastic in mesophyll cells than in the bundle sheath cells (Figure 3D, E).

**Figure 3.**
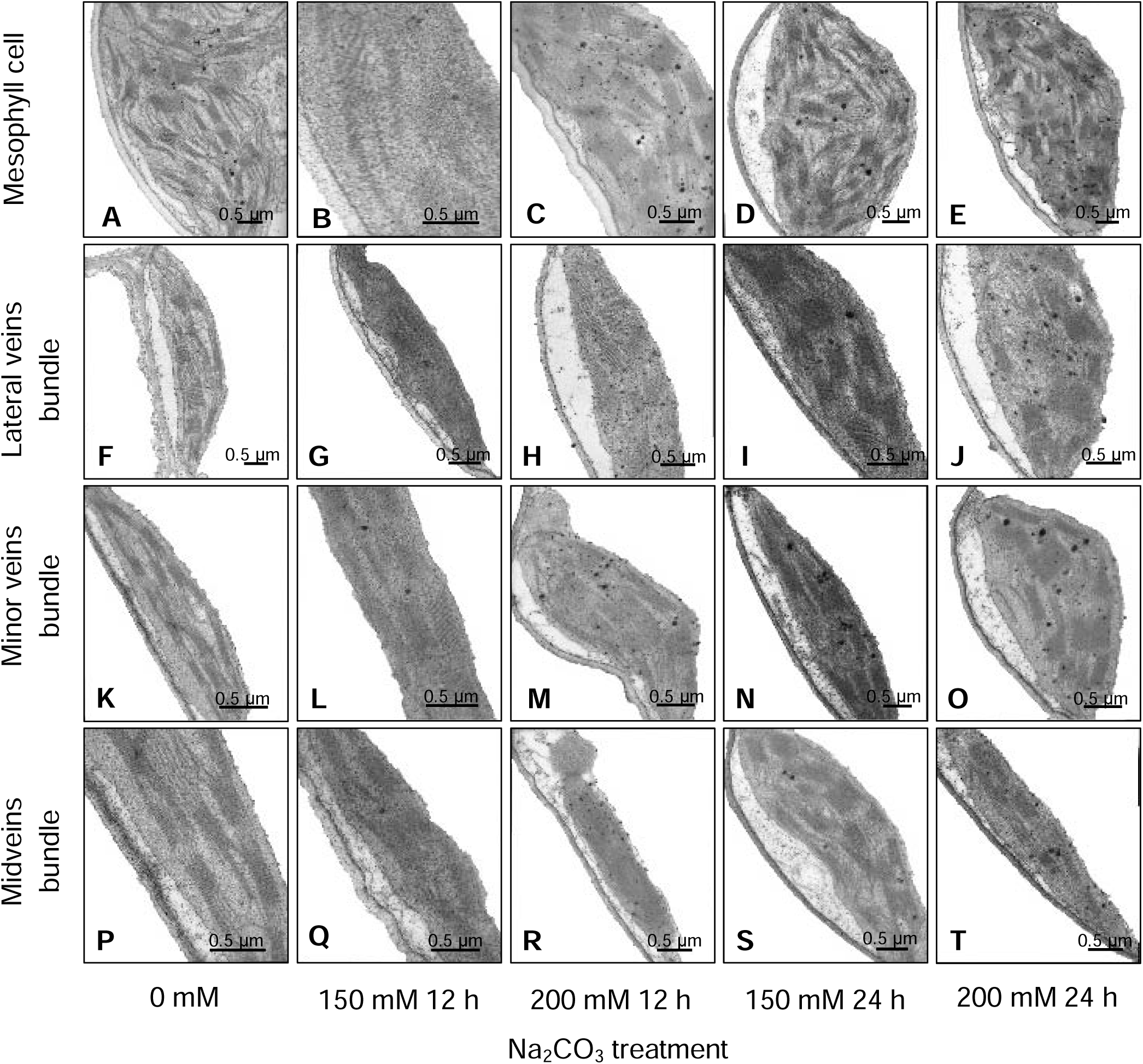
Ultrastructure of chloroplasts in alkaligrass leaves under Na_2_CO_3_ treatment. **A-E.** Chloroplasts in mesophyll cells. **F-J.** Chloroplasts in lateral veins bundle sheath cells. **K-O.** Chloroplasts in minor veins bundle sheath cells. **P-T.** Chloroplasts in midvein bundle sheath cells. Bar = 0.5 μm.

### Na_2_CO_3_ treatment changed antioxidant enzymes in leaves and isolated chloroplasts

To evaluate the level of oxidative stress in leaves and chloroplasts under Na_2_CO_3_ treatment, the O_2_^-^ generation rate, H_2_O_2_ content and four metabolites (i.e. ascorbate (AsA), dehydroascorbate (DHA), glutathione (GSH), and oxidized glutathione (GSSG)), and the activities of nine antioxidant enzymes in ROS scavenging system were monitored (**Figure 4**).

In leaves, the O_2_^-^ generation rate and H_2_O_2_ content increased under the Na_2_CO_3_ treatment (Figure 4A). The contents of several metabolites (e.g., reduced AsA, DHA, and GSSG) did not change, but GSH increased at 24 HAT (Figure 4B, C). Importantly, the activities of superoxide dismutase (SOD), catalase (CAT), and dehydroascorbate reductase (DHAR) decreased at 24 HAT, and CAT activity was inhibited at 12 HAT200 (Figure 4D, E and F). The activities of peroxidase (POD), ascorbate peroxidase (APX), monodehydroascorbate reductase (MDHAR), glutathione peroxidase (GPX), glutathione reductase (GR), and glutathione S-transferase (GST) showed increased patterns under the Na_2_CO_3_ treatment (Figure 4D, E, F, G and H). These results indicate that the superoxide dismutation by SOD and reduction of H_2_O_2_ to H_2_O by CAT decreased, but the APX/POD pathway, AsA-GSH cycle, and GPX pathway were enhanced to cope with the Na_2_CO_3_-induced oxidative stress.

**Figure 4.**
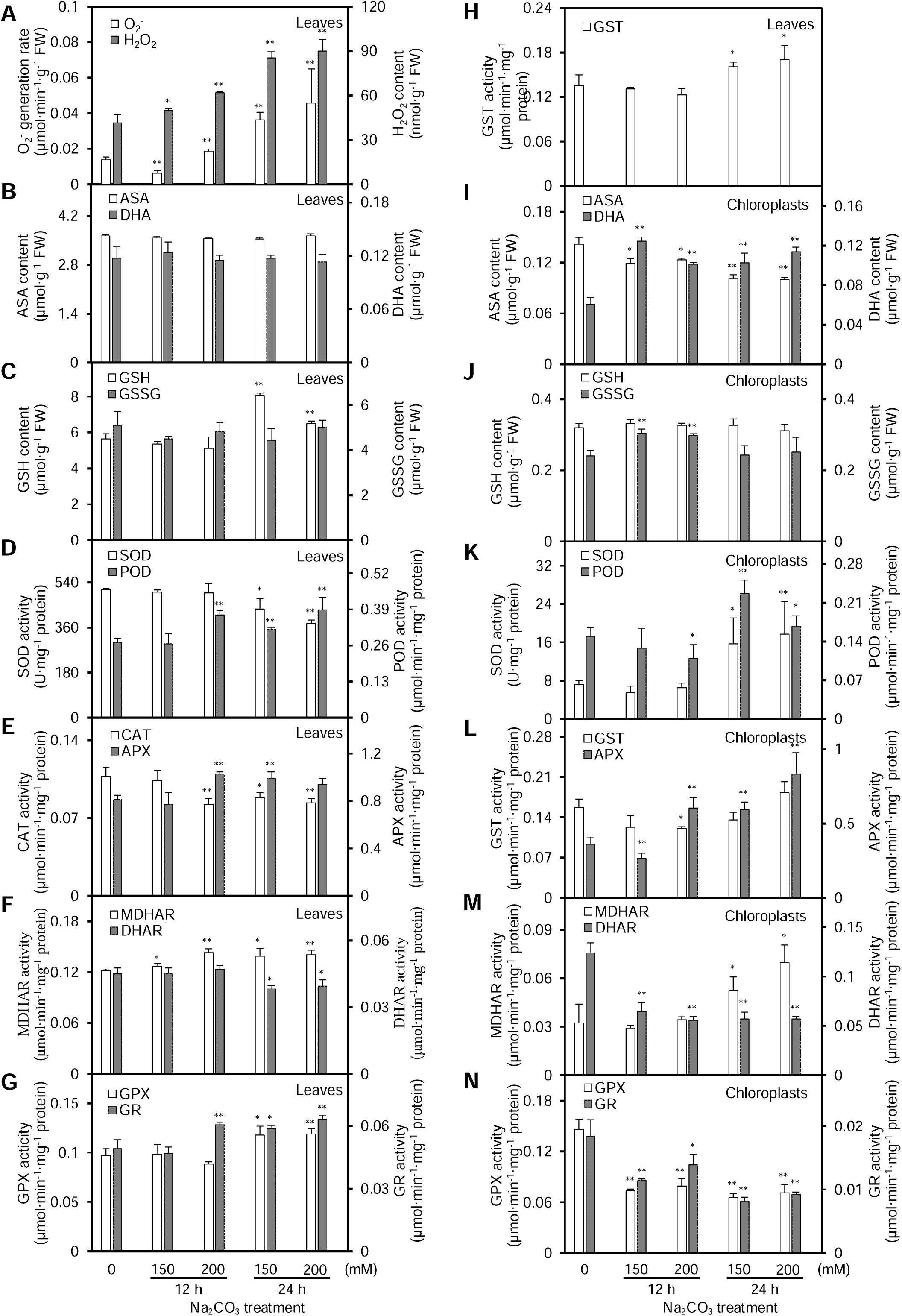
Effects of Na_2_CO_3_ treatments on antioxidant systems in leaves and chloroplasts of alkaligrass. **A.** O_2_^-^ generation rate and H_2_O_2_ content in leaves. **B.** Ascorbate (AsA) and dehydroascorbate (DHA) contents in leaves. **C.** Glutathione (GSH) and oxidized glutathione (GSSG) contents in leaves. **D.** Superoxide dismutase (SOD) and peroxidase (POD) activities in leaves. **E.** Catalase (CAT) and ascorbate peroxidase (APX) activities in leaves. **F.** Monodehydroascorbate reductase (MDHAR) and dehydroascorbate reductase (DHAR) activities in leaves. **G.** Glutathione peroxidase (GPX) and glutathione reductase (GR) activities in leaves. **H.** Glutathione S-transferase (GST) activity in leaves. **I.** AsA and DHA contents in chloroplasts. **J.** GSH and GSSG contents in chloroplasts. **K.** SOD and POD activities in chloroplasts. **L.** GST and APX activities in chloroplasts. **M.** MDHAR and DHAR activities in chloroplasts. **N.** GPX and GR activities in chloroplasts. The values were determined after plants were treated with 0, 150 and 200 mM Na_2_CO_3_ for 12 and 24 h, and were presented as means ± standard deviation (n ≥ 3). The asterisks indicate significant differences (*, P < 0.05; **, P < 0.01).

We isolated chloroplasts with high purity for activity and proteomics analyses (Figure S2A, B). In isolated chloroplasts, the AsA content decreased, but the DHA content increased under Na_2_CO_3_. The contents of GSH and GSSG stayed at relative stable levels at 24 HAT (Figure 4I, J). The activities of SOD, POD, APX, and MDHAR increased at 24 HAT, and APX activity increased at 12 HAT (Figure 4K, L, and M). The activities of DHAR, GPX, and GR were inhibited, however GST activity did not change significantly under Na_2_CO_3_ (Figure 4L, M and N). These results indicate that ROS in chloroplasts were mainly dismutated by SOD, and subsequently reduced in APX/POD pathway under the Na_2_CO_3_ treatment.

### Na_2_CO_3_-responsive proteome revealed modulation of photosynthesis and ROS scavenging to cope with the stress

A total of 104 Na_2_CO_3_-responsive proteins in leaves were identified and classified into ten functional categories (**Figure 5**A, Figure S3, Tables S1-S3). Cluster analysis generated two main clusters (Figure 5B). In Cluster I, 63 Na_2_CO_3_-decreased proteins were involved in photosynthesis, carbohydrate and energy metabolism, protein synthesis and turnover, and cell wall metabolism. In Cluster II, 41 Na_2_CO_3_-increased proteins were related to energy metabolism, Chl metabolism, membrane and transport, and cell cycle. Importantly, subcellular localization prediction suggested that 63 proteins (60.6%) were specially localized in chloroplasts, and six proteins (6%) in either chloroplasts or other subcellular locations (Figure 5C; Tables S3 and S4). The changes of 37 photosynthesis-related proteins indicate that the balance of excitation energy between PSII and PSI was disrupted and the efficiency of electron transfer and CO_2_ assimilation were inhibited. In contrast, photorespiration was induced under the Na_2_CO_3_ treatment. Aside from this, changes of 11 proteins involved in ROS and ion homeostasis as well as signaling pathway were triggered under Na_2_CO_3_ treatment (Table S3).

**Figure 5.**
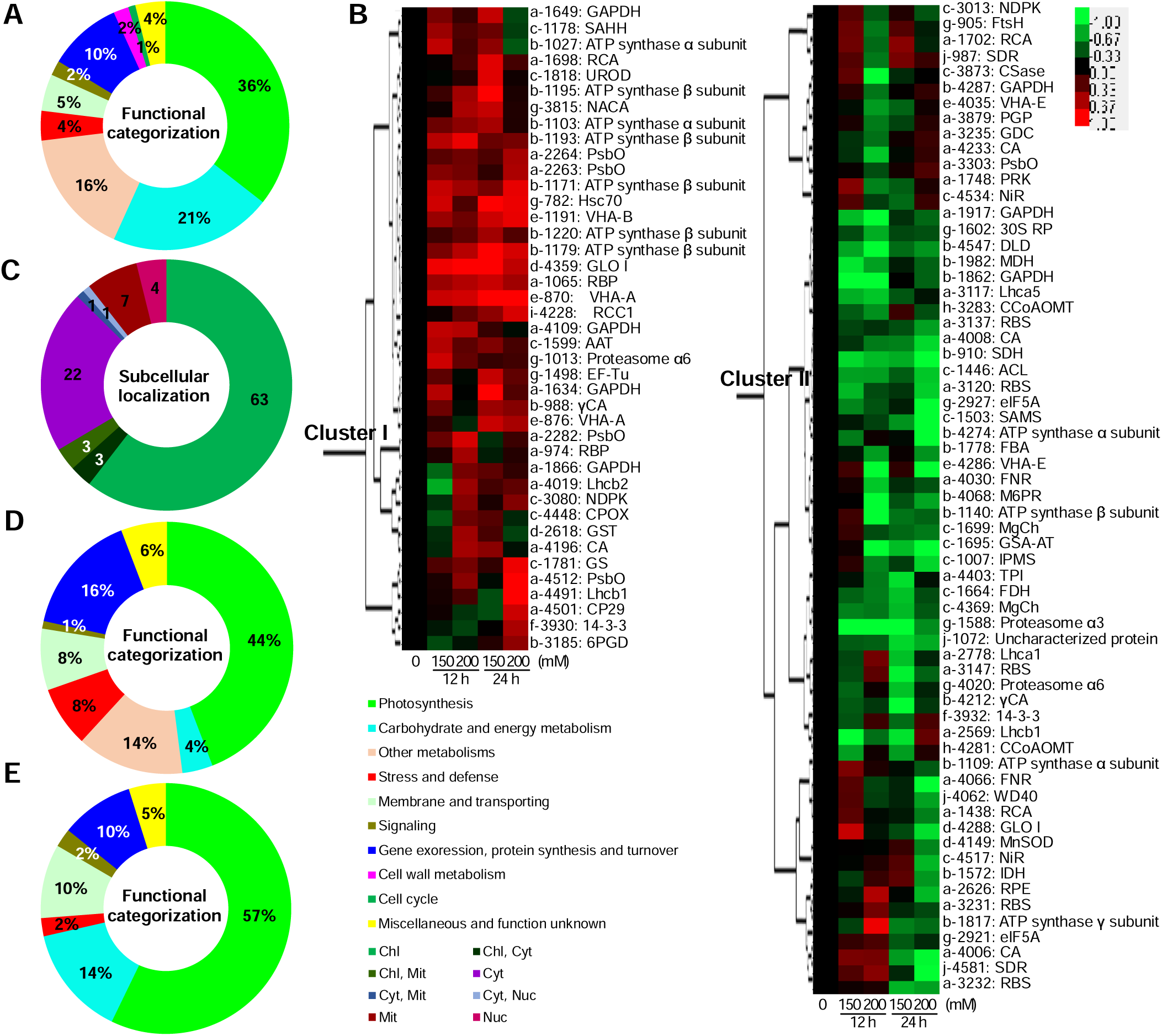
Functional categorization, hierarchical clustering analysis, and subcellular location prediction of the Na_2_CO_3_-responsive proteins. **A.** Functional category of Na_2_CO_3_-responsive leaf proteins. The percentages of proteins in different functional categories are shown in the pie. **B.** Heat map of Na_2_CO_3_-responsive protein species from leaf proteome. Two main clusters (I–II) are shown in the figure, functional categories indicated by lower-case letters (**a**, photosynthesis; **b**, carbohydrate and energy metabolism; **c**, other metabolisms; **d**, stress and defense; **e**, membrane and transporting; **f**, signaling; **g**, protein synthesis and turnover; **h**, cell wall metabolism; **i**, cell cycle; **j**, miscellaneous and function unknown), spot numbers and protein name abbreviations are listed on the right side (detailed information on protein names and abbreviations can be found in Table S1). **C.** Predicted localization of proteins from leaf proteome by internet tools and literatures. The numbers of protein species with different locations are shown in the pie. Chl, chloroplast; Cyt, cytoplasm; Mit, mitochondria; Nuc, nucleus. **D-E.** A total of 102 Na_2_CO_3_-responsive protein species and 84 Na_2_CO_3_-responsive phosphoprotein species in chloroplasts were classified into eight and seven functional categories, respectively. The percentages of proteins in different functional categories are shown in the pie.

Furthermore, we identified 121 Na_2_CO_3_-responsive proteins in chloroplasts (Table S5). Among them, there were 102 chloroplast-localized proteins belonging to eight functional categories (Figure 5D; Tables S6 and S7). Of these, 49 were photosynthetic proteins accounting for 48% of the total. This included five chlorophyll a/b binding proteins, 14 PSII-related proteins, seven PSI-related proteins, nine photosynthetic electron transfer chain proteins, four subunits of ATP synthase and ten Calvin cycle enzymes. Most of them were obviously altered at 12 HAT200 and 24 HAT (Table S6). Besides, nine photosynthetic electron transfer chain proteins and four subunits of chloroplast ATP synthase were changed (Table S6). This indicates that although Na_2_CO_3_ inhibited the light harvesting, the PSII and PSI were not changed much at 12 HAT, but were enhanced at 24 HAT. However, ATP synthesis decreased at 12 HAT200, and then recovered to normal or enhanced at 24 HAT. Changes of the ten Calvin cycle-related proteins imply that carbon assimilation was inhibited at 24 HAT (Table S6). In addition, among the five ROS scavenging enzymes, thioredoxin peroxidase, 2-Cys peroxiredoxin BAS1, and GR increased at 12 HAT and decreased at 24 HAT, while APX and GST decreased at 12 HAT and not changed at 24 HAT (Table S6).

### Phosphoproteomics revealed novel Na_2_CO_3_-responsive phosphorylation sites

We identified 63 Na_2_CO_3_-responsive phosphoproteins in leaves. Of these, 39 proteins showed increased phosphorylation levels and 21 had decreased phosphorylation levels. These proteins were classified into seven functional categories (Figure 5E, Table S8). Thirty-four phosphoproteins were predicted to be chloroplast-located, and involved in light harvesting, PSII, Calvin cycle and ATP synthesis (Table S8).

In chloroplasts from the Na_2_CO_3_-treated leaves, 161 unique phosphopeptides were identified, and 137 were quantified by dimethyl labeling (**Figure 6**A and Table S9). Among them, 50 proteins were found to be Na_2_CO_3_-responsive with 57 phosphorylation sites, including 33 increased and 15 decreased (Figure 6B and **Table 1**). The increased phosphoproteins include seven light harvesting proteins, six PSII proteins, five PSI proteins, three electron transfer chain proteins, two Calvin cycle-related proteins, a Na^+^/H^+^ antiporter, a villin-2, and two thylakoid organization related proteins. The decreased include five ATP synthase subunits and sucrose-phosphate synthase. In addition, two signaling related proteins and six proteins involved in gene expression and protein turnover increased in phosphorylation at 24 HAT (Table 1).

**Table 1.**
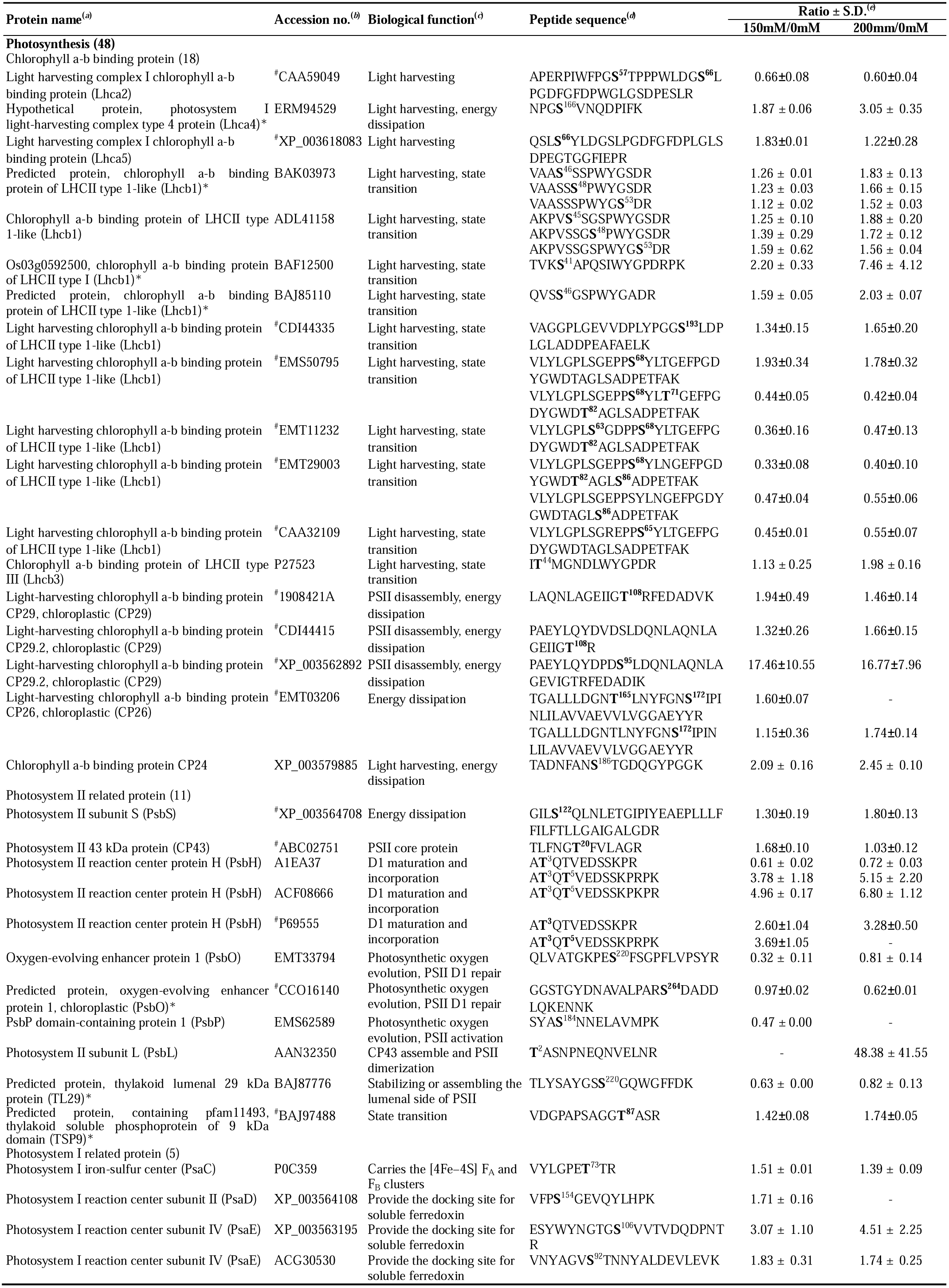
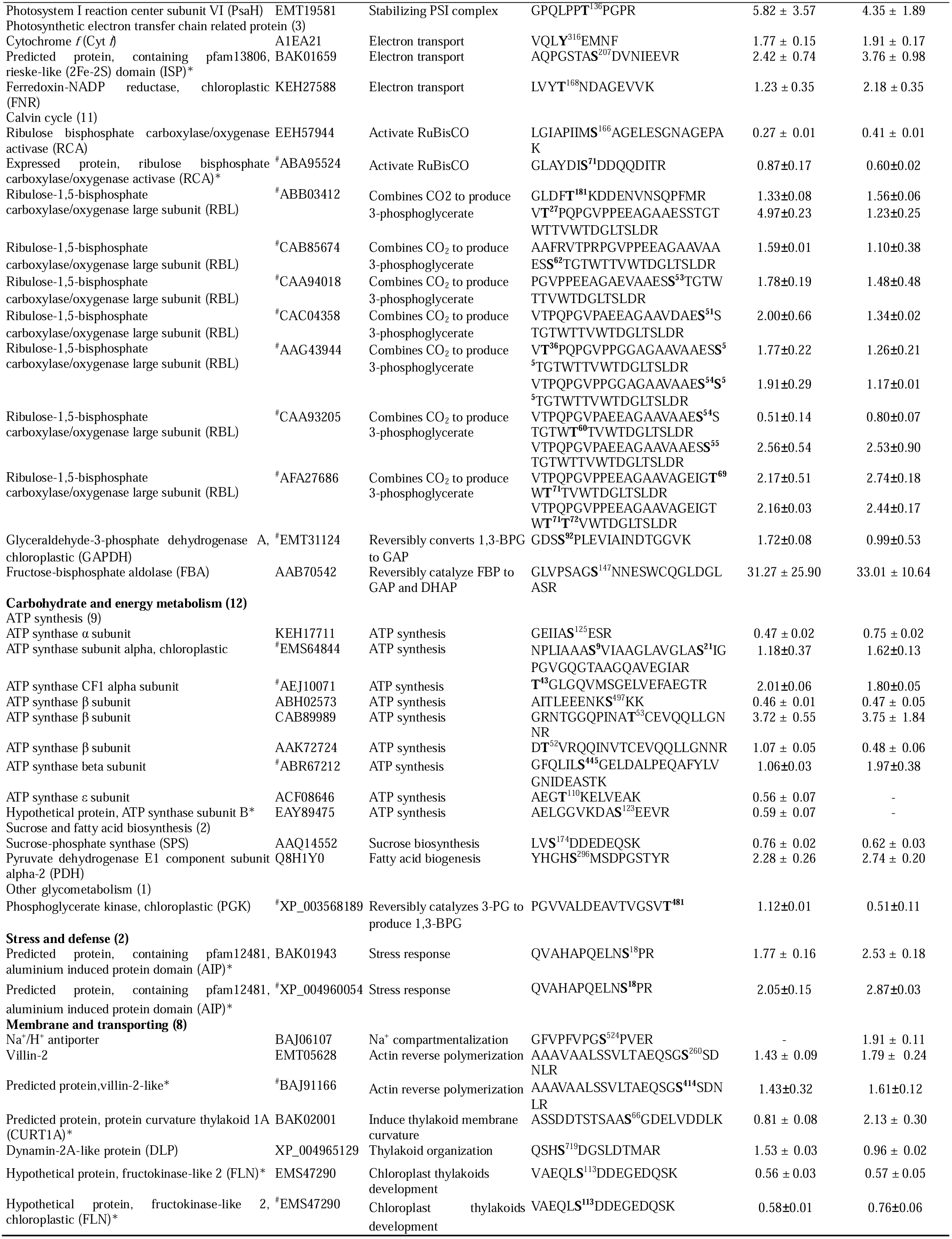
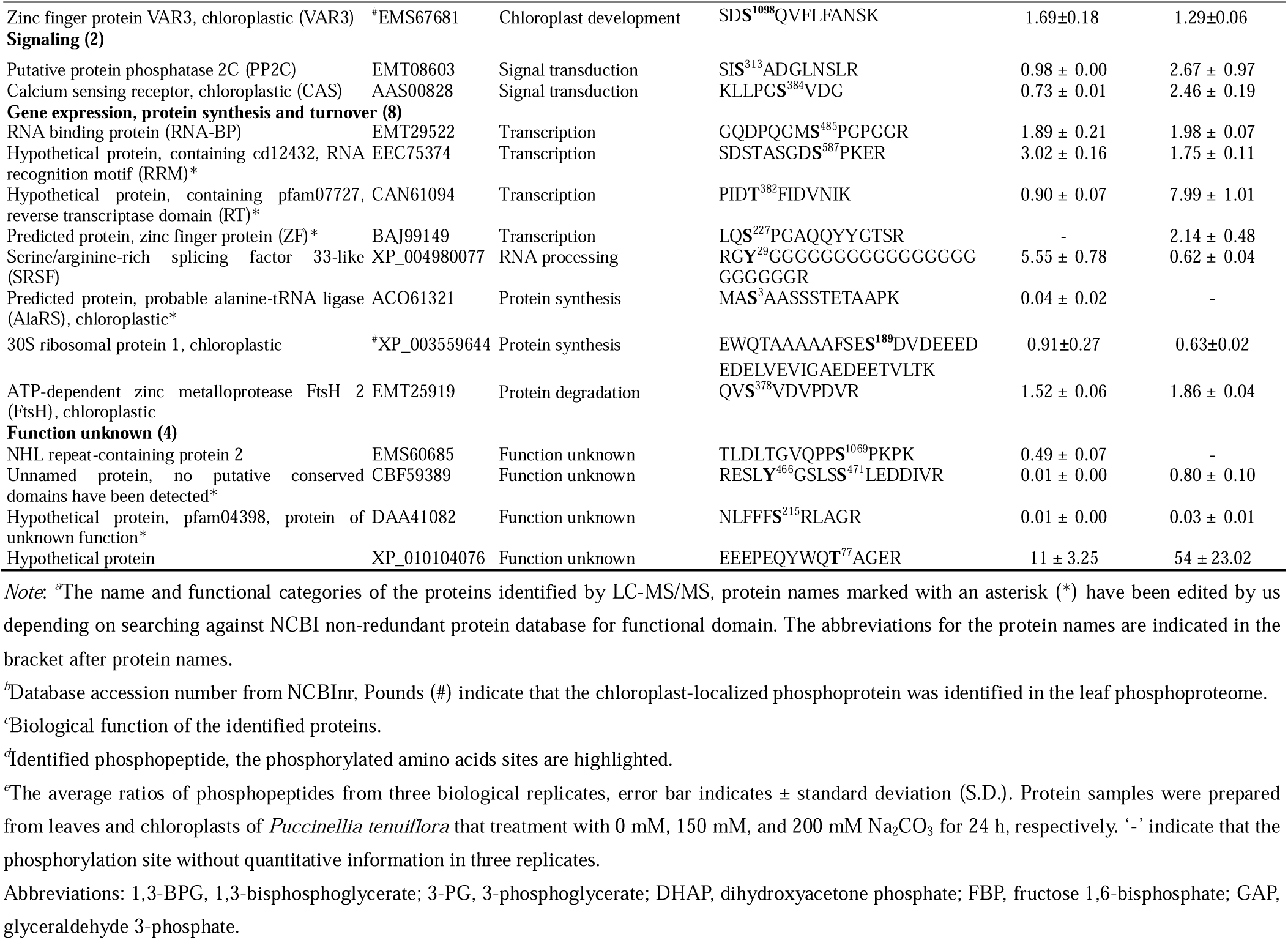
Na_2_CO_3_-responsive phosphoproteins in chloroplasts from alkaligrass leaves

**Figure 6.**
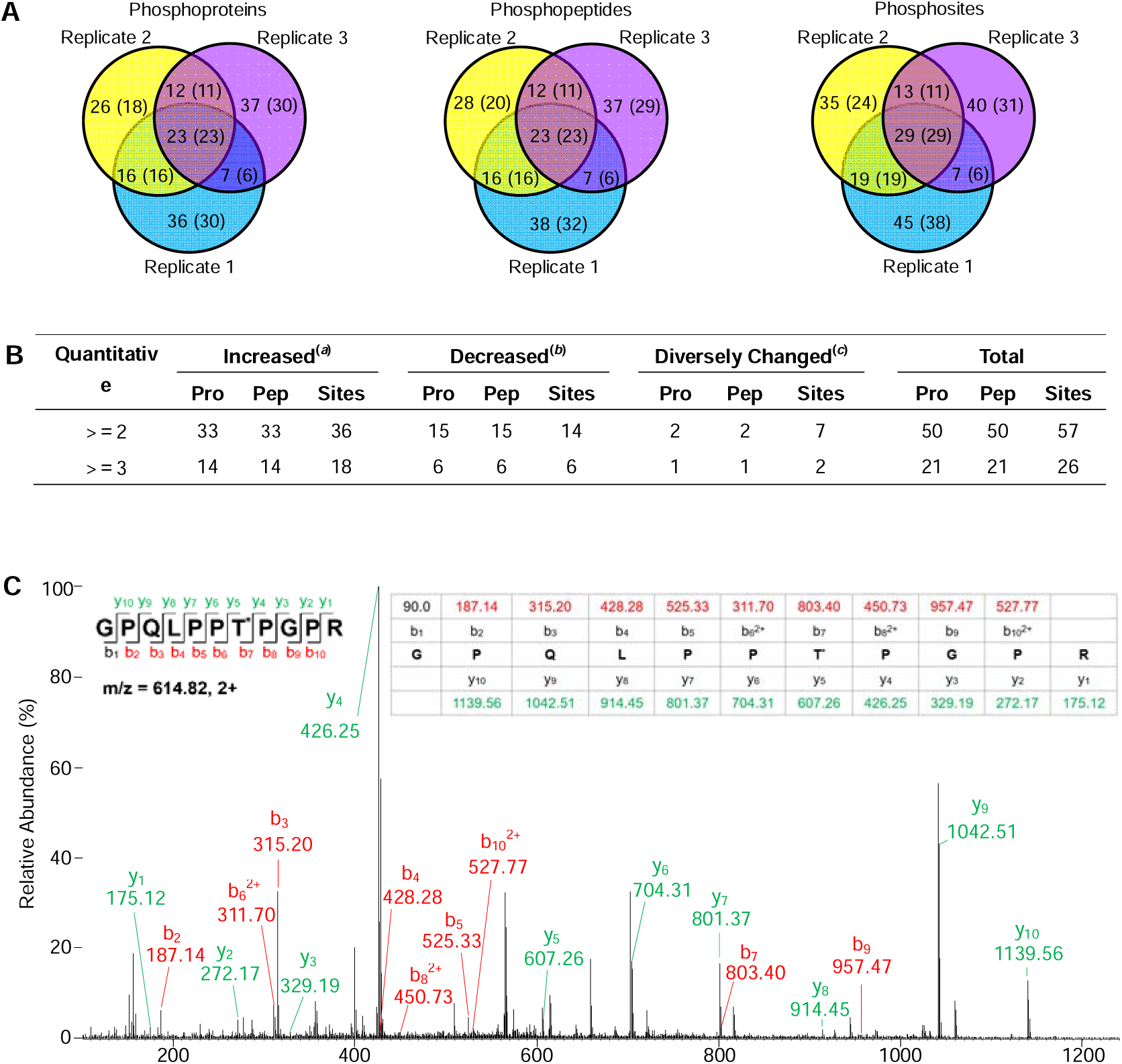
Summary of alkaligrass chloroplast phosphoproteome. **A.** Venn diagrams depicting overlap in phosphoproteins, phosphopeptides, and phosphorylation sites identified in three biological replicates. Numbers in parentheses indicate the quantified phosphoproteins, phosphopeptides, and phosphorylation sites, respectively. **B.** Na_2_CO_3_-responsive phosphoproteins, phosphopeptides, and phosphorylation sites in at least two replicates; ^*a,b*^ Phosphorylation level increased/decreased in certain treatment condition; ^*c*^ Phosphorylation level increased in one treatment but decreased in the other, or the changed peptide with two phosphorylation sites; Pro, protein; Pep, peptide. **C.** Example of a representative MS/MS spectrum of phosphopeptides identified in the chloroplast phosphoproteome (fragmentation spectrum shown of m/z 614.82, +2, dimethyllabeled, Accession No. EMT19581); Asterisk (*) represents the phosphorylation site.

In summary, we identified a total of 84 Na_2_CO_3_-responsive, chloroplast-localized phosphoproteins in leaf and chloroplast phosphoproteomes (Figures 6C and S4; Table 1). We identified 56 novel phosphorylation sites in the Na_2_CO_3_-responsive chloroplastic phosphoproteins, which may be crucial for regulating photosynthesis, membrane and transport, signaling, stress response, and protein synthesis and turnover (Table S10).

### Three-dimensional (3D) structure modeling of phosphoproteins

We built thirteen homology-based 3D models of chloroplast-localized phosphoproteins using the SWISS-MODEL program (Figure S5A, B, C, D, F, G, I, J, K, L, M and N). We also accepted two experimentally solved 3D structures as homology models by the significant amino acid sequence similarity and conserved phosphorylation sites with our phosphoproteins (Figure S5E, H). The 3D models showed the numbers of helices and beta sheets, and the phosphorylation sites of each protein (Figure S5 and Table S11).

### Twenty-eight homologous genes of Na_2_CO_3_-responsive phosphoproteins exhibited diverse expression patterns

In order to evaluate the gene expression patterns of the Na_2_CO_3_-responsive phosphoproteins, 28 homologous genes were analyzed through quantitative real-time (qRT-PCR) analysis with ubiquitin as an internal control (Figure S6, Table S12). Ten down-regulated genes were involved in light harvesting, PSII and PSI assembling, photosynthetic electron transfer, ATP synthesis, and thylakoid organization (Figure S6). Besides, three genes (*i.e. photosystem I reaction center subunit II (PsaD), serine/arginine-rich splicing factor 33-like*, and *Na*^*+*^*/H*^*+*^ *antiporter*) maintained stable levels under the Na_2_CO_3_. Interestingly, 15 up-regulated genes were involved in photosynthesis, Na^+^/H^+^ transport, calcium sensing, gene expression and protein turnover.

### Immunodetection of seven representative Na_2_CO_3_-rsponsive proteins

To further evaluate the protein abundances of representative photosynthetic proteins under Na_2_CO_3_ treatment, Western blotting was conducted using available antibodies. The abundances of PSII subunits (photosystem II 22 kDa protein (PsbS), photosystem II D1 protein (D1) and oxygen evolving enhancer protein (PsbO)) and PSI subunit of PsaD increased at 24 HAT200 (**Figure 7**). RuBisCO large subunit (RBL) decreased at 24 HAT (Figure 7). Photosynthetic electron transfer chain related cytochrome *f* (Cyt *f*) and Calvin cycle related phosphoglycerate kinase (PGK) maintained stable protein abundances at 24 HAT. Calvin cycle related sedoheptulose-1,7-bisphosphatase (SBPase) was used as the loading control (Figure 7).

**Figure 7.**
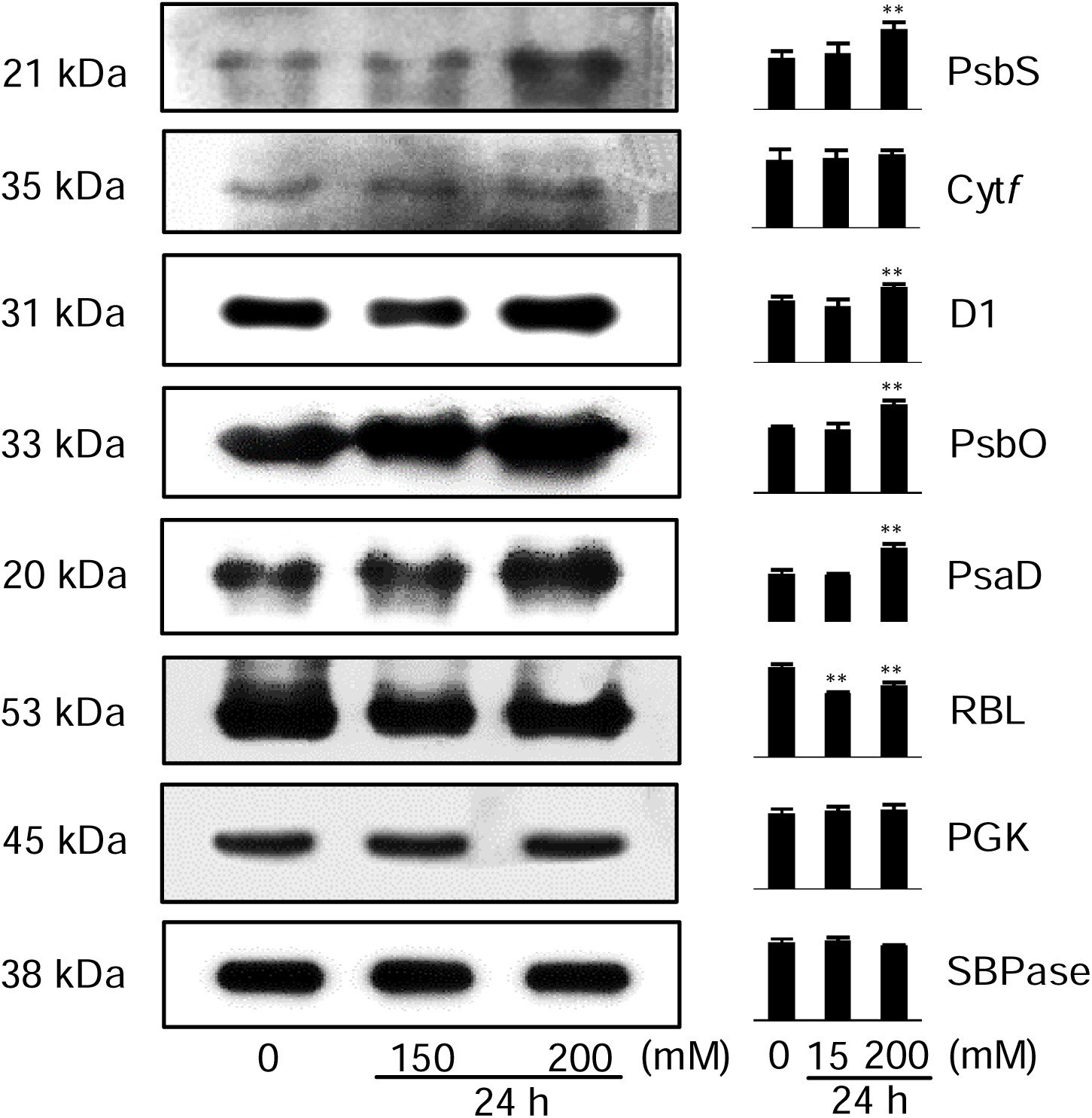
Western blot image of eight alkali-responsive chloroplast proteins. Eight chloroplast proteins from plants under different treatment conditions (0, 150, and 200 mM Na_2_CO_3_ for 24 h) were loaded with equal amounts. They were photosystem II 22 kDa protein (PsbS), cytochrome *f* (Cyt *f*), photosystem II D1 protein (D1), oxygen evolving enhancer protein (PsbO), photosystem I reaction center subunit II (PsaD), RuBisCO large subunit (RBL), phosphoglycerate kinase (PGK), and sedoheptulose-1,7-bisphosphatase (SBPase). These proteins were separated by 15% SDS-PAGE and analyzed by immunoblotting, and SBPase was used to control for equal loading. The relative abundances of these proteins were shown on the right. Results are the average of three independent experiments.

### Over-expression of PtFBA enhanced cell alkali tolerance

In our proteomics results, chloroplast-localized FBA increased significantly at phosphorylation level under Na_2_CO_3_ treatment. Therefore, FBA was selected as a representative Na_2_CO_3_ responsive protein for functional analysis. The full length cDNA of *PtFBA* was ligated into *PpsbAII* expression vector, and then transformed to wild-type (WT) cells of a model cyanobacterium *Synechocystis* 6803, generating an *OX-PtFBA* strain (**Figure 8**A). As expected, PCR analysis confirmed a complete segregation of the FBA over-expression (*OX-PtFBA*) strain (Figure 8B). Transcript analysis of *PtFBA* gene demonstrated the presence of gene product in the *OX-PtFBA* cells (Figure 8C). Western blotting analysis using a generic antibody against FBA also demonstrated that the FBA significantly increased in the *OX-PtFBA* strain when compared with WT (Figure 8D). The growth of the *OX-PtFBA* cells, as deduced from cell density and Chl *a* content, was much higher than the WT strain under the treatment of 0.4 M Na_2_CO_3_ for 4 days, although their growth was similar under normal conditions (Figure 8E, F). Thus, we conclude that overexpression of *PtFBA* resulted in enhanced Na_2_CO_3_ tolerance of *Synechocystis* 6803.

**Figure 8.**
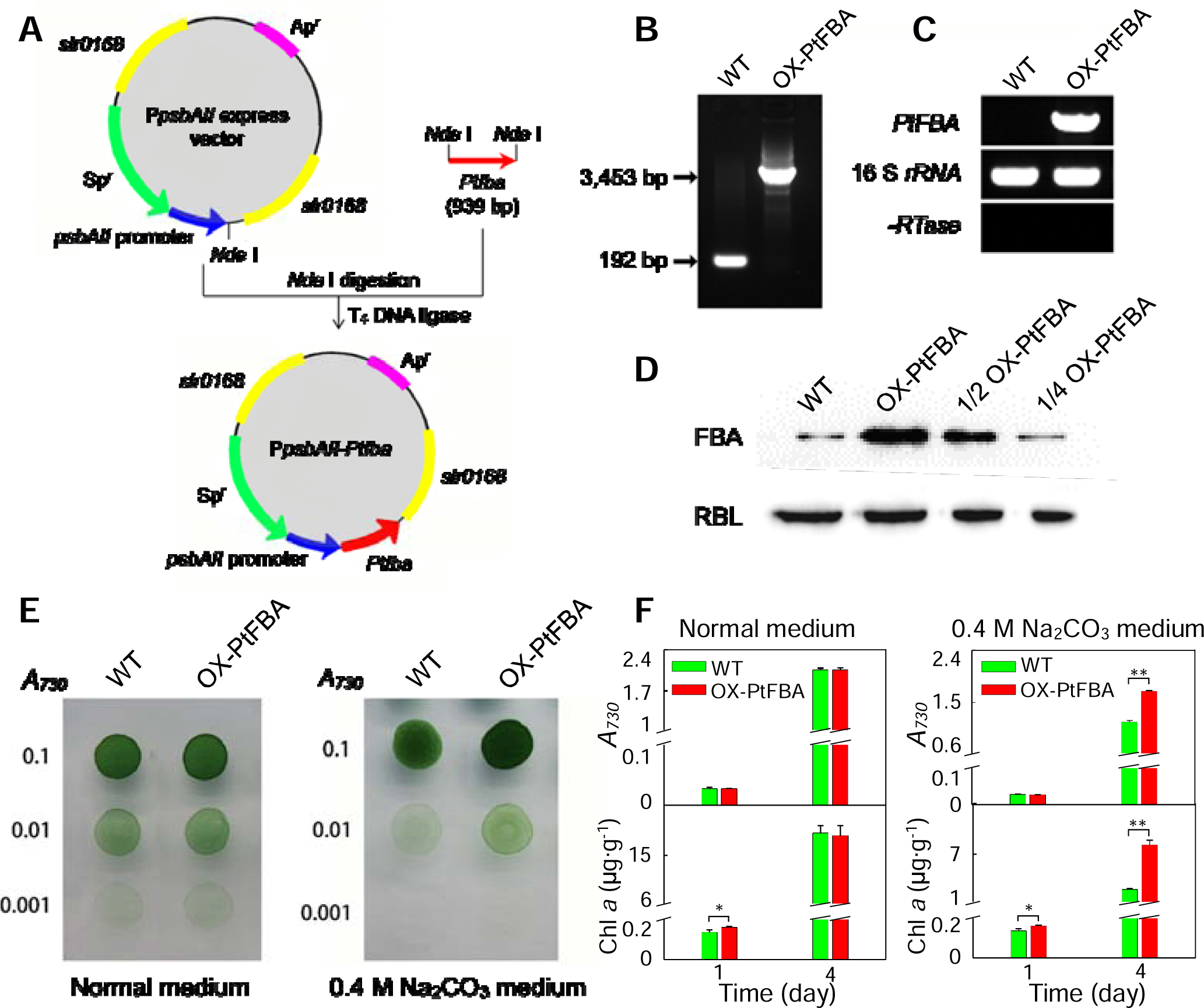
Transgenic analysis of chloroplast-localized *PtFBA* gene in a model cyanobacterium *Synechocystis* sp. strain PCC 6803. **A.** Construction of an over-expression vector P*psbAII-PtFBA*, to generate the *PtFBA* over-expression (OX-PtFBA) strain in *S*. PCC 6803. **B.** PCR segregation analysis of the *OX-PtFBA* cells. **C.** The transcript levels of *PtFBA* in the wild-type (WT) and *OX-PtFBA* strains. The transcript level of *16 S ribosomal RNA* (*rRNA*) in each sample is shown as a control. The absence of contamination of DNA was confirmed by PCR without reverse transcriptase reaction. **D.** Immunodetection of FBA levels in WT and *OX-PtFBA* strains. Protein blotting was performed with antibody against FBA. Lanes were loaded with 15 μg total proteins and RBL was used as a loading control. **E.** Growth of WT and *OX-PtFBA* strains under 0.4 M Na_2_CO_3_. Three μL of cell suspensions with densities corresponding to A_730_ nm values of 0.1 (upper rows), 0.01 (middle rows), and 0.001 (lower rows) were spotted on agar plates with normal medium and 0.4 M Na_2_CO_3_ medium. **F.** A_730_ nm values and chl *a* contents in WT and *OX-PtFBA* strains after cultivation in the medium with 0 M (controls) and 0.4 M Na_2_CO_3_ for 1 and 4 days, respectively. Mean ± S.D., n= 3.

## Discussion

### Diverse photoprotection mechanisms to counteract Na_2_CO_3_-induced photoinhibition

Photosynthesis modulation is critical for plant stress response. PSII supercomplex was very sensitive to environmental changes [28,29]. In Na_2_CO_3_-treated alkaligrass, PSII (*e.g.*, OEC and the reaction center proteins) was oxidatively damaged, resulting in the decreases of photochemical efficiency and electron transport [25,30]. Our results indicate that diverse photoprotection mechanisms were employed in alkaligrass to counteract alkali-induced photoinhibition. First, the accumulation of PsbS, chlorophyll a/b binding protein (CP) 24, and CP29, as well as induction of *CP24* gene at 24 HAT may contribute to PsbS-protonation-dependent conformation conversion of PSII antenna system, suggesting that PsbS-dependent thermal dissipation was enhanced to minimize the potential for photo-oxidative damage under the Na_2_CO_3_ treatment (Figure S8A) [31]. Consistently, CP24 and CP29 also displayed high abundances in salt-sensitive plants (Arabidopsis, oilseed rape (*Brassica napus*), and potato (*Solanum tuberosum*)) and salt-tolerant plants (Indian mustard (*Brassica juncea*), mangrove (*Kandelia candel*), wild tomato (*Solanum chilense*) and Sugar beet) under salt stresses [17,32-37]. Second, the phosphorylation at Ser186 of CP24, Thr165 and Ser172 of CP26, as well as Ser95 and Thr108 of CP29 were enhanced in alkaligrass at 24 HAT (Table 1), while CP24 became dephosphorylated in NaCl-treated *B. distachyon* [21]. The reversible phosphorylation of CP24 was supposed to regulate the alternative mode of phosphorylation-independent thermal dissipation and phosphorylation-dependent energy spillover in lycophytes [38]. Third, the state transition between PSII and PSI was regulated by protein phosphorylation in the Na_2_CO_3_-treated alkaligrass (Figure S8B). A serine/threonine-protein kinase (STN7) and protein phosphatases modulate the reversible phosphorylation of LHCII (*i.e.* Lhcb1 and Lhcb2) and thylakoid soluble phosphoprotein of 9 kDa (TSP9) [23]. The phosphorylation of Lhcb1 was reported to be salinity-increased in *B. distachyon* [21]. Besides, TSP9 interacts with LHCII and the peripheries of PSII and PSI, facilitating dissociation of LHCII from PSII for regulating photosynthetic state transitions [39]. Thus, the Na_2_CO_3_-enhanced gene expression of *Lhcb1*, abundances of Lhcb1, Lhcb2, and STN7, as well as phosphorylation of Lhcb1 and TSP9 may facilitate the state transition in alkaligrass (Figure S8B). Similarly, Lhcb1 and Lhcb2 increased in abundances by salt in salt-tolerant plants (*e.g.,K. candel*, moss (*Physcomitrella patens*)), *B. juncea*, soybean (*Glycine max*), and sugar beet [17,34,35,40,41]. Furthermore, Lhcb3 phosphorylation also increased in alkaligrass (Table 1). The rate of state transition was induced in Arabidopsis *Lhcb3* knockout mutant [42]. This implies that Lhcb3 is probably involved in the state transition, but the underlying regulatory mechanism needs to be further investigated (Figure S8B).

Cyclic electron transport (CET) is critical for protecting photosynthetic apparatus and additional ATP supply [43]. Several decreased electron transport-related proteins indicate that electron transport was slowed down in alkaligrass at 12 HAT (Figure S8C), and this may alleviate the damage of plastoquinone over-reduction. However, at 24 HAT, when the photosynthetic capacity and electron transport rate were inhibited, CET was induced due to the Na_2_CO_3_-increased gene expression of *Cyt f*, accumulated protein levels and enhanced phosphorylation of ferredoxin-NADP(+) reductase (FNR), Cyt *f* and cytochrome *b*_6_*f* complex iron-sulfur subunit (Figure S8C). Among them, FNR phosphorylation may modify its thylakoid membrane localization to regulate the electron transport [44]. Additionally, several CET-related proteins (*e.g.* Cyt *f*, cytochrome *b*_6_*f* complex iron-sulfur subunit, FNR, NAD(P)H-quinone oxidoreductase subunit M (NdhM), and NdhJ) were accumulated in the NaCl-stressed halophytes [14,45]. Therefore, the Na_2_CO_3_-stressed alkaligrass employed NDH-dependent CET to alleviate photo-oxidative damage and provide extra ATP.

Photorespiration is critical for GSH synthesis, nitrogen and carbon assimilation, and feedback regulation of photosynthetic activity to cope with alkali stress (Figure S8D) [4,25,46]. In addition, the decrease of photosynthesis was not resulted from stomatal limitation, but from inhibition of Calvin cycle because most Calvin cycle enzymes decreased significantly (Figure S8D). This is consistent with a previous report in halophytes and salt-tolerant cultivars [4,47].

### Enhancement of PSII repair machinery to minimize photodamage

PSII repair machinery was efficiently and dynamically employed to minimize photodamage in alkaligrass (Figure S8E). Na_2_CO_3_-induced STN7 and STN8 can promote the phosphorylation of PSII core proteins and CP29. The protein phosphorylation loosens the attractive forces among the subunits of PSII-LHCII supercomplex, enabling migration of the damaged PSII to stroma thylakoids for subsequent detachment of damaged D1 from the core complex during the repairing process [48]. Subsequently, the thylakoid lumen 18.3 kDa protein can function as an acid phosphatase to dephosphorylate the damaged D1 protein, and this process is facilitated by the Na_2_CO_3_-accumulated PsbO with GTPase activity. The dephosphorylated D1 is recognized and degraded by the Na_2_CO_3_-induced ATP-dependent zinc metalloprotease FtsH 2 (FtsH) (Figure S8E). Besides, Na_2_CO_3_-induced ToxA-binding protein 1 may contribute to the D1 degradation process through its positive regulatory role in the stabile accumulation of FtsH protease in chloroplast stroma [49]. Simultaneously, nascent copies of D1 protein are synthesized and processed rapidly (Figure S8E). Under Na_2_CO_3_ treatment, the D1 maturation and co-translational insertion into PSII complexes were prompted by the induced *PsbH* gene, accumulation of PsbH and low PSII accumulation 1 protein, as well as the induced phosphorylation of Thr3 or Thr5 in the PsbH [50,51].

OEC (PsbO, PsbP and PsbQ) is peripherally bound to PSII at the luminal side of the thylakoid membrane, which can stabilize the binding of inorganic cofactors, maintain the active Mn-cluster, and enhance oxygen-evolution in PSII [52]. The expression of *PsbO* and *PsbP* decreased in leaves at 24 HAT, and the abundance of OEC was affected in alkaligrass and mangrove [45]. Interestingly, phosphorylation of PsbO and PsbP was Na_2_CO_3_-decreased in alkaligrass (Figure S8C), and PsbP and PsbQ have been reported to be phosphorylated in thylakoid lumen of Arabidopsis [53]. This indicates that the Na_2_CO_3_-regulated OECs change will facilitate the PSII assembly and oxygen evolution. Besides, photosystem II subunit L (PsbL) and TL29 also participate in the assembly of the PSII complex [54,55]. The phosphorylation of PsbL was induced and the phosphorylation of TL29 was inhibited in response to Na_2_CO_3_ (Figure S8E). Although the phosphorylation of PsbL was also reported in Arabidopsis [23], their regulatory mechanisms are not known.

### Reverse phosphorylation of Na_2_CO_3_-responsive proteins regulated the activities of PSI and ATP synthase

In contrast to PSII, PSI drew little attention due to difficulties in accurately measuring its activity [56]. The abundance changes of several PSI proteins (*e.g.* Lhca1 and Lhca5) and Na_2_CO_3_-enhanced phosphorylation of Lhca2 and Lhca4 in alkaligrass allow the regulation of light absorption through the antenna modulation to prevent PSI damage (Figure S8C) [25]. The STN7-regulated phosphorylation of Lhca4 was also induced in Arabidopsis when the plastoquinone overly reduced [57]. Therefore, the enhancement of Lhca4 phosphorylation would be favorable for trapping and dissipation of excitations, working as a photoprotective mechanism of PSI [58].

Among the PSI core proteins, PsaA, PsaB, and PsaC [59] were decreased in alkaligrass (Figure S8C), barley (*Hordeum vulgare*) [60], and other halophytes [33,61]. The phosphorylation of PsaC was enhanced in alkaligrass at 24 HAT (Figure S8C), which has been reported in Arabidopsis, green algae (*Chlamydomonas reinhardtii*), and spikemoss (*Selaginella moellendorffii*) [23]. The decreased abundances of PsaA, PsaB, and PsaC imply that saline-alkali stress inhibited the energy transfer of PSI. In addition, PsaD and PsaE provide docking sites for ferredoxin, and PsaF is important for interaction with the lumenal electron donor plastocyanin, while PsaG and PsaH participate in stabilizing PSI complex [62]. In this study, these proteins were significantly accumulated, and the phosphorylation of PsaD, PsaE, and PsaH were also enhanced at 24 HAT of Na_2_CO_3_ (Figure S8C), although only *PsaH* gene transcription was obviously induced at 24 HAT. Their phosphorylation events have been reported in Arabidopsis, green algae, and *Synechocystis* 6803 [23], and the phosphorylation of PsaD was supposed to regulate the electron transfer from PSI to the electron acceptors in Arabidopsis chloroplast stroma [63]. Therefore, we suggested that the salinity-/alkali-increased abundances or phosphorylation of these proteins facilitate stabilization of the PSI complex (thus protecting it from photodamage), and activate the CET around PSI [62].

The gene expression and protein abundance of different subunits of ATP synthase were altered by Na_2_CO_3_ stress in alkaligrass (Figure S6) and other plant species [4]. The activity of ATP synthase is salinity-responsive, being regulated by the reversible protein phosphorylation [4,64]. Several phosphorylation sites in ATP synthase subunits (α, β, δ, and ε) of spinach (*Spinacia oleracea*) chloroplasts have been reported [64]. In this study, Na_2_CO_3_ inhibited the phosphorylation of α (Ser125), β (Ser497, and Thr52), and ε subunit (Thr110), but enhanced the phosphorylation of α (Ser9, Ser21, and Thr43) and β subunits (Thr53 and Ser445) in alkaligrass at 24 HAT (Figure S8C). This implies that the stability and rotation of F1 head of ATP synthase are modulated for the dynamic regulation of its activity to cope with alkali stress.

### Different ROS homeostasis pathways employed in chloroplasts and other subcellular locations to cope with Na_2_CO_3_ stress

Na_2_CO_3_ treatment disrupted the electron transport in chloroplasts, as well as tricarboxylic acid cycle and respiration chain in mitochondria (Figures 2 and S8F), leading to the increases of H_2_O_2_ and O_2_^-^ in leaves [65]. A previous proteomic study reported that many ROS-scavenging enzymes were altered in salinity-stressed leaves [4]. Our results indicated that parts of AsA-GSH cycle (*i.e.* APX and GPX pathways) were induced in leaves, but most ROS scavenging pathways in chloroplasts were inhibited, except for the APX pathway and SOD pathway (Figure 4). This implies that different pathways are employed in chloroplasts and other subcellular locations in leaves to cope with the short-term Na_2_CO_3_ stress (12 h). While under long-term NaCl or Na_2_CO_3_ stress (7 d), the pathways of SOD, POD, and CAT were all induced in leaves of alkaligrass [24, 25].

Various non-enzymatic antioxidants are important for ROS scavenging [66]. In this study, although the balance of AsA and DHA in chloroplasts was perturbed, the contents of AsA and DHA in leaves were stable. Additionally, the ratio of GSH/GSSG was stable in chloroplasts, and increased in leaves at 24 HAT (Figure 4). Furthermore, Na_2_CO_3_ also increased glyoxalase I, but decreased chloroplast-localized cysteine synthase. Both enzymes are involved in GSH/GSSG balance (Figure S8F). All these imply that the AsA-GSH cycle is inhibited in chloroplasts, but enhanced in other organelles and cytoplasm of leaf cells to cope with the alkali stress. In addition, a chloroplast-localized activator of bc1 complex kinase increased at 24 HAT, which would facilitate tocopherol cyclase phosphorylation to stabilize it at plastoglobules for vitamin E synthesis [67]. The thylakoid membrane-localized vitamin E is a lipid antioxidant. This result is consistent with our previous finding of NaCl-increased vitamin E content and tocopherol cyclase abundance in leaves of alkaligrass [25].

Additionally, the atlas of Na_2_CO_3_-responsive proteins indicated that modulation of Chl synthesis (Figure S8G), chloroplast movement and stability (Figure S8H) were critical for alkali adaptation, and ABA-dependent alkali-responsive pathways were employed to regulate both nuclear and chloroplastic gene expression and protein processes for osmoprotectant synthesis and signaling pathways in alkaligrass (Figure S8I and J).

## Conclusion

Although NaCl-responsive mechanisms have been well-studied in various halophytes using proteomics approaches [4], the Na_2_CO_3_-responsive proteins and corresponding regulatory mechanisms in halophytes were rarely explored. This study is the first detailed investigation of Na_2_CO_3_-responsive proteins in chloroplasts using proteomics and phosphoproteomics approaches, which revealed several crucial Na_2_CO_3_-responsive pathways in halophyte chloroplasts (Figure S8). Our study showed that maintenance of energy balance between PSII and PSI, efficiency of PSII damage repair, cyclic electron transport, dynamic thylakoid membrane architecture, as well as osmotic and ROS homeostasis were essential for photosynthetic modulation in response to Na_2_CO_3_. Both the nuclear- and chloroplast-encoded proteins were critical for the Na_2_CO_3_-responsive chloroplast function. More importantly, the newly-identified protein phosphorylation sites suggest that the reversible protein phosphorylation is important for regulating multiple signaling and metabolic pathways in chloroplasts to cope with the Na_2_CO_3_ stress. Some of these Na_2_CO_3_-responsive proteins and phosphoproteins are potential saline-alkali stress biomarkers for further functional characterization and biotechnological application.

## Materials and methods

### Plant material treatment and biomass analysis

Seeds of alkaligrass (*Puccinellia tenuiflora* (Turcz.) scribn. et Merr.) were sowed on vermiculite and grown in Hoagland solution in pots under fluorescent light (220 μmol m^−2^ s^−1^, 12 h day and 12 h night) at 25 °C day and 20 °C night, and 75% humidity for 50 days. Seedlings were treated with 0 mM, 150 mM, and 200 mM Na_2_CO_3_ (pH 11) for 12 h and 24 h, respectively (Figure S7). After treatment, leaves were harvested, either used immediately or stored at −80 °C for experiments. **S**hoot length and leaf fresh weight of seedlings were immediately measured. Dry weight, relative water content, as well as ion contents of K^+^, Na^+^, Ca^2+^ and Mg^2+^ were determined according to the method of Zhao et al. [26].

### Membrane integrity, osmolytes, and ABA analysis

The malondialdehyde content and relative electrolyte leakage were determined using previous methods [68,69]. Free proline and total soluble sugar contents were quantified with a spectrometer at 520 nm and 630 nm, respectively [26]. The content of endogenous ABA was measured by an indirect ELISA method [70].

### Photosynthesis and chloroplast ultrastructure analysis

Photosynthesis and Chl fluorescence parameters were measured using previous methods [25,69]. Net photosynthetic rate, stomatal conductance, and transpiration rate were determined at 10:00 a.m. using a portable photosynthesis system LI-COR 6400 (LI-COR Inc., Lincoln, Nebraska, USA). Chl fluorescence parameters were recorded using a pulse-amplitude-modulated (PAM) Chl fluorometer (Dua-PAM-100) (Heintz Walz, Effeltrich, Germany) and an emitter-detector-cuvette assembly with a unit 101ED (ED-101US). For the rapid fluorescence induction kinetics analysis, the OJIP were measured at room temperature (25 °C) with a portable fluorometer PAM-2500 (Walz, Effeltrich, Germany). The fluorescence measurement and calculation were performed according to the protocol of Strasser et al. [71]. Chl contents were determined according to the method of Wang et al. [69]. The ultrastructure of chloroplasts were observed under a JEOL-2000 transmission electron microscope (JEOL, Tokyo, Japan) according to Suo et al. [72].

### Measurements of ROS and antioxidant contents, and antioxidant enzyme activities

The O_2_^-^ generation rate and H_2_O_2_ content were measured according to Zhao et al. [26]. Reduced AsA, total AsA, GSSG, and total GSH contents were determined according to methods of Law et al. [73]. The activities of SOD, POD, CAT, APX, GR, and GST were measured as previously described [25,26]. The activities of MDHAR, DHAR, and GPX were assayed according to Zhao et al. [26].

### Chloroplast isolation, protein extraction and purity assessment

Intact chloroplasts were isolated according to Ni et al. [74], and the chloroplast protein for iTRAQ and phosphoproteomics analysis was extracted according to Wang et al. [75]. The purity of chloroplast protein was assessed by Western blot analysis with antibodies against marker proteins for different subcellular compartments according to Dai et al. [76]. Primary antibodies against marker proteins and protein loading amounts were listed in Table S13.

### Proteomic analysis chloroplasts and leaves

The protein samples were extracted, fractioned, lyophilized, and resuspended for MS/MS analysis according to Zhao et al. [26]. The peptides of control (0 mM Na_2_CO_3_), 12 HAT150, 12 HAT200, 24 HAT150, and 24 HAT200 were labeled with iTRAQ reagents 116, 117, 118, 119, and 121 (AB Sciex Inc., Foster City, CA, USA), respectively. Three biological replicates were performed. The LC-MS/MS analysis was performed by Triple TOF™ 5600 LC-MS/MS (AB Sciex Inc., Concord, Canada) according to Zhao et al. [26]. The MS/MS data were submitted to database searching using the Paragon algorithm of ProteinPilot (version 4.0, AB Sciex Inc.). The databases used were the Uniprot Liliopsida database (320 685 entries) and the National Center for Biotechnology Information (NCBI) non-redundant database (7,262,093 sequences). The proteomics data was available in the Proteomics Identifications (PRIDE) database [77] under accession number PXD005491.

Total protein samples from leaves were prepared and analyzed using two-dimensional gel electrophoresis according to the method of Suo et al. [72] and Dai et al. [76]. The MS and MS/MS spectra were acquired on a MALDI-TOF/TOF mass spectrometer (AB Sciex Inc.) [69]. The MS/MS spectra were searched against the NCBI non-redundant green plant database (http://www.ncbi.nlm.nih.gov/) (3,082,218 sequences) using the search engine Mascot (version 2.3.0, Matrix Science, London, UK) (http://www.matrixscience.com) according to Meng et al. [78]. The proteomics data was deposited in the PRIDE database [77] under accession number PXD005455.

### Phosphoproteomic analysis of proteins from chloroplasts and leaves

After digestion, the chloroplast peptide samples of control, 24 HAT150, and 24 HAT200 were labeled with stable isotope dimethyl labeling in light, intermediate, and heavy, respectively, according to the method of Boersema et al. [79]. In addition, the peptides from each leaf protein sample were labeled with iTRAQ reagents (113 and 116 for control, 114 and 117 for 24 HAT150, as well as 115 and 119 for 24 HAT200) according to the manufacturer’s instructions, respectively (AB Sciex Inc.). The phosphopeptides were enriched using TiO_2_ micro-column [80]. LC-MS/MS analysis was performed using a nanoAcquity ultraperformance LC (Waters, Milford, MA, USA) coupled with an Orbitrap Fusion Tribrid mass spectrometer (Thermo Fisher Scientific, Watham, MA, USA) [81].

For database searching, raw data files from chloroplast phosphoproteome were processed using Mascot server (version 2.3.0, Matrix Science) and searched against the NCBI green plant database (7,262,093 sequences) using an in-house Mascot Daemon (version 2.4, Matrix Science) [81]. For phosphopeptide relative quantification, Mascot Distiller (version 2.5.1.0, Matrix Science) was used. Precursor ion protocol was used for peptide quantification and the ratios were calculated using the peak areas of extracted ion chromatograms based on the trapezium integration method [82]. For leaf phosphoproteins, MS/MS data and peak lists were extracted using ProteinPilot (version 4.0, AB Sciex), searched against the NCBI green database (7,262,093 sequences) at a 95% confidence interval (unused ProtScore > 1.3). After database searching, reliable quantification of individual phosphopeptide was achieved by the mean ± S.D. of triplicate experiments, the peptides with more than 1.5-fold changes in at least two replicates were considered to be changed at phosphorylation level. The chloroplast and leaf phosphoproteomics data have been deposited to PRIDE [77] under the accession numbers of PXD005472 and PXD005471, respectively.

### Protein classification, hierarchical clustering, subcellular location prediction, and 3D structure analysis

Functional domains of proteins were analyzed using BLAST programs (http://www.ncbi.nlm.nih.gov/BLAST/), and then were clustered by cluster 3.0 (http://bonsai.hgc.jp/~mdehoon/software/cluster/software.htm). The prediction of protein subcellular location was determined according to Suo et al. [72]. The SWISS-MODEL comparative protein modeling server (http://swissmodel.expasy.org/) was employed to generate 3D structural models of phosphoproteins [83].

### qRT-PCR analysis of homologous gene expression

Total RNA was isolated from leaves using the pBiozol plant total RNA extraction reagent (BioFlux, Hangzhou, China). A first-strand cDNA was obtained from 1 μg of total RNA using a PrimeScript^®^ RT reagent kit (Takara Bio, Inc., Otsu, Japan). The sequences of candidate genes were obtained from the local alkaligrass EST database using a BLASTn program. qRT-PCR amplification was performed using the specific primer pairs (Table S12) on a 7500 real time PCR system (Applied Biosystems Inc., USA). The amplification process was performed according the method of Suo et al. [72].

### Western blot analysis

Western blotting was conducted according to Dai et al. [76]. The primary antibodies were raised in rabbits against the Arabidopsis PsbS, Cyt *f*, D1, PsbO, PsaD, RBL, PGK, and SBPase. Signals were detected with ECL Plus™ reagent (GE Healthcare) according to the manufacturer’s instruction. Relative abundances were analyzed using the Image Master 2D Platinum Software (version 5.0, GE Healthcare). For the immunodetection of the FBA level in the WT and *OX-PtFBA Synechocystis* 6803, antibodies against FBA and RBL was used, 15 μg total protein aliquots of WT and *OX-PtFBA* (including the indicated serial dilutions) were loaded, and RBL was used as a loading control.

### Overexpression of *PtFBA* in *Synechocystis* 6803

Full length cDNA of *PtFBA* was amplified by PCR using appropriate primers (Table S14). The P*psbAII* expression vector was used to generate *OX-PtFBA* strain. A fragment containing the *PtFBA* gene was amplified by PCR (Table S14) and then inserted into Nde I sites of P*psbAII* to form the *PpsbAII-PtFBA* expression vector construct, which was used to transform the WT *Synechocystis* 6803 using a natural transfer method [84]. The transformants were spread on BG-11 agar plates containing10 μg·ml^−1^ of spectinomycin, then incubated in 2% (v/v) CO_2_ in air and illumination at 40 μmol photons m^−2^·s^−1^. The *OX-PtFBA* cells in the transformants was segregated to homogeneity (by successive streak purification) as determined by PCR amplification (Table S14), reverse transcription (RT-PCR) analysis (Table S14), and immunoblotting [85]. Cell growth and Chl *a* content analysis were conducted according to Gao et al. [86].

## Statistical analysis

All the results are presented as means ± standard deviation of at least three replicates. The physiological and proteomics data were analyzed by Student’s *t* test. A *p*-value < 0.05 was considered statistically significant.

## Authors’ contribution

SD and ZQ conceived of the study, and participated in its design and coordination and helped to draft the manuscript. JS and HZ carried out physiological experiments. HZ, QZ, BS, JY, JL, and LG participated in proteomics analysis. JS, JM, YS, and XZ carried out the phosphoproteomics analysis. JC carried out the chloroplast ultrastructure analysis. JS and LP carried out the immunoassays. NZ, YZ, YL, and WM carried out the molecular genetic studies. SC, TW, YM, and SG helped to draft the manuscript. All authors read and approved the final manuscript.

## Acknowledgments

The project was supported by The Foundation of Shanghai Science and Technology Committee (No. 17391900600),The Program for Professor of Special Appointment (Eastern Scholar) from The Shanghai Bureau of Higher Education (2011 and 2017), and The National Natural Science Foundation of China (No. 31270310) to Shaojun Dai. Peter Scott from University of Florida is acknowledged for critical reading and editing of the manuscript.

## Supplementary material

**Figure S1 Morphological changes of alkaligrass seedlings grown under Na**_**2**_**CO**_**3**_ **treatment**

50-day old seedlings were treated with different concentrations, 0 mM, 150 mM and 200 mM of Na_2_CO_3_ (pH 11) for 0, 12 and 24 h, respectively. Bar = 3.23 cm.

**Figure S2 Integrity and purity determination of chloroplasts isolated from alkaligrass leaves**

**A.** Autofluorescence of the purified chloroplasts were visualized under a fluorescence microscope, Bar=10 μm. **B.** Western blot analysis of proteins from purified chloroplasts. Proteins from the leaves and purified chloroplasts were separated on 12.5% SDS-PAGE, and immunodetected with rabbit polyclonal antibodies against VHA (vacuolar-type H^+^-ATPase, vacuole marker), CAT (catalase, peroxisome marker), COXII (mitochondrial cytochrome oxidase subunit II, mitochondrial marker), Sar1 (secretion-associated and ras-related protein 1, endoplasmic reticulum marker), and FBPase (cytosolic fructose 1,6-biphosphatase, cytosolic marker), respectively.

**Figure S3 Coomassie brilliant blue-stained two-dimensional gel electrophoresis gels**

Protein was extracted from leaves of *Puccinellia tenuiflora* under Na_2_CO_3_ treatment conditions and separated on 24 cm IPG strips (pH 4-7 linear gradient) using isoelectric focusing in the first dimension, followed by 12.5% SDS-PAGE gels in the second dimension. A total of 104 Na_2_CO_3_-responsive proteins identified by mass spectrometry were marked with numbers on the representative gels. Detailed information can be found in Supplementary Table S2. **A.** 0 mM Na_2_CO_3_. **B.** 150 mM-12 h Na_2_CO_3_. **C.** 200 mM-12 h Na_2_CO_3_. **D.** 150 mM-24 h Na_2_CO_3_. **E.** 200 mM-24 h Na_2_CO_3_.

**Figure S4 A list of MS/MS spectra of phosphopeptides derived from alkaligrass leaf and chloroplast phosphoproteins**

**Figure S5 Homology based structural models of the chloroplast phosphoproteins from alkaligrass**

The ribbon images of **A-D, F, G**, and **I-N** are generated by SWISS-MODEL program based on the comparative protein structure modeling approach, and the 3D models based on their own amino acid sequences. While images of **E** and **H** are the experimentally resolved 3D structure that has a significant amino acid sequence similarity to our phosphoproteomic identified sequence by searching the SWISS-MODEL template library. By using the Swiss-PdbViewer (version 3.7), the phosphorylation sites are shown with balls in red (enhanced phosphorylation) and green (inhibited phosphorylation). Domains from the phosphoprotein are highlighted in different colors. **A.** Lhca2, light-harvesting chlorophyll a/b binding protein of LHCI type II (PDB ID: 4xk8). **B.** Lhca4, photosystem I light-harvesting complex type 4 protein (PDB ID: 4y28). **C.** Lhcb1, chlorophyll a/b binding protein of LHCII type 1 (PDB ID: 1rwt). **D.** PsbS, photosystem II subunit S (PDB ID: 4ri2). **E.** PsbO, oxygen-evolving enhancer protein 1 (PDB ID: 3jcu). **F.** TL29, thylakoid lumenal 29 kDa protein (PDB ID: 3rrw). **G.** PsaC, photosystem I subunit VII (PDB ID: 4y28). **H.** PsaD, photosystem I reaction center subunit II (PDB ID: 2wsc). **I.** PsaE, photosystem I reaction center subunit IV (I and II representing two protein species of PsaE) (PDB ID: 4y28). **J.** Cyt f, cytochrome f (PDB ID: 1q90). **K.** FNR, ferredoxin-NADP reductase (PDB ID: 1qg0). **L.** ATP synthase alpha subunit (PDB ID: 5cdf). **M.** ATP synthase beta subunit (PDB ID: 1fx0). **N.** PDH, pyruvate dehydrogenase E1 component subunit alpha (PDB ID: 1ni4). Detailed information can be found in Table S11.

**Figure S6 Cluster analysis of the homologous gene expression patterns of alkaligrass chloroplast phosphoproteins under 24h of Na**_**2**_**CO**_**3**_ **treatment**

Heat map of the 28 homologous genes from *Puccinellia tenuiflora* leaves under 24 h of Na_2_CO_3_ treatment (0, 150, and 200 mM), three main clusters (I–III) are shown in the figure. The scale bar indicates log_2_ transformed relative expression levels of genes. The up-regulated and down-regulated genes are presented in red and green, respectively. Homologous gene names are listed on the right side.

**Figure S7 Workflow of plant treatment and sampling**

Seeds of *Puccinellia tenuiflora* were sowed on vermiculite and grown in Hoagland solution in pots. Seedlings about 50-day-old were treated with 0, 150, and 200 mM Na_2_CO_3_ (pH 11) for 12 and 24 h, respectively. After treatment, leaves from both control and treatments were harvested for experiments.

**Figure S8 Schematic presentation of Na**_**2**_**CO**_**3**_ **responsive mechanisms in leaves from alkaligrass**

The Na_2_CO_3_ responsive proteins from both leaf and chloroplast proteomes were integrated into subcellular pathways. **A.** Thermal dissipation. **B.** State transition. **C.** Photosynthetic electron transfer. **D.** Photorespiration and Calvin cycle. **E.** PSII repair cycle. **F.** ROS scavenging. **G.** Chlorophyll biosynthesis. **H.** Chloroplast movement and thylakoid membrane stability. **I.** Ionic and osmotic homeostasis. **J.** Gene expression, protein turnover and transport. The scale bar indicates log (base 2) transformed protein abundance, enzyme activity, and substrate content ratios (compared with 0 mM Na_2_CO_3_ treatment). The ratio was ranging from −3.0 to 3.0 with 7 different colors from green to red. P in a red (increased in phosphorylation level), green (decreased in phosphorylation level), and blue (can be phosphorylated by other protein, but did not identified in our phosphoproteomic study) circle indicates phosphorylated protein. The solid line indicates single-step reaction, dashed line indicates multistep reaction, and the dotted line indicates movement of proteins or other substances. Abbreviations: 1,3-BPG, 1,3-bisphosphoglyceric acid; 13^1^-Hydroxy-Mg-Proto ME, 13^1^-hydroxy-magnesium-protoporphyrin IX 13-monomethyl ester; 3-PGA, 3-phosphoglyceric acid; 30S/50S, 30S/50S ribosomal protein; 40S/60S, eukaryotic small/large ribosomal subunit; ABA, abscisic acid; ABC1K, activator of bc1 complex kinases; ALA, 5-aminolaevulinic acid; AlaRS, alanine-tRNA ligase; APX, ascorbate peroxidase; AsA, ascorbic acid; CA, carbonic anhydrase; CAT, catalase; Coprogen III, corproporphyrinogen III; CP, chlorophyll a/b binding protein; CPOX, coproporphyrinogen-III oxidase; CSase, cysteine synthase; CURT1A, curvature thylakoid 1A protein; Cyt *f*, cytochrome *f*; D1/D2, photosystem II D1/D2 protein; DHA, dehydroascorbic acid; DHAP, dihydroxyacetone phosphate; DHAR, dehydroascorbate reductase; DLD, dihydrolipoyl dehydrogenase; DLP, dynamin-2A-like protein; DnaJ, chaperone protein DnaJ-like; E4P, erythrose 4-phosphate; EF-Tu, elongation factor Tu; eIF4B, eukaryotic initiation factor 4B; eIF5A, eukaryotic translation initiation factor 5A1; F6P, fructose 1,6-diphosphate; FBA, fructose-bisphosphate aldolase; FBP, fructose 1,6-bisphosphate; Fd, ferredoxin; FNR, ferredoxin-NADP(+) reductase; FtsH, ATP-dependent zinc metalloprotease FtsH 2; GAP, glyceraldehyde 3-phosphate; GAPDH, glyceraldehyde 3-phosphate dehydrogenase; GDC, glycine decarboxylase; GGDR, geranylgeranyl diphosphate reductase; GGPP, geranylgeranyl pyrophosphate; GLO I, glyoxalase I; GPX, glutathione peroxidase; GR, glutathione reductase; GS-, xenobiotic substrates; GSA, glutamate 1-semialdehyde aminotransferase; GSA-AT, glutamate-1-semialdehyde 2,1-aminomutase; GSH, glutathione; GSSG, oxidized glutathione; GST, glutathione S-transferase; Hsc70, heat shock cognate 70 kDa protein; Hsp70/90, 70/90 kDa heat shock-related protein; IDH, isocitrate dehydrogenase (NADP)-like; IPP, isopentenyl pyrophosphate; ISP, rieske-like iron-sulfur (2Fe-2S) protein; LHCII, light harvesting complex II; Lhca1/2/4/5, light harvesting chlorophyll a/b binding protein1/2/4/5; Lhcb1/2/3, light harvesting chlorophyll a/b binding protein 1/2/3; LIL3, light harvesting-like protein 3; LPA1, low PSII accumulation 1 protein; MDA, malondialdehyde; MDH, malate dehydrogenase; MDHA, monodehydroascorbate; MDHAR, monodehydroascorbate reductase; Mg-pro IX, Mg-proporphyrin IX; Mg-pro ME, Mg-protoporphyrin IX monomethyl ester; MgCh, magnesium-protoporyphyrin IX chelatase ChlI subunit; MgCy, magnesium-protoporphyrin IX monomethyl ester [oxidative] cyclase; MgMT, magnesium-protoporphyrin O-methyltransferase; NACA, nascent polypeptide associated complex subunit alpha-like protein; NDF1, NDH-dependent cyclic electron flow 1; NdhJ/M/O, NAD(P)H-quinone oxidoreductase subunit J/M/O; OAA, oxaloacetate; P-EAMeT, phosphoethanolamine N-methyltransferase; PAP, plastid-lipid-associated protein; PC, plastocyanin; PG, phosphoglycolate; PGP, phosphoglycolate phosphatase; Phytyl-PP, phytyl pyrophosphate; PPIase, peptidyl-prolyl cis-trans isomerase CYP20-2 domain; PPOX, protoporphyrinogen oxidase; PQ, plastoquinone; PQH_2_, plastohydroquinone; PRK, phosphoribulokinase; Prx, 2-Cys peroxiredoxin BAS1; PsaB, photosystem I P700 chlorophyll A apoprotein A2; PsaC, photosystem I subunit VII; PsaD, photosystem I reaction center subunit II; PsaE, photosystem I reaction center subunit IV; PsaF, photosystem I reaction center subunit III; PsaG, photosystem I subunit V; PsaH, photosystem I reaction center subunit VI; PSAT, phosphoserine aminotransferase; PsbH, photosystem II reaction center protein H; PsbL, Photosystem II subunit L; PsbO, oxygen-evolving enhancer protein 1; PsbP, photosystem II 23kDa oxygen evolving protein; PsbQ, oxygen evolving enhancer protein 3 domain; PsbS, photosystem II 22 kDa protein; R5P, ribose 5-phosphate; RaiA, ribosome-associated inhibitor A domain; RBL, RuBisCO large chain; RBP, RuBisCO large subunit-binding protein; RBS, RuBisCO small chain; RCA, RuBisCO activase; RCC1, regulator of chromosome condensation repeat; RNA-BP, RNA binding protein; RNP, 31 kDa ribonucleoprotein; ROS, reactive oxygen species; RPE, ribulose-phosphate 3-epimerase; RRM, RNA recognition motif; RT, reverse transcriptase domain; Ru5P, ribulose-5-phosphate; RuBP, ribulose-1,5-bisphosphate; S7P, sedoheptulose 7-phosphate; SAHH, S-adenosyl-L-homocysteine hydrolase; SBP, sedoheptulose 1,7-bisphosphate; SDH, succinate dehydrogenase flavoprotein; SecY, preprotein translocase subunit SecY; SHMT, serine hydroxymethyltransferase; SOD, manganese superoxide dismutase; SRSF, serine/arginine-rich splicing factor 33-like; STN7/8, serine/threonine-protein kinase STN7/8; TCA, tricarboxylic acid; Tic62, Tic 62 domain containing protein; TK, transketolase; TL15, thylakoid lumenal 15 kDa protein; TL29, thylakoid lumenal 29 kDa protein; TLP18.3, thylakoid lumen 18.3 kDa protein; TOC/TIC, translocon at the outer/inner envelope membrane of chloroplasts; ToxABP1, chloroplast-localized ToxA binding protein 1; TPI, triosephosphate isomerase; TPx, thioredoxin peroxidase; Trx, thioredoxin; TSP9, thylakoid soluble phosphoprotein; URO III, uroporphyrinogen III; UROD, uroporphyrinogen decarboxylase; VHA, vacuolar H^+^-ATPase; Xu5P, xylulose 5-phosphate; YCF54, YCF54 domain containing protein; ZF, zinc finger protein.

**Table S1 Detailed information of Na**_**2**_**CO**_**3**_**-responsive proteins in alkaligrass leaves identified by MALDI-TOF MS/MS**

**Table S2 Protein spots with multiple proteins identified in a single spot on two-dimensional gels of alkaligrass leaves**

**Table S3 Na**_**2**_**CO**_**3**_**-responsive proteins in alkaligrass leaves**

**Table S4 Subcellular localization prediction of the Na**_**2**_**CO**_**3**_**-responsive proteins in alkaligrass leaves**

**Table S5 Detailed information on Na**_**2**_**CO**_**3**_**-responsive proteins in alkaligrass chloroplasts revealed by iTRAQ-based proteomic analysis**

**Table S6 Na**_**2**_**CO**_**3**_**-responsive proteins in alkaligrass chloroplasts**

**Table S7 Subcellular localization prediction of the Na**_**2**_**CO**_**3**_**-responsive proteins in alkaligrass chloroplasts**

**Table S8 Na**_**2**_**CO**_**3**_**-responsive phosphoproteins in alkaligrass leaves**

**Table S9 Phosphoproteins identified in the chloroplast phosphoproteome of alkaligrass under Na**_**2**_**CO**_**3**_ **treatment**

**Table S10 A comparison of the phosphorylation sites/phosphoproteins identified in alkaligrass chloroplasts with phosphoproteins reported in plant species**

**Table S11 Summary of homology models of phosphoproteins in alkaligrass chloroplasts**

**Table S12 List of primer pairs used in quantitative real-time PCR**

**Table S13 Information on antibodies used for chloroplast purity assessment**

**Table S14 Primers used for overexpression of *PtFBA* in *Synechocystis* 6803**

## References

[1] Yang C, Chong J, Li C, Kim C, Shi D, Wang D. Osmotic adjustment and ion balance traits of an alkali resistant halophyte *Kochia sieversiana* during adaptation to salt and alkali conditions. Plant Soil 2007; 294: 263–76.

[2] Yang C, Xu H, Wang L, Liu J, Shi D, Wang D. Comparative effects of salt-stress and alkali-stress on the growth, photosynthesis, solute accumulation, and ion balance of barley plants. Photosynthetica 2009; 47: 79–86.

[3] Tuteja N. Mechanisms of high salinity tolerance in plants. Methods Enzymol 2007; 428: 419–38.

[4] Zhang H, Han B, Wang T, Chen S, Li H, Zhang Y, et al. Mechanisms of plant salt response: insights from proteomics. J Proteome Res 2012; 11: 49–67.

[5] Suo J, Zhao Q, David L, Chen S, Dai S. Salinity response in chloroplasts: insights from gene characterization. Int J Mol Sci 2017; 18: 1011.

[6] Silveira JAG, Carvalho FEL. Proteomics, photosynthesis and salt resistance in crops: an integrative view. J Proteomics 2016; 143: 24–35.

[7] Majeran W, Friso G, Ponnala L, Connolly B, Huang M, Reidel E, et al. Structural and metabolic transitions of C_4_ leaf development and differentiation defined by microscopy and quantitative proteomics in maize. Plant Cell 2010; 22: 3509–42.

[8] Olinares PD, Ponnala L, van Wijk KJ. Megadalton complexes in the chloroplast stroma of *Arabidopsis thaliana* characterized by size exclusion chromatography, mass spectrometry, and hierarchical clustering. Mol Cell Proteomics 2010; 9: 1594–615.

[9] Zhao Q, Chen S, Dai S. C4 photosynthetic machinery: insights from maize chloroplast proteomics. Front Plant Sci 2013; 4: 85.

[10] Zybailov B, Friso G, Kim J, Rudella A, Rodriguez VR, Asakura Y, et al. Large scale comparative proteomics of a chloroplast Clp protease mutant reveals folding stress, altered protein homeostasis, and feedback regulation of metabolism. Mol Cell Proteomics 2009; 8: 1789–810.

[11] Muneer S, Park YG, Manivannan A, Soundararajan P, Jeong BR. Physiological and proteomic analysis in chloroplasts of *Solanum lycopersicum* L. under silicon efficiency and salinity stress. Int J Mol Sci 2014; 15: 21803–24.

[12] Kamal AH, Cho K, Kim DE, Uozumi N, Chung KY, Lee SY, et al. Changes in physiology and protein abundance in salt-stressed wheat chloroplasts. Mol Biol Rep 2012; 39: 9059–74.

[13] Chang L, Guo A, Jin X, Yang Q, Wang D, Sun Y, et al. The beta subunit of glyceraldehyde 3-phosphate dehydrogenase is an important factor for maintaining photosynthesis and plant development under salt stress-Based on an integrative analysis of the structural, physiological and proteomic changes in chloroplasts in *Thellungiella halophila*. Plant Sci 2015; 236: 223–38.

[14] Fan P, Feng J, Jiang P, Chen X, Bao H, Nie L, et al. Coordination of carbon fixation and nitrogen metabolism in Salicornia europaea under salinity: comparative proteomic analysis on chloroplast proteins. Proteomics 2011; 11: 4346–67.

[15] Joaquin-Ramos A, Huerta-Ocampo JA, Barrera-Pacheco A, De Leon-Rodriguez A, Baginsky S, Barba de la Rosa AP. Comparative proteomic analysis of amaranth mesophyll and bundle sheath chloroplasts and their adaptation to salt stress. J Plant Physiol 2014; 171: 1423–35.

[16] Meng F, Luo Q, Wang Q, Zhang X, Qi Z, Xu F, et al. Physiological and proteomic responses to salt stress in chloroplasts of diploid and tetraploid black locust (*Robinia pseudoacacia* L.). Sci Rep 2016; 6: 23098.

[17] Wang L, Liang W, Xing J, Tan F, Chen Y, Huang L, et al. Dynamics of chloroplast proteome in salt-stressed mangrove *Kandelia candel* (L.) Druce. J Proteome Res 2013; 12: 5124–36.

[18] Zörb C, Herbst R, Forreiter C, Schubert S. Short-term effects of salt exposure on the maize chloroplast protein pattern. Proteomics 2009; 9: 4209–20.

[19] Chen Y, Hoehenwarter W. Changes in the phosphoproteome and metabolome link early signaling events to rearrangement of photosynthesis and central metabolism in salinity and oxidative stress response in Arabidopsis. Plant Physiol 2015; 169: 3021–33.

[20] Liu Z, Li Y, Cao H, Ren D. Comparative phospho-proteomics analysis of salt-responsive phosphoproteins regulated by the MKK9-MPK6 cascade in Arabidopsis. Plant Sci 2015; 241: 138–50.

[21] Lv DW, Subburaj S, Cao M, Yan X, Li X, Appels R, et al. Proteome and phosphoproteome characterization reveals new response and defense mechanisms of *Brachypodium distachyon* leaves under salt stress. Mol Cell Proteomics 2014; 13: 632–52.

[22] Yu B, Li J, Koh J, Dufresne C, Yang N, Qi S, et al. Quantitative proteomics and phosphoproteomics of sugar beet monosomic addition line M14 in response to salt stress. J Proteomics 2016; 143: 286–97.

[23] Grieco M, Jain A, Ebersberger I, Teige M. An evolutionary view on thylakoid protein phosphorylation uncovers novel phosphorylation hotspots with potential functional implications. J Exp Bot 2016; 67: 3883–96.

[24] Yu J, Chen S, Wang T, Sun G, Dai S. Comparative proteomic analysis of *Puccinellia tenuiflora* leaves under Na_2_CO_3_ stress. Int J Mol Sci 2013; 14: 1740–62.

[25] Yu J, Chen S, Zhao Q, Wang T, Yang C, Diaz C, et al. Physiological and proteomic analysis of salinity tolerance in *Puccinellia tenuiflora*. J Proteome Res 2011; 10: 3852–70.

[26] Zhao Q, Suo J, Chen S, Jin Y, Ma X, Yin Z, et al. Na_2_CO_3_-responsive mechanisms in halophyte *Puccinellia tenuiflora* roots revealed by physiological and proteomic analyses. Sci Rep 2016; 6: 32717.

[27] Zhang X, Wei L, Wang Z, Wang T. Physiological and molecular features of *Puccinellia tenuiflora* tolerating salt and alkaline-salt stress. J Integr Plant Biol 2013; 55: 262–76.

[28] Nishiyama Y, Murata N. Revised scheme for the mechanism of photoinhibition and its application to enhance the abiotic stress tolerance of the photosynthetic machinery. Appl Microbiol Biotechnol 2014; 98: 8777–96.

[29] Wei X, Su X, Cao P, Liu X, Chang W, Li M, et al. Structure of spinach photosystem II-LHCII supercomplex at 3.2 A resolution. Nature 2016; 534: 69–74.

[30] Wang Y, Sun G, Suo B, Chen G, Wang J, Yan Y. Effects of Na_2_CO_3_ and NaCl stresses on the antioxidant enzymes of chloroplasts and chlorophyll fluorescence parameters of leaves of *Puccinellia tenuiflora* (Turcz.) scribn.et Merr. Acta Physiol Plant 2008; 30: 143–50.

[31] Ruban AV. Nonphotochemical chlorophyll fluorescence quenching: mechanism and effectiveness in protecting plants from photodamage. Plant Physiol 2016; 170: 1903–16.

[32] Evers D, Legay S, Lamoureux D, Hausman JF, Hoffmann L, Renaut J. Towards a synthetic view of potato cold and salt stress response by transcriptomic and proteomic analyses. Plant Mol Biol 2012; 78: 503–14.

[33] Pang Q, Chen S, Dai S, Chen Y, Wang Y, Yan X. Comparative proteomics of salt tolerance in *Arabidopsis thaliana* and *Thellungiella halophila*. J Proteome Res 2010; 9: 2584–99.

[34] Yang L, Zhang Y, Zhu N, Koh J, Ma C, Pan Y, et al. Proteomic analysis of salt tolerance in sugar beet monosomic addition line M14. J Proteome Res 2013; 12: 4931–50.

[35] Yousuf PY, Ahmad A, Aref IM, Ozturk M, Hemant, Ganie AH, et al. Salt-stress-responsive chloroplast proteins in *Brassica juncea* genotypes with contrasting salt tolerance and their quantitative PCR analysis. Protoplasma 2016; 253: 1565–75.

[36] Zhou S, Sauvé RJ, Liu Z, Reddy S, Bhatti S, Hucko SD, et al. Identification of salt-induced changes in leaf and root proteomes of the wild tomato, *Solanum chilense*. J Am Soc Hortic Sci 2011; 136: 288–302.

[37] Jia H, Shao M, He Y, Guan R, Chu P, Jiang H. Proteome dynamics and physiological responses to short-term salt stress in *Brassica napus* leaves. PLoS One 2015; 10: e0144808.

[38] Ferroni L, Angeleri M, Pantaleoni L, Pagliano C, Longoni P, Marsano F, et al. Light-dependent reversible phosphorylation of the minor photosystem II antenna Lhcb6 (CP24) occurs in lycophytes. Plant J 2014; 77: 893–905.

[39] Fristedt R, Carlberg I, Zygadlo A, Piippo M, Nurmi M, Aro EM, et al. Intrinsically unstructured phosphoprotein TSP9 regulates light harvesting in *Arabidopsis thaliana*. Biochemistry 2009; 48: 499–509.

[40] Wang L, Liu X, Liang M, Tan F, Liang W, Chen Y, et al. Proteomic analysis of salt-responsive proteins in the leaves of mangrove *Kandelia candel* during short-term stress. PLoS One 2014; 9: e83141.

[41] Wang X, He Y. Proteomic analysis of the response to high-salinity stress in *Physcomitrella patens*. Planta 2008; 228: 167–77.

[42] Damkjaer JT, Kereiche S, Johnson MP, Kovacs L, Kiss AZ, Boekema EJ, et al. The photosystem II light-harvesting protein Lhcb3 affects the macrostructure of photosystem II and the rate of state transitions in Arabidopsis. Plant Cell 2009; 21: 3245–56.

[43] Rumeau D, Peltier G, Cournac L. Chlororespiration and cyclic electron flow around PSI during photosynthesis and plant stress response. Plant Cell Environ 2007; 30: 1041–51.

[44] Hodges M, Miginiac-Maslow M, Le Marechal P, Remy R. The ATP-dependent post translational modification of ferredoxin: NADP^+^ oxidoreductase. Biochim Biophys Acta 1990; 1052: 446–52.

[45] Wang L, Pan D, Li J, Tan F, Hoffmann-Benning S, Liang W, et al. Proteomic analysis of changes in the *Kandelia candel* chloroplast proteins reveals pathways associated with salt tolerance. Plant Sci 2015; 231: 159–72.

[46] Osmond CB, Grace SC. Perspectives on photoinhibition and photorespiration in the field: quintessential inefficiencies of the light and dark reactions of photosynthesis? J Exp Bot 1995; 46: 1351–62.

[47] Boex-Fontvieille E, Daventure M, Jossier M, Hodges M, Zivy M, Tcherkez G. Phosphorylation pattern of Rubisco activase in Arabidopsis leaves. Plant Biol (Stuttg) 2014; 16: 550–7.

[48] Gururani MA, Venkatesh J, Tran LS. Regulation of photosynthesis during abiotic stress-induced photoinhibition. Mol Plant 2015; 8: 1304–20.

[49] Manning VA, Chu AL, Scofield SR, Ciuffetti LM. Intracellular expression of a host-selective toxin, ToxA, in diverse plants phenocopies silencing of a ToxA-interacting protein, ToxABP1. New Phytol 2010; 187: 1034–47.

[50] Komenda J, Tichy M, Eichacker LA. The PsbH protein is associated with the inner antenna CP47 and facilitates D1 processing and incorporation into PSII in the cyanobacterium *Synechocystis* PCC 6803. Plant Cell Physiol 2005; 46: 1477–83.

[51] Peng L, Ma J, Chi W, Guo J, Zhu S, Lu Q, et al. Low PSII accumulation 1 is involved in efficient assembly of photosystem II in *Arabidopsis thaliana*. Plant Cell 2006; 18: 955–69.

[52] Ifuku K. The PsbP and PsbQ family proteins in the photosynthetic machinery of chloroplasts. Plant Physiol Biochem 2014; 81: 108–14.

[53] Reiland S, Messerli G, Baerenfaller K, Gerrits B, Endler A, Grossmann J, et al. Large-scale Arabidopsis phosphoproteome profiling reveals novel chloroplast kinase substrates and phosphorylation networks. Plant Physiol 2009; 150: 889–903.

[54] Granlund I, Storm P, Schubert M, Garcia-Cerdan JG, Funk C, Schroder WP. The TL29 protein is lumen located, associated with PSII and not an ascorbate peroxidase. Plant Cell Physiol 2009; 50: 1898–910.

[55] Suorsa M, Regel RE, Paakkarinen V, Battchikova N, Herrmann RG, Aro EM. Protein assembly of photosystem II and accumulation of subcomplexes in the absence of low molecular mass subunits PsbL and PsbJ. Eur J Biochem 2004; 271: 96–107.

[56] Sonoike K. Photoinhibition of photosystem I. Physiol Plant 2011; 142: 56–64.

[57] Ihnatowicz A, Pesaresi P, Lohrig K, Wolters D, Muller B, Leister D. Impaired photosystem I oxidation induces STN7-dependent phosphorylation of the light-harvesting complex I protein Lhca4 in *Arabidopsis thaliana*. Planta 2008; 227: 717–22.

[58] Passarini F, Wientjes E, van Amerongen H, Croce R. Photosystem I light-harvesting complex Lhca4 adopts multiple conformations: red forms and excited-state quenching are mutually exclusive. Biochim Biophys Acta 2010; 1797: 501–8.

[59] Jensen PE, Bassi R, Boekema EJ, Dekker JP, Jansson S, Leister D, et al. Structure, function and regulation of plant photosystem I. Biochim Biophys Acta 2007; 1767: 335–52.

[60] Rasoulnia A, Bihamta MR, Peyghambari SA, Alizadeh H, Rahnama A. Proteomic response of barley leaves to salinity. Mol Biol Rep 2011; 38: 5055–63.

[61] Cheng T, Chen J, Zhang J, Shi S, Zhou Y, Lu L, et al. Physiological and proteomic analyses of leaves from the halophyte *Tangut Nitraria* reveals diverse response pathways critical for high salinity tolerance. Front Plant Sci 2015; 6: 30.

[62] Yang H, Liu J, Wen X, Lu C. Molecular mechanism of photosystem I assembly in oxygenic organisms. Biochim Biophys Acta 2015; 1847: 838–48.

[63] Hansson M, Vener AV. Identification of three previously unknown in vivo protein phosphorylation sites in thylakoid membranes of *Arabidopsis thaliana*. Mol Cell Proteomics 2003; 2: 550–9.

[64] Schmidt C, Zhou M, Marriott H, Morgner N, Politis A, Robinson CV. Comparative cross-linking and mass spectrometry of an intact F-type ATPase suggest a role for phosphorylation. Nat Commun 2013; 4: 1985.

[65] Suzuki N, Koussevitzky S, Mittler R, Miller G. ROS and redox signalling in the response of plants to abiotic stress. Plant Cell Environ 2012; 35: 259–70.

[66] Miller G, Suzuki N, Ciftci-Yilmaz S, Mittler R. Reactive oxygen species homeostasis and signalling during drought and salinity stresses. Plant Cell Environ 2010; 33: 453–67.

[67] Martinis J, Glauser G, Valimareanu S, Kessler F. A chloroplast ABC1-like kinase regulates vitamin E metabolism in Arabidopsis. Plant Physiol 2013; 162: 652–62.

[68] Li H, Sun Q, Zhao S, Zhang W. Principles and techniques of plant physiological biochemical experiment. 3rd ed. Beijing: Higher education press; 2000.

[69] Wang X, Chen S, Zhang H, Shi L, Cao F, Guo L, et al. Desiccation tolerance mechanism in resurrection fern-ally *Selaginella tamariscina* revealed by physiological and proteomic analysis. J Proteome Res 2010; 9: 6561–77.

[70] Yang J, Zhang J, Wang Z, Zhu Q, Wang W. Hormonal changes in the grains of rice subjected to water stress during grain filling. Plant Physiol 2001; 127: 315–23.

[71] Strasserf RJ, Srivastava A, Govindjee. Polyphasic cholofphyll *a* fluorescence transient in plants and cyanobacteria. Photochem Photobiol 1995; 61: 32–42.

[72] Suo J, Zhao Q, Zhang Z, Chen S, Cao J, Liu G, et al. Cytological and proteomic analyses of *Osmunda cinnamomea* germinating spores reveal characteristics of fern spore germination and rhizoid tip-growth. Mol Cell Proteomics 2015; 14: 2510–34.

[73] Law MY, Charles SA, Halliwell B. Glutathione and ascorbic acid in spinach (*Spinacia oleracea*) chloroplasts. The effect of hydrogen peroxide and of paraquat. Biochem J 1983; 210: 899–903.

[74] Ni RJ, Shen Z, Yang CP, Wu YD, Bi YD, Wang BC. Identification of low abundance polyA-binding proteins in Arabidopsis chloroplast using polyA-affinity column. Mol Biol Rep 2010; 37: 637–41.

[75] Wang X, Li X, Deng X, Han H, Shi W, Li Y. A protein extraction method compatible with proteomic analysis for the euhalophyte *Salicornia europaea*. Electrophoresis 2007; 28: 3976–87.

[76] Dai S, Li L, Chen T, Chong K, Xue Y, Wang T. Proteomic analyses of *Oryza sativa* mature pollen reveal novel proteins associated with pollen germination and tube growth. Proteomics 2006; 6: 2504–29.

[77] Vizcaino JA, Csordas A, del-Toro N, Dianes JA, Griss J, Lavidas I, et al. 2016 update of the PRIDE database and its related tools. Nucleic Acids Res 2016; 44: D447–56.

[78] Meng X, Zhao Q, Jin Y, Yu J, Yin Z, Chen S, et al. Chilling-responsive mechanisms in halophyte *Puccinellia tenuiflora* seedlings revealed from proteomics analysis. J Proteomics 2016; 143: 365–81.

[79] Boersema PJ, Foong LY, Ding VM, Lemeer S, van Breukelen B, Philp R, et al. In-depth qualitative and quantitative profiling of tyrosine phosphorylation using a combination of phosphopeptide immunoaffinity purification and stable isotope dimethyl labeling. Mol Cell Proteomics 2010; 9: 84–99.

[80] Zhang X, Ye J, Jensen ON, Roepstorff P. Highly efficient phosphopeptide enrichment by calcium phosphate precipitation combined with subsequent IMAC enrichment. Mol Cell Proteomics 2007; 6: 2032–42.

[81] Ma J, Wang D, She J, Li J, Zhu JK, She YM. Endoplasmic reticulum-associated N-glycan degradation of cold-upregulated glycoproteins in response to chilling stress in Arabidopsis. New Phytol 2016; 212: 282–96.

[82] She YM, Rosu-Myles M, Walrond L, Cyr TD. Quantification of protein isoforms in mesenchymal stem cells by reductive dimethylation of lysines in intact proteins. Proteomics 2012; 12: 369–79.

[83] Wei S, Bian Y, Zhao Q, Chen S, Mao J, Song C et al. Salinity-induced palmella formation mechanism in halotolerant algae *Dunaliella salina* revealed by quantitative proteomics and phosphoproteomics. Front. Plant Sci., 2017; 8: 810.

[84] Long Z. Effects of different light treatments on the natural transformation of *Synechocystis* sp. strain PCC 6803. African Journal of Microbiology Research 2011; 5: 3603–10.

[85] Zhao J, Rong W, Gao F, Ogawa T, Ma W. Subunit Q is required to stabilize the large complex of NADPH dehydrogenase in *Synechocystis* sp. strain PCC 6803. Plant Physiol 2015; 168: 443–51.

[86] Gao F, Zhao J, Wang X, Qin S, Wei L, Ma W. NdhV Is a subunit of NADPH dehydrogenase essential for cyclic electron transport in *Synechocystis* sp. strain PCC 6803. Plant Physiol 2016; 170: 752–60.

