## Supplementary figures and images for "Na_2_CO_3_-Responsive Photosynthetic and ROS Scavenging Mechanisms in Chloroplasts of Alkaligrass Revealed by Phosphoproteomics"

### Supplemental Figure 1

0 mM

150 mM

200 mM

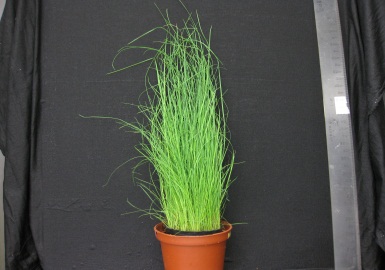

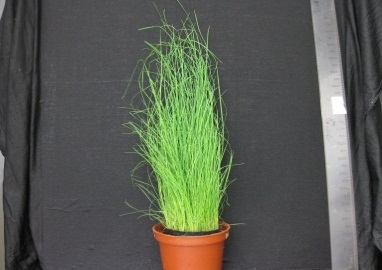

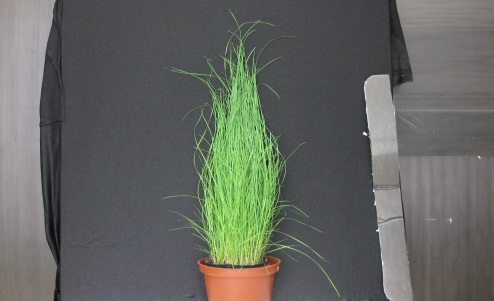

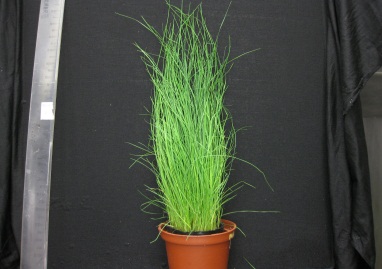

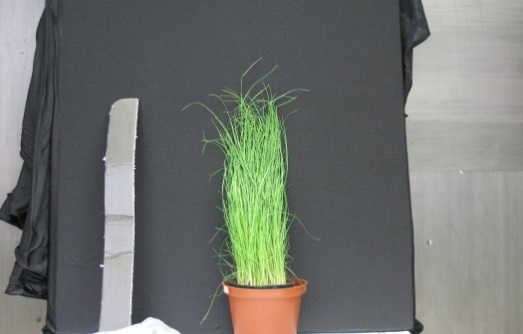

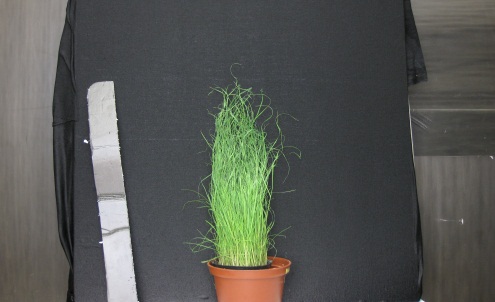

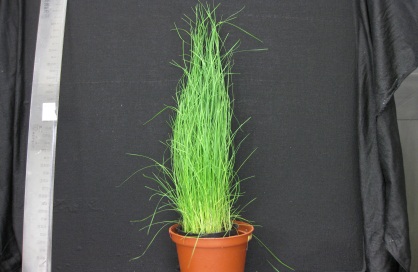

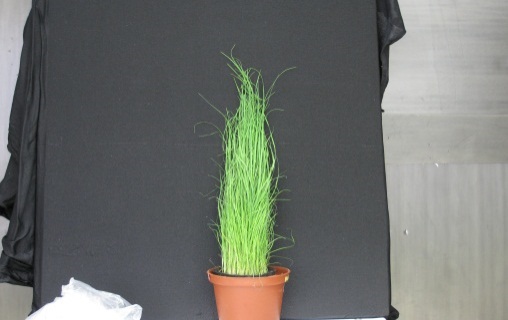

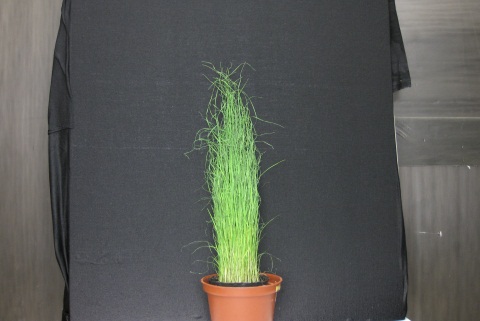

0 h	12 h	24 h

3.23 cm

### Supplemental Figure 2

**B**

Leaf

Chloroplast

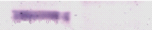

VHA

CAT

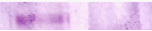

COXII

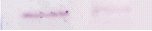

Sar1

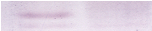

FBPase

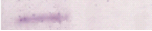

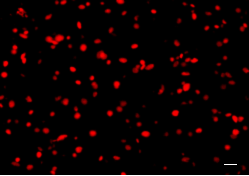

**A**

10 μm

### Supplemental Figure 3

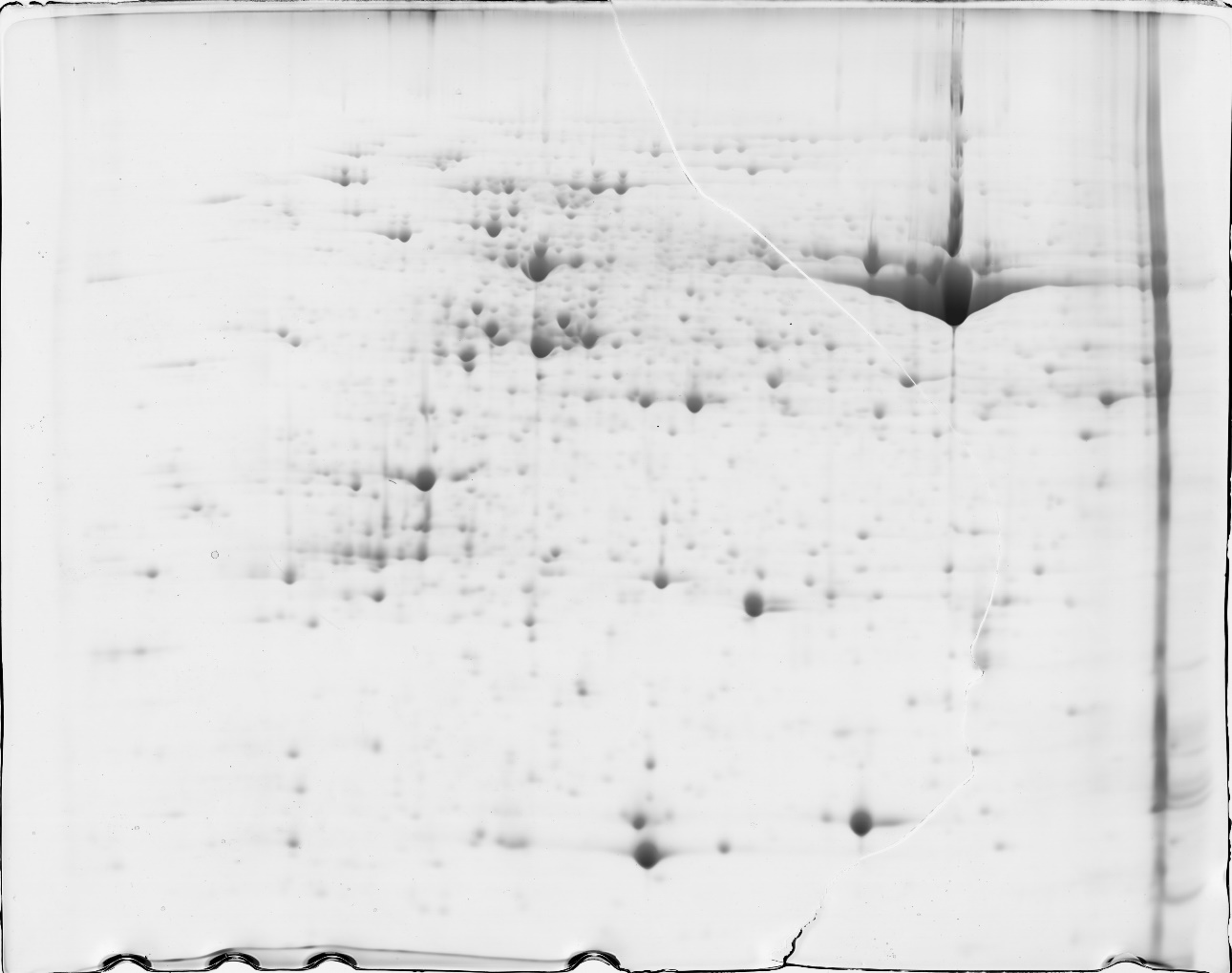

4006

4008

4196

1698

1456

1687

3120

3147

3232

1634

1649

1917

4109

1903

4403

1820

2626

3298

2778

2569

4019

4491

4501

4512

4030

4066

2605

2616

2618

4240

4002

4288

4359

4272

895

1027

1141

1020

1065

1171

1817

1140

1179

1193

1195

4547

1446

1606

910

1982

1862

3003

4212

4151

1695

1818

4115

1686

1599

3602

1503

4281

1178

1781

1929

1960

3873

1007

3496

2947

4068

4187

3906

3759

3027

3816

3904

4581

1001

3815

4492

782

912

1013

1588

4098

1859

3930

4062

870

4286

3804

3905

1352

4228

2612

4158

530

**B**

**4**

**7**

**pI**

**14.4**

**18.4**

**25.0**

**35.0**

**45.0**

**66.2**

**116.0**

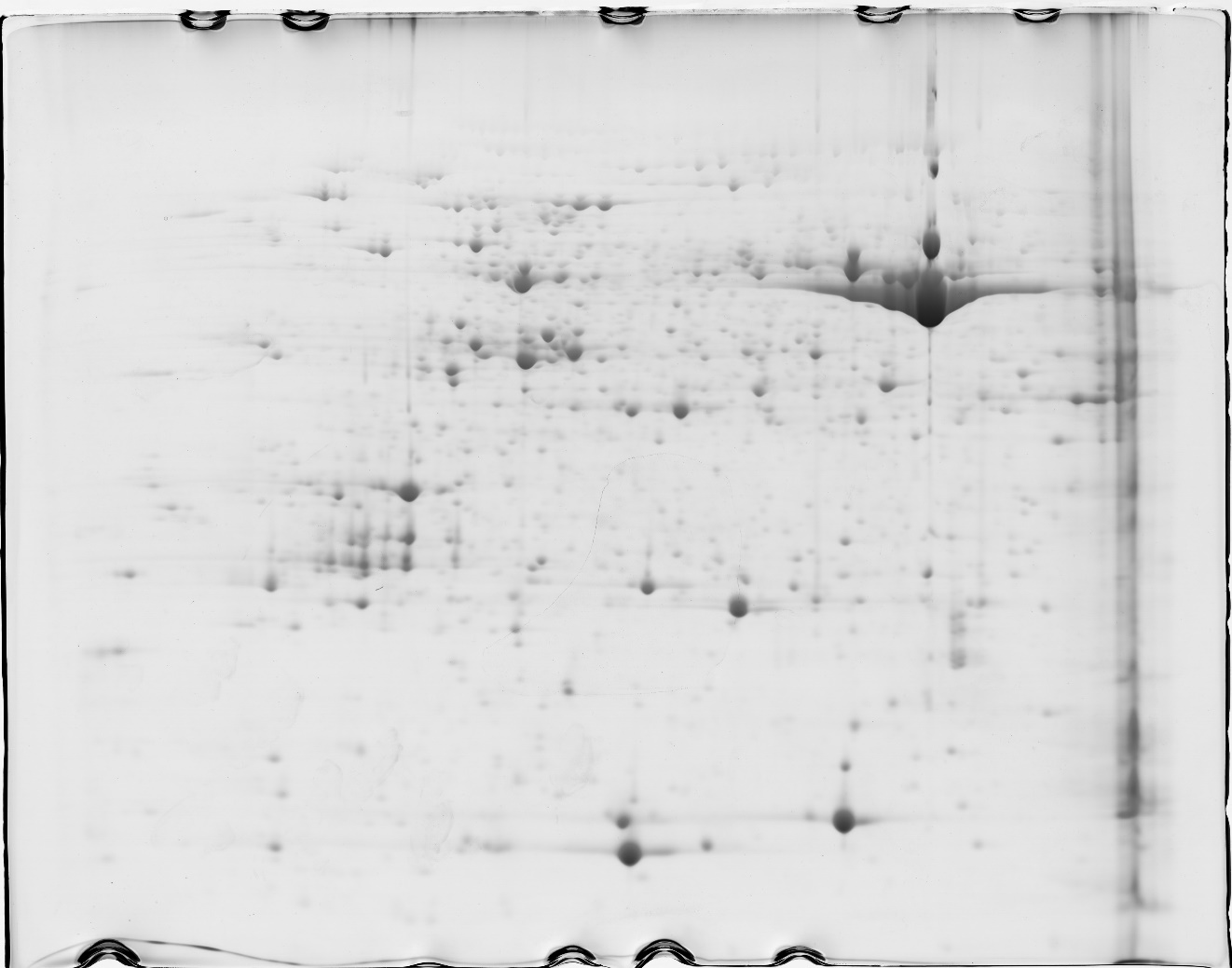

4006

4008

4196

**A**

4233

1698

974

3120

3137

3147

3232

1634

1649

1917

4109

4403

1820

2626

2778

3117

4493

2569

4019

4491

4501

2263

2264

2282

4512

4030

4066

2605

2616

2618

4240

4002

4288

4359

4272

895

1027

4274

1103

1020

1065

1171

1817

1140

1179

1193

1195

1220

4547

1446

910

1982

1862

988

4212

4151

1695

1818

4369

1599

1503

4281

1178

1781

1929

1960

3873

1007

4068

4517

3906

4579

1664

3816

3904

4427

4581

1498

2927

1602

3815

4492

782

1013

1375

1588

4020

4098

1859

3930

4062

870

876

4286

1191

3804

3905

1352

4228

2612

4158

530

1072

4143

**4**

**7**

**14.4**

**18.4**

**25.0**

**35.0**

**45.0**

**66.2**

**116.0**

**pI**

**MM**

(KDa)

**MM**

(KDa)

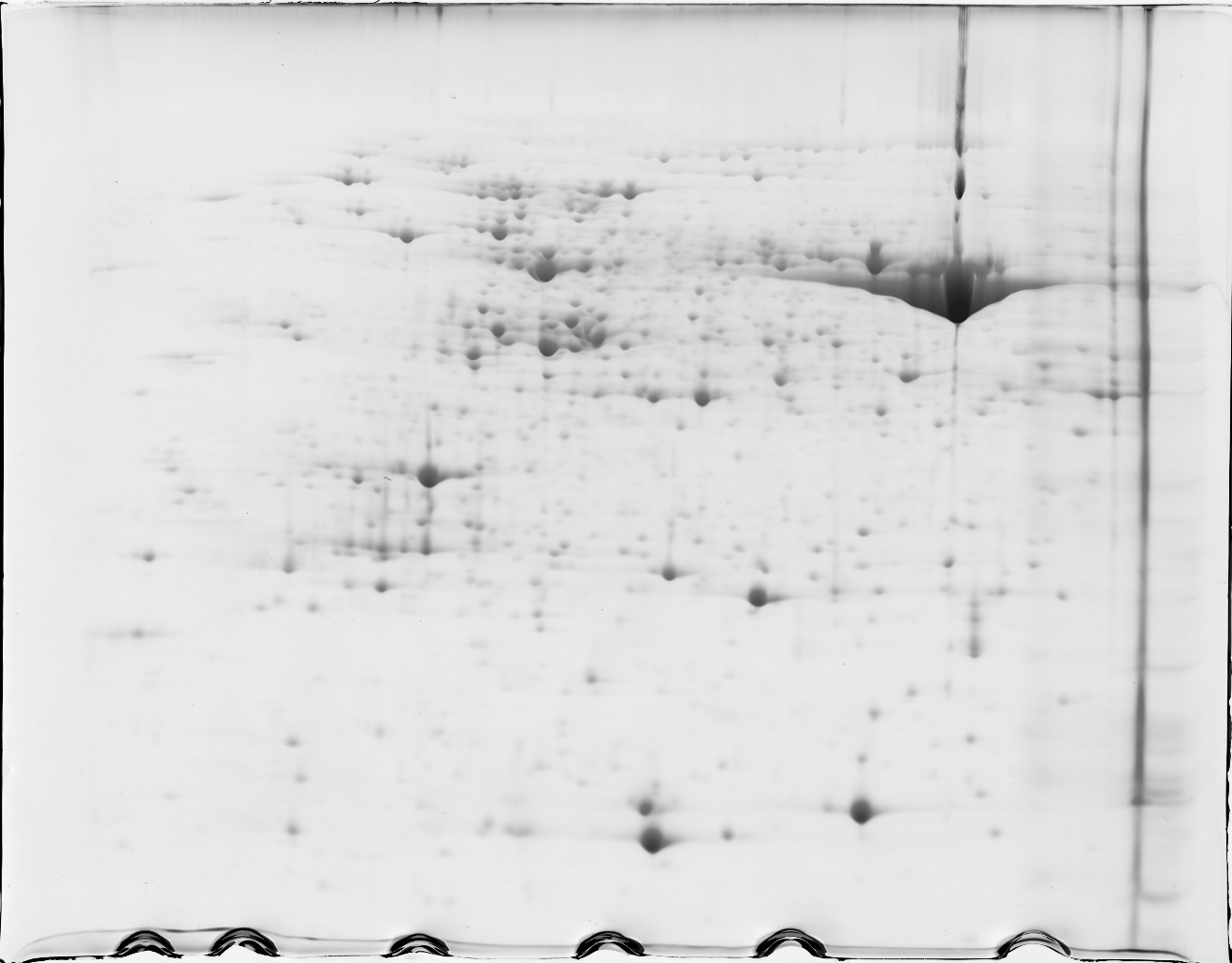

4006

4196

1698

974

1687

3147

3232

1452

1649

1917

1903

4403

1820

2626

3298

2778

4493

2569

4019

4491

4501

4501

2282

4512

4501

4066

2605

2616

3582

4240

4002

4288

4359

4272

895

1027

4274

1103

1020

1065

1817

1140

1179

1195

1820

1606

910

1982

1760

1862

988

4212

1695

1818

4115

1686

4369

3602

1503

3093

4281

1178

1781

1929

1960

3873

4068

4517

4187

3759

4579

1634

1664

3027

3904

4501

4581

1001

1498

2850

2927

3815

4492

782

912

1588

4020

4098

1859

3930

3785

4501

870

876

4286

1191

3804

3905

4501

1352

2612

4158

530

1072

3310

4233

**D**

**4**

**7**

**pI**

**14.4**

**18.4**

**25.0**

**35.0**

**45.0**

**66.2**

**116.0**

**MM**

(KDa)

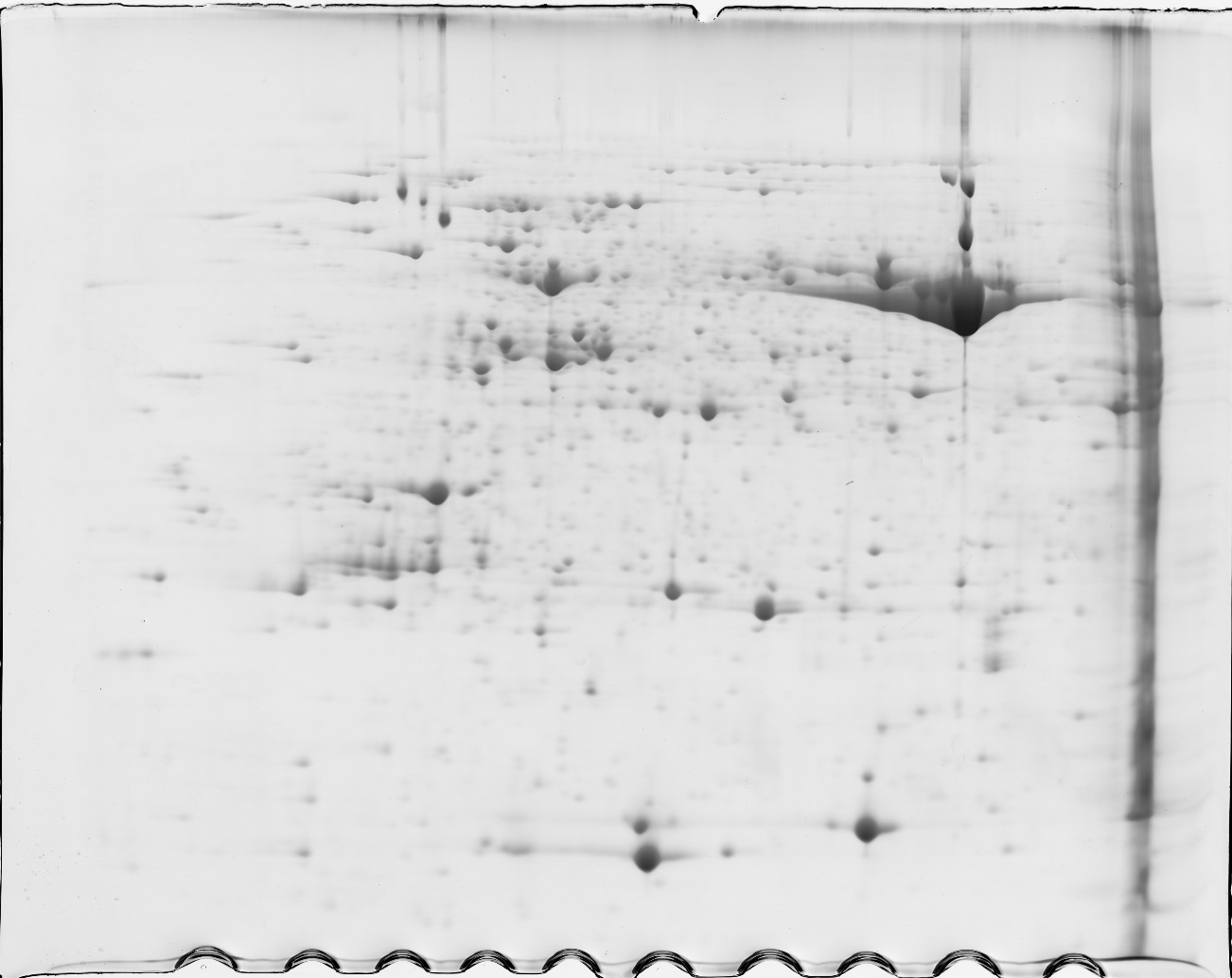

4006

4196

4233

974

1687

3147

3066

3232

1634

1917

4109

1903

1820

2626

3298

2778

3117

4493

2569

4019

4491

2282

3104

4030

4066

2605

2616

2618

4240

4002

4288

4359

4272

895

4274

1065

1171

1817

1140

1179

1193

1195

4547

1606

910

1760

1862

4081

3003

988

4212

4151

3734

3083

1695

1818

4115

1503

3093

4281

1929

1960

3873

3496

2947

4068

4517

3906

4579

3006

3027

3816

3904

4427

4581

1001

1498

2850

2927

1602

3815

4492

782

912

1588

4020

4098

1859

3930

3785

870

876

3871

4286

3804

3905

1352

2612

4158

530

4143

**C**

**4**

**7**

**pI**

**14.4**

**18.4**

**25.0**

**35.0**

**45.0**

**66.2**

**116.0**

**MM**

(KDa)

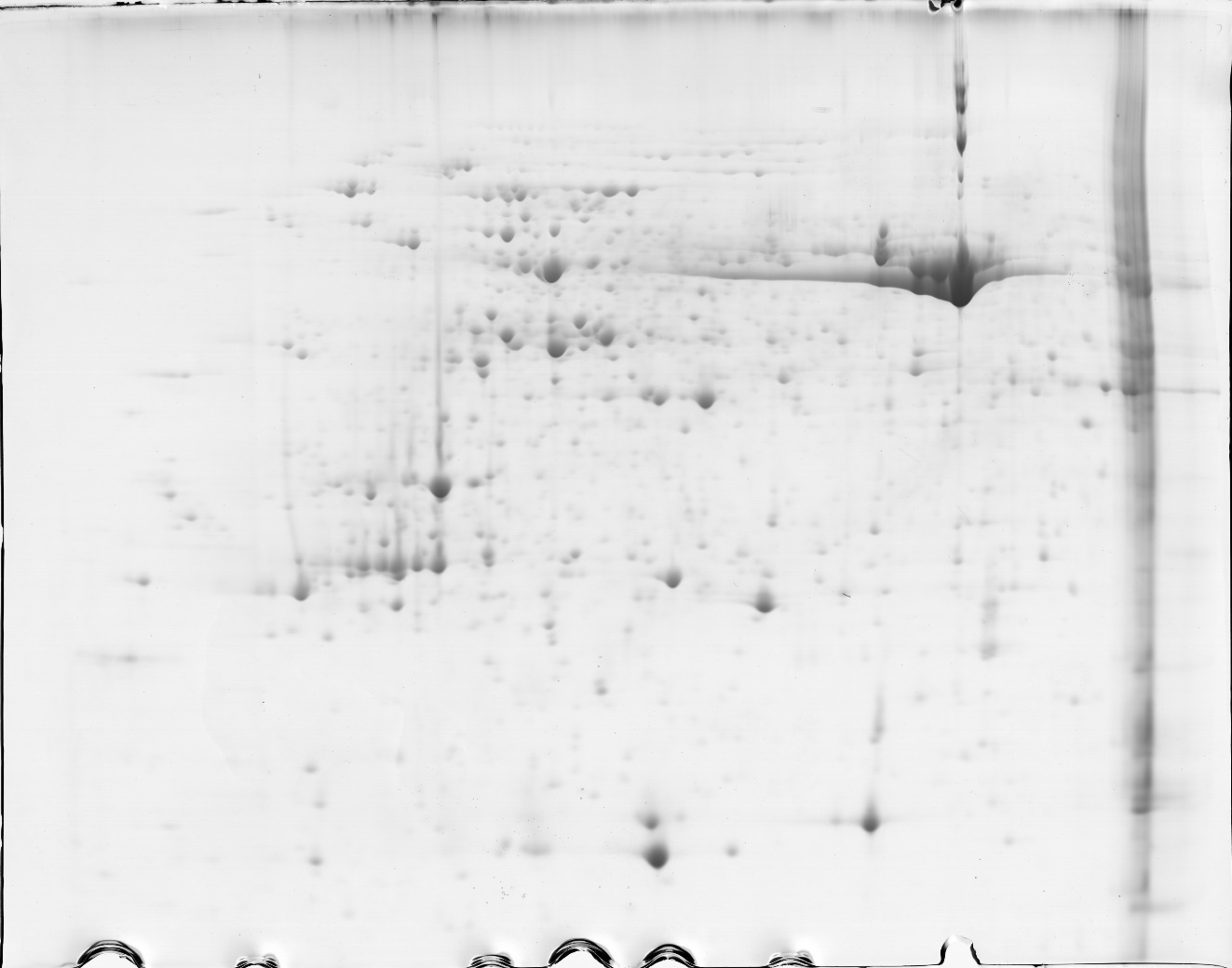

4006

4008

1456

3120

3137

3147

3066

3232

1634

1649

4109

4403

1820

2626

3298

3117

4493

2569

4019

4491

4501

2263

2264

3104

4512

4066

2605

2616

2618

3582

4240

4002

4288

4359

895

1027

4274

1141

1020

1171

1817

1140

1179

1195

1446

1606

910

1862

1917

4081

3003

4212

4151

3734

3083

1695

1818

1686

1503

4281

1178

1781

1929

1960

3873

1007

3496

2947

4068

4517

3906

3759

4579

1452

3006

3816

3904

4427

4581

1001

2927

4492

782

1375

1588

4098

1859

3930

3785

4062

870

876

3871

4286

1191

3905

1352

4228

2612

4158

530

1072

3310

4143

4196

4233

**E**

**4**

**7**

**pI**

**14.4**

**18.4**

**25.0**

**35.0**

**45.0**

**66.2**

**116.0**

**MM**

(KDa)

### Supplemental Figure 6

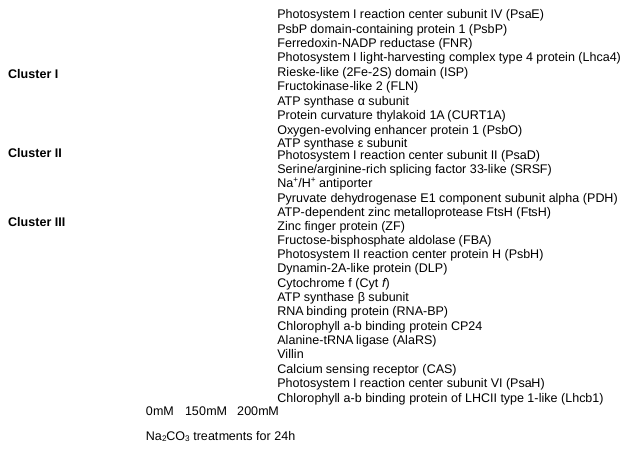

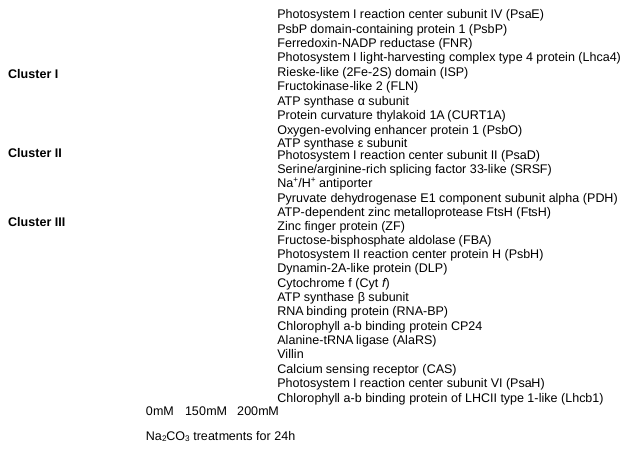

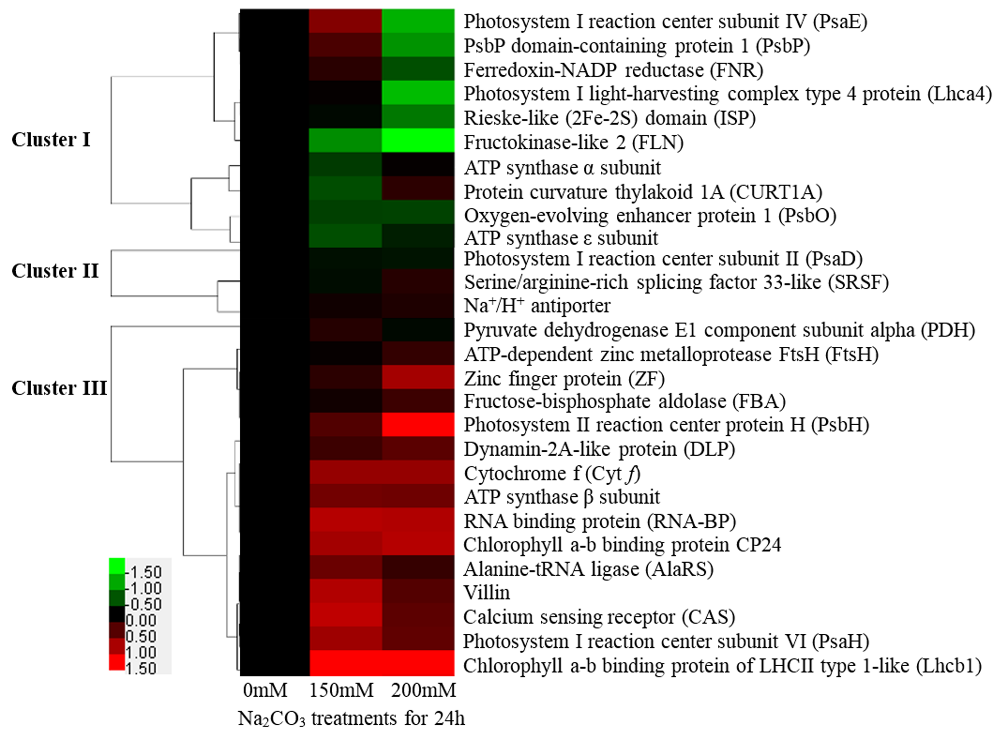
