## Supplemental Figure 4 for "Na_2_CO_3_-Responsive Photosynthetic and ROS Scavenging Mechanisms in Chloroplasts of Alkaligrass Revealed by Phosphoproteomics"

1. Chloroplast phosphoproteome

Accession No. ERM94529

NPGS_p_VNQDPIFK

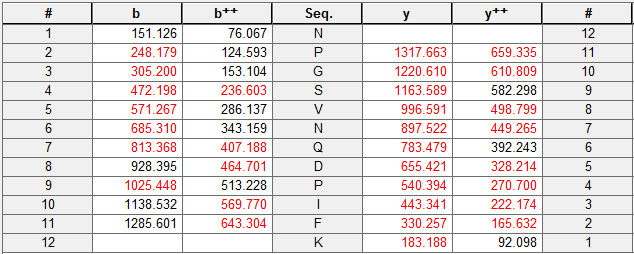

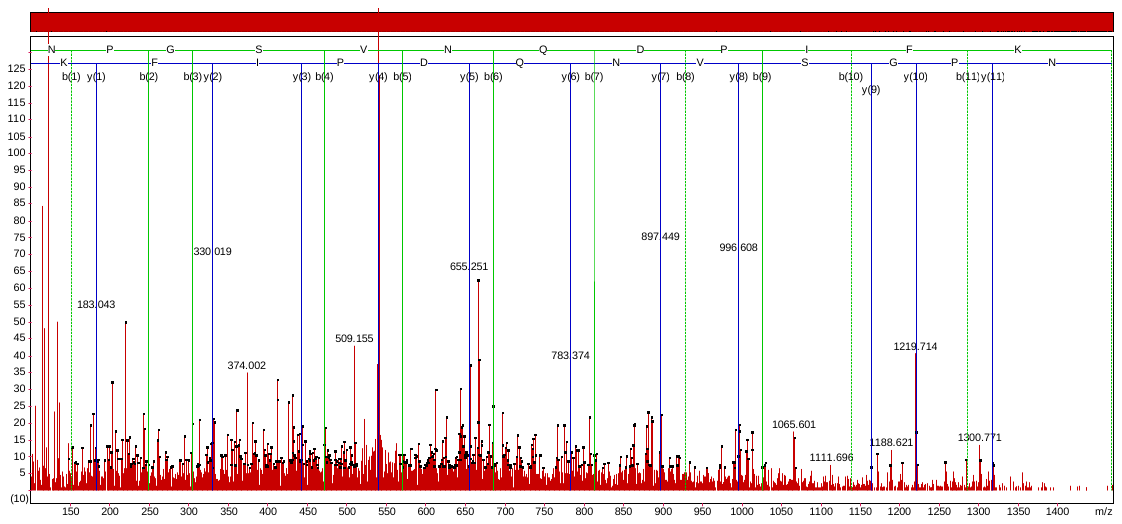

Accession No. BAK03973

VAAS_p_SSPWYGSDR

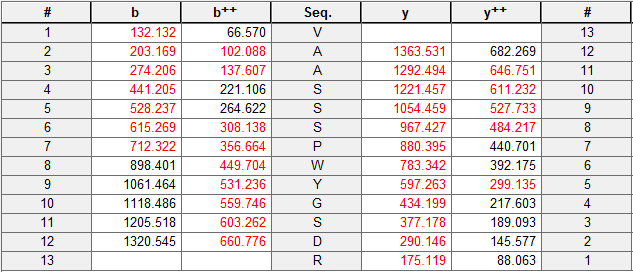

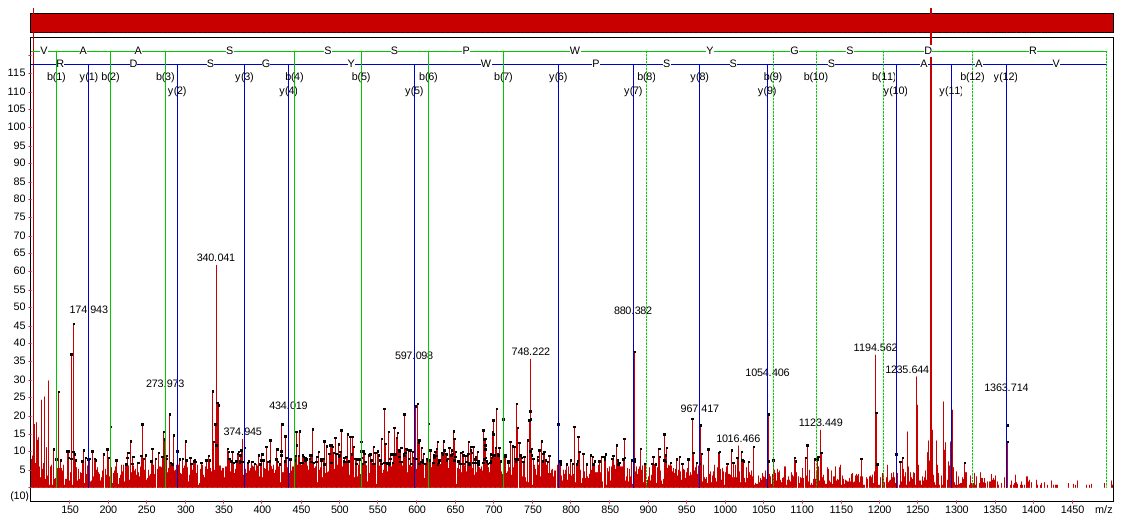

VAASSS_p_PWYGSDR

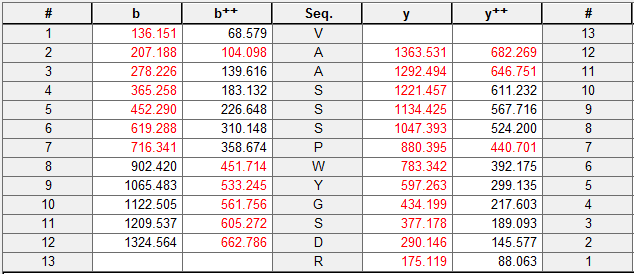

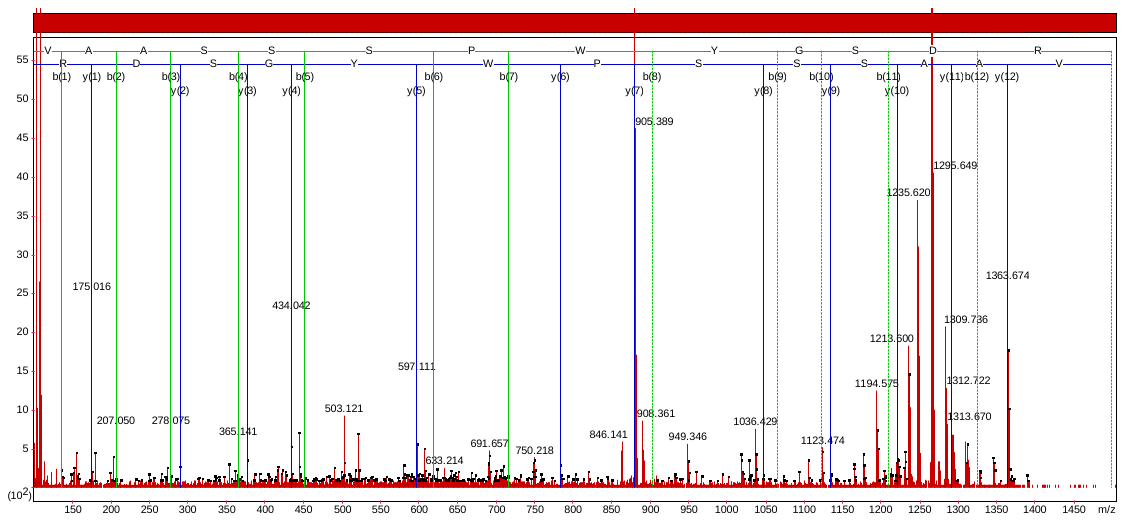

VAASSSPWYGS_p_DR

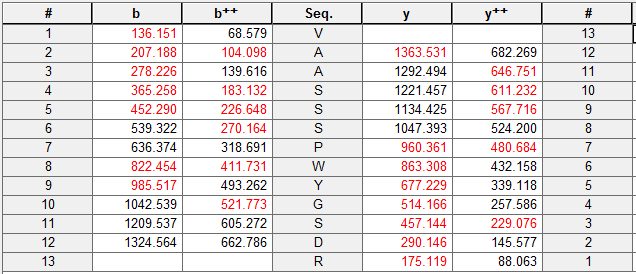

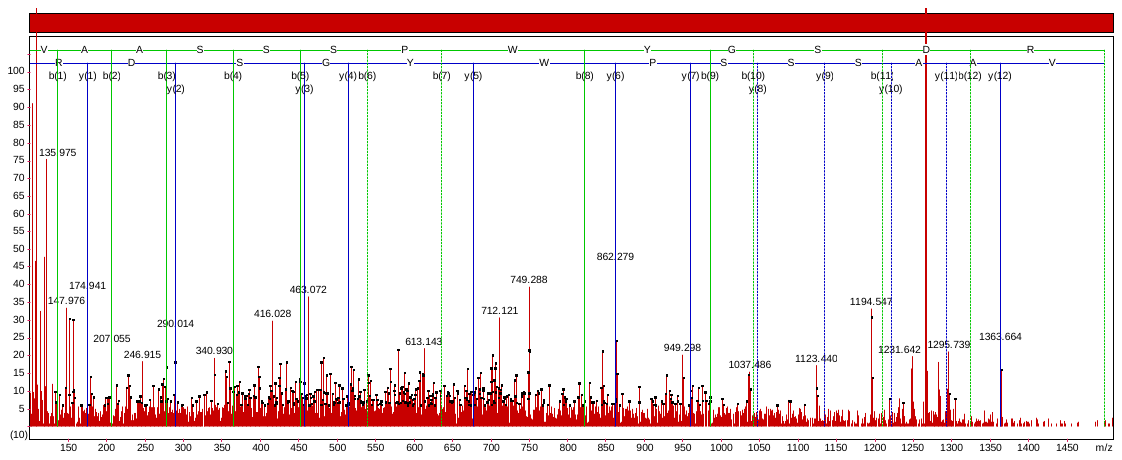

Accession No. ADL41158

AKPVS_p_SGSPWYGSDR

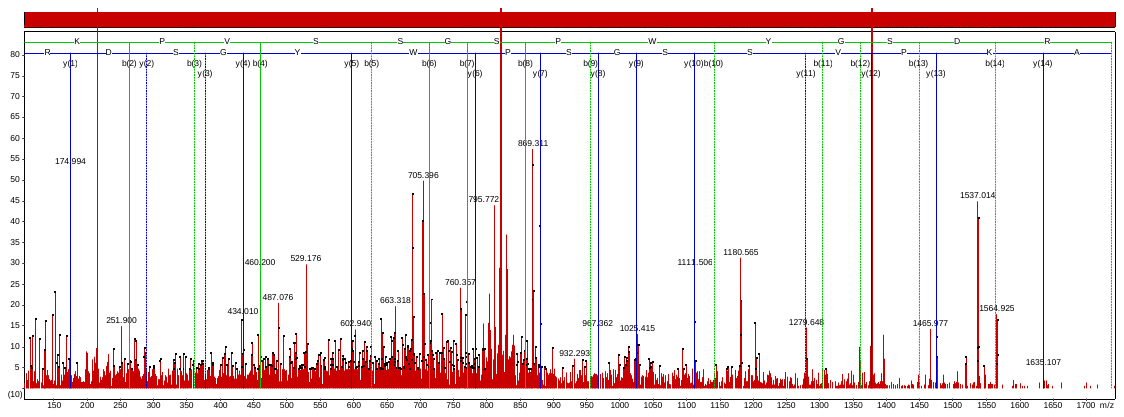

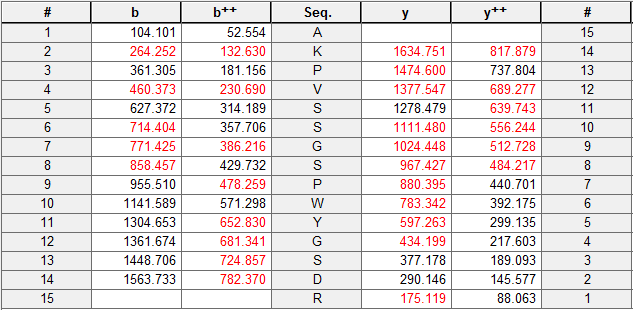

AKPVSSGS_p_PWYGSDR

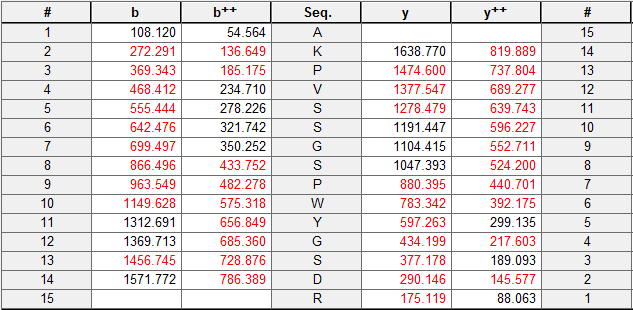

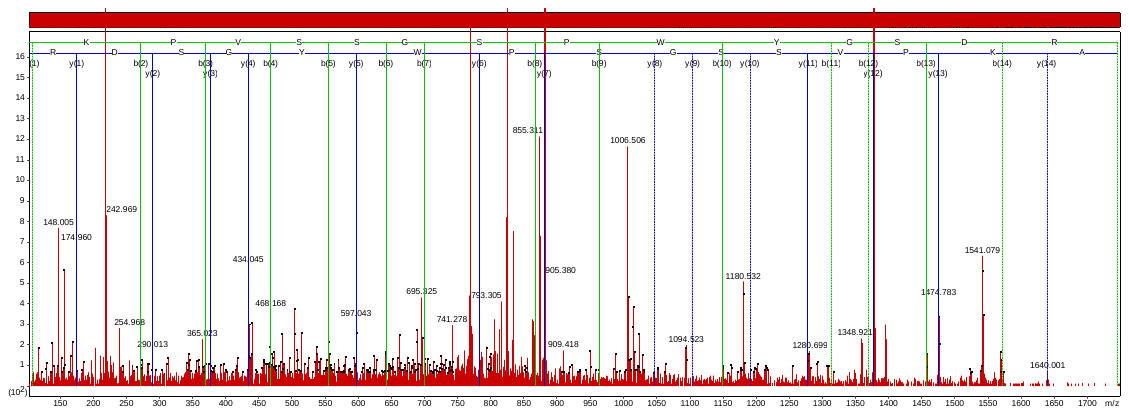

AKPVSSGSPWYGS_p_DR

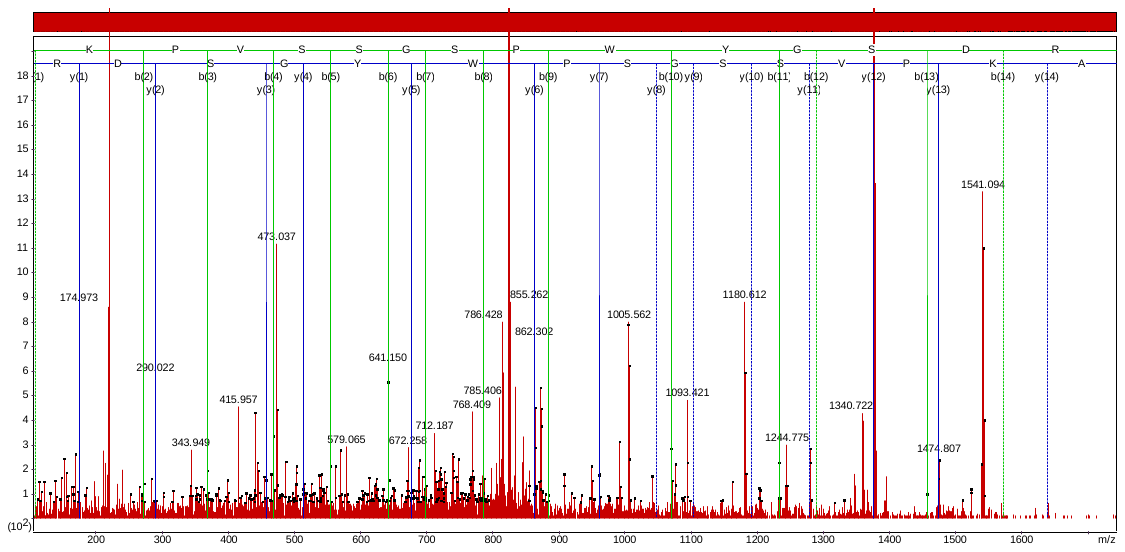

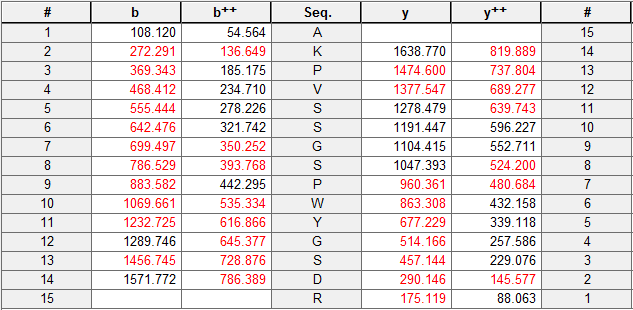

Accession No. BAF12500

TVKS_p_APQSIWYGPDRPK

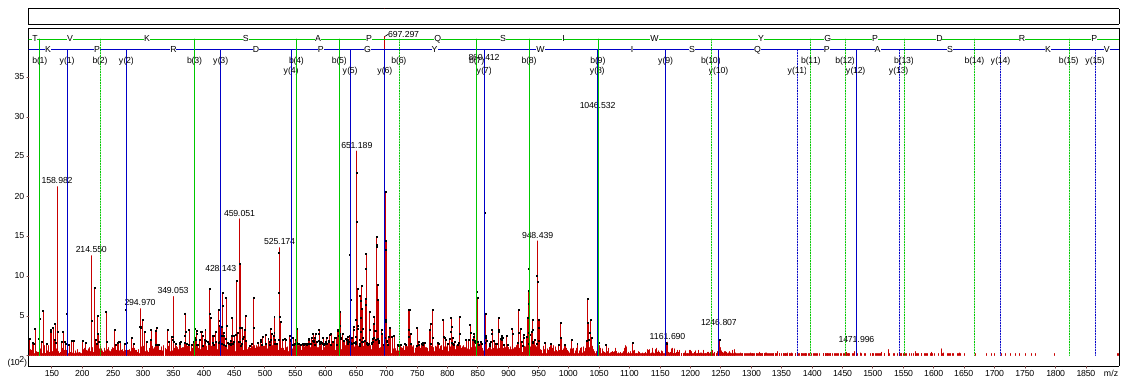

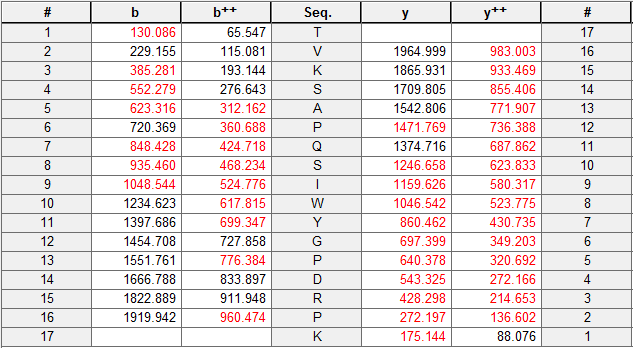

Accession No. BAJ85110

QVSS_p_GSPWYGADR

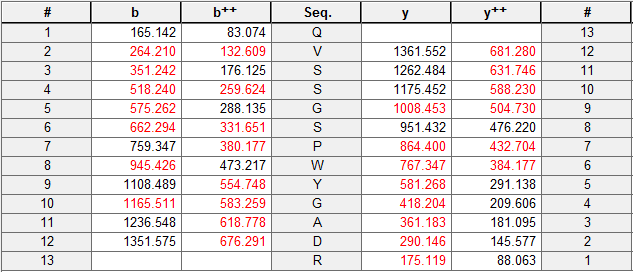

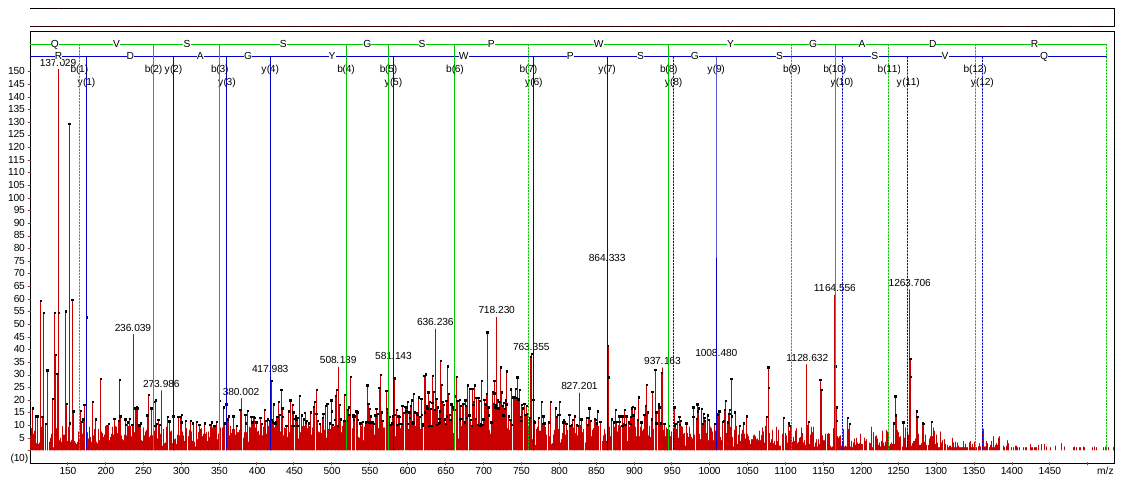

Accession No. P27523

IT_p_MGNDLWYGPDR

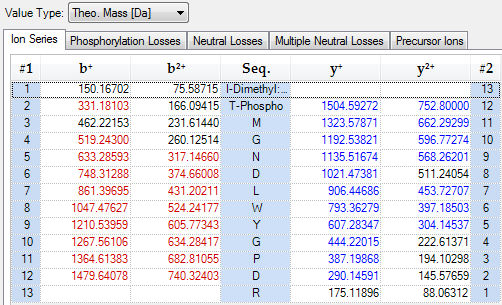

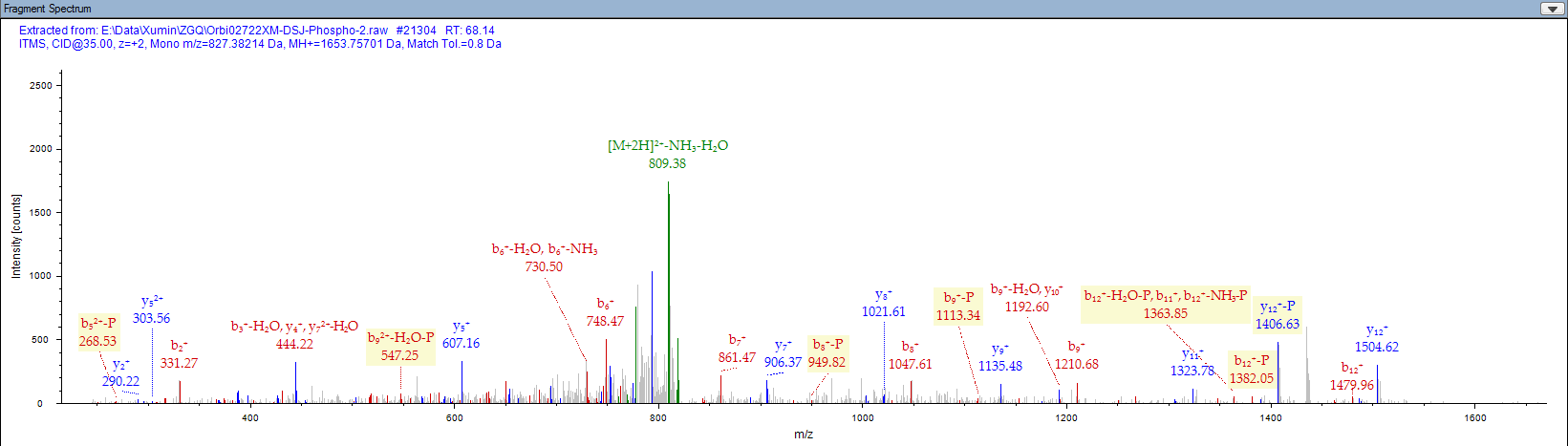

Accession No. XP_003579885

TADNFANS_p_TGDQGYPGGK

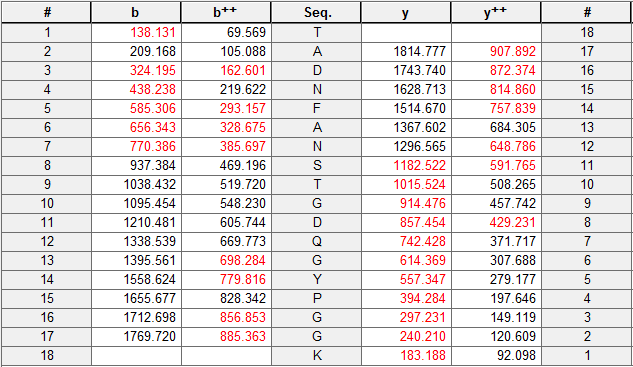

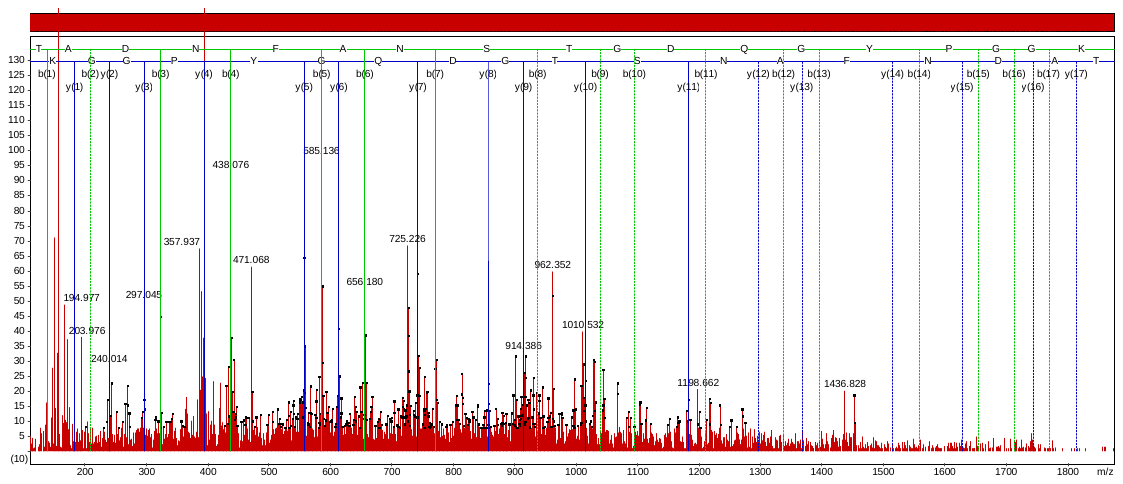

Accession No. A1EA37

AT_p_QTVEDSSKPR

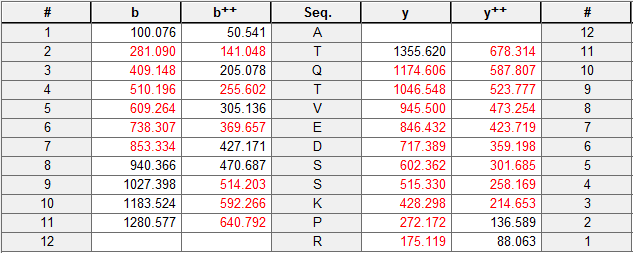

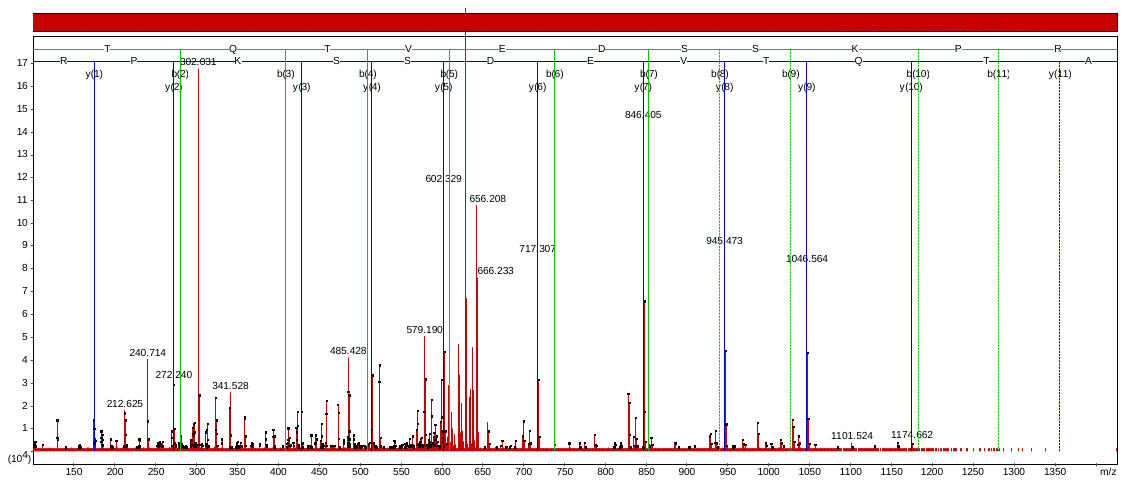

AT_p_QT_p_VEDSSKPRPK

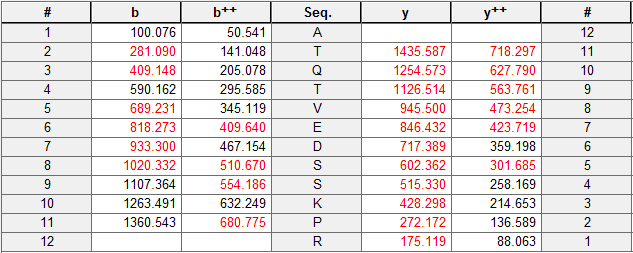

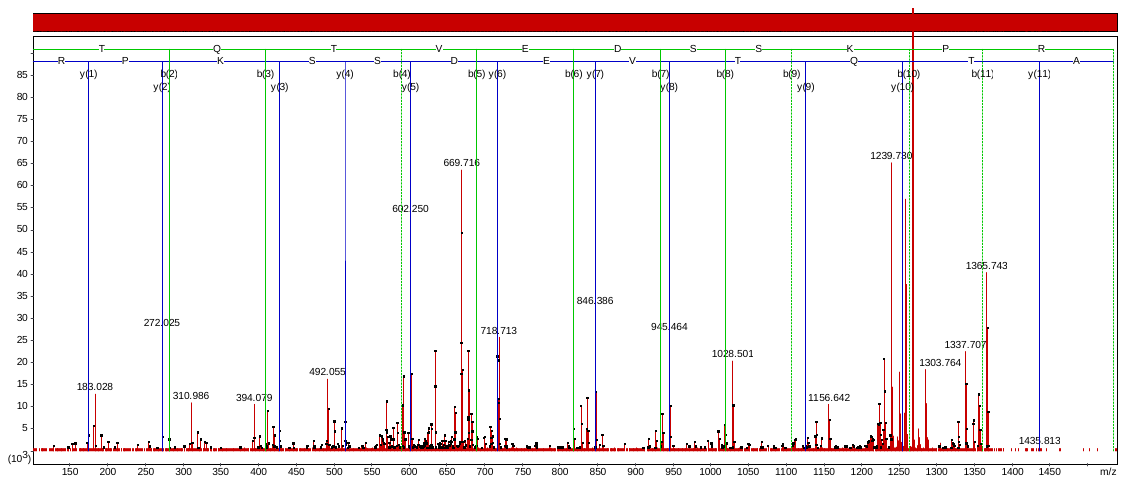

Accession No. ACF08666

AT_p_QT_p_VEDSSKPKPR

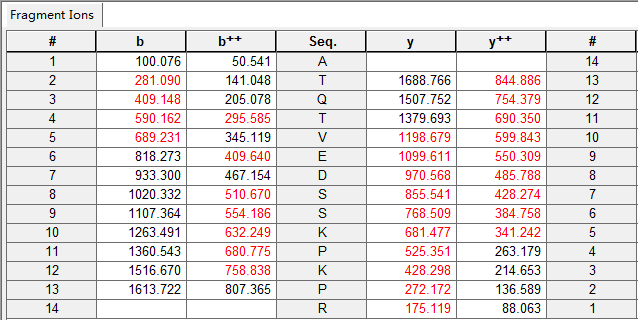

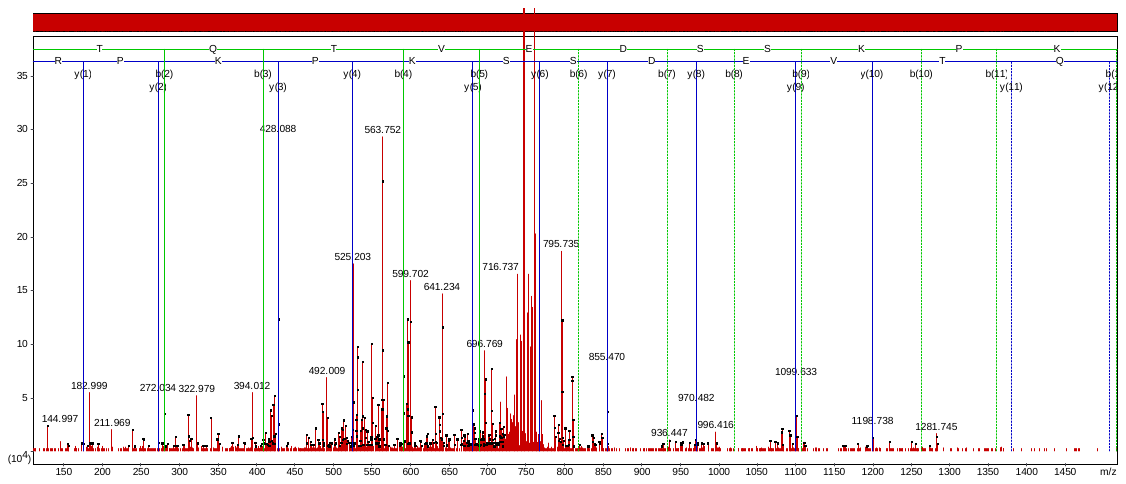

Accession No. EMT33794

QLVATGKPES_p_FSGPFLVPSYR

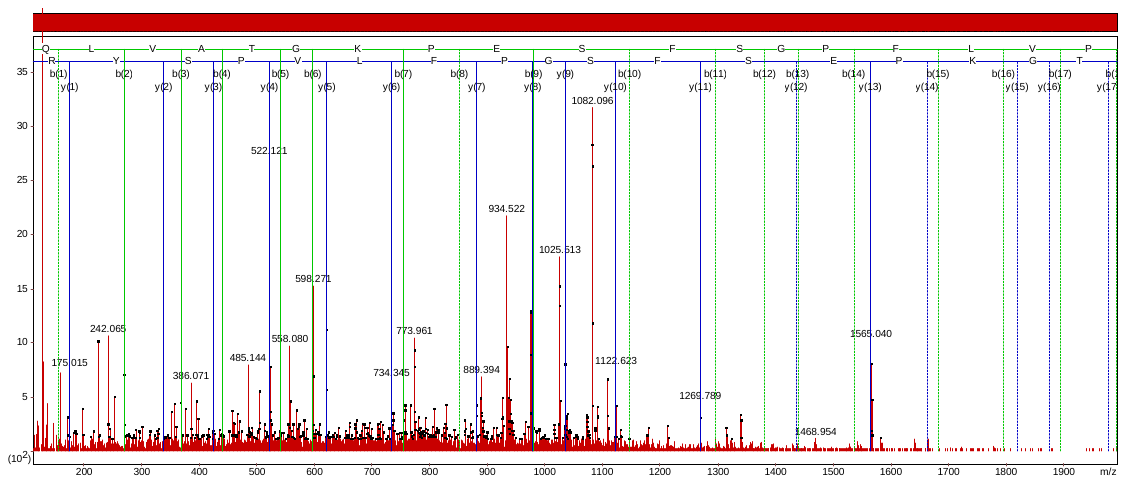

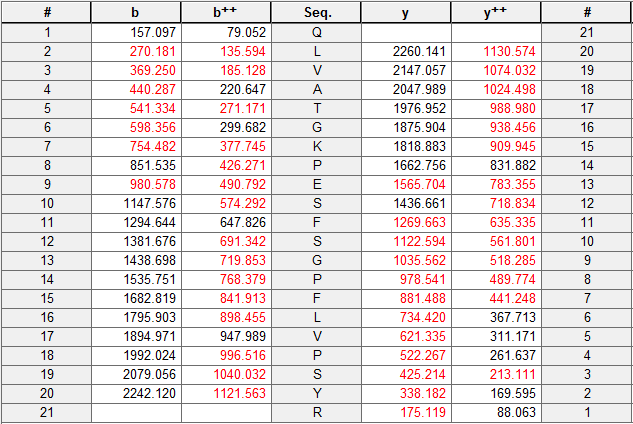

Accession No. EMS62589

SYAS_p_NNELAVMPK

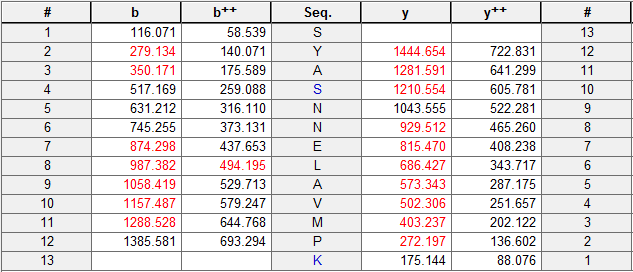

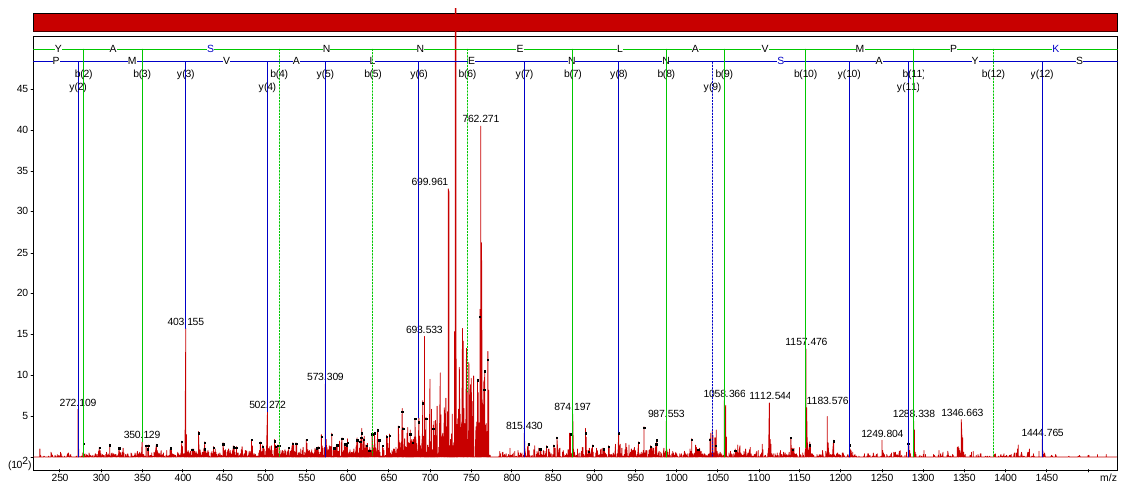

Accession No. AAN32350

T_p_ASNPNEQNVELNR

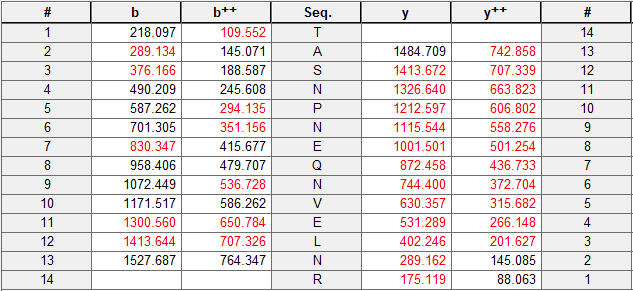

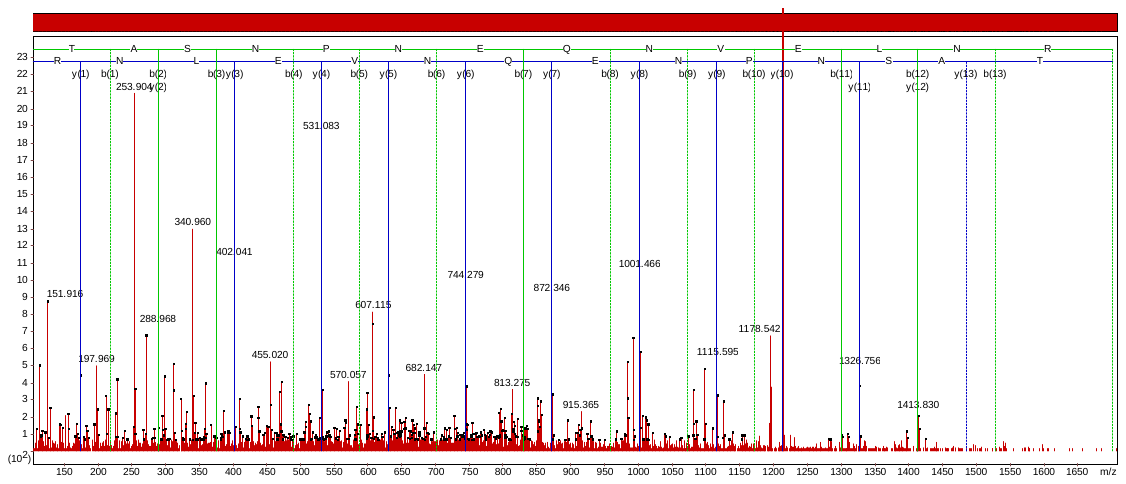

Accession No. BAJ87776

TLYSAYGSS_p_GQWGFFDK

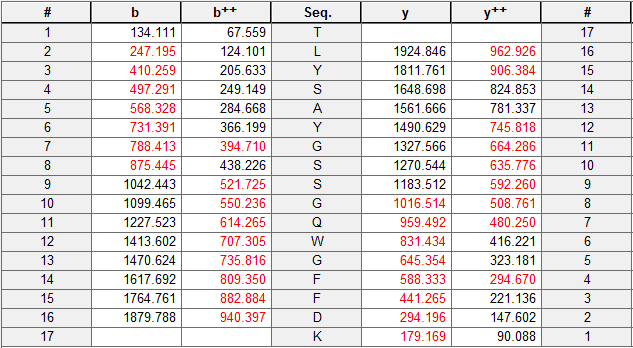

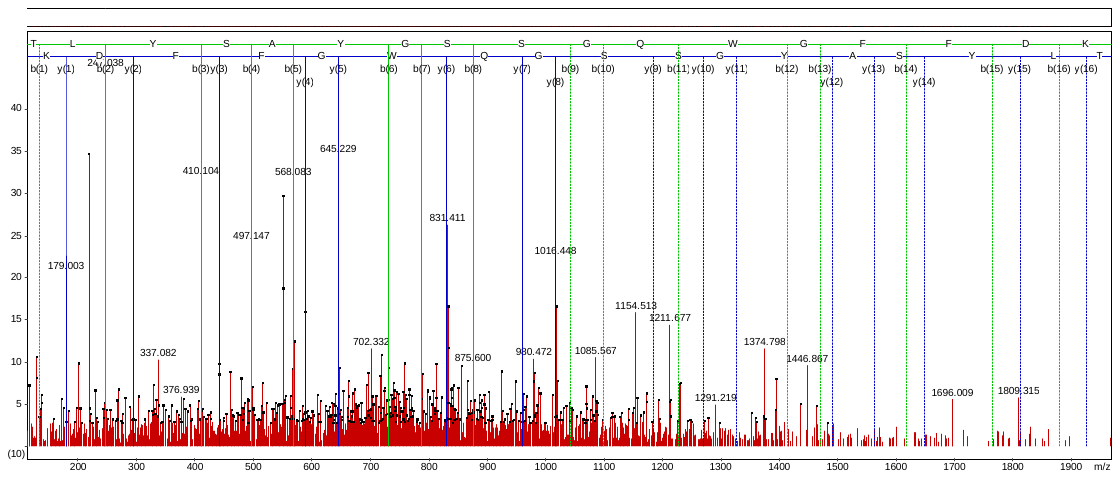

Accession No. P0C359

VYLGPET_p_TR

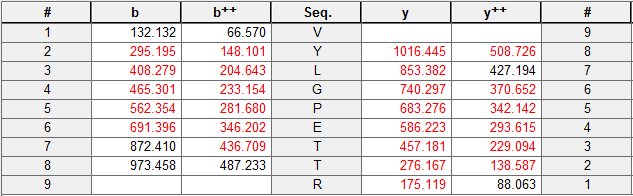

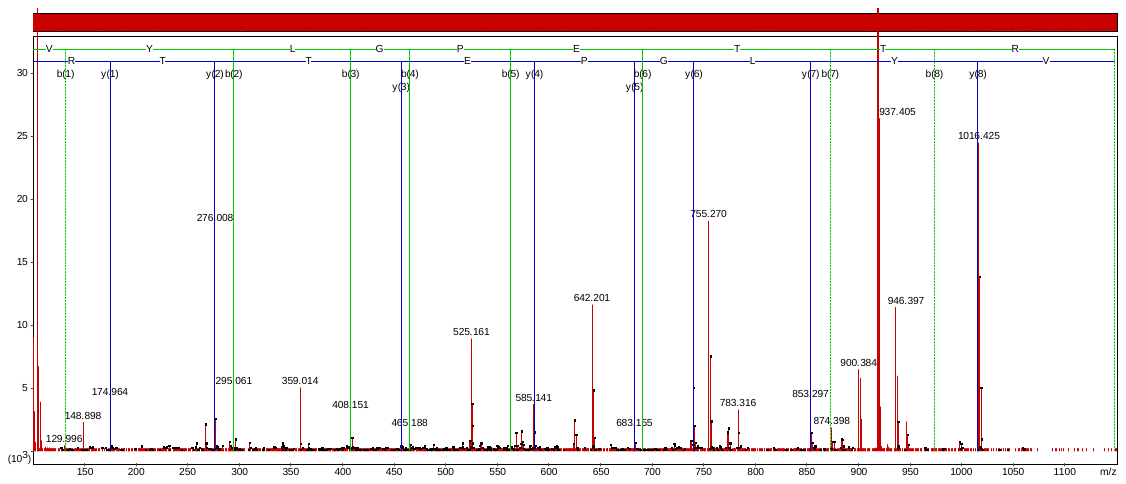

Accession No. XP_003564108

VFPS_p_GEVQYLHPK

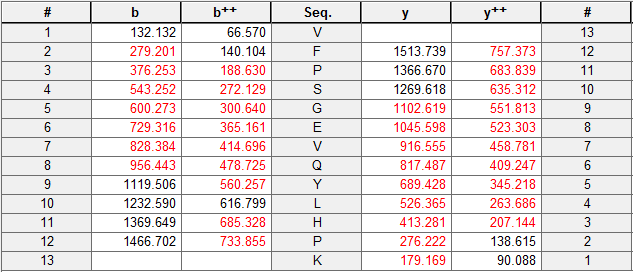

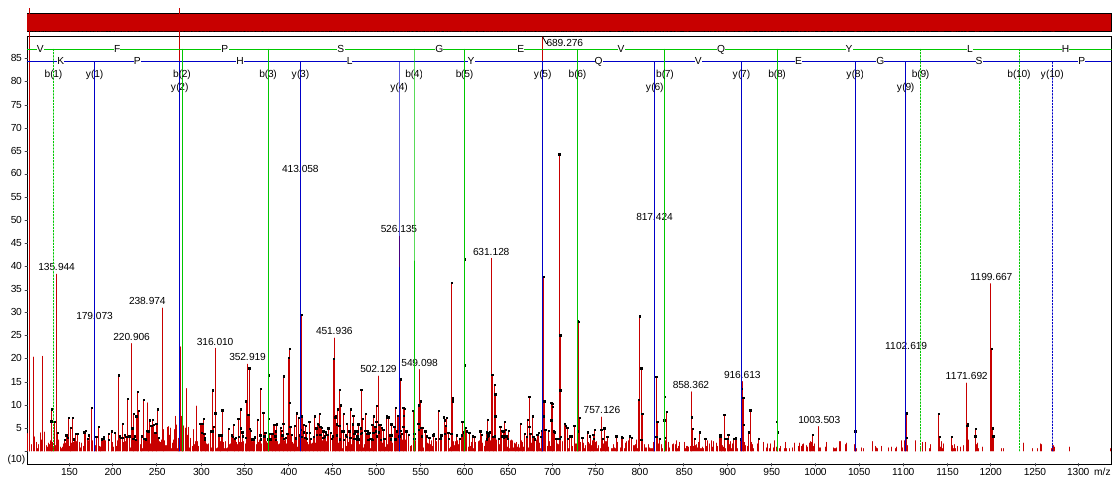

Accession No. XP_003563195

ESYWYNGTGS_p_VVTVDQDPNTR

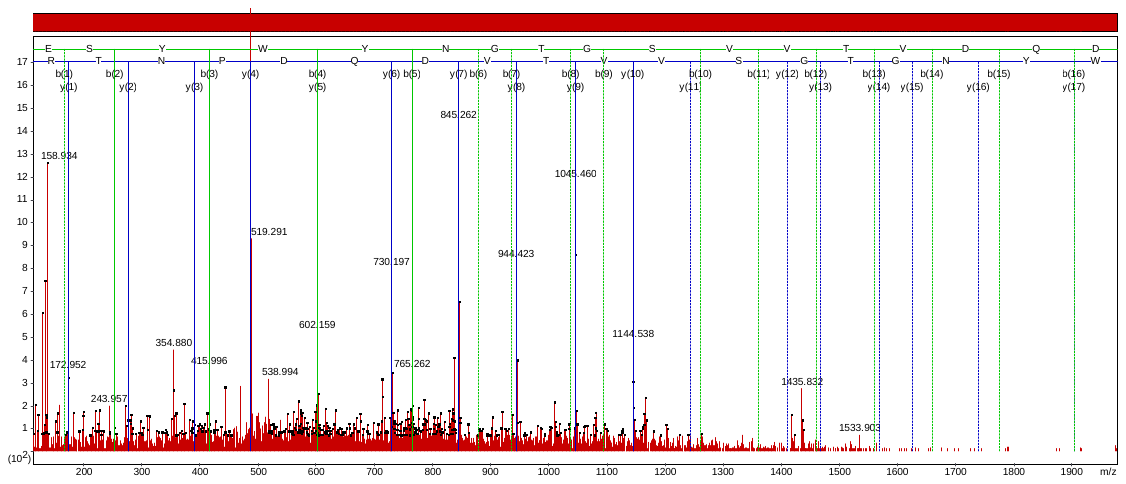

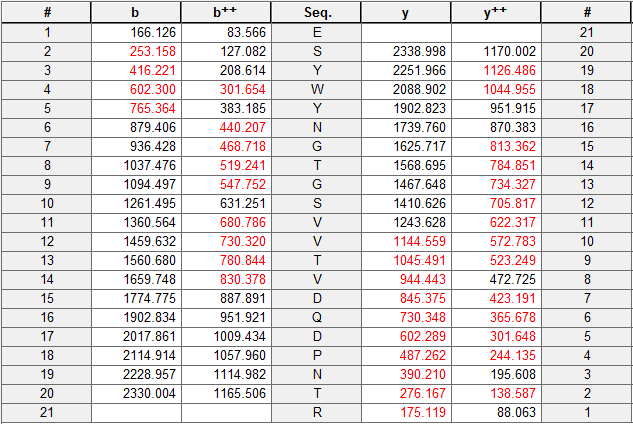

Accession No. ACG30530

VNYAGVS_p_TNNYALDEVLEVK

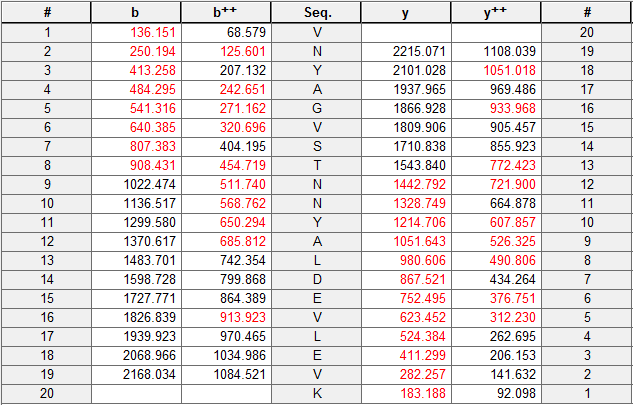

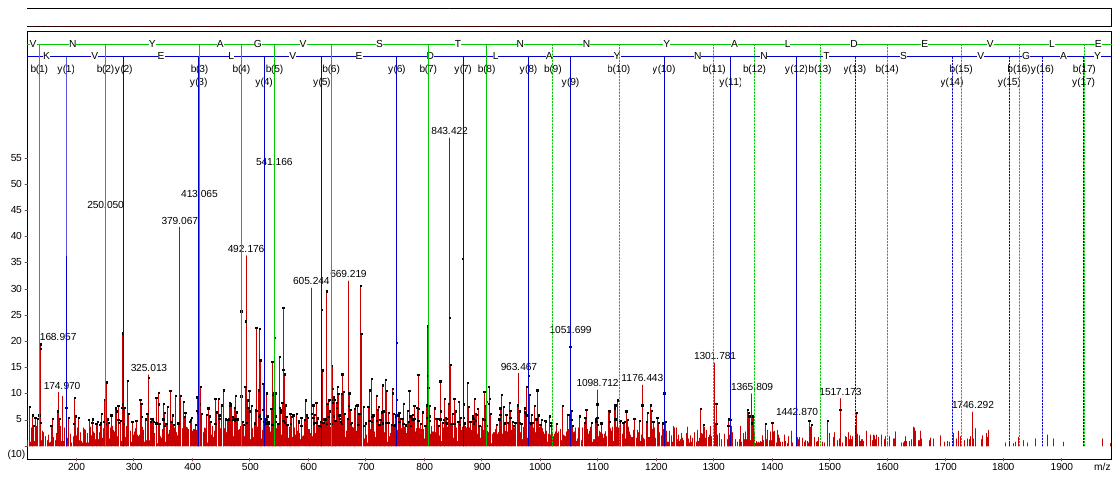

Accession No. EMT19581

GPQLPPT_p_PGPR

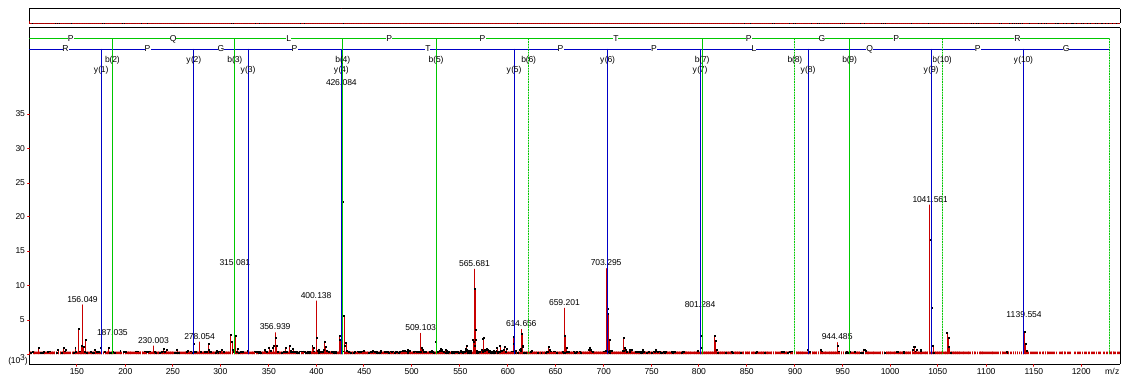

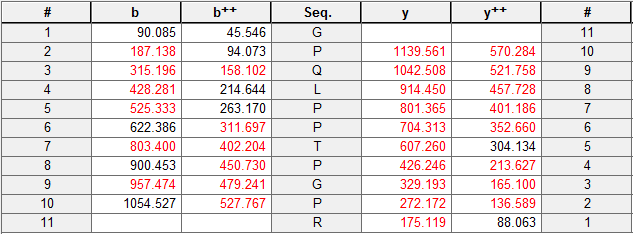

Accession No. A1EA21

VQLY_p_EMNF

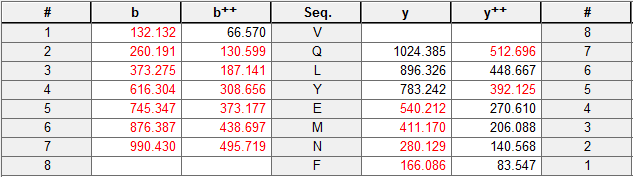

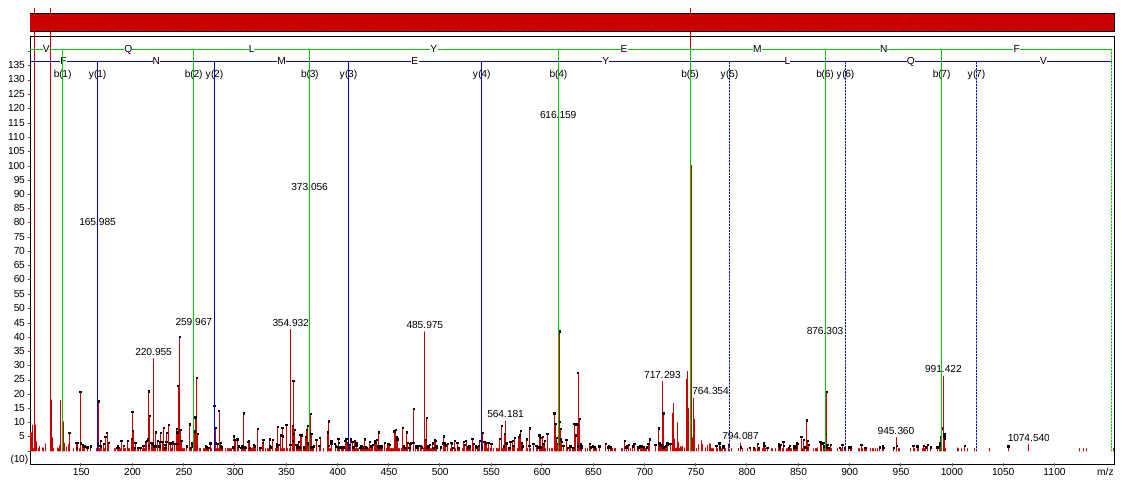

Accession No. BAK01659

AQPGSTAS_p_DVNIEEVR

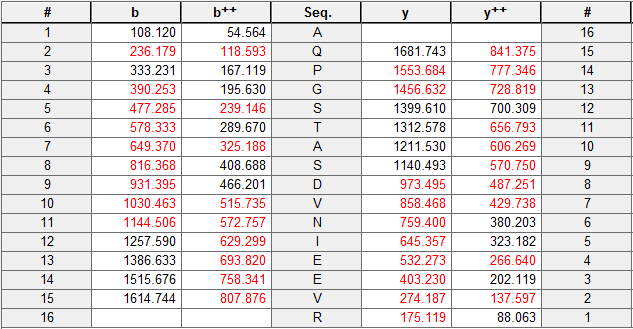

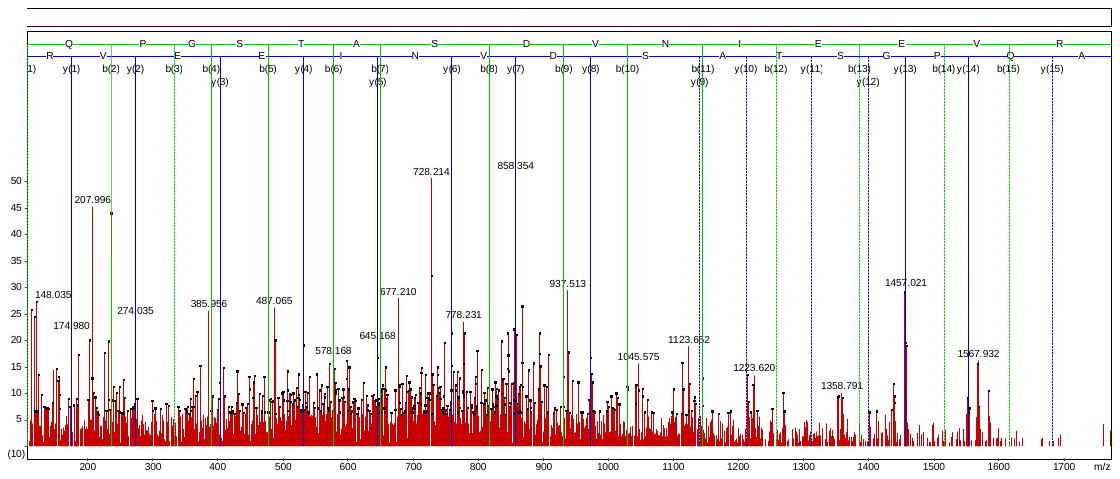

Accession No. KEH27588

LVYT_p_NDAGEVVK

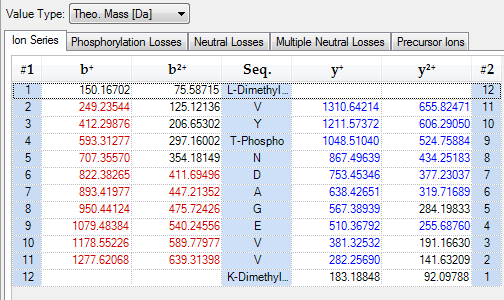

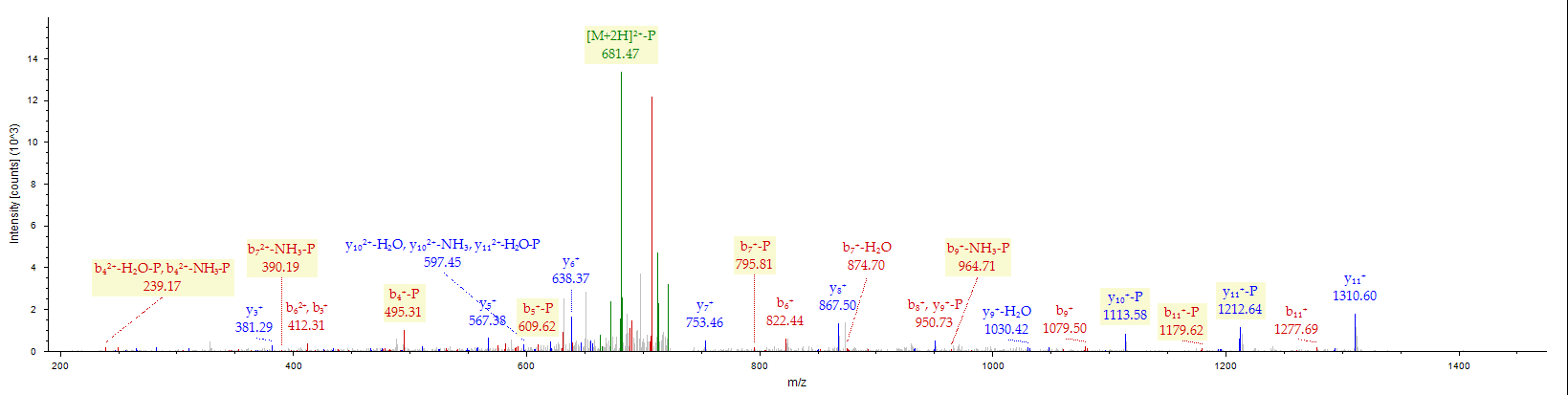

Accession No. EEH57944

LGIAPIIMS_p_AGELESGNAGEPAK

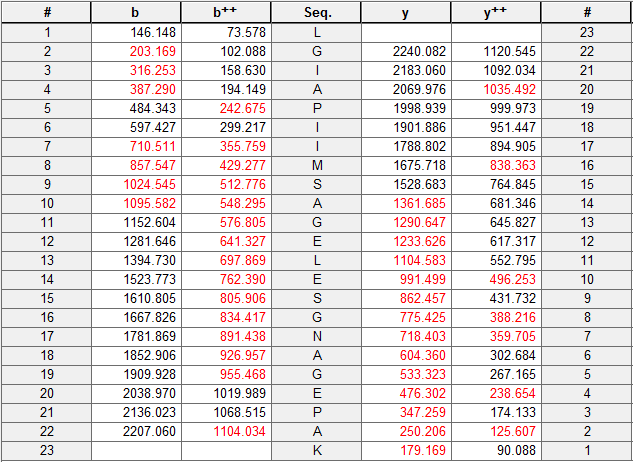

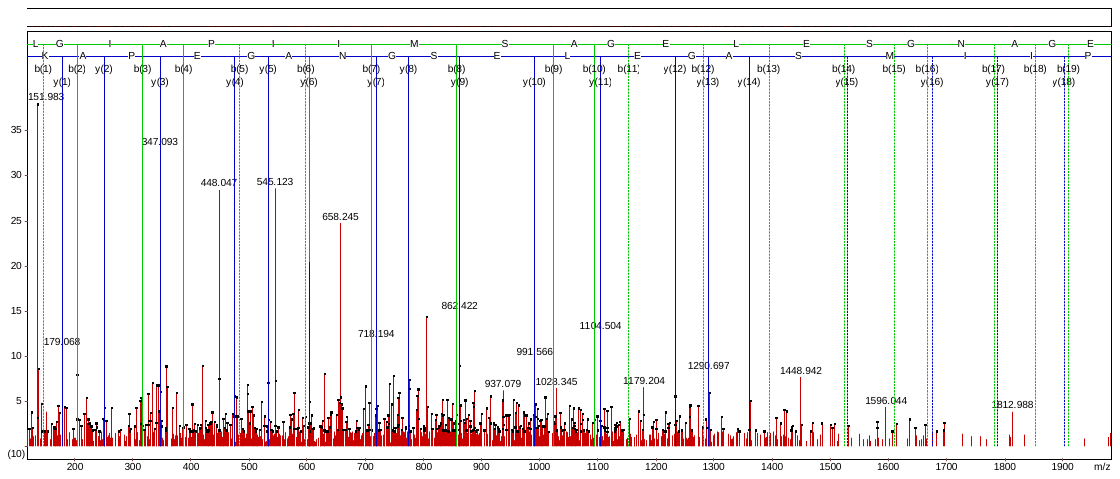

Accession No. AAB70542

GLVPSAGS_p_NNESWCQGLDGLASR

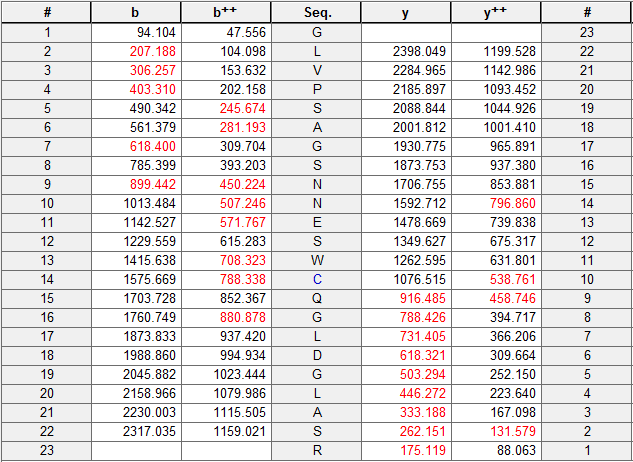

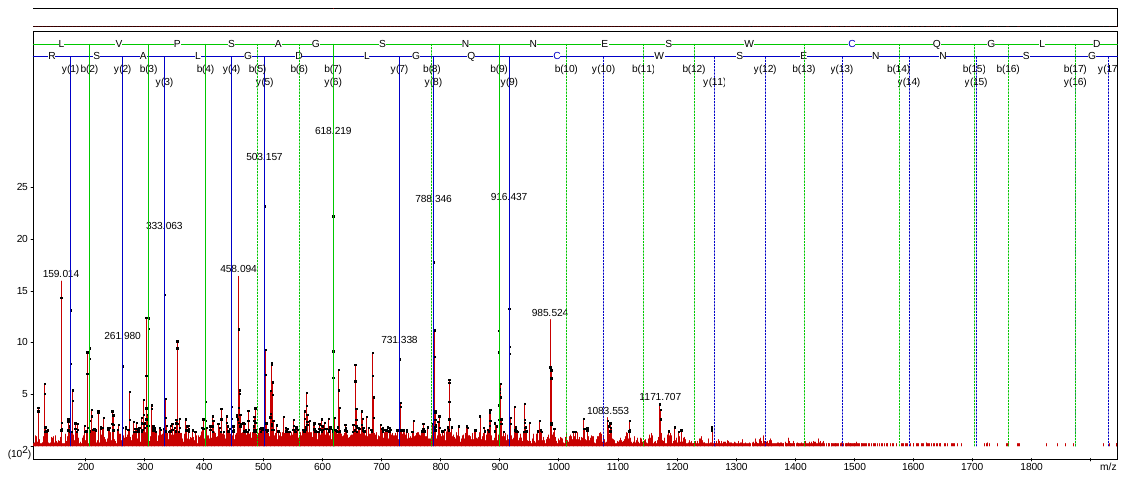

Accession No. KEH17711

GEIIAS_p_ESR

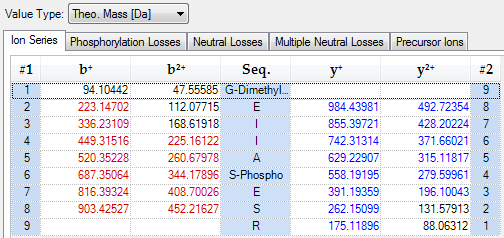

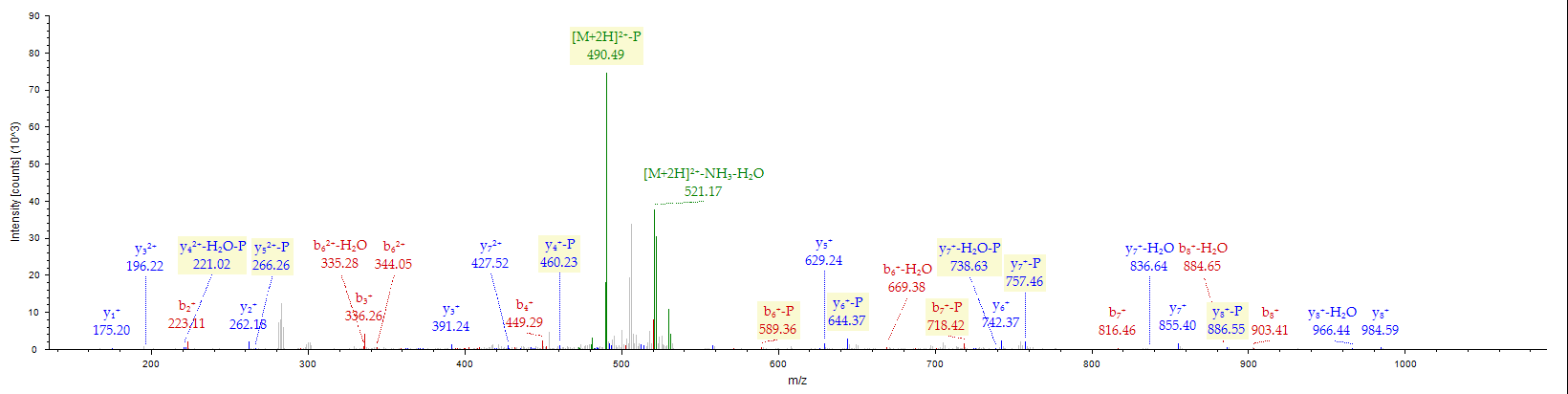

Accession No. ABH02573

AITLEEENKS_p_KK

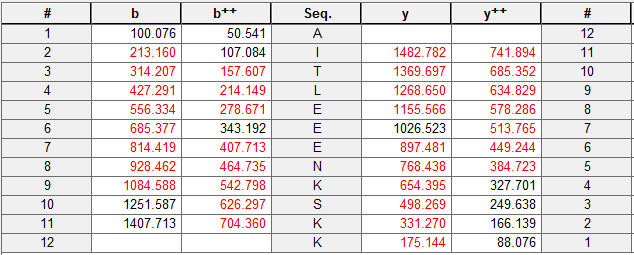

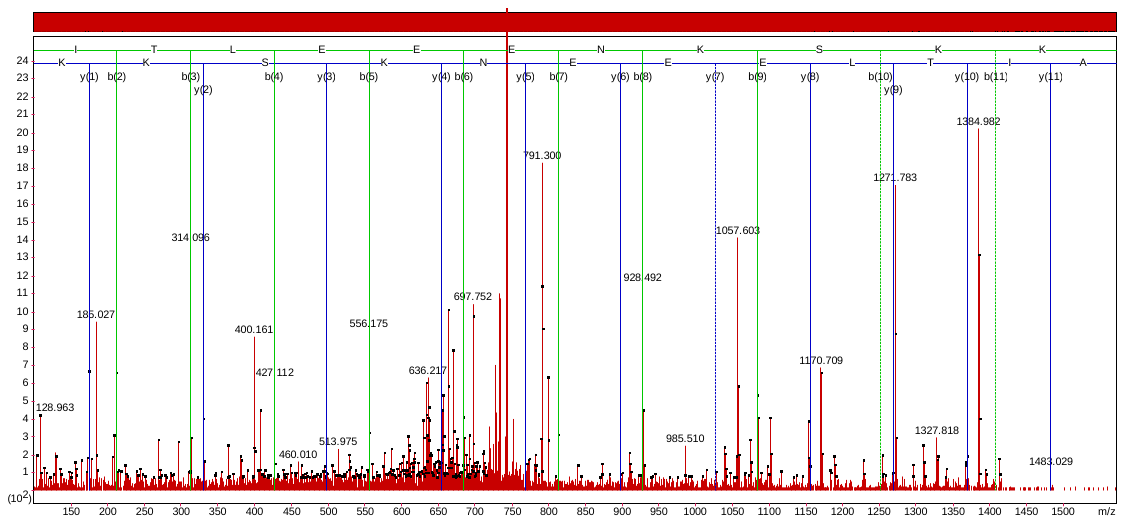

Accession No. CAB89989

GRNTGGQPINAT_p_CEVQQLLGNNR

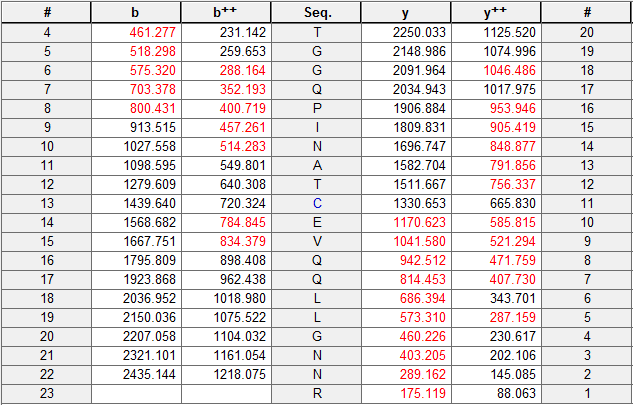

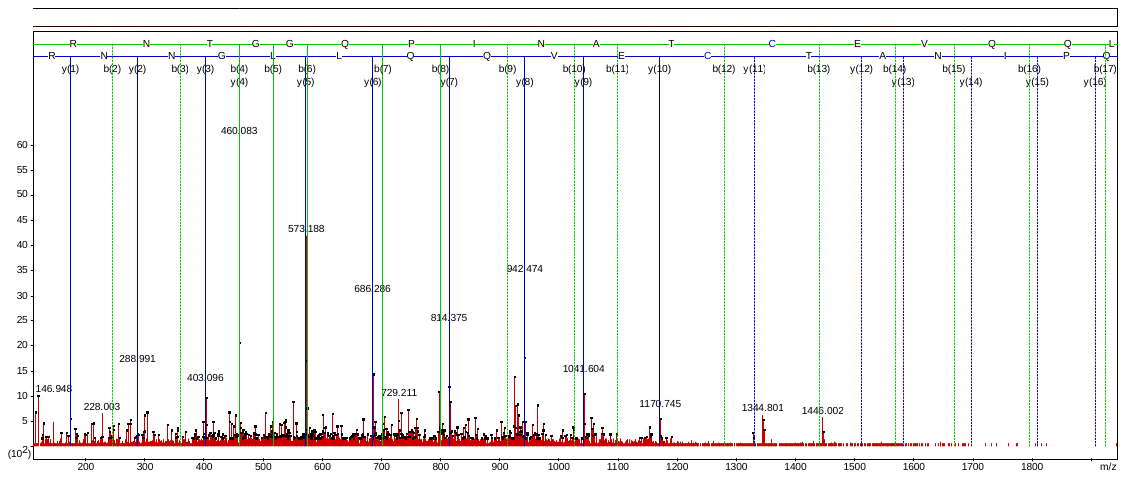

Accession No. AAK72724

DT_p_VRQQINVTCEVQQLLGNNR

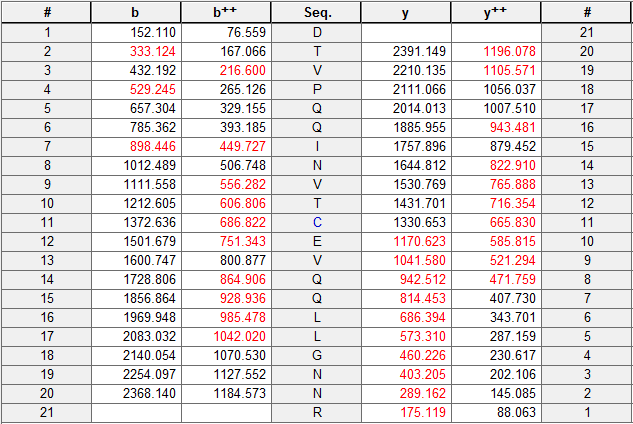

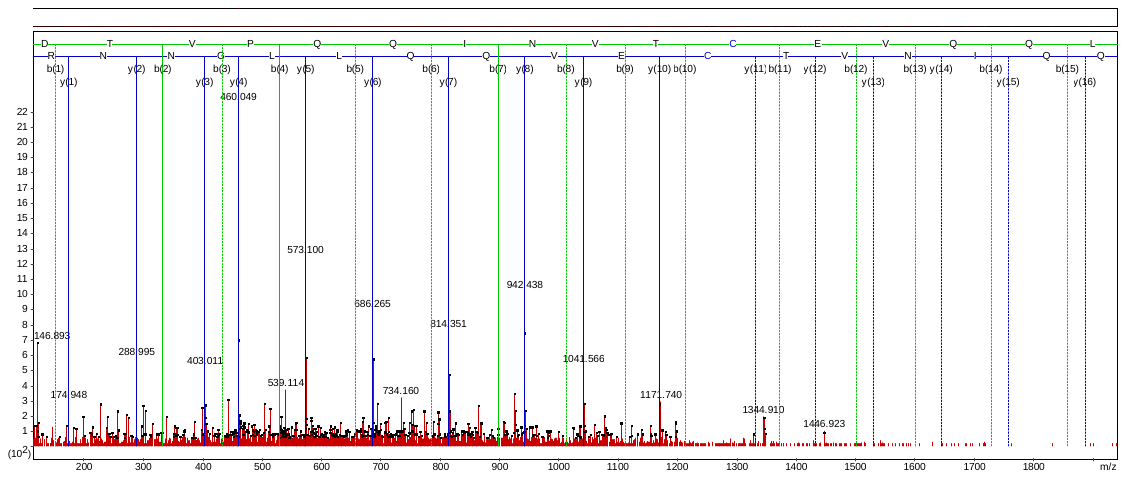

Accession No. ACF08646

AEGT_p_KELVEAK

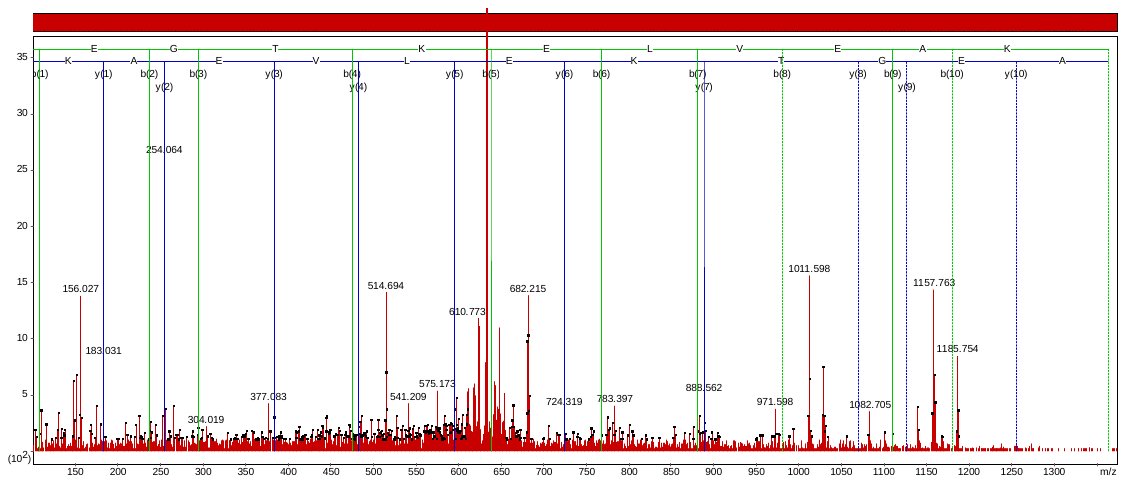

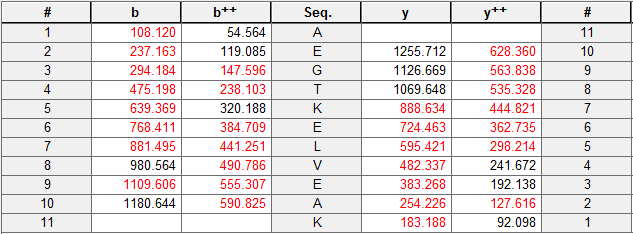

Accession No. EAY89475

AELGGVKDAS_p_EEVR

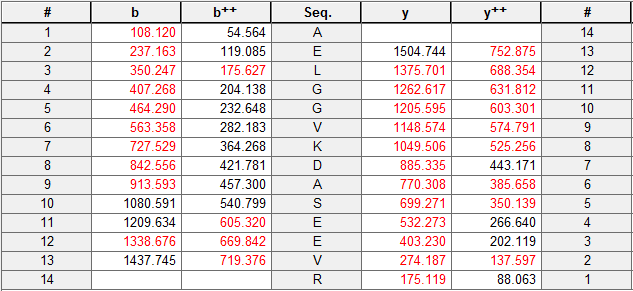

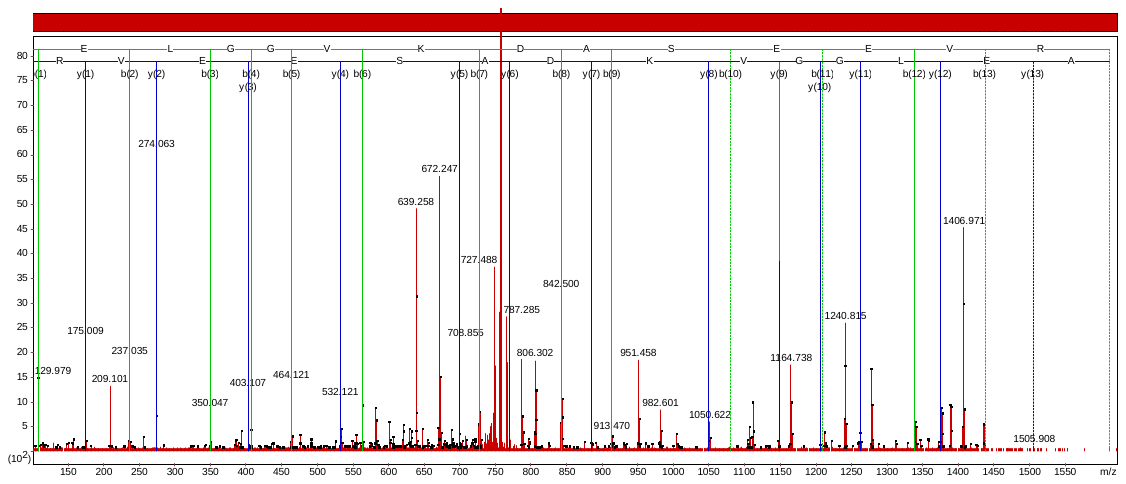

Accession No. AAQ14552

LVS_p_DDEDEQSK

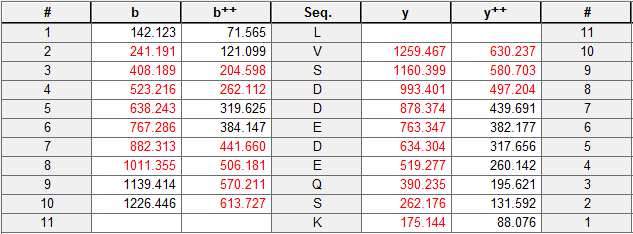

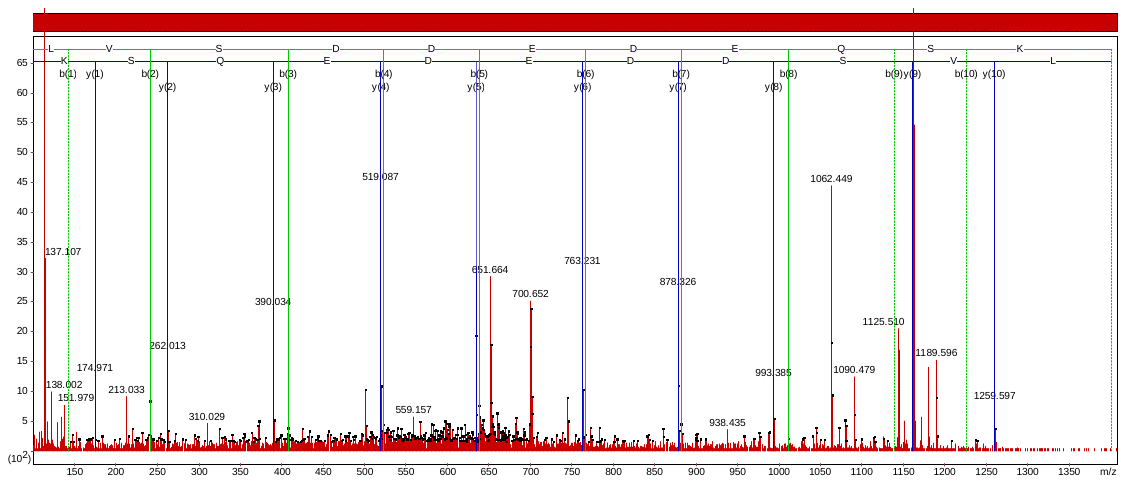

Accession No. Q8H1Y0

YHGHS_p_MSDPGSTYR

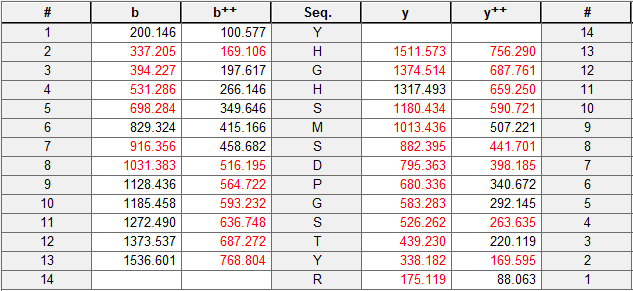

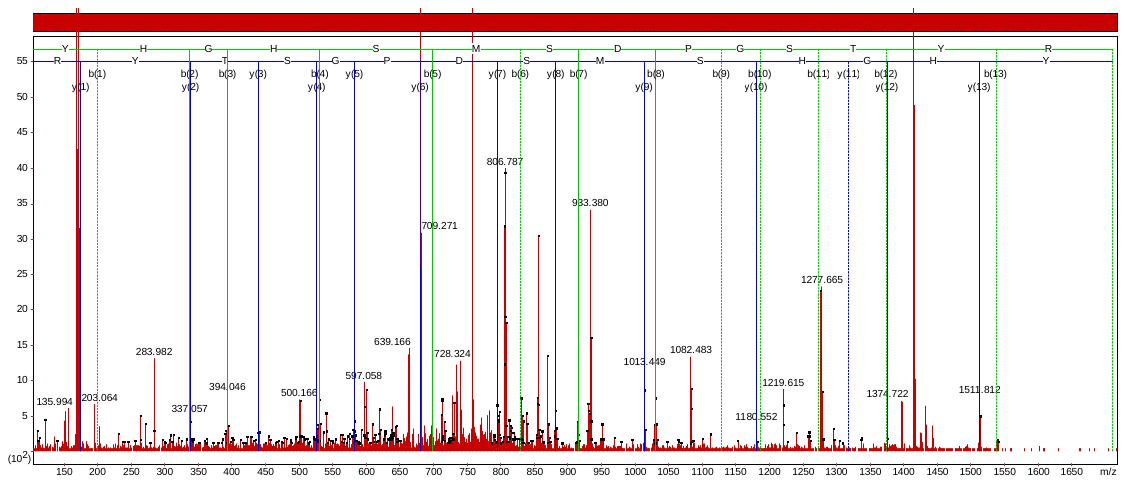

Accession No. BAK01943

QVAHAPQELNS_p_PR

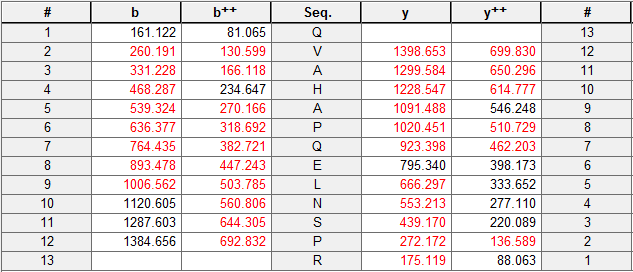

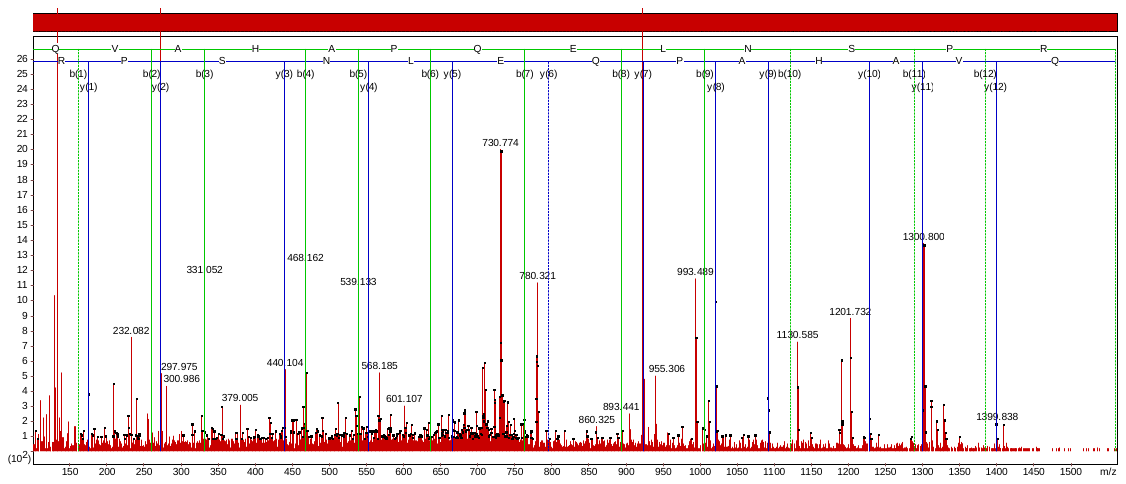

Accession No. BAJ06107

GFVPFVPGS_p_PVER

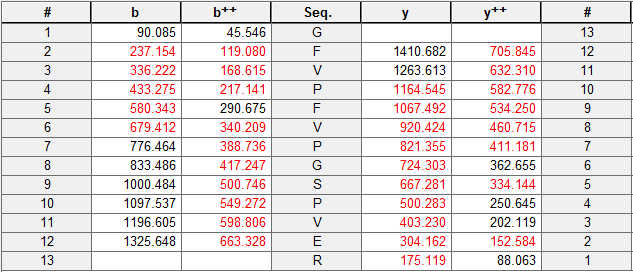

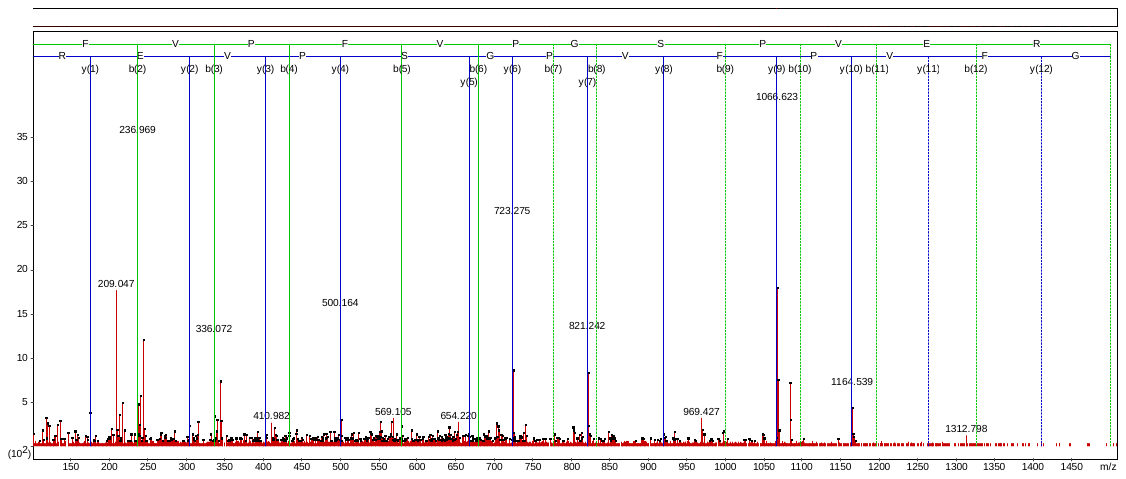

Accession No. EMT05628

AAAVAALSSVLTAEQSGS_p_SDNLR

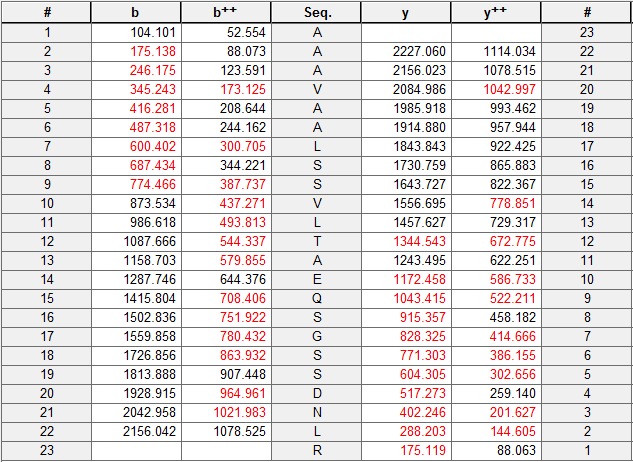

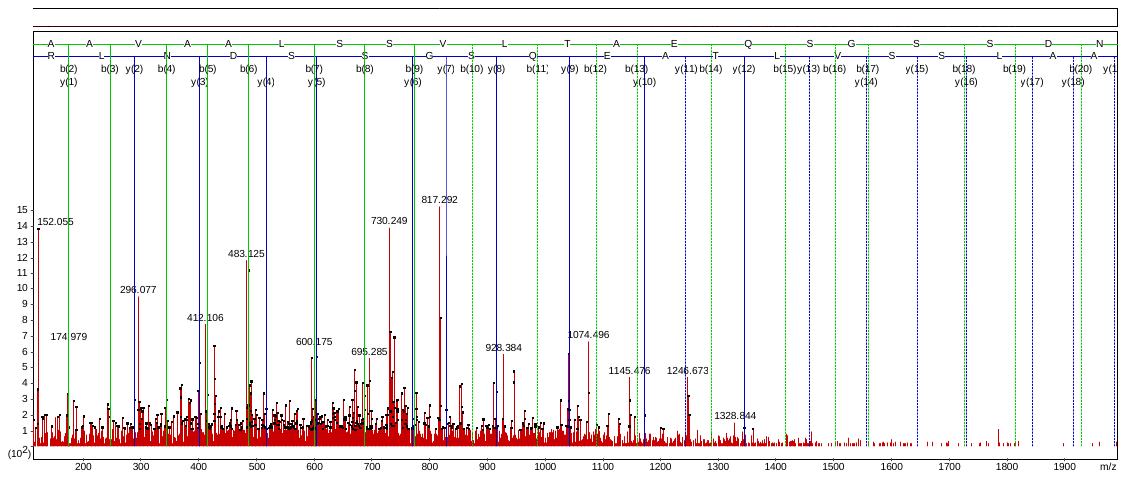

Accession No. BAK02001

ASSDDTSTSAAS_p_GDELVDDLK

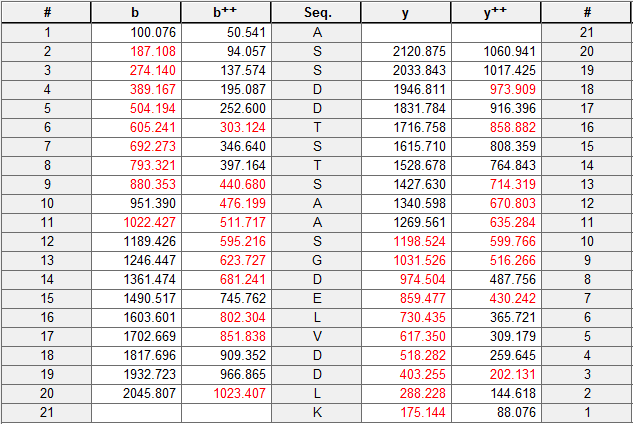

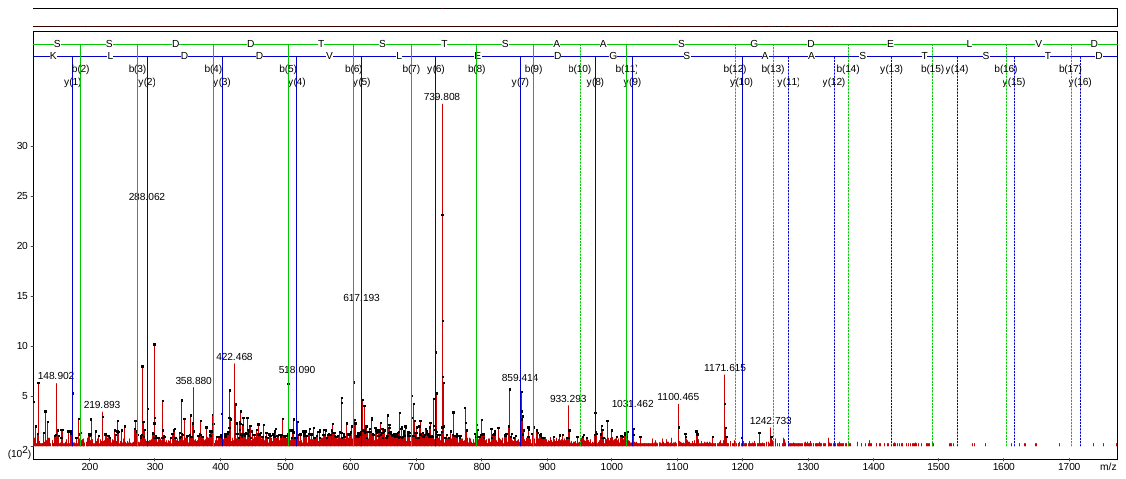

Accession No. XP_004965129

QSHS_p_DGSLDTMAR

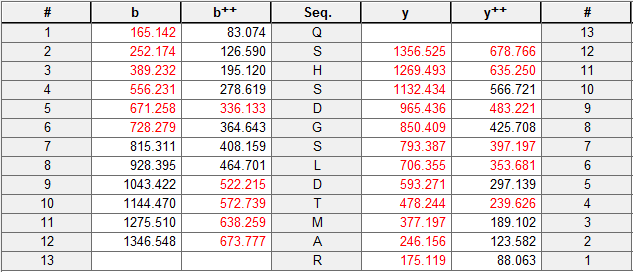

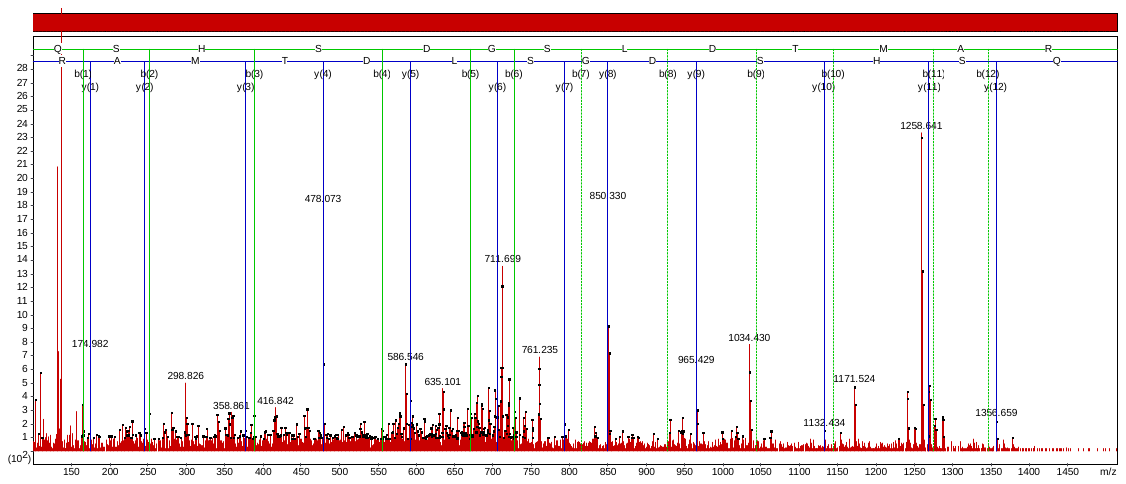

Accession No. EMS47290

VAEQLS_p_DDEGEDQSK

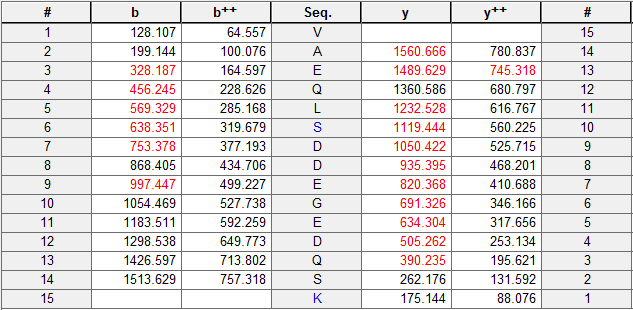

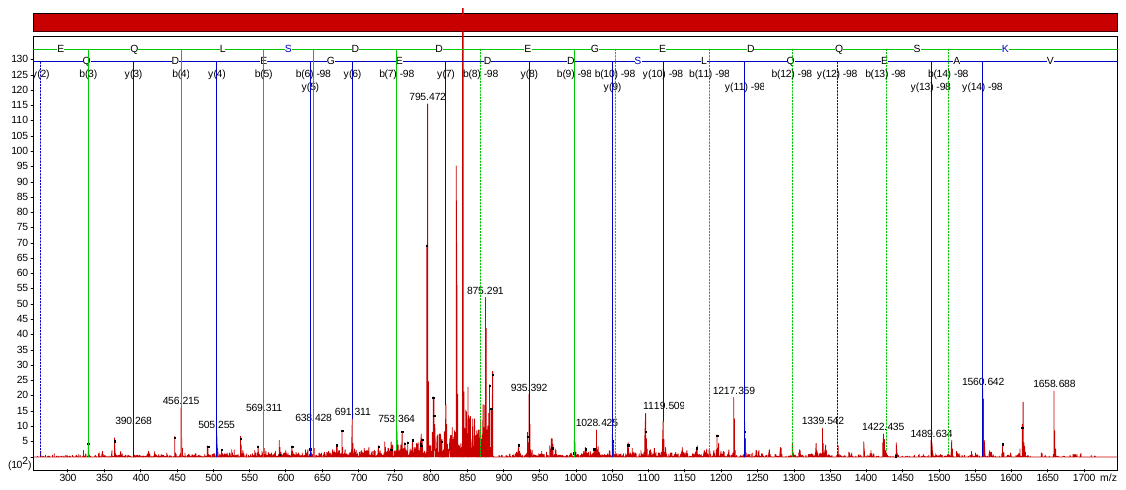

Accession No. EMT08603

SIS_p_ADGLNSLR

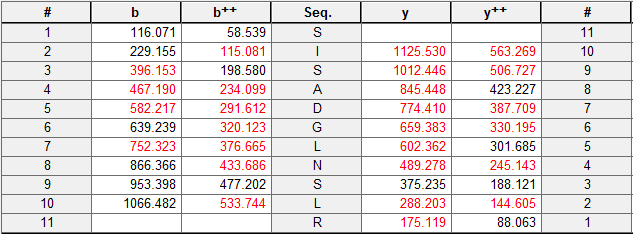

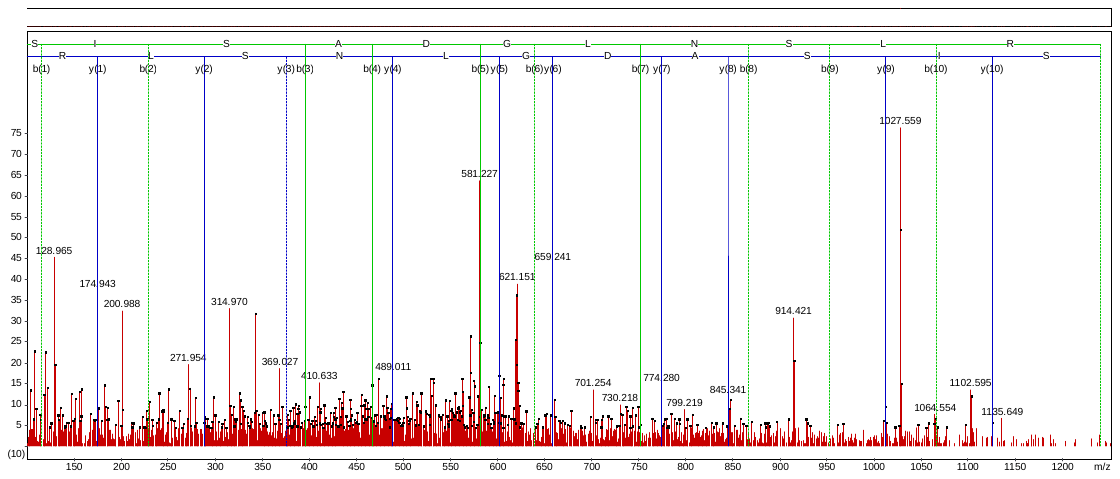

Accession No. AAS00828

KLLPGS_p_VDG

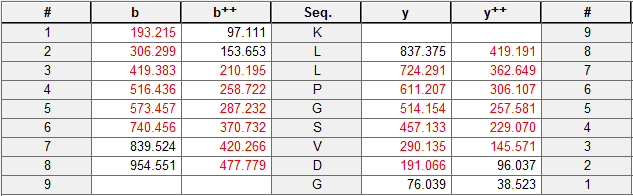

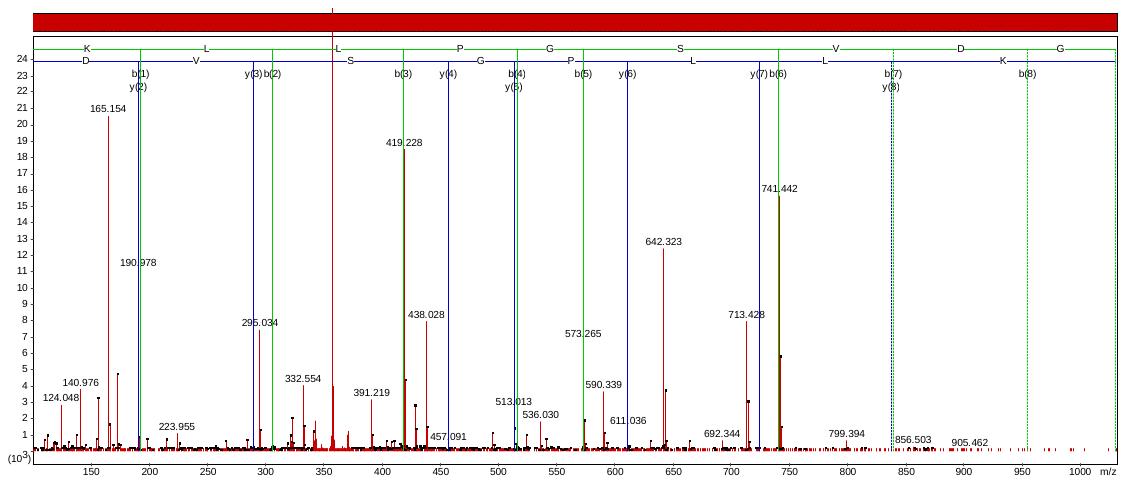

Accession No. EMT29522

GQDPQGMS_p_PGPGGR

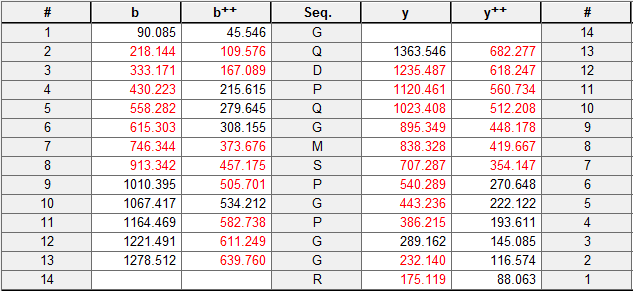

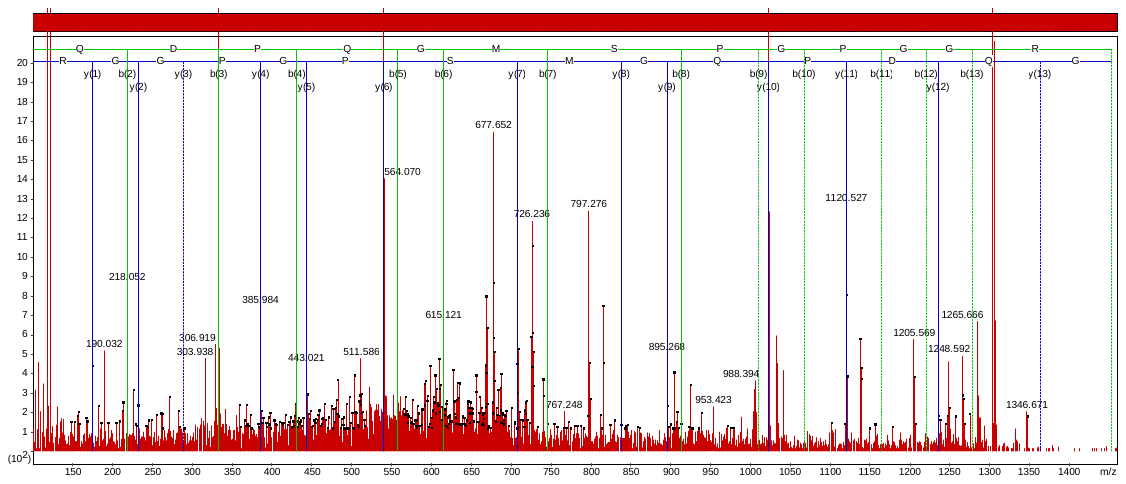

Accession No. EEC75374

SDSTASGDS_p_PKER

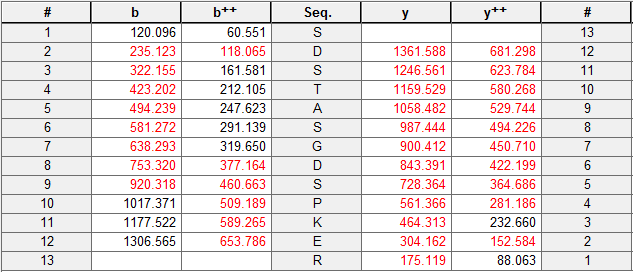

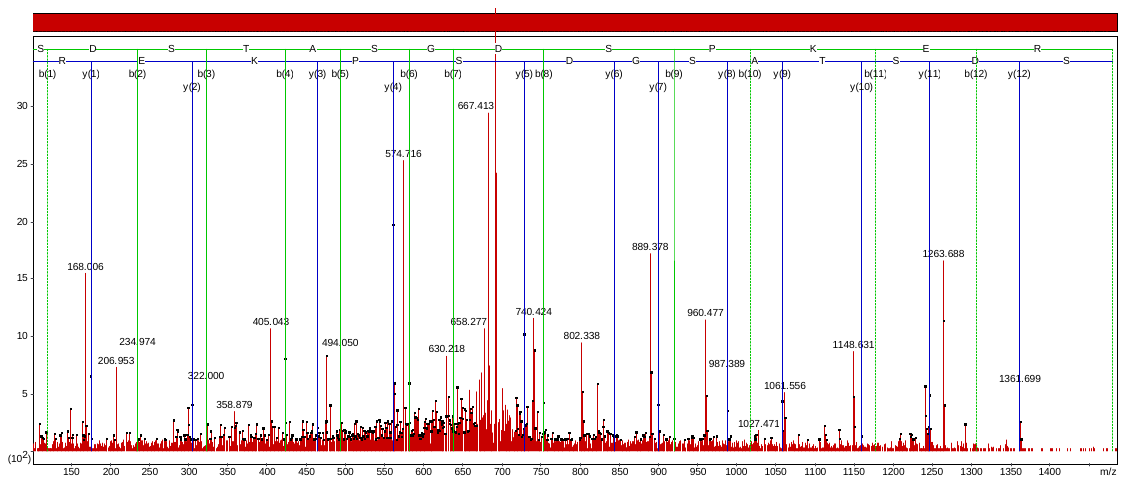

Accession No. CAN61094

PIDT_p_FIDVNIK

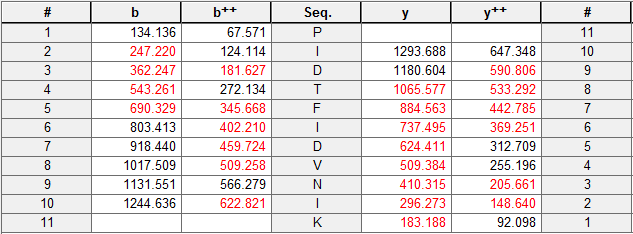

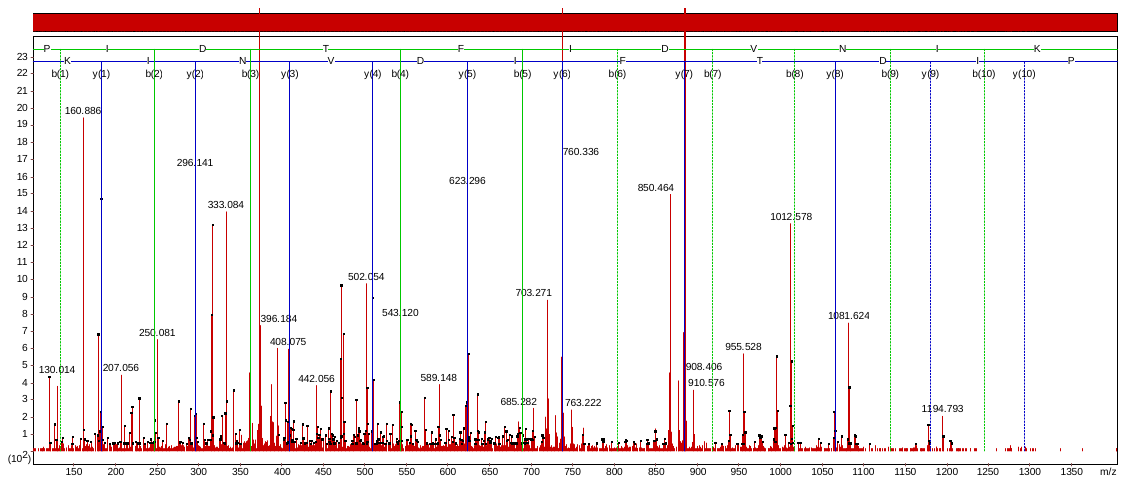

Accession No. BAJ99149

LQS_p_PGAQQYYGTSR

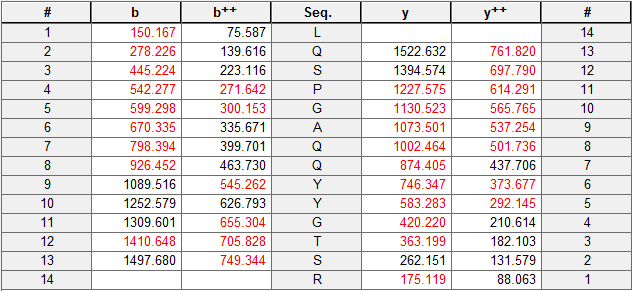

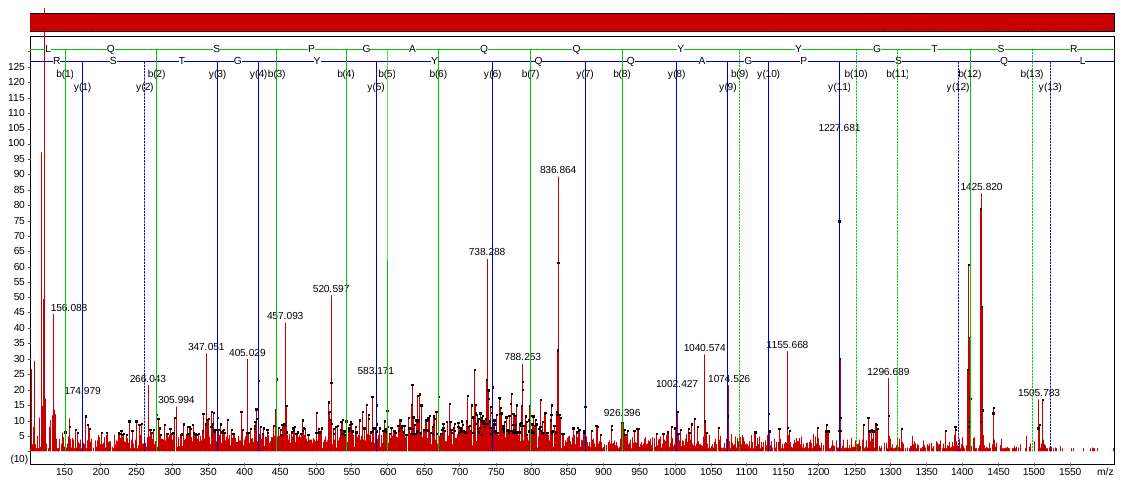

Accession No. XP_004980077

RGY_p_GGGGGGGGGGGGGGGGGGGGGGR

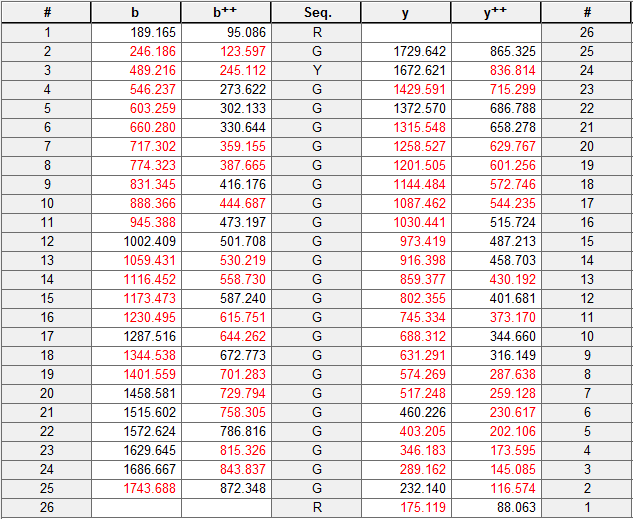

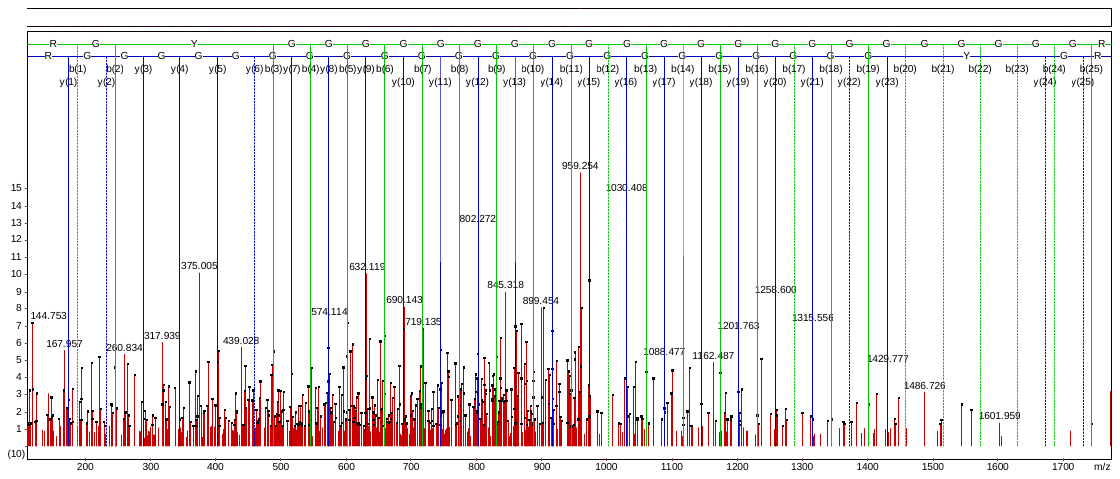

Accession No. ACO61321

MAS_p_AASSSTETAAPK

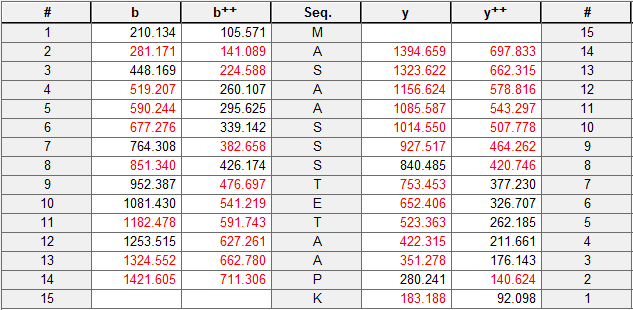

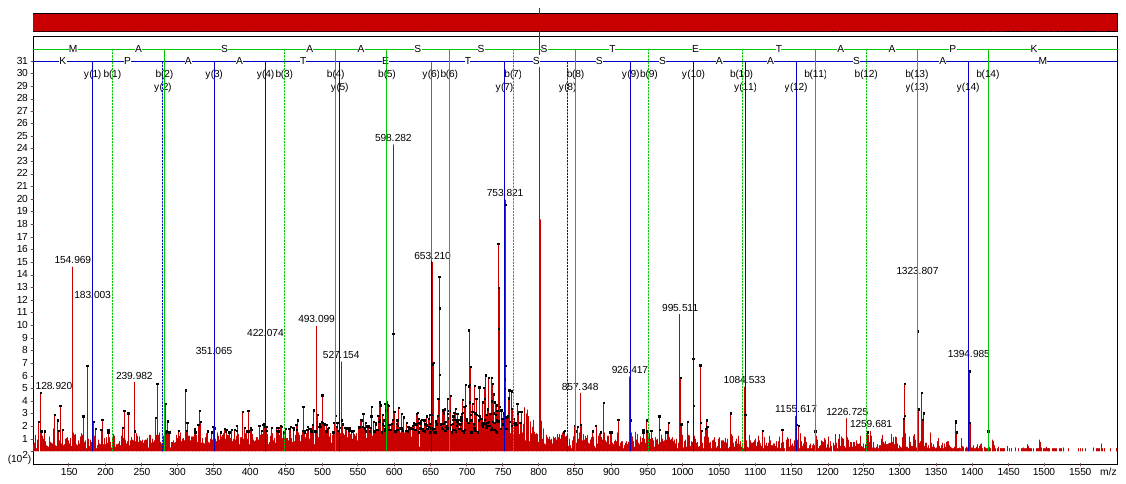

Accession No. EMT25919

QVS_p_VDVPDVR

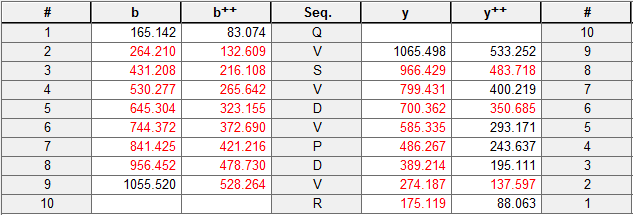

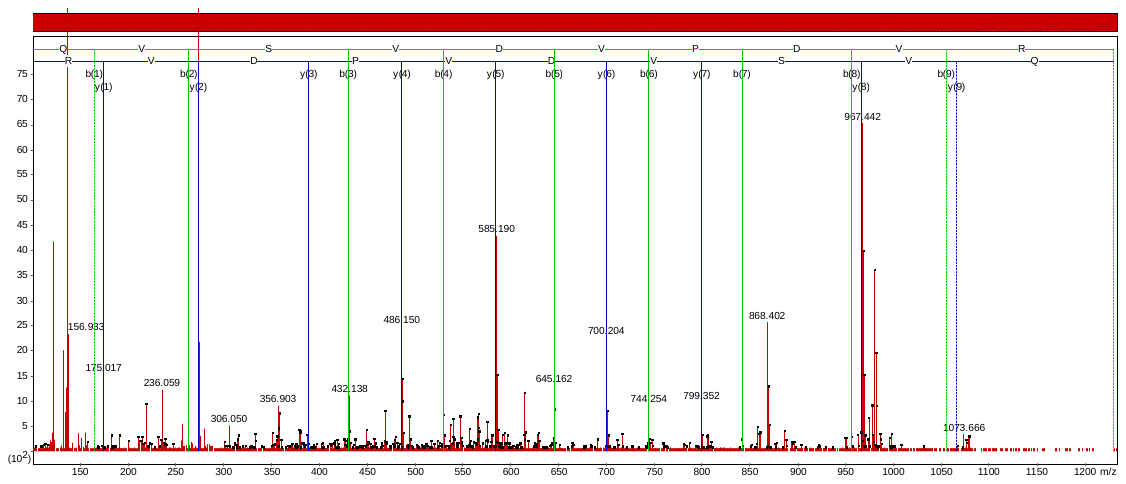

Accession No. EMS60685

TLDLTGVQPPS_p_PKPK

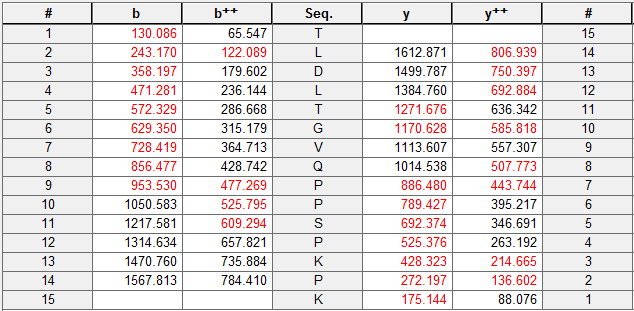

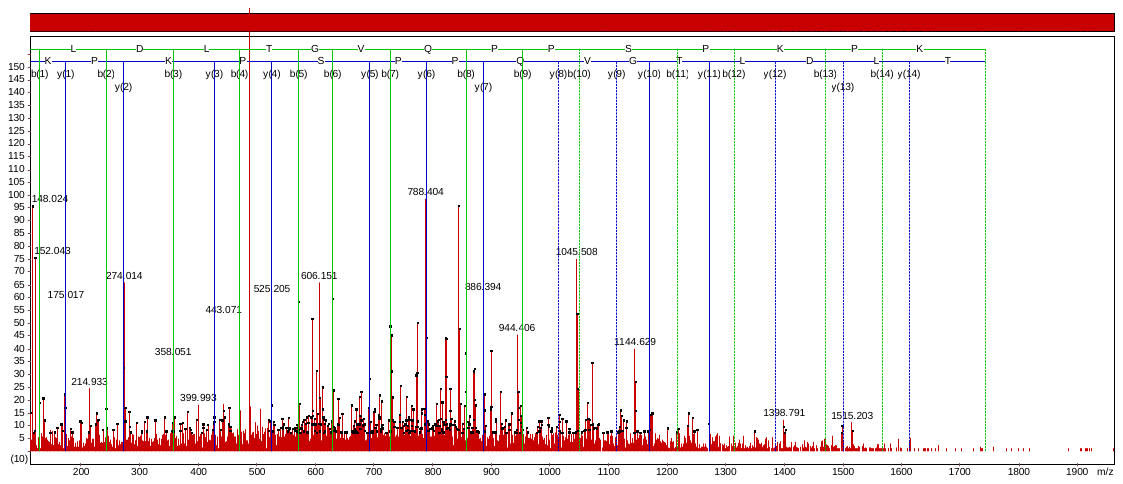

Accession No. CBF59389

RESLY_p_GSLSS_p_LEDDIVR

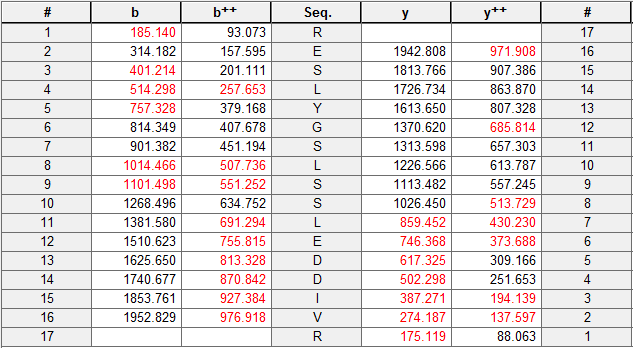

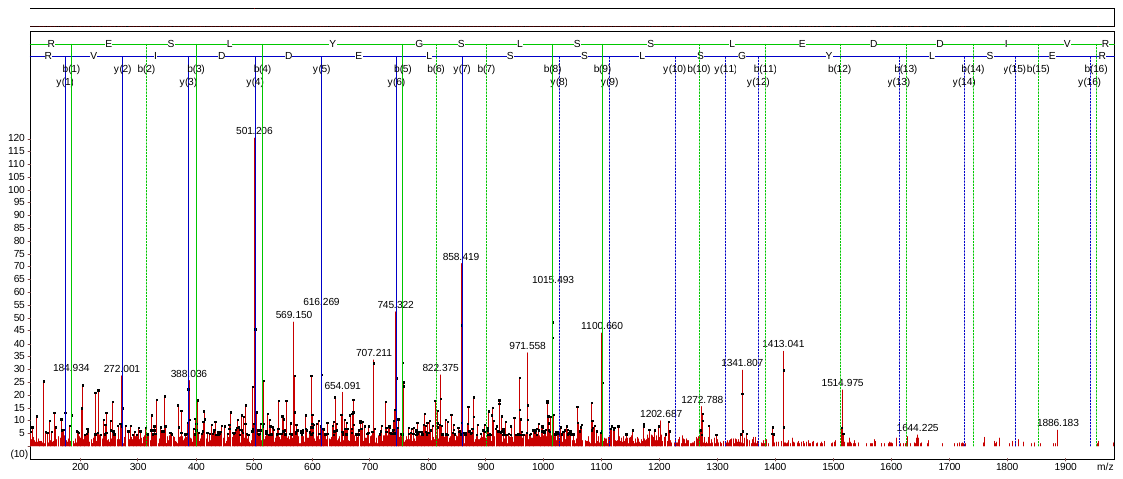

Accession No. DAA41082

NLFFFS_p_RLAGR

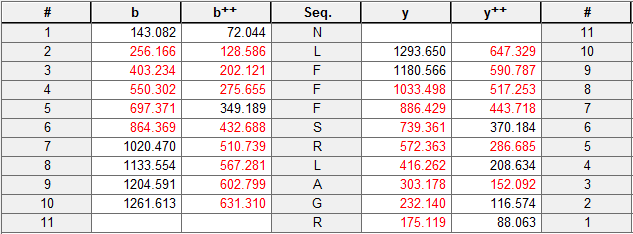

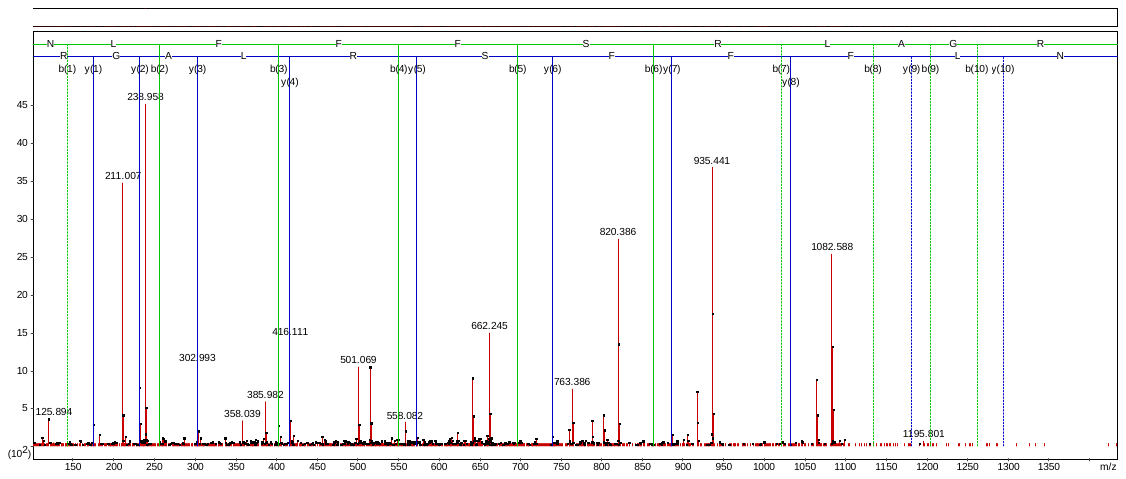

Accession No. XP_010104076

EEEPEQYWQT_p_AGER

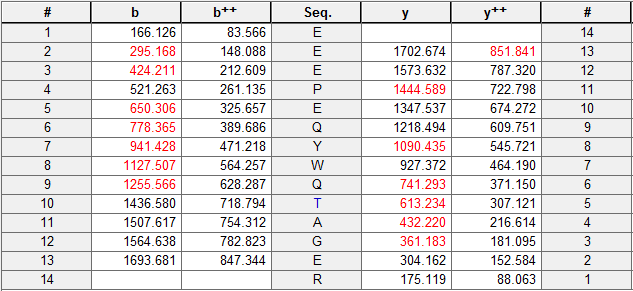

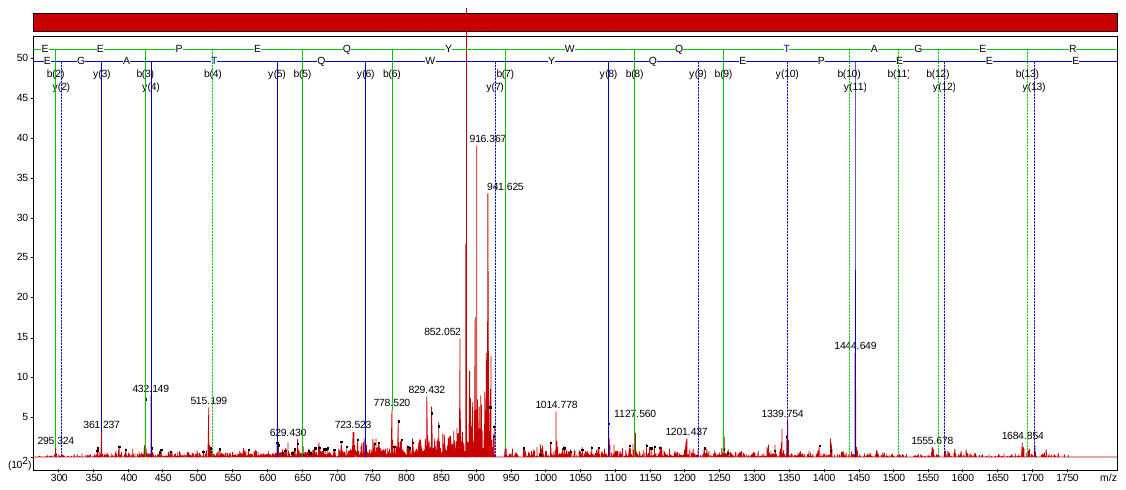

1. Leaf phosphoproteome

Accession No. CAA59049/ gi|666054

APERPIWFPGS[Pho]TPPPWLDGS[Pho]LPGDFGFDPWGLGSDPESLR

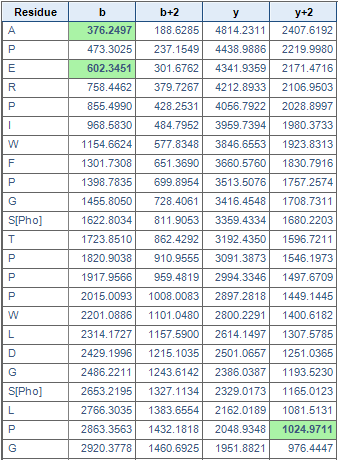

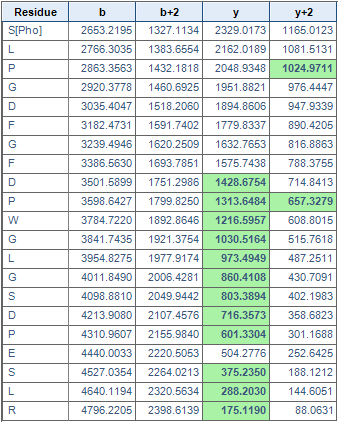

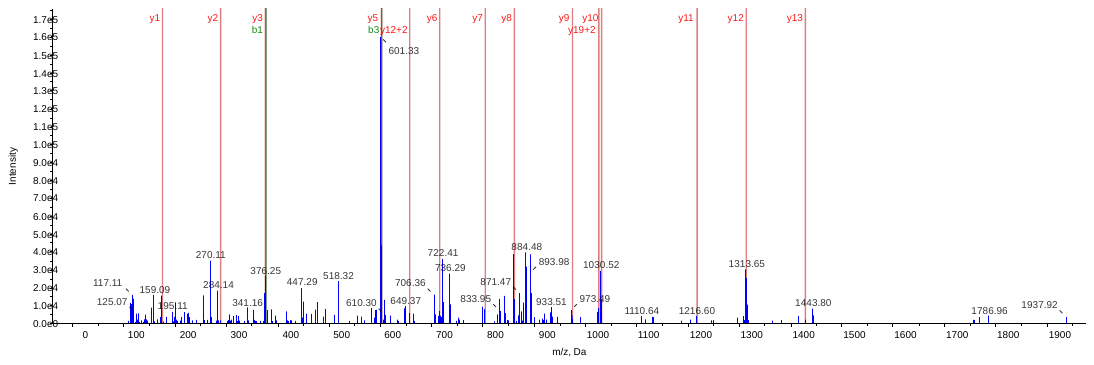

Accession No. XP_003618083/ gi|357495589

QSLS[Pho]YLDGSLPGDFGFDPLGLSDPEGTGGFIEPR

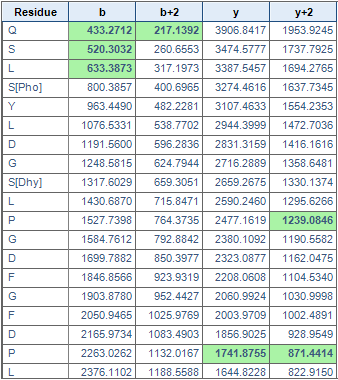

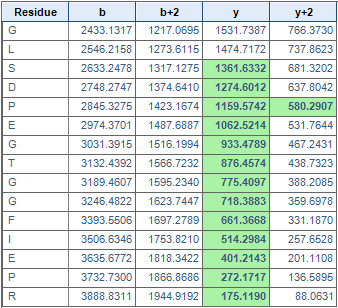

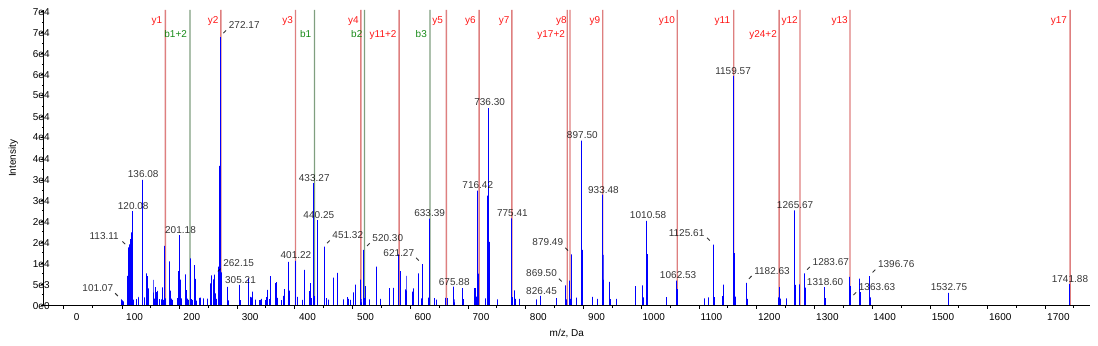

Accession No. CDI44335/ gi|550555057

VAGGPLGEVVDPLYPGGS[Pho]LDPLGLADDPEAFAELK

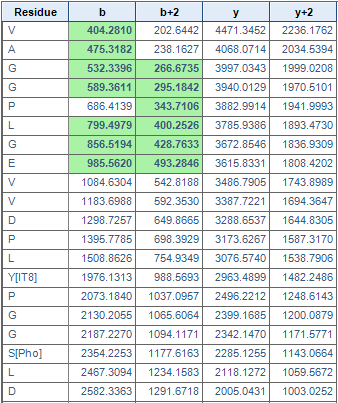

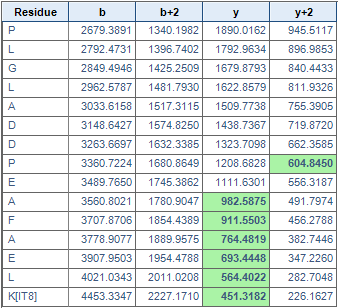

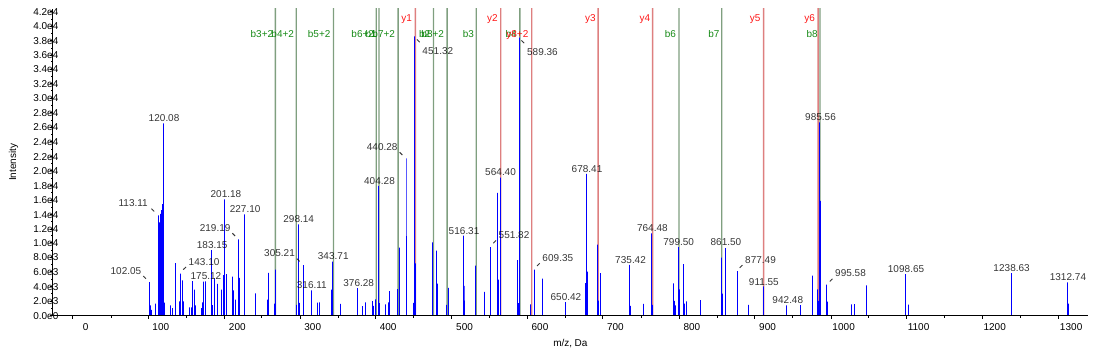

Accession No. EMS50795/gi|473952979

VLYLGPLSGEPPS[Pho]YLTGEFPGDYGWDTAGLSADPETFAK

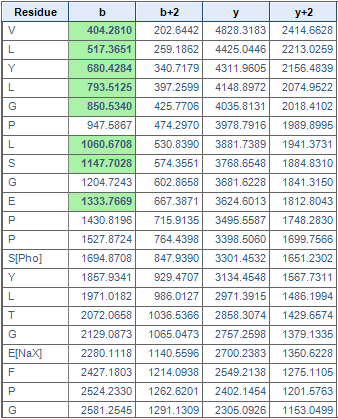

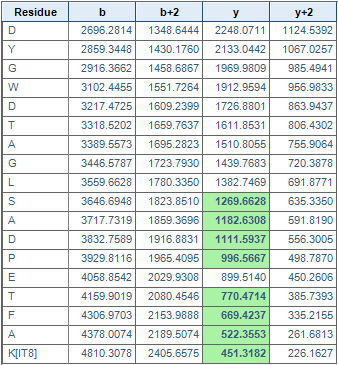

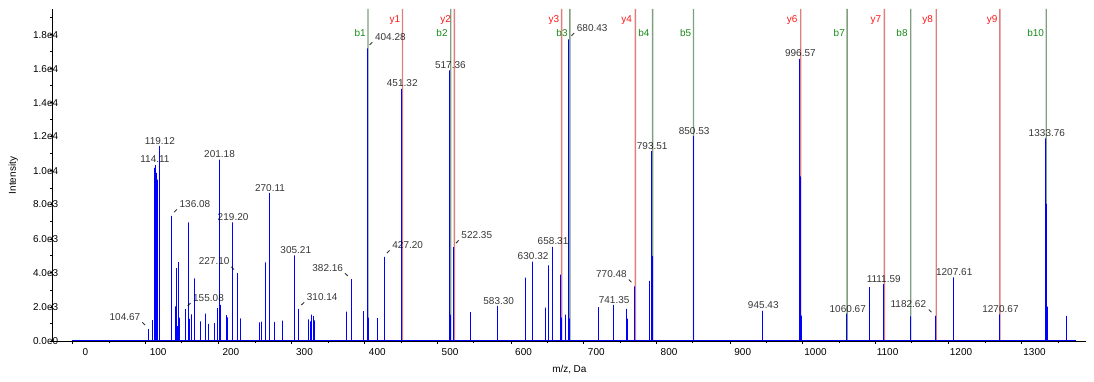

VLYLGPLSGEPPS[Pho]YLT[Pho]GEFPGDYGW[HKy]DT[Pho]AGLSADPETFAK[IT8]

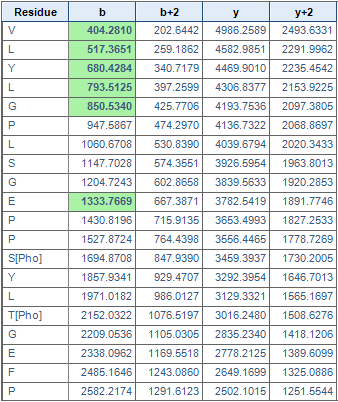

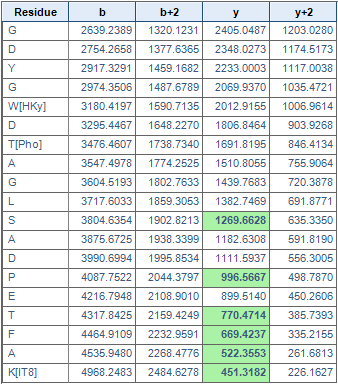

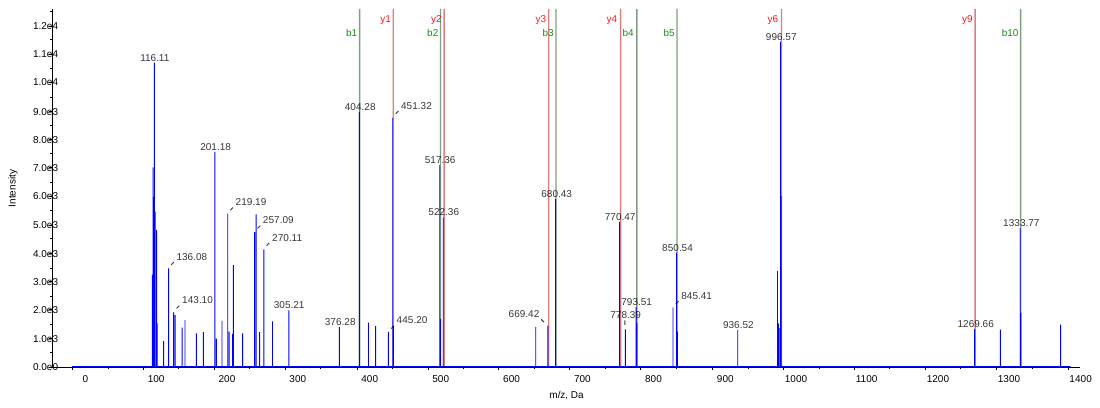

Accession No. EMT11232/ gi|475549279

VLYLGPLS[Pho]GDPPS[Pho]YLTGEFPGDYGWDT[Pho]AGLSADPETFAK

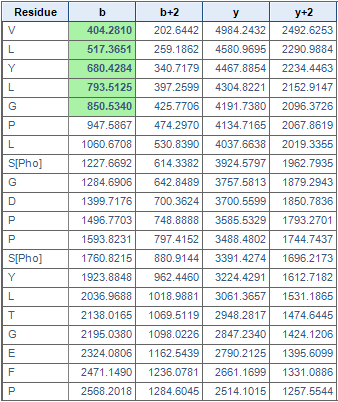

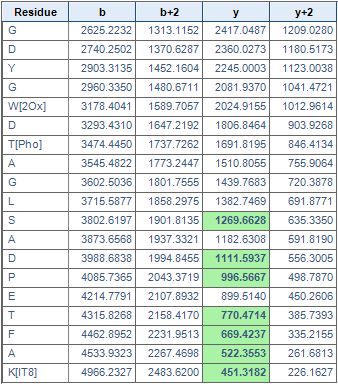

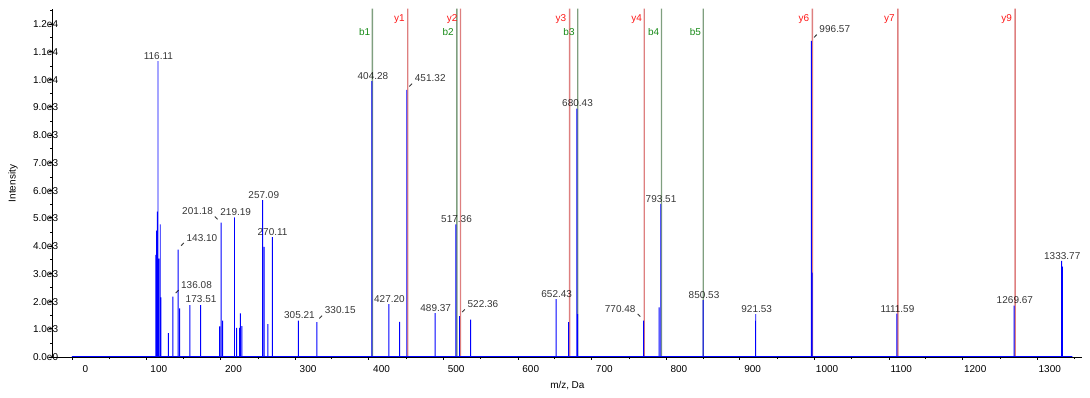

Accession No. EMT29003/ gi|475614191

VLYLGPLSGEPPS[Pho]YLNGEFPGDYGWDT[Pho]AGLS[Pho]ADPETFAK

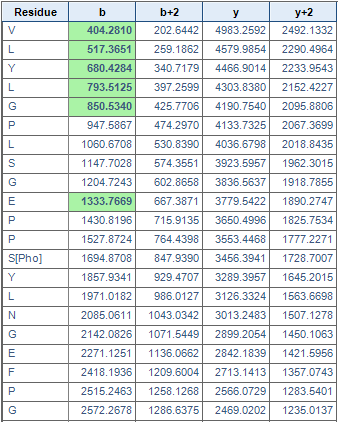

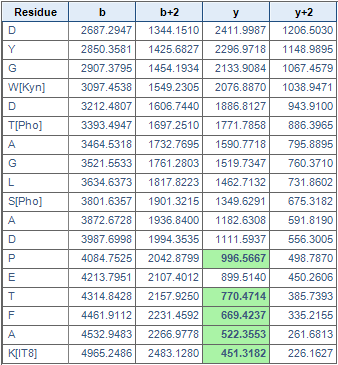

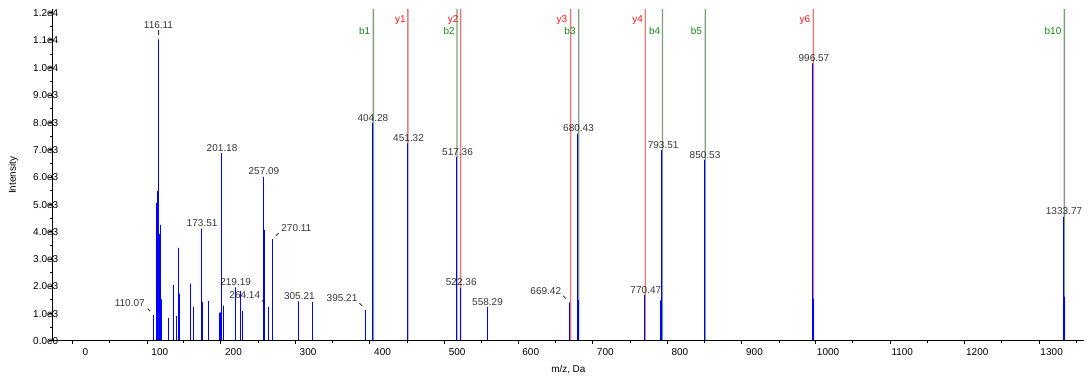

VLYLGPLSGEPPSYLNGEFPGDYGWDTAGLS[Pho]ADPETFAK

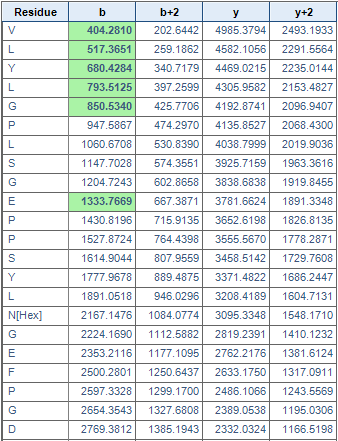

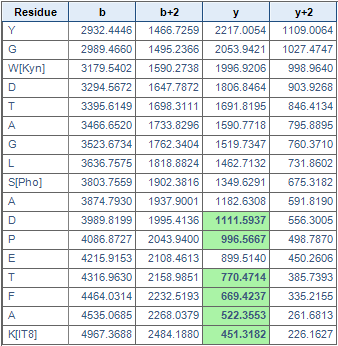

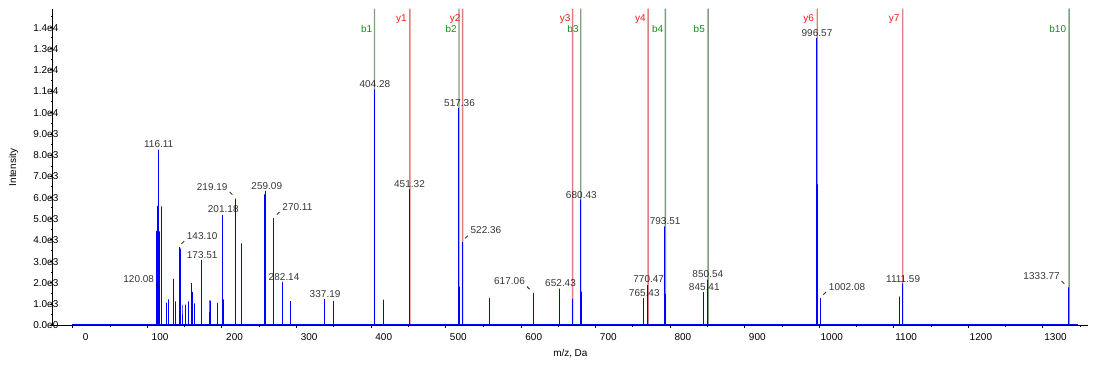

Accession No. CAA32109/gi|20182

VLYLGPLSGREPPS[Pho]YLTGEFPGDYGWDTAGLSADPETFAK

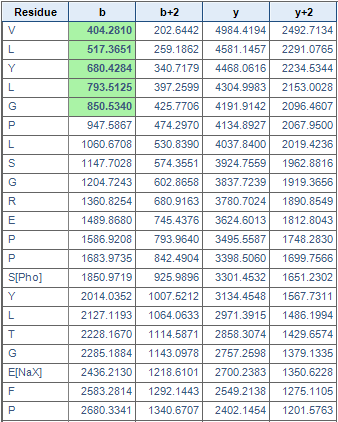

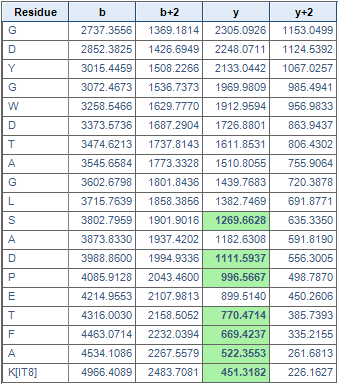

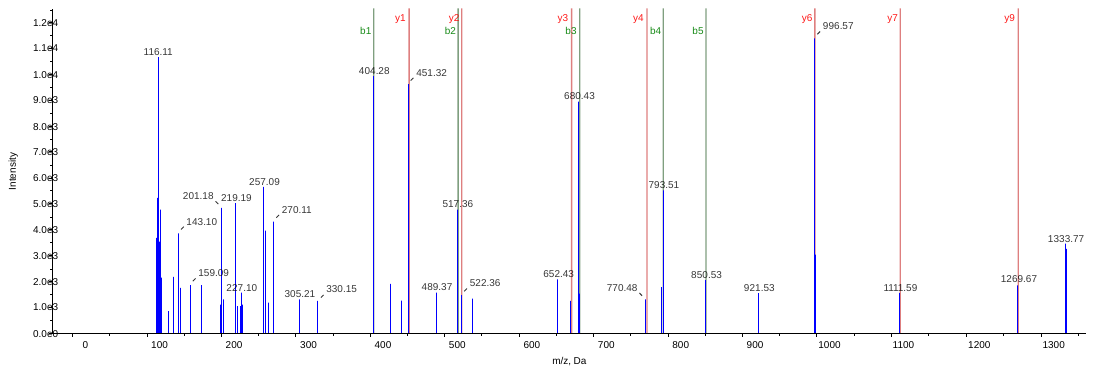

Accession No. 1908421A/ gi|445116

LAQNLAGEIIGT[Pho]RFEDADVK

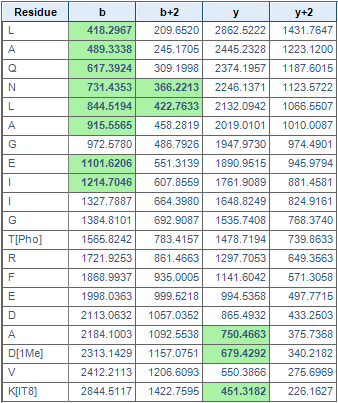

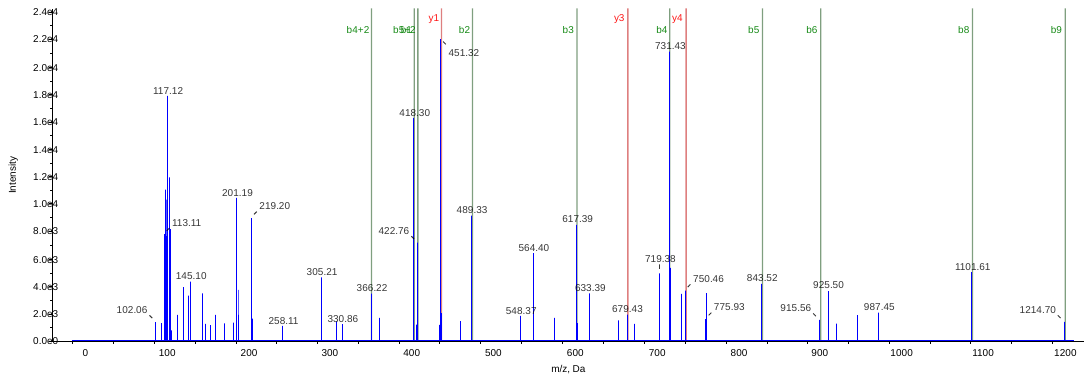

Accession No. CDI44415/gi|550554240

PAEYLQYDVDSLDQNLAQNLAGEIIGT[Pho]R

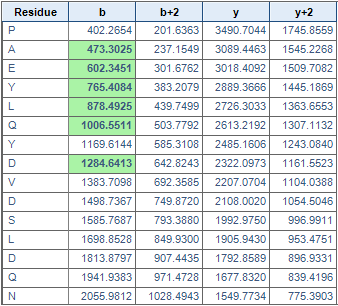

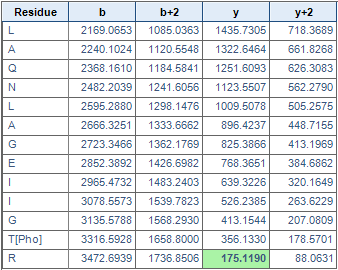

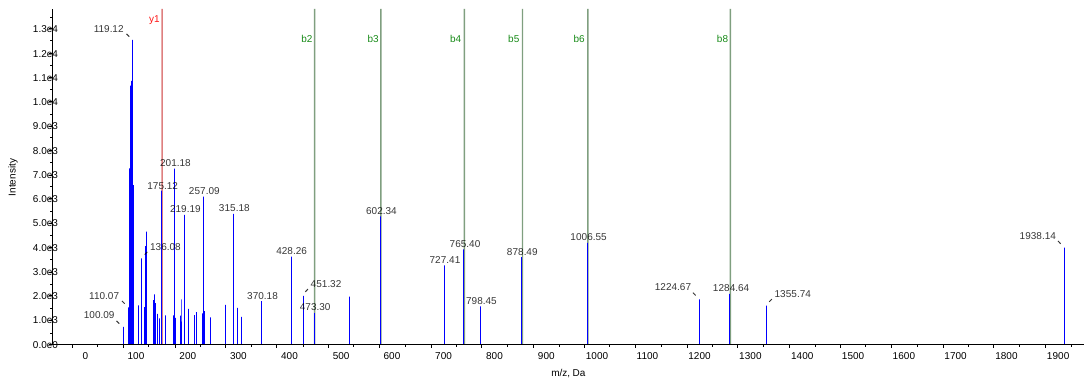

Accession No. XP_003562892/gi|357122377

PAEYLQYDPDS[Pho]LDQNLAQNLAGEVIGTRFEDADIK

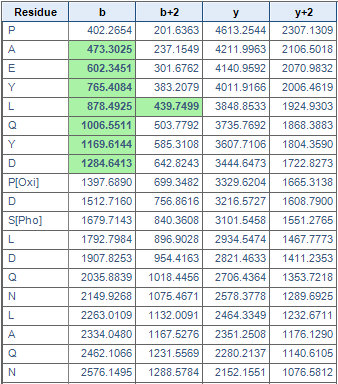

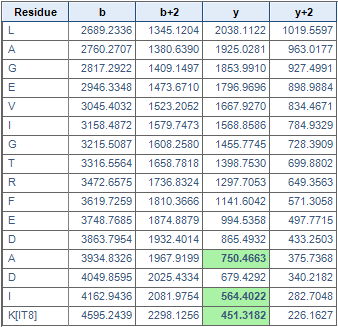

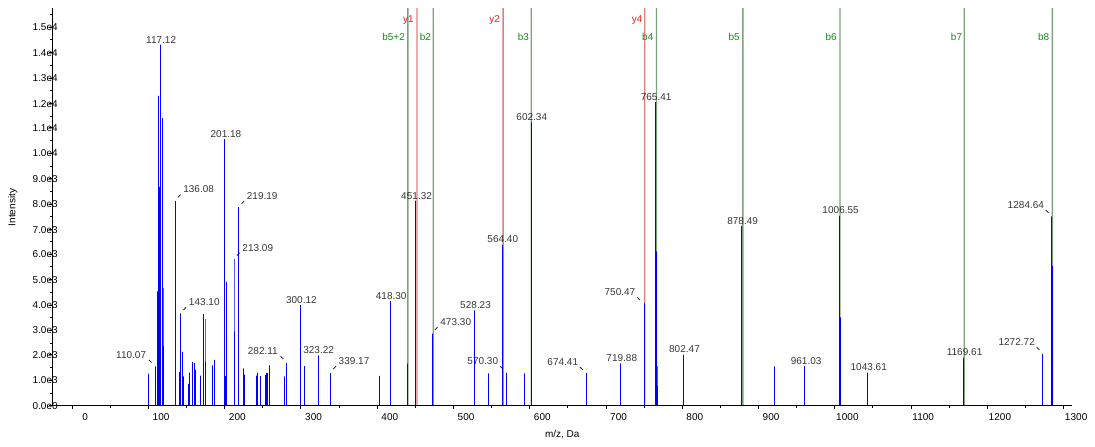

Accession No. EMT03206/gi|475485989

TGALLLDGNT[Pho]LNYFGNS[Pho]IPINLILAVVAEVVLVGGAEYYR

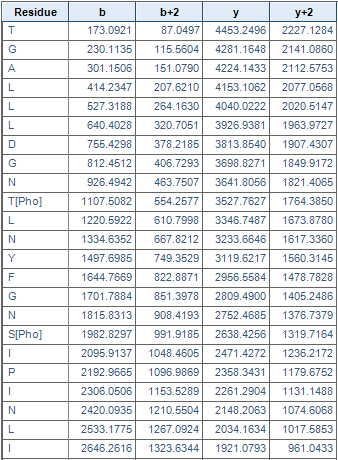

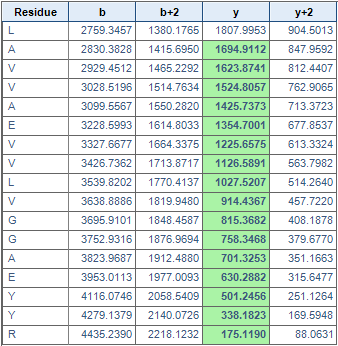

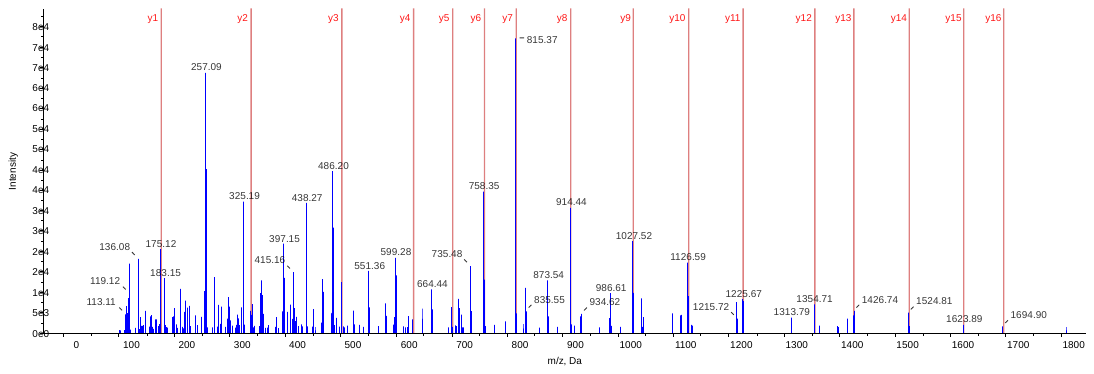

TGALLLDGNTLNYFGNS[Pho]IPINLILAVVAEVVLVGGAEYYR

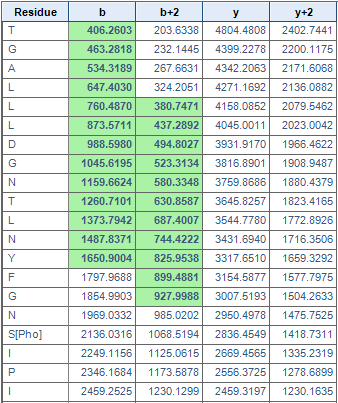

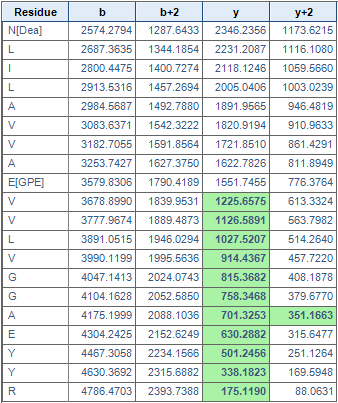

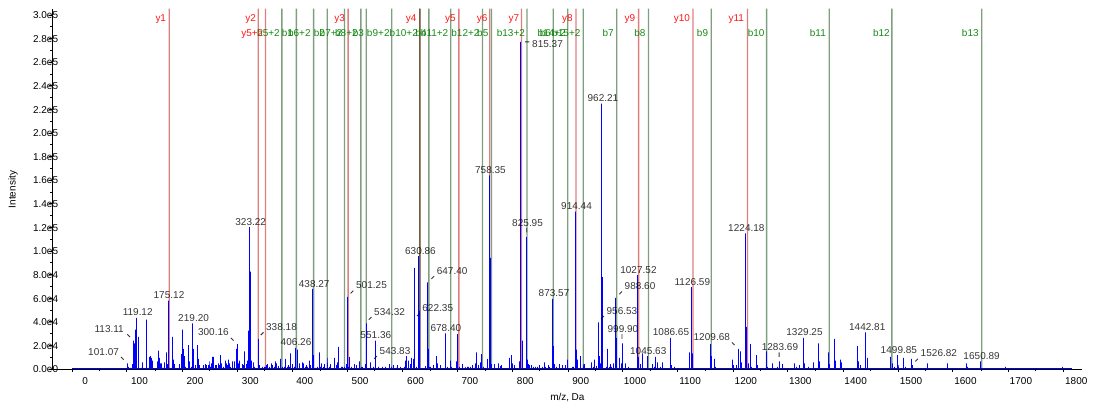

Accession No. XP_003564708/gi|357126063

GILS[Pho]QLNLETGIPIYEAEPLLLFFILFTLLGAIGALGDR

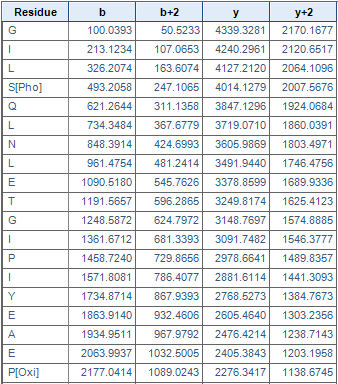

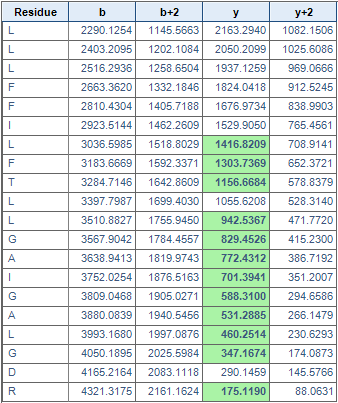

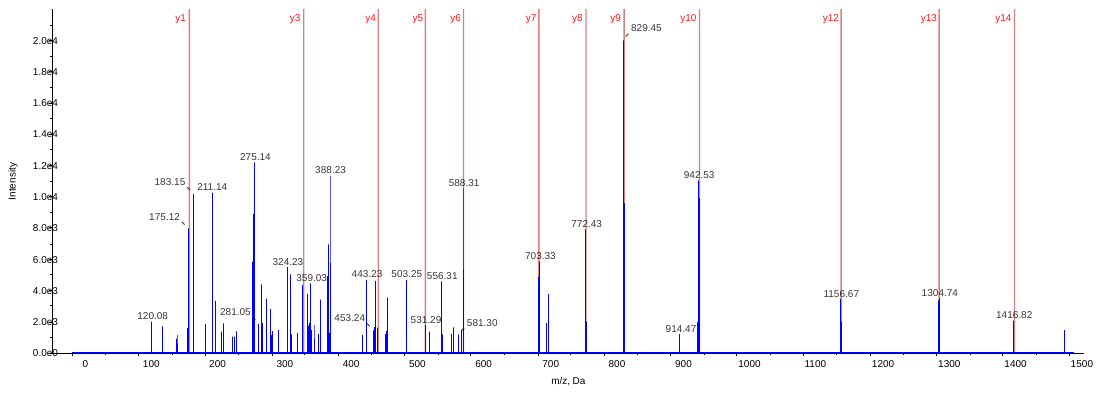

Accession No. ABC02751/gi|83317154

TLFNGT[Pho]FVLAGR

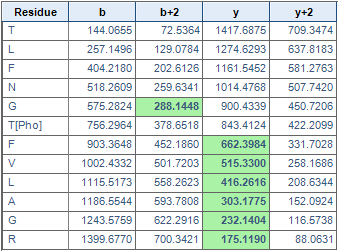

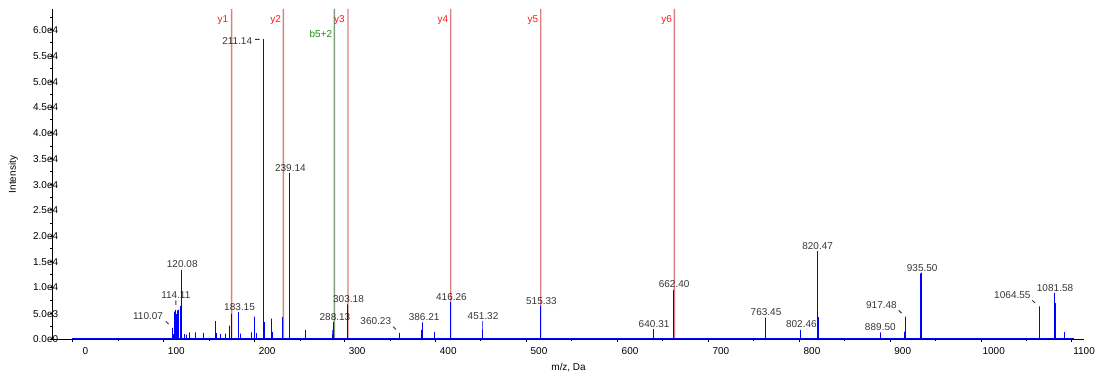

Accession No. P69555/gi|61230121

AT[Pho]QTVEDSSKPR

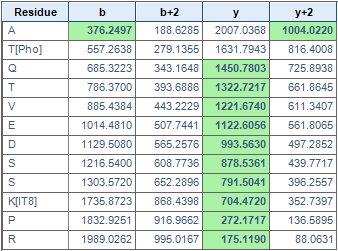

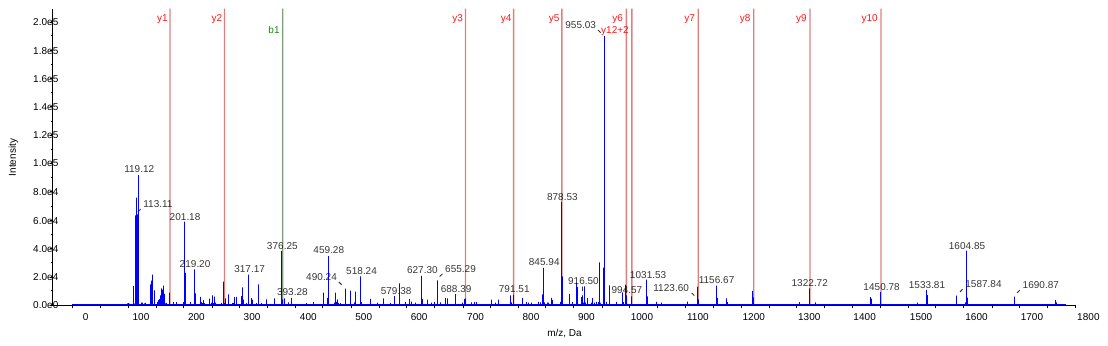

AT[Pho]QT[Pho]VEDSSKPRPK

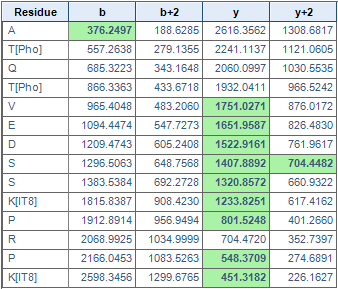

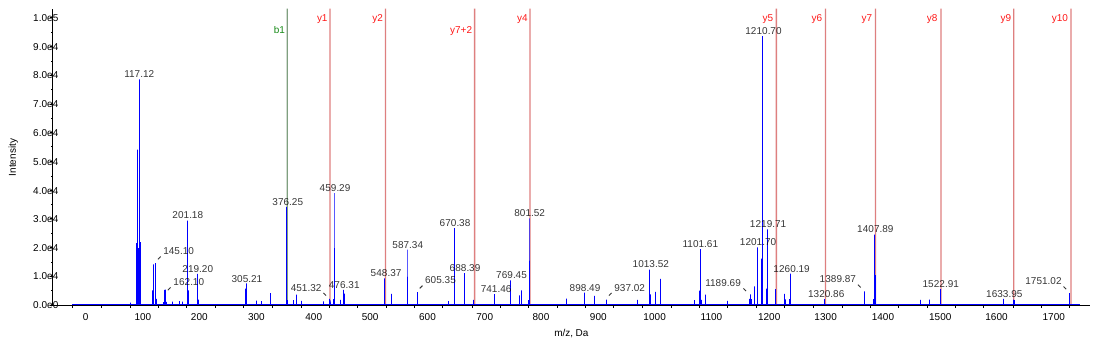

Accession No. CCO16140/gi|412991295

GGSTGYDNAVALPARS[Pho]DADDLQKENNK

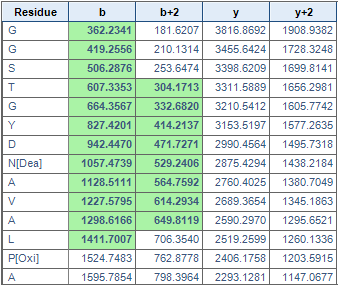

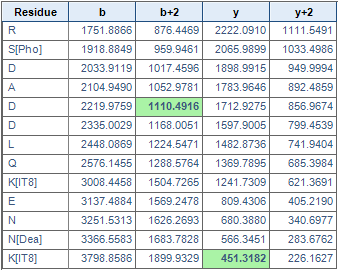

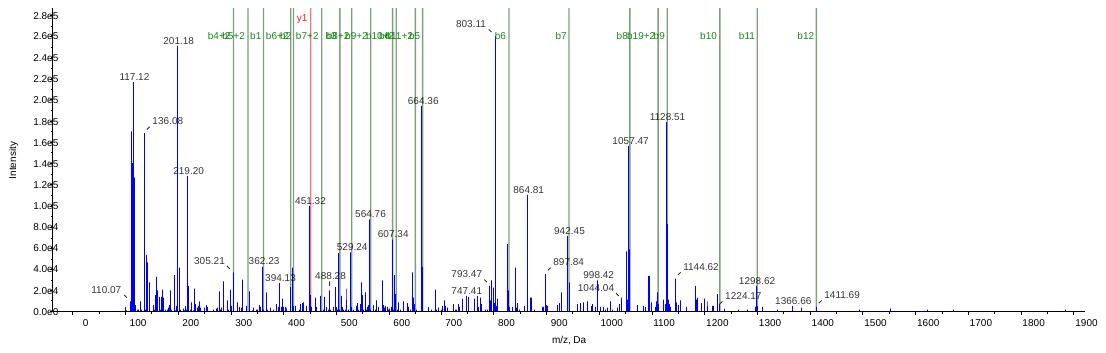

Accession No. BAJ97488/gi|326528933

VDGPAPSAGGT[Pho]ASR

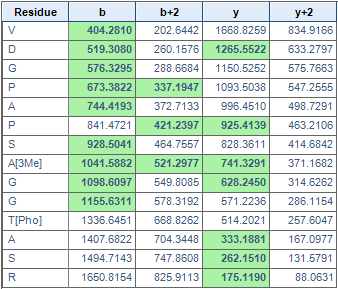

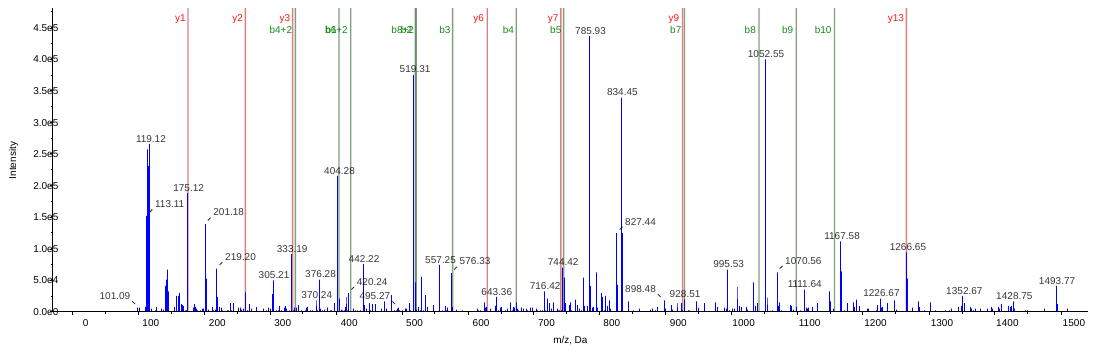

Accession No. ABA95524/gi|77552727

GLAYDIS[Pho]DDQQDITR

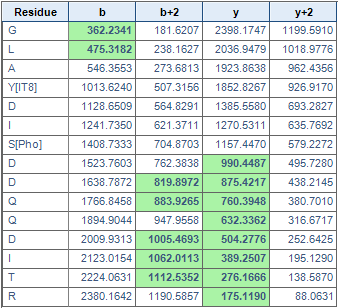

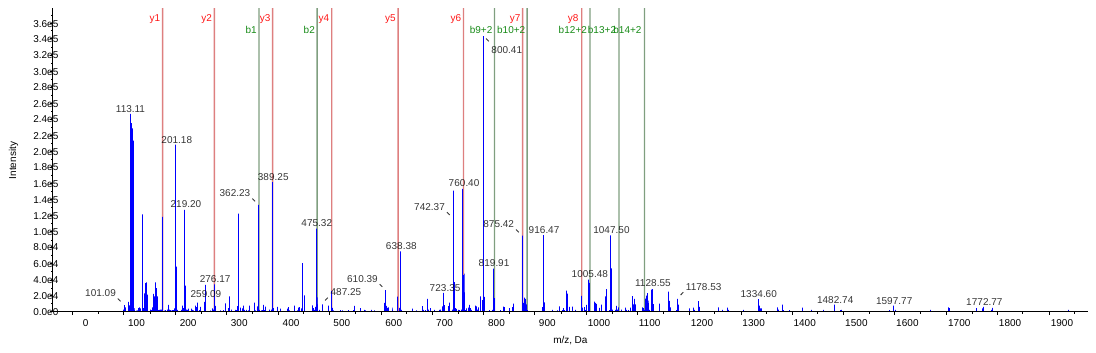

Accession No. ABB03412/gi|77798470

GLDFT[Pho]KD[Pho]DENVNSQPFMR

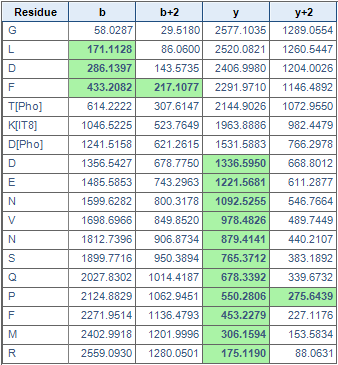

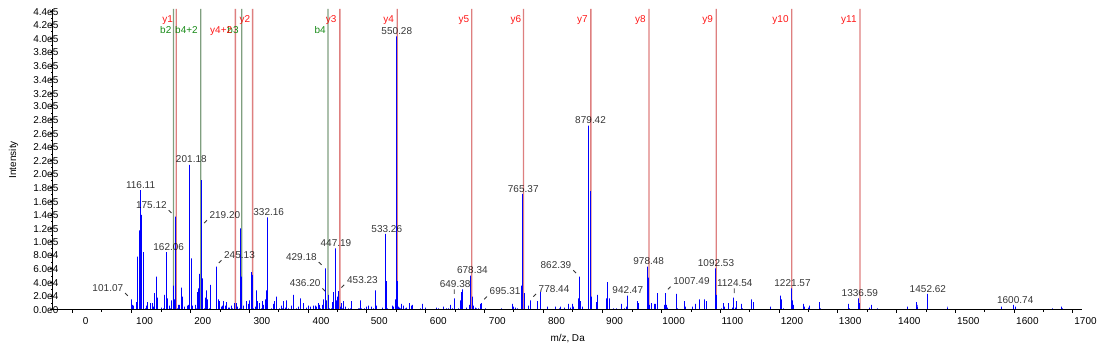

VT[Pho]PQPGVPPEEAGAAESSTGTWTTVWTDGLTSLDR

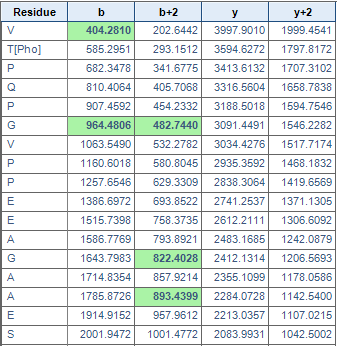

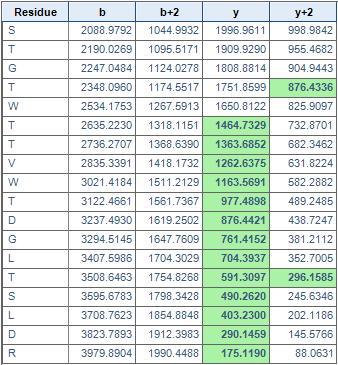

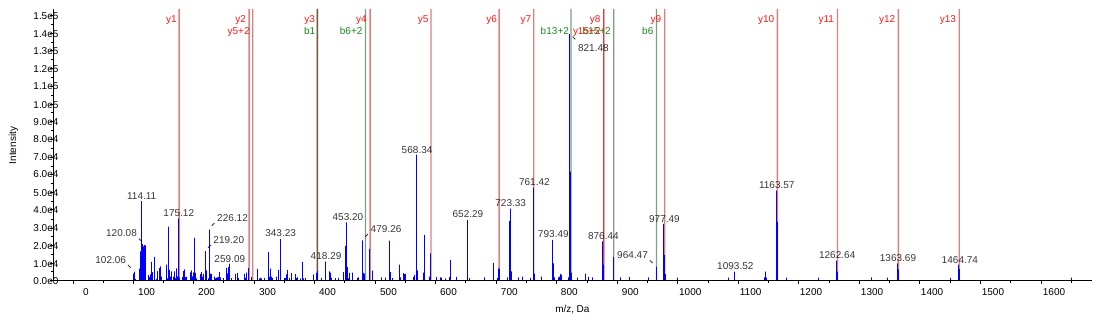

Accession No. CAB85674/gi|7414457

AAFRVTPRPGVPPEEAGAAVAAESS[Pho]TGTWTTVWTDGLTSLDR

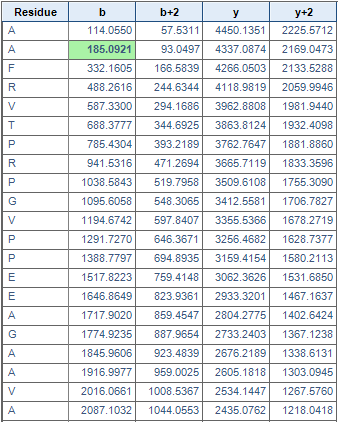

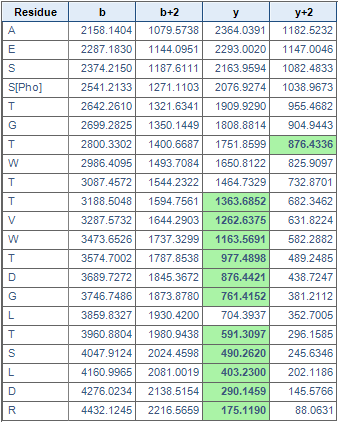

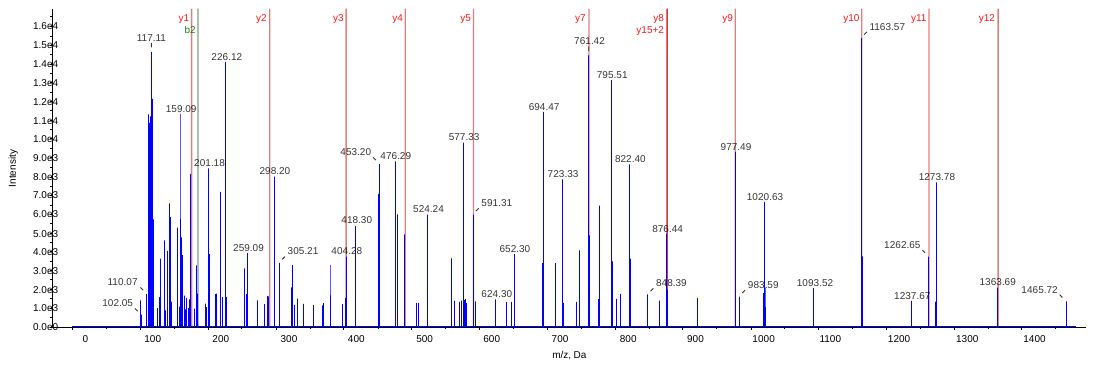

Accession No. CAA94018/gi|1771824

PGVPPEEAGAEVAAESS[Pho]TGTWTTVWTDGLTSLDR

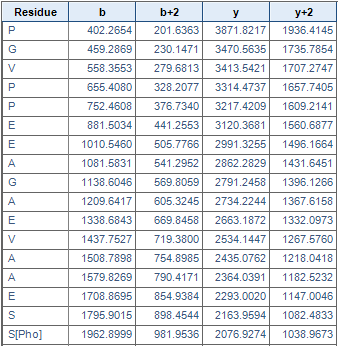

Accession No. CAC04358/gi|9909841

VTPQPGVPAEEAGAAVDAES[Pho]STGTWTTVWTDGLTSLDR

Accession No. AAG43944/gi|12004215

VT[Pho]PQPGVPPGGAGAAVAAESS[Pho]TGTWTTVWTDGLTSLDR

VTPQPGVPPGGAGAAVAAES[Pho]S[Pho]TGTWTTVWTDGLTSLDR

Accession No. CAA93205/gi|1518392

VTPQPGVPAEEAGAAVAAES[Pho]STGTWT[Pho]TVWTDGLTSLDR

VTPQPGVPAEEAGAAVAAESS[Pho]TGTWTTVWTDGLTSLDR

Accession No. AFA27686/gi|374975191

VTPQPGVPPEEAGAAVAGEIGT[Pho]WT[Pho]TVWTDGLTSLDR

VTPQPGVPPEEAGAAVAGEIGTWT[Pho]T[Pho]VWTDGLTSLDR

Accession No. EMT31124/gi|475620495

GDSS[Pho]PLEVIAINDTGGVK

Accession No. XP_003568189/gi|357133147

PGVVALDEAVTVGSVT[Pho]

Accession No. EMS64844/gi|474384779

NPLIAAAS[Pho]VIAAGLAVGLAS[Pho]IGPGVGQGTAAGQAVEGIAR

Accession No. AEJ10071/gi|338825698

T[Pho]GLGQVMSGELVEFAEGTR

Accession No. ABR67212/gi|150035733

GFQLILS[Pho]GELDALPEQAFYLVGNIDEASTK

Accession No. XP_004960054/gi|514742084

QVAHAPQELNS[Pho]PR

Accession No. EMS47290/gi|473841507

VAEQLS[Pho]DDEGEDQSK

Accession No. BAJ91166/gi|326505854

AAAVAALSSVLTAEQSGS[Pho]SDNLR

Accession No. EMS67681/gi|474425093

SDS[Pho]QVFLFANSK

Accession No. XP_003559644/gi|357115740

EWQTAAAAAFSES[Pho]DVDEEEDEDELVEVIGAEDEETVLTK

Accession No. XP_006300324/ gi|565485369

LSELLGVEVVMANDS[Pho]IGEEVQKLVAALPEGGVLLLENVR

Accession No. XP_003564482/ gi|357125604

AHGTAVGLPSDDDMGNS[Pho]EVGHNALGAGR

Accession No. BAD36412/ gi|51091643

LSS[Pho]IDAQLR

Accession No. EMT08717/ gi|475535263

ATGAFILTAS[Pho]HNPGGPTEDFGIK

Accession No. EMT06309/ gi|475519222

FDWDHPVHLQPMS[Pho]PTTTK

Accession No. EMT21987/ gi|475592099

FDWDHPMHLQPTS[Pho]PTAVK

Accession No. EMS63629/ gi|474368000

SDDATEVSETDS[Pho]PGDSLR

Accession No. BAN63108/ gi|524456011

QYS[Pho]SGGTEK

QYSS[Pho]GGTEK

GAAS[Pho]LSGK

Accession No. EMS54049/ gi|474063387

AMNKPAEYDS[Pho]DDEIIGTAR

Accession No. AAC05084/ gi|2754825

DEDPLLDS[Pho]DDERPESFDDELR

Accession No. AGW21711/ gi|544370429

S[Pho]LGQNPTEAELQAMINEVDADGNGTIDFPEFLNLMAR

SLGQNPT[Pho]EAELQAMINEVDADGNGTIDFPEFLNLMAR

SLGQNPTEAELQAMINEVDADGNGT[Pho]IDFPEFLNLMAR

Accession No. ABR18064/ gi|148909959

SLGQNPTEAELQDMIS[Pho]EVDADGNGTIDFPEFLNLMAR

Accession No. AAR99410/ gi|41072339

SLGQNPTEAELQDMINEVDADGNGTIDFPEFLNLT[Pho]AR

Accession No. XP_004970381/ gi|514783586

EEWT[Pho]AEVFDVDLLR

Accession No. AAT01376/ gi|46576015

AS[Pho]AEVLGK

Accession No. EMT11740/ gi|475552158

SSS[Pho]FGQQTSGFQQSDSFKQR

SSS[Pho]FGQQTSGFQQSDS[Pho]FKQR

Accession No. ABA95233/ gi|77552436

SLGANLFDRPQPNS[Pho]PTVYDWLYSDETR

Accession No. XP_002465460/ gi|242036131

GT[Pho]GAPVYLAAVLEYLAAEVLELAGNAAR

Accession No. XP_003569004/gi|357134801

VGS[Pho]GAPVY[Pho]LAAVLEYLAAEVLELAGNAAR

PVY[Pho]LAAVLEYLAAEVLELAGNAAR

PVYLAAVLEY[Pho]LAAEVLELAGNAAR

Accession No. XP_006479831/gi|568852331

VGAGAPVYLS[Pho]AVLEYLAAEVLELAGNAAR
