## Supplemental Figure 5 for "Na_2_CO_3_-Responsive Photosynthetic and ROS Scavenging Mechanisms in Chloroplasts of Alkaligrass Revealed by Phosphoproteomics"

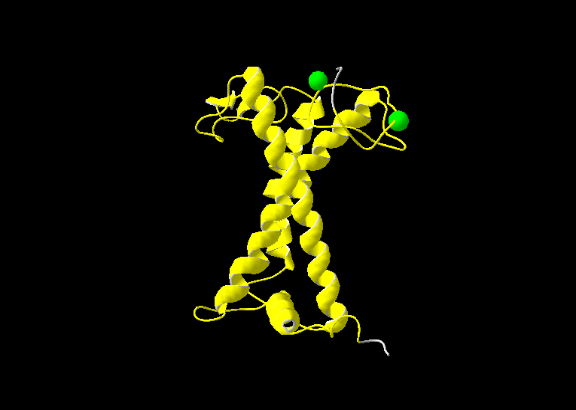

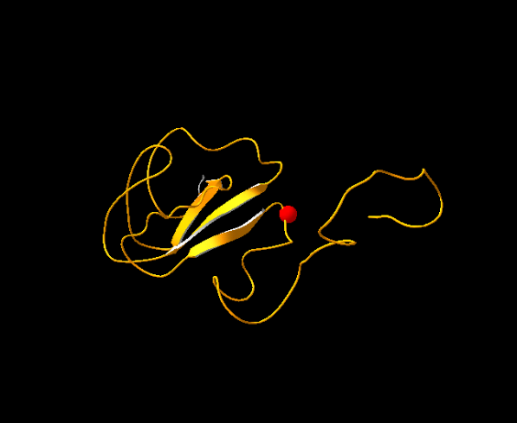

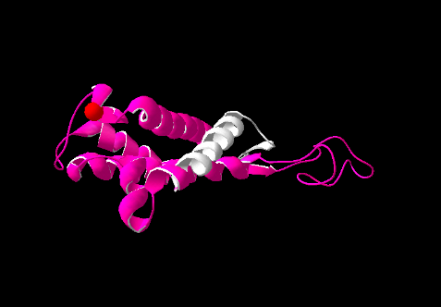

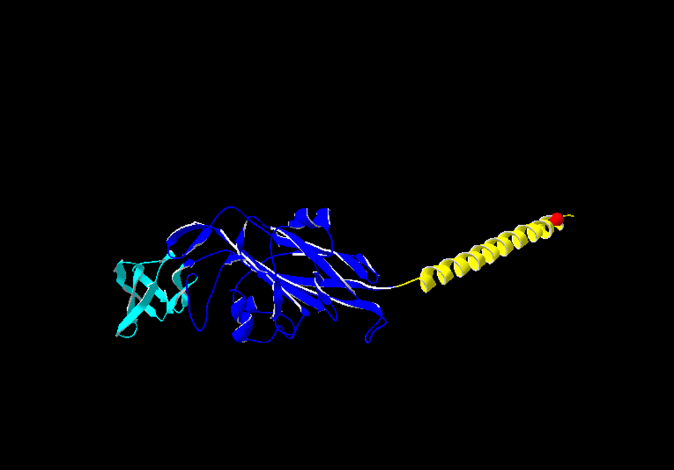

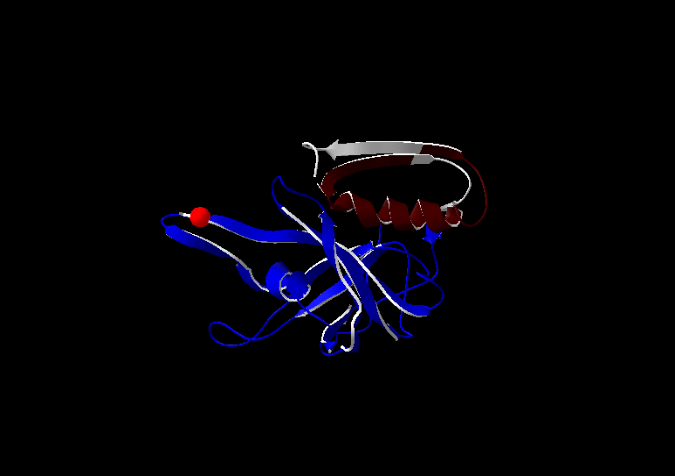

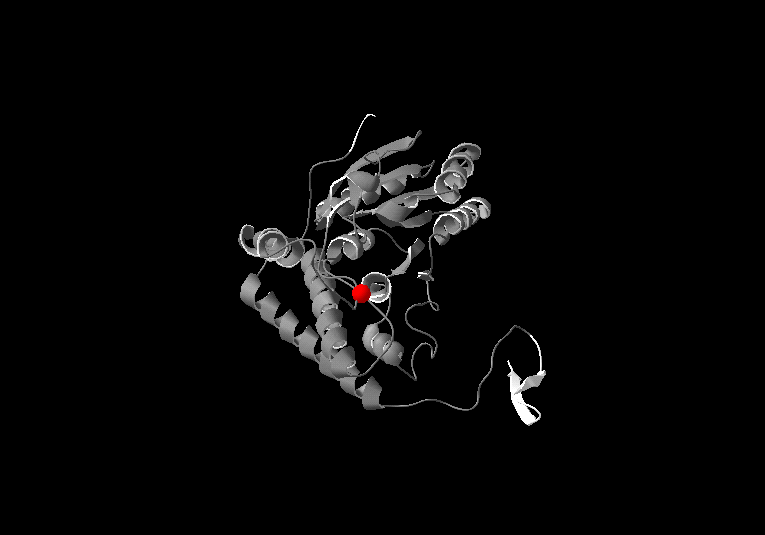

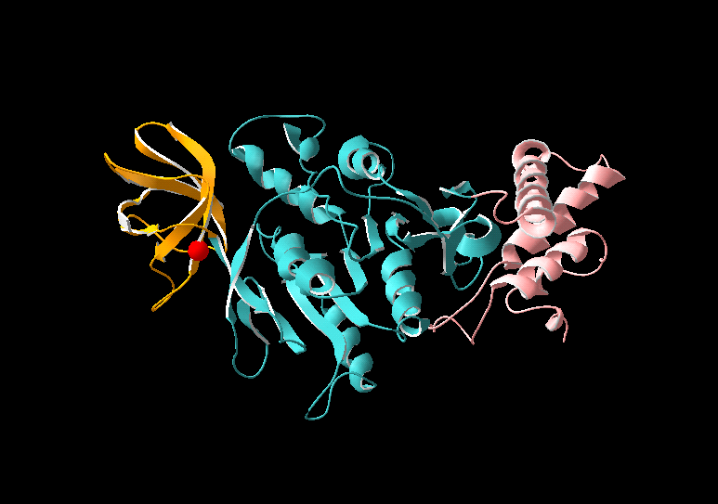

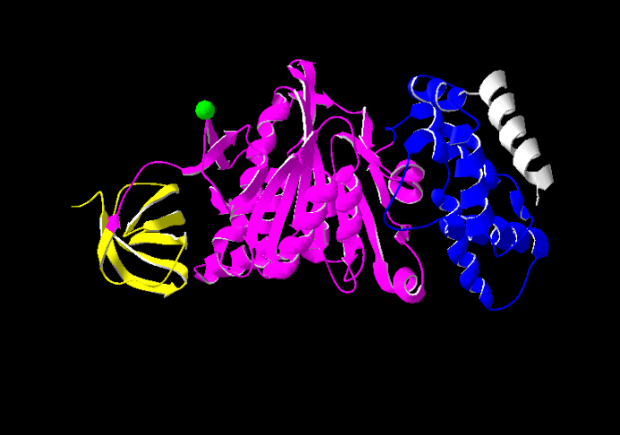

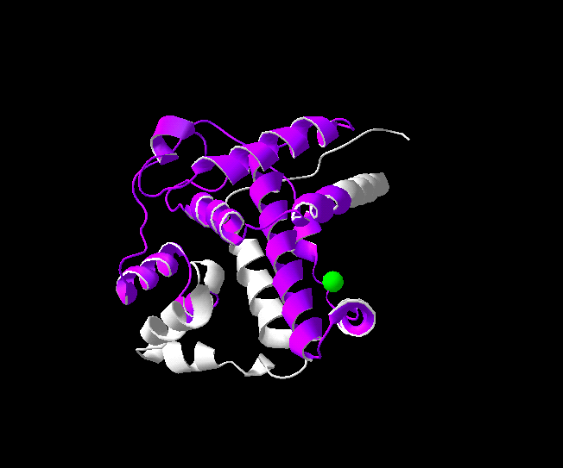

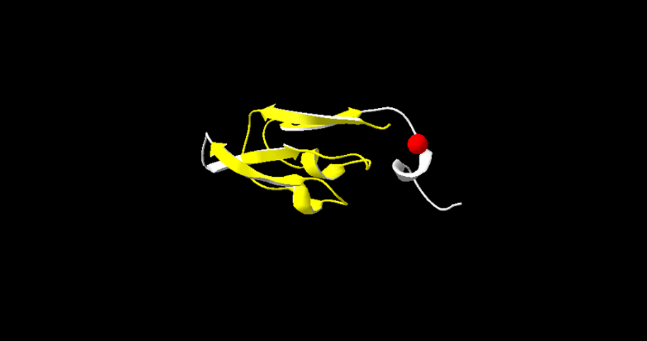

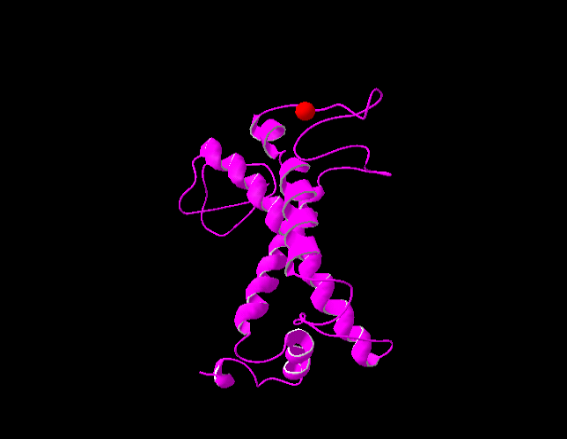

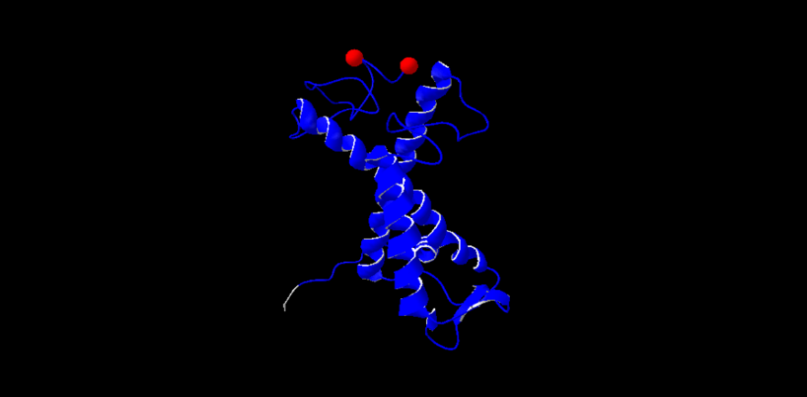

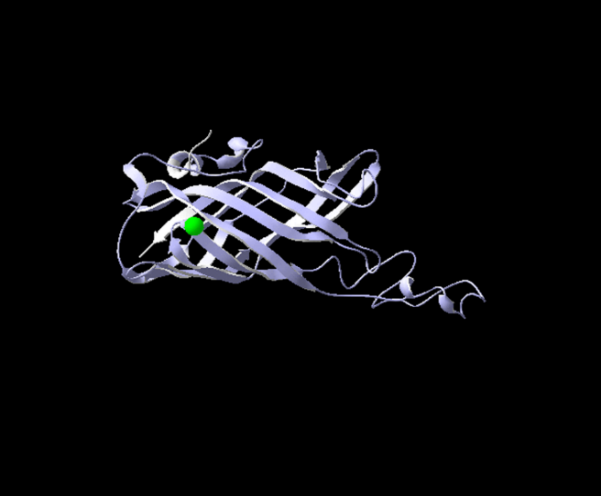

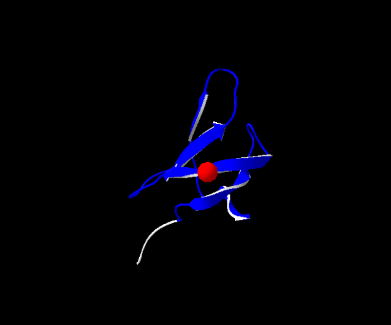

**S166**

B

**Chl a-b binding protein**

**domain**

**Lhca4**

**S53**

**S48**

C

**Chl a-b binding protein**

**domain**

**Lhcb1**

E

**S220**

**PsbO**

**Manganese-stabilising domain**

**Haem peroxidase domain**

F

**S220**

**TL29**

H

**T73**

**4Fe-4S ferredoxin-type domain**

D

**S154**

**PsaC**

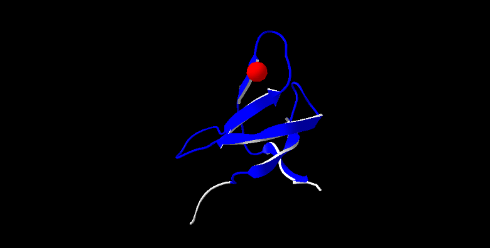

I

**S92**

**Electron transport accessory protein-like domain**

**S106**

**Photosystem I subunit PsaD**

**PsaD**

**I**

**II**

**PsaE**

J

**Flavoprotein pyridine nucleotide cytochrome reductase**

**ATP synthase alpha subunit, central domain**

**Y316**

**Cytochrome *f* large domain**

**T168**

**N-terminal domain**

**S125**

**Transmembrane anchor**

**Cytochrome *f* small domain**

**Ferredoxin reductase-type FAD-binding domain**

**ATP synthase α subunit**

**Cyt *f***

**C-terminal domain**

L

K

M

**Central domain**

**N-terminal domain**

**T53**

**C-terminal domain**

**ATP synthase β subunit**

**FNR**

N

**Thiamin diphosphate-binding fold**

**S296**

**PDH**

**S66**

A

**S122**

G

**Chl a-b binding protein domain**

**Lhca2**

**Chl a-b binding protein domain**

**PsbS**

**S57**
