## Supplemental Figure 7 for "Na_2_CO_3_-Responsive Photosynthetic and ROS Scavenging Mechanisms in Chloroplasts of Alkaligrass Revealed by Phosphoproteomics"

200 mM Na_2_CO_3_, 24 h

150 mM Na_2_CO_3_, 24 h

200 mM Na_2_CO_3_, 12 h

Planting

Sampling

Control

50 days

10:00

10:00

22:00

150 mM Na_2_CO_3_, 12 h

49.5 days

49.5 days

49 days

49 days

12 h

24 h

24 h

12 h

0 mM Na_2_CO_3_

150 mM Na_2_CO_3_

200 mM Na_2_CO_3_
