## Supplemental Figure 8 for "Na_2_CO_3_-Responsive Photosynthetic and ROS Scavenging Mechanisms in Chloroplasts of Alkaligrass Revealed by Phosphoproteomics"

Chloroplast

**h**

P

CP43

D2

D1

P

P

LHCII

P

CP29

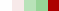

P

**Thylakoid membrane stability**

CURT1A

**f**

**e**

P

P

P

**ROS scavenging**

ToxABP1

PsbS

P

PAP

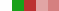

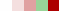

P

P

P

CP26

ABC1K

D1

P

P

Lhcb1

**PSII**

Lhcb2

**PSII**

CP29

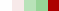

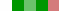

**h**

**c**

**a**

P

TSP9

P

Lhca4

P

P

P

P

**b**

Lhca2

P

Lhca5

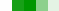

PC

P

Lhcb3

P

Lhcb1

PsbP

**j**

P

P

Villin-2

UROD

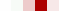

**Protein transport**

Pyruvate

P

**g**

GSA

GSA-AT

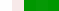

ALA

Protoporphyrinogen IX

NADPH

Tic62

P

P

P

P

H^+^

PPOX

P

**d**

ATP synthase β

Hsp90

Mitochondrion

P

RRM

Ribosome

50S

30S

P

P

CA

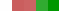

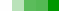

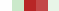

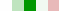

P

Hsc70

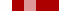

P

P

GS^-^

RNA-BP

P

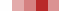

**j**

Citrate

OAA

Isocitrate

2-Oxo-glutarate

Succinate

NADH

NADH

Fumarate

SDH

Malate

FADH_2_

MDH

**TCA Cycle**

CO_2_

CO_2_

IDH

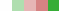

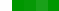

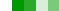

Photodamage

STN7

Pi

Pi

O_2_ + H^+^

H_2_O

PQH_2_

PQ

ISP

Cyt *f*

P

P

Fd

NdhM

NdhJ

NDF1

NdhO

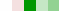

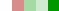

FNR

PsaC

PsaD

PsaE

PsaF

PsaG

PsaH

PsaB

P

P

P

P

H^+^

e^-^

P

ATP synthase

NADP^+^

NADPH

GAP

F6P

E4P

FBA

P

TK

PRK

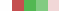

Ru5P

RPE

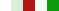

Xu5P

TK

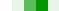

TPI

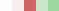

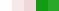

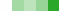

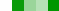

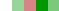

RBS

3-PGA

RuBP

IPP

Phytyl-PP

GGPP

GGDR

LIL3

Carotene

ctDNA

RT

P

RNP

ZF

P

EF-Tu

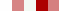

RaiA

AlaRS

P

30S

50S

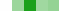

TL15

P

PPIase

HSP70

DnaJ

URO III

SecY

NADP^+^

FtsH

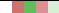

Na_2_CO_3_

TOC

TIC

NACA

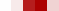

Acetyl CoA

CoA-SH

NADH

DLD

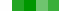

NAD^+^

ATP

ADP

Pyruvate

H^+^

Glycine

NADH

Serine

NH_3_

H_2_O

CO_2_

NAD^+^

SHMT

GDC

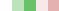

PG

O_2_

CO_2_

Pi

DHA

MDHAR

NAD(P)H

APX

GR

NAD(P)H

NAD(P)^+^

GST

DHAR

H_2_O

GSSG

H_2_O

TPx

PrxR

Trx

H_2_O_2_

GPX

GSH

AsA

MDHA

NAD(P)^+^

O_2_^-^

SOD

Protoporphyrin IX

CSase

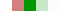

O-acetyl-serine

Mg^2+^

Serine

Cysteine

ABA

Chl *a*

Chl *b*

13^1^-Hydroxy-Mg-Proto ME

Chlorophyllide *a*

Mg-Proto ME

PSAT

Methylglyoxal

H_2_O

NAD(P)H

O_2_^-^

NAD(P)^+^

H_2_O_2_

PrxR

Trx

AsA

CAT

GSH

GLO I

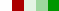

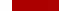

S-lactoylglutathione

SOD

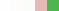

DHAR

P-EAMeT

S-adenosyl-homocysteine

Homocysteine

Phosphocholine

Choline

eIF5A

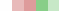

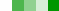

NAD(P)^+^

NAD(P)H

GST

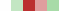

Phosphoethanolamine

SAHH

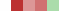

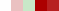

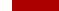

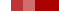

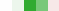

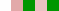

VHA

H^+^

H^+^

Proteasome

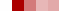

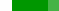

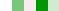

Glycine betaine

MDA

Na^+^

K^+^

P

Na^+^/H^+^ antiporter

STN7/8

Disassembly

LHCII

Pi

LHCII

D1

FtsH

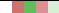

P

LPA1

**PSII**

D1

P

PsbL

TL29

P

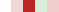

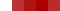

RBP

RBL

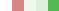

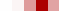

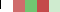

RCA

DLP

P

H^+^

**PSI**

PsbQ

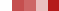

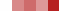

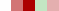

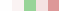

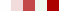

PsbO

e^-^

Glycine

**Photorespiration**

NADP^+^

ATP

ADP + Pi

H^+^

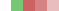

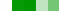

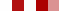

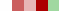

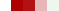

GAPDH

FBA

P

DHAP

FBP

SBP

S7P

R5P

1,3-BPG

**PSII**

PsbH

P

β

ε

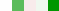

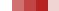

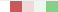

α

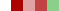

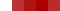

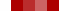

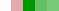

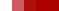

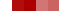

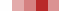

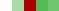

γ

B

H^+^

Actin

**Chloroplast movement**

P

CP24

Heat

e^-^

**Photosynthetic electron transfer**

Lhca1

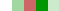

Inactive RuBisCO

**PSII repair cycle**

TLP18.3

D1

**PSII**

MgMT

Mg-Pro IX

MgCy

YCF54

MgCh

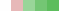

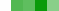

Coprogen III

CPOX

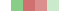

Amino acid

Tic62

**Ionic and osmotic homeostasis**

Proline

ABA

Na^+^

[Ca^2+^] _cyt_

3-phosphonooxypyruvate

Phosphoserine

Glutamate

MDHA

ATP

Na^+^

Na^+^/H^+^ antiporter

**Calvin cycle**

Soluble sugar

Peptide

40S

60S

40S

H_2_O

Protein

Protein

Peptide

Peptide

Amino acid

Protein

**Thermal dissipation**

**State transition**

**Chlorophyll biosynthesis**

Glycolate

PGP

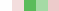

Pre-mRNA

DNA

Nucleus

Chromosome

Transcription

RCC1

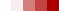

Replication

ADP + Pi^+^

Vacuole

Cell membrane

SRSF

**Gene expression and protein turnover**

**ROS scavenging**

Thylakoid membrane

Light

mRNA

**f**

eIF4B

P

mRNA

**d**

**f**

P

60S

P

Ribosome

GS^-^

APX

MDHAR

GPX

GR

Leaf protein abundance

Chloroplast protein abundance

Leaf enzyme activity

Chloroplast enzyme activity

Chloroplast substrate content

Leaf substrate content

-3.0 -2.0 -1.0 0.0 1.0 2.0 3.0

GSSG

DHA

**i**

**Figure S8** Schematic presentation of Na_2_CO_3_ responsive mechanisms in leaves of *Puccinellia tenuiflora*. The Na_2_CO_3_ responsive proteins from both leaf and chloroplast proteomes were integrated into subcellular pathways. (a) Thermal dissipation. (b) State transition. (c) Photosynthetic electron transfer. (d) Photorespiration and Calvin cycle. (e) PSII repair cycle. (f) ROS scavenging. (g) Chlorophyll biosynthesis. (h) Chloroplast movement and thylakoid membrane stability. (i) Ionic and osmotic homeostasis. (j) Gene expression, protein turnover and transport. The scale bar indicates log (base 2) transformed protein abundance, enzyme activity, and substrate content ratios (compared with 0mM Na_2_CO_3_ treatment), the ratio was ranging from -3.0 to 3.0 with 7 different colors from green to red. The darker the color, the higher of protein abundances, enzyme activities, and substrate contents were stand for. P in a red (increased in phosphorylation level), green (decreased in phosphorylation level), and blue (can be phosphorylated by other protein, but did not identified in our phosphoproteomic study) circle indicates phosphorylated protein. The solid line indicates single-step reaction, dashed line indicates multistep reaction, and the dotted line indicates movement of proteins or other substances. Abbreviations: 1,3-BPG, 1,3-bisphosphoglyceric acid; 13^1^-Hydroxy-Mg-Proto ME, 13^1^-hydroxy-magnesium-protoporphyrin IX 13-monomethyl ester; 3-PGA, 3-phosphoglyceric acid; 30S/50S, 30S/50S ribosomal protein; 40S/60S, eukaryotic small/large ribosomal subunit; ABA, abscisic acid; ABC1K, activator of bc1 complex kinases; ALA, 5-aminolaevulinic acid; AlaRS, alanine-tRNA ligase; APX, ascorbate peroxidase; AsA, ascorbic acid; CA, carbonic anhydrase; CAT, catalase; Coprogen III, corproporphyrinogen III; CP, chlorophyll a-b binding protein; CPOX, coproporphyrinogen-III oxidase; CSase, cysteine synthase; CURT1A, curvature thylakoid 1A protein; Cyt *f*, cytochrome *f*; D1/D2, photosystem II D1/D2 protein; DHA, dehydroascorbic acid; DHAP, dihydroxyacetone phosphate; DHAR, dehydroascorbate reductase; DLD, dihydrolipoyl dehydrogenase; DLP, dynamin-2A-like protein; DnaJ, chaperone protein DnaJ-like; E4P, erythrose 4-phosphate; EF-Tu, elongation factor Tu; eIF4B, eukaryotic initiation factor 4B; eIF5A, eukaryotic translation initiation factor 5A1; F6P, fructose 1,6-diphosphate; FBA, fructose-bisphosphate aldolase; FBP, fructose 1,6-bisphosphate; Fd, ferredoxin; FNR, ferredoxin-NADP(+) reductase; FtsH, ATP-dependent zinc metalloprotease FtsH 2; GAP, glyceraldehyde 3-phosphate; GAPDH, glyceraldehyde 3-phosphate dehydrogenase; GDC, glycine decarboxylase; GGDR, geranylgeranyl diphosphate reductase; GGPP, geranylgeranyl pyrophosphate; GLO I, glyoxalase I; GPX, glutathione peroxidase; GR, glutathione reductase; GS-, xenobiotic substrates; GSA, glutamate 1-semialdehyde aminotransferase; GSA-AT, glutamate-1-semialdehyde 2,1-aminomutase; GSH, glutathione; GSSG, oxidized glutathione; GST, glutathione S-transferase; Hsc70, heat shock cognate 70 kDa protein; Hsp70/90, 70/90 kDa heat shock-related protein; IDH, isocitrate dehydrogenase (NADP)-like; IPP, isopentenyl pyrophosphate; ISP, rieske-like iron-sulfur (2Fe-2S) protein; LHCII, light harvesting complex II; Lhca1/2/4/5, light harvesting chlorophyll a/b binding protein1/2/4/5; Lhcb1/2/3, light harvesting chlorophyll a-b binding protein 1/2/3; LIL3, light harvesting-like protein 3; LPA1, low PSII accumulation 1 protein; MDA, malondialdehyde; MDH, malate dehydrogenase; MDHA, monodehydroascorbate; MDHAR, monodehydroascorbate reductase; Mg-pro IX, Mg-proporphyrin IX; Mg-pro ME, Mg-protoporphyrin IX monomethyl ester; MgCh, magnesium-protoporyphyrin IX chelatase ChlI subunit; MgCy, magnesium-protoporphyrin IX monomethyl ester [oxidative] cyclase; MgMT, magnesium-protoporphyrin O-methyltransferase; NACA, nascent polypeptide associated complex subunit alpha-like protein; NDF1, NDH-dependent cyclic electron flow 1; NdhJ/M/O, NAD(P)H-quinone oxidoreductase subunit J/M/O; OAA, oxaloacetate; P-EAMeT, phosphoethanolamine N-methyltransferase; PAP, plastid-lipid-associated protein; PC, plastocyanin; PG, phosphoglycolate; PGP, phosphoglycolate phosphatase; Phytyl-PP, phytyl pyrophosphate; PPIase, peptidyl-prolyl cis-trans isomerase CYP20-2 domain; PPOX, protoporphyrinogen oxidase; PQ, plastoquinone; PQH_2_, plastohydroquinone; PRK, phosphoribulokinase; Prx, 2-Cys peroxiredoxin BAS1; PsaB, photosystem I P700 chlorophyll A apoprotein A2; PsaC, photosystem I subunit VII; PsaD, photosystem I reaction center subunit II; PsaE, photosystem I reaction center subunit IV; PsaF, photosystem I reaction center subunit III; PsaG, photosystem I subunit V; PsaH, photosystem I reaction center subunit VI; PSAT, phosphoserine aminotransferase; PsbH, photosystem II reaction center protein H; PsbL, Photosystem II subunit L; PsbO, oxygen-evolving enhancer protein 1; PsbP, photosystem II 23kDa oxygen evolving protein; PsbQ, oxygen evolving enhancer protein 3 domain; PsbS, photosystem II 22 kDa protein; R5P, ribose 5-phosphate; RaiA, ribosome-associated inhibitor A domain; RBL, RuBisCO large chain; RBP, RuBisCO large subunit-binding protein; RBS, RuBisCO small chain; RCA, RuBisCO activase; RCC1, regulator of chromosome condensation repeat; RNA-BP, RNA binding protein; RNP, 31 kDa ribonucleoprotein; ROS, reactive oxygen species; RPE, ribulose-phosphate 3-epimerase; RRM, RNA recognition motif; RT, reverse transcriptase domain; Ru5P, ribulose-5-phosphate; RuBP, ribulose-1,5-bisphosphate; S7P, sedoheptulose 7-phosphate; SAHH, S-adenosyl-L-homocysteine hydrolase; SBP, sedoheptulose 1,7-bisphosphate; SDH, succinate dehydrogenase flavoprotein; SecY, preprotein translocase subunit SecY; SHMT, serine hydroxymethyltransferase; SOD, manganese superoxide dismutase; SRSF, serine/arginine-rich splicing factor 33-like; STN7/8, serine/threonine-protein kinase STN7/8; TCA, tricarboxylic acid; Tic62, Tic 62 domain containing protein; TK, transketolase; TL15, thylakoid lumenal 15 kDa protein; TL29, thylakoid lumenal 29 kDa protein; TLP18.3, thylakoid lumen 18.3 kDa protein; TOC/TIC, translocon at the outer/inner envelope membrane of chloroplasts; ToxABP1, chloroplast-localized ToxA binding protein 1; TPI, triosephosphate isomerase; TPx, thioredoxin peroxidase; Trx, thioredoxin; TSP9, thylakoid soluble phosphoprotein; URO III, uroporphyrinogen III; UROD, uroporphyrinogen decarboxylase; VHA, vacuolar H^+^-ATPase; Xu5P, xylulose 5-phosphate; YCF54, YCF54 domain containing protein; ZF, zinc finger protein.
