## Supplemental Table 3 for "Na_2_CO_3_-Responsive Photosynthetic and ROS Scavenging Mechanisms in Chloroplasts of Alkaligrass Revealed by Phosphoproteomics"

**Table S3 Na_2_CO_3_-responsive proteins in alkaligrass leaves**

| **Spot**  **no.^(^*^a^*^)^** | **Protein name^(^*^b^*^)^** | **Accession no.^(^*^c^*^)^** | **Loc^(^*^d^*^)^** | **Biological function^(^*^e^*^)^** | **Thr.**  **MM(Da)**  **/p*I*****^(^*^f^*^)^** | **Exp.**  **MM(Da)**  **/p*I*****^(^*^g^*^)^** | **Sco****^(^*^h^*^)^** | **QM****^(^*^i^*^)^** | **V% ± S.D.^(^*^j^*^)^** |
| --- | --- | --- | --- | --- | --- | --- | --- | --- | --- |
| **Photosynthesis (37)** | | | | | | | | | |
| Chlorophyll a-b binding protein (6)  **  **  **  ** | | | | | | | | | |
| 2778 | Chlorophyll a-b binding protein (Lhca1) | ACO06087 | Chl | PSI Light harvesting | 31,427  /6.12 | 23,380  /5.30 | 108 | 2 |  |
| 3117 | Light harvesting chlorophyll a-b binding protein (Lhca5) | AFS34654 | Chl | PSI Light harvesting | 20,709 /6.34 | 16,059  /4.91 | 121 | 5 | **  **  **  * |
| 2569 | Light harvesting chlorophyll a-b binding protein (Lhcb1) | ADL41158 | Chl | Light harvesting, state transitions | 28,367/  5.14 | 27,171  /4.86 | 161 | 2 | **  **  **  ** |
| 4491 | Light harvesting chlorophyll a-b binding protein (Lhcb1) | ADL41158 | Chl | Light harvesting, state transitions | 28,367/  5.14 | 27,323  /4.72 | 81 | 2 | **  **  **  * |
| 4019 | Light harvesting chlorophyll a-b binding protein of LHCII type II (Lhcb2) | AAC15992 | Chl | Light harvesting, state transitions | 28,564  /5.62 | 27,852  /6.81 | 86 | 2 | **  **  **  ** |
| 4501 | Light harvesting chlorophyll a-b binding protein CP29, photosystem II (CP29) | CAA44777 | Chl | PSII disassembly, energy dissipation | 30,839  /5.33 | 28,338  /4.83 | 201 | 4 | **  **  ** |
| Photosystem II related protein (5) | | | | | | | | | |
| 2263 | Oxygen-evolving enhancer protein 1 (PsbO) | ABQ52657 | Chl | Photosynthetic oxygen evolution, PSII D1 repair | 34,719  /6.08 | 31,545  /4.92 | 209 | 3 | **  **  **  ** |
| 2264 | Oxygen-evolving enhancer protein 1 (PsbO) | ABQ52657 | Chl | Photosynthetic oxygen evolution, PSII D1 repair | 34,719  /6.08 | 31,049  /5.00 | 350 | 4 | **  **  **  ** |
| 2282 | Oxygen-evolving enhancer protein 1 (PsbO) | ABQ52657 | Chl | Photosynthetic oxygen evolution, PSII D1 repair | 34,719  /6.08 | 31,284  /5.09 | 170 | 3 | **  **  **  ** |
| 3303 | Oxygen-evolving enhancer protein 1 (PsbO) | ABQ52657 | Chl | Photosynthetic oxygen evolution, PSII D1 repair | 34,719  /6.08 | 13,459  /5.68 | 132 | 5 | **  **  ** |
| 4512 | Oxygen-evolving enhancer protein 1 (PsbO) | ABQ52657 | Chl | Photosynthetic oxygen evolution, PSII D1 repair | 34,719  /6.08 | 31,545  /4.89 | 195 | 3 | **  **  *  * |
| Photosynthetic electron transfer chain (2) | | | | | | | | | |
| 4030 | Ferredoxin-NADP(+) reductase (FNR) | CAD30024 | Chl | Electron transport | 39,181  /8.29 | 34,767  /6.95 | 147 | 3 | **  **  **  ** |
| 4066 | Ferredoxin-NADP(+) reductase (FNR) | AAA34029 | Chl | Electron transport | 41,399  /8.67 | 36,321  /6.35 | 140 | 2 | **  **  **  ** |
| Calvin cycle (22) | | | | | | | | | |
| 4006 | Predicted protein, containing cd00884, carbonic anhydrase domain (CA)* | BAK00501 | Chl | Catalyze the hydration of CO_2_ to form bicarbonate | 25,214  /8.04 | 24,844  /6.68 | 117 | 2 | **  **  **  ** |
| 4008 | Carbonic anhydrase isoform X1 (CA) | XP_010232070 | Chl | Catalyze the hydration of CO_2_ to form bicarbonate | 28,104  /8.34 | 24,904  /6.72 | 284 | 5 | **  **  **  ** |
| 4196 | Carbonic anhydrase isoform X1 (CA) | XP_010232070 | Chl | Catalyze the hydration of CO_2_ to form bicarbonate | 28,104  /8.34 | 24,450  /6.22 | 221 | 3 | **  **  * |
| 4233 | Carbonic anhydrase isoform X1 (CA) | XP_010232070 | Chl | Catalyze the hydration of CO_2_ to form bicarbonate | 28,104  /8.34 | 25,427  /5.81 | 221 | 4 | **  * |
| 1698 | Ribulose-1,5-bisphosphate carboxylase activase A isoform X1 (RCA) | XP_010237358 | Chl | Activate RuBisCO | 52,371  /5.68 | 42,408  /5.41 | 380 | 4 | **  **  ** |
| 1438 | Ribulose-1,5-bisphosphate carboxylase activase (RCA) | AAP72270 | Chl | Activate RuBisCO | 22,493  /4.98 | 50,539  /5.05 | 163 | 2 | **  **  * |
| 1702 | Ribulose-1,5-bisphosphate carboxylase activase (RCA) | XP_010037804 | Chl | Activate RuBisCO | 49,294  /6.4 | 42,610  /5.55 | 310 | 4 | **  **  *  * |
| 1065 | RuBisCO large subunit-binding protein alpha subunit (RBP) | XP_003558045 | Chl | Assemble RuBisCO | 61,688  /5.31 | 67,272  /5.09 | 89 | 2 | **  **  **  ** |
| 974 | RuBisCO large subunit-binding protein beta subunit (RBP) | EMS68298 | Chl | Assemble RuBisCO | 83,354  /7.79 | 72,116  /5.82 | 412 | 8 | **  **  **  ** |
| 3120 | Ribulose-1,5-bisphosphate carboxylase small chain PW9 (RBS) | XP_003573910 | Chl | Combines CO_2_ to produce 3-phosphoglycerate | 19,728  /8.97 | 15,995  /5.65 | 139 | 3 | **  **  **  ** |
| 3137 | Ribulose-1,5-bisphosphate carboxylase small chain PW9 (RBS) | XP_003573910 | Chl | Combines CO_2_ to produce 3-phosphoglycerate | 19,728  /8.97 | 15,197  /6.30 | 159 | 3 | **  **  **  ** |
| 3147 | Ribulose-1,5-bisphosphate carboxylase small chain PW9 (RBS) | XP_003573910 | Chl | Combines CO_2_ to produce 3-phosphoglycerate | 19,728  /8.97 | 15,442  /5.71 | 87 | 2 | **  **  **  ** |
| 3231 | Ribulose-1,5-bisphosphate carboxylase small chain PW9 (RBS) | XP_003573910 | Chl | Combines CO_2_ to produce 3-phosphoglycerate | 19,728  /8.97 | 14,347  /5.25 | 108 | 3 | **  **  * |
| 3232 | Ribulose-1,5-bisphosphate carboxylase small chain PW9 (RBS) | XP_003573910 | Chl | Combines CO_2_ to produce 3-phosphoglycerate | 19,728  /8.97 | 14,474  /6.30 | 91 | 2 | **  **  ** |

Table S3 *(continued from previous page.)*

| **Spot**  **no.^(^*^a^*^)^** | **Protein name^(^*^b^*^)^** | **Accession no.^(^*^c^*^)^** | **Loc^(^*^d^*^)^** | **Biological function^(^*^e^*^)^** | **Thr.**  **MM(Da)**  **/p*I*^(^*^f^*^)^** | **Exp.**  **MM(Da)**  **/p*I*^(^*^g^*^)^** | **Sco^(^*^h^*^)^** | **QM^(^*^i^*^)^** | **V% ± S.D.^(^*^j^*^)^** |
| --- | --- | --- | --- | --- | --- | --- | --- | --- | --- |
| 1866 | Glyceraldehyde 3-phosphate dehydrogenase subunit A (GAPDH) | ABD37955 | Chl | Reversibly converts 1,3-BPG to GAP | 41,524  /9.15 | 38,929  /6.56 | 127 | 3 | **  **  **  ** |
| 1917 | Glyceraldehyde 3-phosphate dehydrogenase A (GAPDH) | XP_003579898 | Chl | Reversibly converts 1,3-BPG to GAP | 43,1110  /7.01 | 38,001  /6.72 | 322 | 5 | **  **  **  ** |
| 1634 | Predicted protein, glyceraldehyde 3-phosphate dehydrogenase B (GAPDH)* | BAJ86633 | Chl | Reversibly converts 1,3-BPG to GAP | 47,357  /5.9 | 44,397  /5.81 | 255 | 4 | **  **  ** |
| 1649 | Predicted protein, glyceraldehyde 3-phosphate dehydrogenase B (GAPDH)* | BAJ86633 | Chl | Reversibly converts 1,3-BPG to GAP | 47,357  /5.9 | 43,978  /6.01 | 142 | 3 | **  **  **  ** |
| 4109 | Predicted protein, glyceraldehyde 3-phosphate dehydrogenase B (GAPDH)* | BAJ86633 | Chl | Reversibly converts 1,3-BPG to GAP | 47,357  /5.9 | 28,628  /6.62 | 432 | 7 | **  **  **  * |
| 4403 | Triosephosphate isomerase (TPI) | P46225 | Chl | Reversible converts DHAP to GAP | 31,955  /6 | 27,258  /5.38 | 139 | 2 | **  ** |
| 2626 | Ribulose-phosphate 3-epimerase (RPE) | XP_003558725 | Chl | Reversible converts R5P to Ru5P | 29,438  /7.7 | 26,148  /6 | 221 | 2 | **  **  ** |
| 1748 | Phosphoribulokinase (PRK) | CAA41020 | Chl | Catalyze the formation of RuBP | 45,406  /5.84 | 41,089  /5.03 | 317 | 6 | **  **  **  ** |
| Photorespiration (2) | | | | | | | | | |
| 3879 | Phosphoglycolate phosphatase 1B (PGP) | XP_006652446 | Chl | Photorespiration, catalyze the generation of glycolate | 39,724  /6.02 | 33,355  /5.02 | 178 | 5 | **  **  **  ** |
| 3235 | Glycine decarboxylase (GDC) | AAM92707 | Mit | Photorespiration, convert glycine to THF | 21,578  /4.99 | 14,359  /4.60 | 143 | 2 | **  **  ** |
| **Carbohydrate and energy metabolism (22)** | | | | | | | | | |
| Energy metabolism (11) | | | | | | | | | |
| 1027 | ATP synthase CF1 α subunit | AAU06224 | ^#^Chl | ATP synthesis | 55,549  /6.03 | 68,995  /5.86 | 198 | 5 | **  **  ** |
| 4274 | ATP synthase CF1 α subunit | YP_007026550 | ^#^Chl | ATP synthesis | 55,761  /6.11 | 31,308  /5.43 | 699 | 10 | **  **  * |
| 1103 | ATP synthase CF1 α subunit | YP_007026550 | ^#^Chl | ATP synthesis | 55,761  /6.11 | 65,180  /5.97 | 455 | 8 | **  **  **  ** |
| 1109 | ATP synthase CF1 α subunit | YP_007026550 | ^#^Chl | ATP synthesis | 55,761  /6.11 | 64,974  /6.13 | 464 | 8 | **  **  *  * |
| 1171 | ATP synthase CF1 β subunit | CAA25114 | ^#^Chl | ATP synthesis | 53,842  /5.11 | 62,853  /5.13 | 236 | 6 | **  **  **  ** |
| 1195 | ATP synthase CF1 β subunit | NP_114266 | ^#^Chl | ATP synthesis | 53,881  /5.06 | 61,770  /5.44 | 755 | 12 | **  **  **  * |
| 1140 | ATP synthase CF1 β subunit | NP_039390 | ^#^Chl | ATP synthesis | 54,037  /5.47 | 63,955  /5.38 | 176 | 7 | **  **  **  ** |
| 1179 | ATP synthase CF1 β subunit | ABH02572 | ^#^Chl | ATP synthesis | 52,989  /5.16 | 62,556  /5.28 | 238 | 6 | **  **  **  **  **  **  **  ** |
| 1193 | ATP synthase CF1 β subunit | ABH02572 | ^#^Chl | ATP synthesis | 52,989  /5.16 | 61,284  /5.24 | 239 | 6 |  |
| 1220 | ATP synthase β subunit | CAA52636 | Chl, Mit | ATP synthesis | 59,326  /5.56 | 60,419  /5.50 | 523 | 9 | **  **  **  ** |
| 1817 | ATP synthase γ subunit | EMS56225 | ^#^Chl | ATP synthesis | 52,111  /7.52 | 39,700  /6.06 | 56 | 2 | **  **  **  ** |
| Mitochondrion respiratory chain (2) | | | | | | | | | |
| 988 | Carbonic anhydrase-like 2 gamma subunit (γCA) | XP_010235460 | Mit | Mitochondria respiration | 27,782  /8.29 | 70,094  /5.61 | 257 | 5 | **  **  **  * |
| 4212 | Carbonic anhydrase-like 2 gamma subunit (γCA) | XP_010235460 | Mit | Mitochondria respiration | 27,782  /8.29 | 29,236  /6.49 | 115 | 4 | **  **  *  * |
| TCA cycle (4) | | | | | | | | | |
| 4547 | Dihydrolipoyl dehydrogenase, mitochondrial (DLD) | XP_003568861 | Mit | Convert dihydrolipoic acid and NAD+ into lipoic acid and NADH | 52,727  /6.93 | 66,009  /6.97 | 116 | 3 | **  **  **  ** |
| 1572 | Os01g0654500, isocitrate dehydrogenase (IDH)* | NP_001043749 | Cyt, Mit | Convert isocitrate to 2-oxo-glutarate | 46,356  /6.34 | 46,479  /6.35 | 72 | 4 | **  **  **  ** |

Table S3 *(continued from previous page.)*

| **Spot**  **no.^(^*^a^*^)^** | **Protein name^(^*^b^*^)^** | **Accession no.^(^*^c^*^)^** | **Loc^(^*^d^*^)^** | **Biological function^(^*^e^*^)^** | **Thr.**  **MM(Da)**  **/p*I*^(^*^f^*^)^** | **Exp.**  **MM(Da)**  **/p*I*^(^*^g^*^)^** | **Sco^(^*^h^*^)^** | **QM^(^*^i^*^)^** | **V% ± S.D.^(^*^j^*^)^** |
| --- | --- | --- | --- | --- | --- | --- | --- | --- | --- |
| 910 | Succinate dehydrogenase flavoprotein subunit (SDH) | XP_002310225 | Chl, Mit | Catalyzes the oxidation of succinate to fumarate | 70,557  /6.4 | 75,737  /6.26 | 220 | 4 | **  **  **  ** |
| 1982 | Malate dehydrogenase (MDH) | XP_003567846 | Mit | Catalyzes the oxidation of malate to oxaloacetate | 35,600  /8.54 | 37,067  /6.79 | 99 | 3 | **  **  **  ** |
| Glycolysis (3) | | | | | | | | | |
| 1778 | Predicted protein, fructose-bisphosphate aldolase (FBA)* | BAJ85287 | Cyt | Reversibly converts FBP to DHAP and GAP | 38,102  /6.06 | 40,063  /6.84 | 191 | 3 | **  **  **  ** |
| 1862 | Glyceraldehyde 3-phosphate dehydrogenase (GAPDH) | AFS33113 | Cyt | Reversibly converts 1,3-BPG to GAP | 36,801  /6.4 | 38,870  /6.85 | 256 | 4 | **  **  ** |
| 4287 | Glyceraldehyde 3-phosphate dehydrogenase 2 (GAPDH) | P08477 | Cyt | Reversibly converts 1,3-BPG to GAP | 33,443  /6.2 | 31,760  /5.14 | 440 | 6 | **  **  ** |
| Pentose phosphate pathway (1) | | | | | | | | | |
| 3185 | Predicted protein, containing pfam03446, 6-phosphogluconate dehydrogenase NAD binding domain (6PGD)* | BAJ91168 | ^#^Chl, Cyt | Catalyzes the decarboxylating reduction of 6PG into Ru5P | 30,560  /5.97 | 15,185  /4.97 | 103 | 2 | **  **  **  ** |
| Mannitol biosynthesis (1) | | | | | | | | | |
| 4068 | NADPH-dependent mannose 6-phosphate reductase (M6PR) | BAD07953 | Cyt | Reversibly converts mannose 6-phosphate to mannitol 1-phosphate | 35,388  /5.88 | 35,401  /6.37 | 77 | 2 | **  **  ** |
| **Other metabolisms (17)** | | | | | | | | | |
| Chlorophyll metabolism (5) | | | | | | | | | |
| 1695 | Glutamate-1-semialdehyde 2,1-aminomutase (GSA-AT) | P18492 | Chl | Converts GSA to ALA | 49,690  /6.39 | 42,880  /5.87 | 391 | 7 | **  **  ** |
| 1818 | Uroporphyrinogen decarboxylase (UROD) | Q42855 | Chl | Converts URO III to Coprogen III | 36,758  /5.84 | 39,670  /6.36 | 125 | 2 | ** |
| 4448 | Coproporphyrinogen-III oxidase (CPOX) | EMS67249 | Chl | Catalyze the formation of protoporphyrinogen IX | 34,864  /6.02 | 37,012  /5.73 | 396 | 6 | **  **  **  ** |
| 1699 | Magnesium-protoporyphyrin IX chelatase ChlI subunit (MgCh) | EAY90669 | Chl | Insertion of Mg^2+^ into protoporphyrin IX | 44,823  /5.51 | 42,812  /5.03 | 122 | 4 | **  **  **  ** |
| 4369 | Magnesium-protoporyphyrin IX chelatase ChlI subunit (MgCh) | XP_003557120 | Chl | Insertion of Mg^2+^ into protoporphyrin IX | 45,420  /5.9 | 29,455  /5.80 | 434 | 9 | **  **  **  ** |
| Nitrogen metabolism (2) | | | | | | | | | |
| 4517 | Ferredoxin-nitrite reductase (NiR) | BAD53072 | Chl | Catalyzes the assimilation of nitrite to NH_4_^+^ | 70,256  /6.88 | 73,033  /6.27 | 84 | 3 | **  **  * |
| 4534 | Ferredoxin-nitrite reductase (NiR) | BAD53072 | Chl | Catalyzes the assimilation of nitrite to NH_4_^+^ | 70,256  /6.88 | 73,265  /6.41 | 84 | 3 | **  **  **  ** |
| Nucleotide metabolism (2) | | | | | | | | | |
| 3013 | Unknown, containing cd04413, nucleoside diphosphate kinase group I-like domain (NDPK)* | ACN31682 | Chl, Mit | Catalyzes the production of (d)NTPs; Oxidative stress response | 26,014  /9.04 | 17,845  /6.63 | 149 | 3 | **  **  **  ** |
| 3080 | Os12g0548300, nucleoside diphosphate kinase 2 (NDPK)* | NP_001066971 | Chl | Catalyzes the production of (d)NTPs; Oxidative stress response | 23,626  /9.51 | 16,700  /5.53 | 94 | 2 | **  **  **  ** |
| Amino acid metabolism (6) | | | | | | | | | |
| 1599 | Aspartate aminotransferase (AAT) | EMS51671 | Chl | Catalyzes the formation of aspartate and α-ketoglutarate | 52,199  /6.77 | 45,750  /5.85 | 178 | 4 | **  **  **  ** |
| 1781 | Glutamine synthetase (GS) | S18603 | Cyt | Catalyzes glutamate and NH_4_^+^ to form glutamine | 40,932  /5.4 | 40,254  /5.5 | 136 | 3 | **  **  **  ** |
| 1007 | Predicted protein, containing cd07940, 2-isopropylmalate synthase, N-terminal catalytic TIM barrel domain (IPMS)* | BAJ93533 | Chl | Reversibly convert acetyl-CoA and 3-methyl-2-oxobutanoate to (2S)-2-isopropylmalate | 67,880  /6.28 | 70,316  /5.72 | 173 | 7 | **  **  **  ** |
| 1503 | S-adenosylmethionine synthetase (SAMS) | CAJ01702 | Cyt | Participate in lignin biosynthesis | 43,243  /5.49 | 48,505  /5.93 | 283 | 4 | **  **  **  ** |
| 1178 | S-adenosyl-L-homocysteine hydrolase (SAHH) | CAJ01706 | Cyt | Reversible hydration of SAH into homocysteine | 49,960  /5.81 | 62,359  /5.91 | 388 | 5 | **  **  **  ** |
| 3873 | Cysteine synthase (CSase) | EMS61355 | Chl | Catalyzes the formation of cysteine | 98,315  /5.04 | 35,804  /5.09 | 445 | 7 | **  **  ** |
| Formate metabolism (1) | | | | | | | | | |
| 1664 | Predicted protein, formate dehydrogenase (FDH)* | BAJ95739 | Mit | Catalyzes the oxidation of formate into CO_2_ | 41,693  /6.51 | 43,770  /6.95 | 262 | 5 | **  **  **  ** |

Table S3 *(continued from previous page)*

| **Spot**  **no.^(^*^a^*^)^** | **Protein name^(^*^b^*^)^** | **Accession no.^(^*^c^*^)^** | **Loc^(^*^d^*^)^** | **Biological function^(^*^e^*^)^** | **Thr.**  **MM(Da)**  **/p*I*^(^*^f^*^)^** | **Exp.**  **MM(Da)**  **/p*I*^(^*^g^*^)^** | **Sco^(^*^h^*^)^** | **QM^(^*^i^*^)^** | **V% ± S.D.^(^*^j^*^)^** |
| --- | --- | --- | --- | --- | --- | --- | --- | --- | --- |
| Fatty acid metabolism (1) | | | | | | | | | |
| 1446 | ATP citrate lyase A-1 (ACL) | NP_172537 | Cyt | Involved in fatty acid biosynthesis | 46,991  /5.39 | 50,300  /5.45 | 61 | 2 | **  **  **  ** |
| **Stress and defense (4)** | | | | | | | | | |
| 4149 | Manganese superoxide dismutase (MnSOD) | AAA33512 | Mit | ROS scavenging | 25,563  /7.11 | 24,944  /6.35 | 124 | 2 | **  ** |
| 2618 | Predicted protein, containing cd03187, glutathione S-transferase C-terminal alpha helical domain (GST)* | BAJ85232 | Cyt | ROS scavenging | 25,155  /5.51 | 26,400  /6.30 | 73 | 2 | **  **  **  ** |
| 4288 | Glyoxalase I (GLO I) | AAS98483 | Cyt | Catalyzes methylglyoxal detoxification | 29,720  /4.99 | 32,315  /5.16 | 82 | 3 | **  **  **  ** |
| 4359 | Glyoxalase I (GLO I) | AAS98483 | Cyt | Catalyzes methylglyoxal detoxification | 29,720  /4.99 | 26,933  /5.56 | 171 | 6 | **  **  **  ** |
| **Membrane and transporting (5)** | | | | | | | | | |
| 870 | Vacuolar proton-ATPase subunit A (VHA-A) | ABD85016 | Cyt | Sequestrate Na^+^ into the vacuole | 68,754  /5.23 | 78,292  /5.43 | 272 | 10 | **  **  **  ** |
| 876 | Vacuolar proton-ATPase subunit A (VHA-A) | ABD85016 | Cyt | Sequestrate Na^+^ into the vacuole | 68,754  /5.23 | 78,168  /5.46 | 243 | 9 | **  **  *  * |
| 1191 | Vacuolar proton ATPase subunit B (VHA-B) | NP_195563 | Cyt | Sequestrate Na^+^ into the vacuole | 54,271  /5.03 | 62,162  /5.19 | 96 | 5 | **  **  **  ** |
| 4035 | Vacuolar proton ATPase subunit E (VHA-E) | AAD10335 | Cyt | Sequestrate Na^+^ into the vacuole | 26,359  /6.57 | 31,403  /6.99 | 207 | 5 | **  **  * |
| 4286 | Vacuolar proton ATPase subunit E (VHA-E) | AAD10335 | Cyt | Sequestrate Na^+^ into the vacuole | 26,359  /6.57 | 30,511  /5.13 | 337 | 5 | **  **  **  * |
| **Signaling (2)** | | | | | | | | | |
| 3930 | 14-3-3-like protein (14-3-3) | AAP48904 | Cyt | Signal transduction | 28,864  /4.79 | 30,449  /4.78 | 108 | 2 | **  **  * |
| 3932 | 14-3-3-like protein (14-3-3)* | BAB11739 | Cyt | Signal transduction | 29,491  /4.75 | 31,808  /4.78 | 162 | 3 | **  **  **  * |
| **Protein synthesis and turnover (10)** | | | | | | | | | |
| 1602 | 30S ribosomal protein S1 (30S RP) | ABF95619 | Chl | Protein synthesis | 43,282  /4.7 | 45,462  /4.57 | 129 | 3 | **  **  **  ** |
| 2921 | Eukaryotic translation initiation factor 5A1 (eIF5A) | AAZ95171 | Cyt | Protein synthesis | 17,580  /5.7 | 19,768  /5.67 | 70 | 3 | **  **  **  ** |
| 2927 | Eukaryotic translation initiation factor 5A1 (eIF5A) | AAZ95171 | Cyt | Protein synthesis | 17,983  /5.45 | 19,705  /5.38 | 90 | 3 | **  **  **  ** |
| 1498 | Elongation factor Tu (EF-Tu) | Q43467 | Chl | Protein synthesis | 52,177  /6.21 | 48,352  /5.32 | 68 | 2 | **  **  ** |
| 3815 | Predicted protein, nascent polypeptide associated complex subunit alpha-like protein 1 (NACA)* | XP_001785878 | Nuc | Protein folding | 21,592  /4.36 | 29,635  /4.25 | 155 | 5 | **  **  **  * |
| 782 | Heat shock cognate 70 kDa protein 2-like (Hsc70) | XP_006664117 | Cyt | Protein folding | 71,546  /5.09 | 83,532  /5.23 | 262 | 6 | **  **  **  ** |
| 905 | ATP-dependent zinc metalloprotease FtsH 2 (FtsH) | EMS55427 | Chl | Degradation of photodamaged D1 protein | 71,987  /5.7 | 75,856  /5.25 | 696 | 10 | **  **  **  ** |
| 1588 | Proteasome subunit alpha type-3 (Proteasome α3) | XP_004970311 | Cyt, Nuc | Protein degradation | 27,465  /5.75 | 45,967  /5.28 | 105 | 3 | **  **  **  ** |
| 1013 | Predicted protein, proteasome subunit alpha type-6 (Proteasome α6)* | BAJ85935 | Nuc | Protein degradation | 27,547  /6.33 | 69,983  /5.28 | 166 | 3 | **  **  *  * |
| 4020 | Proteasome subunit alpha type-6 (Proteasome α6) | Q9LSU3 | Nuc | Protein degradation | 27,670  /6.19 | 27,919  /6.86 | 35 | 2 | **  **  ** |
| **Cell wall metabolism (2)** | | | | | | | | | |
| 3283 | Predicted protein, caffeoyl-CoA O-methyltransferase (CCoAOMT)* | BAK01687 | ^#^Chl, Cyt | Lignin biosynthesis | 29,520  /5.09 | 13,996  /4.60 | 111 | 3 | **  **  **  ** |
| 4281 | Predicted protein, caffeoyl-CoA O-methyltransferase (CCoAOMT)* | BAK01687 | ^#^Chl, Cyt | Lignin biosynthesis | 29,520  /5.09 | 28,052  /5.27 | 215 | 3 | **  ** |
| **Cell cycle (1)** | | | | | | | | | |
| 4228 | Predicted protein, containing pfam00415, regulator of chromosome condensation repeat (RCC1)* | BAJ88198 | ^#^Nuc | Cell cycle | 47,167  /5.35 | 22,143  /5.90 | 394 | 8 | **  **  ** |

Table S3 *(continued from previous page.)*

| **Spot**  **no.^(^*^a^*^)^** | **Protein name^(^*^b^*^)^** | **Accession no.^(^*^c^*^)^** | **Loc^(^*^d^*^)^** | **Biological function^(^*^e^*^)^** | **Thr.**  **MM(Da)**  **/p*I*^(^*^f^*^)^** | **Exp.**  **MM(Da)**  **/p*I*^(^*^g^*^)^** | **Sco^(^*^h^*^)^** | **QM^(^*^i^*^)^** | **V% ± S.D.^(^*^j^*^)^** |
| --- | --- | --- | --- | --- | --- | --- | --- | --- | --- |
| **Miscellaneous and function unknown (4)** | | | | | | | | | |
| 4062 | Unknown, containing cd00200, WD40 domain (WD40)* | ACL54313 | Cyt | Miscellaneous | 36,700  /6.13 | 36,540  /6.46 | 142 | 3 | **  **  **  ** |
| 4581 | Os07g0212200, containing cd05265, atypical short-chain dehydrogenase/reductase domain (SDR)* | NP_001059177 | Chl | Function unknown | 40,983  /7.68 | 38,841  /5.84 | 77 | 4 | **  **  **  ** |
| 987 | Os07g0212200, containing cd05265, atypical short-chain dehydrogenase/reductase domain (SDR)* | NP_001059177 | Chl | Function unknown | 41,268  /7.68 | 71,210  /5.25 | 112 | 4 | **  **  **  ** |
| 1072 | Uncharacterized protein | XP_003573626 | Chl | Function unknown | 33,171  /8.64 | 66,849  /5.48 | 193 | 4 | **  **  **  ** |

*Note: ^a^*Assigned spot number as indicated in Figure S3.

*^b^*The name and functional categories of the proteins identified by MALDI TOF-TOF mass spectrometry, protein names marked with an asterisk (*) have been edited by us depending on searching against NCBI non-redundant protein database for functional domain. The abbreviations for the protein names are indicated in the bracket after protein names.

*^c^*Database accession numbers from NCBInr.

*^d^*Protein subcellular localization (Loc) predicted by softwares (YLoc, LocTree3, Plant-mPLoc, ngLOC, and ChloroP). Only the consistent predictions from at least two tools were accepted as a confident result. Pounds (#) indicate the subcellular localizations were predicted based on literatures listed in Supplemental Table S6. Chl, chloroplast; Cyt, cytoplasm; Mit, mitochondria; Nuc, nucleus.

*^e^*The biological function of the identified proteins.

*^f,g^*Theoretical (*f*) and experimental (*g*) molecular mass (Da) and p*I* of identified proteins. Experimental values were calculated using ImageMaster 2D platinum software. Theoretical values were retrieved from the protein database.

*^h^*The Mascot score obtained after searching against the NCBInr database.

*^i^*The number of unique peptides identified for each protein.

*^j^*Change patterns of the Na_2_CO_3_-responsive proteins. The mean values of protein spot volumes relative to total volume of all the spots (V%). Error bar indicates ± standard deviation (S.D.). The asterisks indicate significant differences (*, P < 0.05; **, P < 0.01).

Abbreviations: 1,3-BPG, 1,3-bisphosphoglyceric acid; 6PG, 6-phosphogluconate; ALA, 5-aminolaevulinic acid; Coprogen III, corproporphyrinogen III; DHAP, dihydroxyacetone phosphate; FBP, fructose 1,6-bisphosphate; GAP, glyceraldehyde 3-phosphate; GSA, glutamate 1-semialdehyde aminotransferase; R5P, ribose 5-phosphate; ROS, reactive oxygen species; Ru5P, ribulose-5-phosphate; RuBP, ribulose-1,5-bisphosphate; THF, N^5^,N^10^-methylene tetrahydrofolate; URO III, uroporphyrinogen III.
