## Supplemental Table 6 for "Na_2_CO_3_-Responsive Photosynthetic and ROS Scavenging Mechanisms in Chloroplasts of Alkaligrass Revealed by Phosphoproteomics"

**Table S6 Na_2_CO_3_-responsive proteins in alkaligrass chloroplasts**

| **Protein name****^(^*^a^*^)^** | **Accession no.^(^*^b^*^)^** | **Loc^(c)^** | **Biological function^(^*^d^*^)^** | **MM(kD)/p*I*****^(^*^e^*^)^** | **PA^(^*^f^*^)^** |
| --- | --- | --- | --- | --- | --- |
| **Photosynthesis (45)** | | | | | |
| Chlorophyll a-b binding protein (5) | | | | | |
| Light harvesting chlorophyll a-b binding protein (Lhca1) | ACO06087 | Chl | Light harvesting for PSI | 30.93  /6.12 | 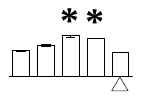 |
| Predicted protein, containing PLN00147, light harvesting complex I chlorophyll a-b binding protein Lhca5 (Lhca5)* | BAJ92159 | Chl | Light harvesting for PSI | 29.35  /8.68 | 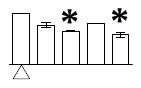 |
| Predicted protein, containing PLN00025, light harvesting chlorophyll a-b binding protein of LHCII type I-like (Lhcb1)* | BAK03973 | Chl | Light harvesting, state transition | 28.09  /5.28 | 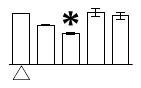 |
| Predicted protein, light harvesting chlorophyll a-b binding protein Lhcb6, chloroplastic (CP24)* | BAK04677 | Chl | Light harvesting, energy dissipation | 27.39  /6.75 | 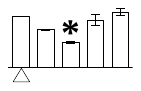 |
| Light harvesting chlorophyll a-b binding protein Lhcb4, chloroplastic(CP29) | 1908421A | Chl | Light harvesting, PSII disassembly, energy dissipation | 31.31  /5.32 | 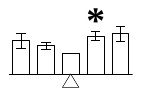 |
| Photosystem II related protein (14) | | | | | |
| Predicted protein, serine/threonine-protein kinase STN7 (STN7)* | BAJ86343 | Chl | Phosphorylation LHCII, CP29 | 62.43  /9.17 | 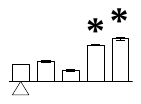 |
| Predicted protein, serine/threonine-protein kinase STN8 (STN8)* | BAJ88887 | Chl | Phosphorylation PSII core proteins | 63.72  /6.07 | 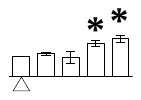 |
| Predicted protein, photosystem II 22 kDa protein (PsbS)* | BAJ96994 | Chl | Energy dissipation | 28.21  /8.77 | 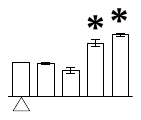 |
| Photosystem II D1 protein (D1) | AAU94352 | Chl | PSII core protein | 38.92  /5.2 | 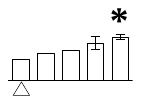 |
| Photosystem II CP43 chlorophyll apoprotein (CP43) | ADO65461 | Chl | PSII core protein | 52.11  /6.7 | 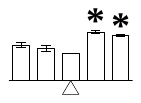 |
| Hypothetical protein OsJ_22738, photosystem II stability/assembly factor HCF136 (HCF136)* | EEE66398 | Chl | Stability/assembly PSII | 171.13  /7.09 | 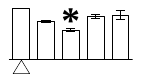 |
| Predicted protein, containing pfam04536, thylakoid lumen 18.3 kDa protein domain (TLP18.3)* | BAK01555 | Chl | Dephosphorylated the damaged D1 | 30.42  /6.34 | 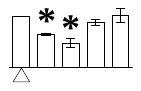 |
| Hypothetical protein, chloroplast-localized Ptr ToxA-binding protein 1 (ToxABP1)* | BAJ85363 | Chl | Stabilize FtsH in chloroplast stroma | 32.34  /9.13 | 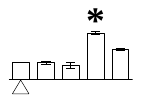 |
| Photosystem II reaction center protein H (PsbH) | ACJ70783 | Chl | D1 maturation and incorporation | 7.77  /8.09 | 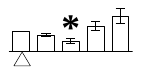 |
| Predicted protein, Low PSII accumulation 1 protein (LPA1)* | BAJ94646 | Chl | Integration of D1 protein into PSII | 49.39  /9.45 | 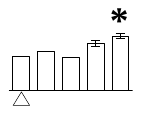 |
| Predicted protein, photosystem II oxygen evolving enhancer protein 1 (PsbO)* | BAJ85472 | Chl | Photosynthetic oxygen evolution, PSII D1 repair | 34.47  /5.74 | 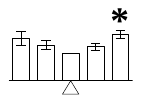 |
| Photosystem II 23kDa oxygen evolving protein (PsbP) | CAA40669 | Chl | Photosynthetic oxygen evolution, PSII activation | 27.27  /8.83 | 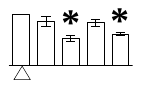 |
| Predicted protein, PsbP-like protein 1 (PsbP)* | BAJ93986 | Chl | Photosynthetic oxygen evolution, PSII activation | 25.10  /9.46 | 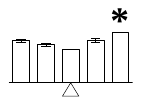 |
| Predicted protein, containing pfam05757, oxygen evolving enhancer protein 3 domain (PsbQ)* | BAJ90882 | Chl | Photosynthetic oxygen evolution, PSII activation | 26.36  /9.86 | 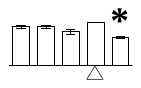 |
| Photosystem I related protein (7) | | | | | |
| Photosystem I P700 chlorophyll A apoprotein A2 (PsaB) | AAM12498 | Chl | Core of the PSI reaction center, binding the electron transport cofactors P700 | 82.59  /6.63 | 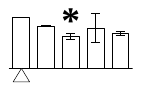 |
| Photosystem I subunit VII (PsaC) | AAZ04109 | Chl | Carries the [4Fe–4S] F_A_ and F_B_ clusters | 8.90  /6.51 | 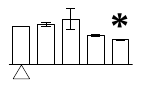 |
| Predicted protein, photosystem I reaction center subunit II (PsaD)* | BAK07564 | Chl | Provide the docking site for soluble ferredoxin | 21.93  /9.81 | 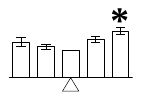 |
| Predicted protein, photosystem I reaction center subunit III (PsaF)* | BAK02382 | Chl | Interaction with the lumenal electron donor plastocyanin | 22.23  /9.37 | 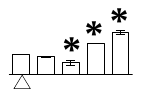 |
| Photosystem I reaction center subunit III (PsaF) | P13192 | Chl | Interaction with the lumenal electron donor plastocyanin | 24.84  /9.46 | 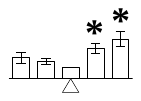 |
| Photosystem I subunit V (PsaG) | Q2L3V4 | Chl | Stabilizing the PSI complex | 15.11  /9.59 | 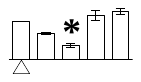 |
| Photosystem I reaction center subunit VI (PsaH) | CAA34218 | Chl | Stabilizing the PSI complex | 14.88  /10.28 | 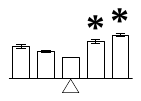 |
| Photosynthetic electron transfer chain related protein (9) | | | | | |
| Cytochrome *f* (Cyt *f*) | A1EA21 | Chl | Electron transport | 35.38  /8.61 | 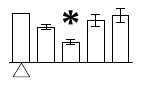 |
| Predicted protein, containing cd03467, rieske-like iron-sulfur (2Fe-2S) protein domain (ISP)* | BAJ90855 | Chl | Electron transport | 29.68  /6.12 | 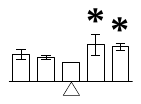 |
| Ferredoxin-NADP(H) oxidoreductase (FNR) | CAD30025 | Chl | Electron transport | 40.23  /6.91 | 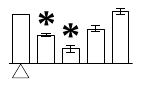 |
| Predicted protein, Ferredoxin-NADP+ reductase, leaf isozyme (FNR)* | BAJ99653 | Chl | Electron transport | 39.60  /8.29 | 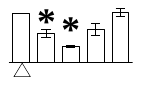 |

Table S6 *(continued from previous page.)*

| **Protein name^(^*^a^*^)^** | **Accession no.^(^*^b^*^)^** | **Loc^(c)^** | **Biological function^(^*^d^*^)^** | **MM(kD)/p*I*^(^*^e^*^)^** | **PA^(^*^f^*^)^** |
| --- | --- | --- | --- | --- | --- |
| NAD(P)H-quinone oxidoreductase subunit J (NdhJ) | ADD63040 | Chl | Cyclic electron transport | 22.23  /8.8 | 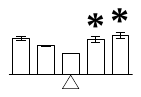 |
| Predicted protein, NAD(P)H-quinone oxidoreductase subunit M (NdhM)* | BAJ92345 | Chl | Cyclic electron transport | 23.65  /7.06 | 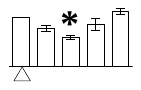 |
| Predicted protein, NAD(P)H-quinone oxidoreductase subunit M (NdhM)* | BAJ88809 | Chl | Cyclic electron transport | 24.02  /6.84 | 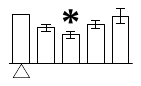 |
| Hypothetical protein SORBIDRAFT_03g046510, NAD(P)H-quinone oxidoreductase subunit O (NdhO)* | EES02088 | Chl | Cyclic electron transport | 17.23  /9.49 | 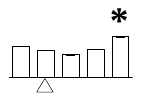 |
| Predicted protein, NDH-dependent cyclic electron flow 1 (NDF1)* | BAJ91603 | Chl | Cyclic electron transport | 38.04  /6.26 | 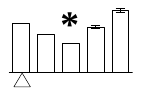 |
| Calvin cycle (10) | | | | | |
| Unnamed protein product, ribulose-1,5-bisphosphate carboxylase/oxygenase large subunit-binding protein alpha subunit (RBP)* | CAA30699 | Chl | Assemble RuBisCO | 57.52  /4.83 | 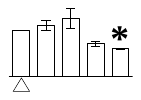 |
| Ribulose-1,5-bisphosphate carboxylase/oxygenase large subunit-binding protein beta subunit (RBP) | Q43831 | Chl | Assemble RuBisCO | 53.41  /4.87 | 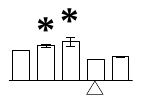 |
| Predicted protein, ribulose-1,5-bisphosphate carboxylase/oxygenase small chain (RBS)* | BAJ87130 | Chl | Combines CO_2_ to produce 3-phosphoglycerate | 19.47  /8.81 | 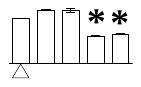 |
| Ribulose-1,5-bisphosphate carboxylase/oxygenase large chain (RBL) | AAA32046 | Chl | Combines CO_2_ to produce 3-phosphoglycerate | 52.91  /6.03 | 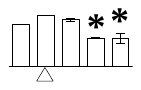 |
| Ribulose-1,5-bisphosphate carboxylase/oxygenase large subunit (RBL) | AAX44980 | Chl | Combines CO_2_ to produce 3-phosphoglycerate | 53.13  /6.22 | 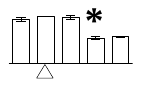 |
| Predicted protein, glyceraldehyde-3-phosphate dehydrogenase A (GAPDH)* | BAK08149 | Chl | Catalyze the conversion of 1,3-BPG to make GAP | 42.70  /7.60 | 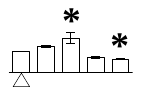 |
| Predicted protein, glyceraldehyde-3-phosphate dehydrogenase B (GAPDH)* | BAJ87214 | Chl | Catalyze the conversion of 1,3-BPG to make GAP | 46.87  /6.02 | 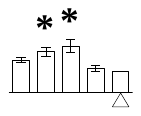 |
| Unnamed protein product, transketolase (TK)* | CAY37266 | Chl | Conversion S7P to R5P and Xu5P | 79.95  /5.9 | 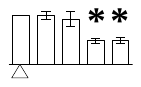 |
| Predicted protein, transketolase (TK)* | BAJ93658 | ^#^Chl | Conversion S7P to R5P and Xu5P | 73.57  /5.45 | 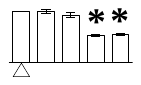 |
| Phosphoribulokinase (PRK) | ACG42204 | Chl | Catalyze the Ru5P phosphorylation to form RuBP | 45.71  /5.74 | 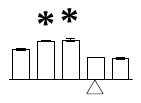 |
| **Energy metabolism (4)** | | | | | |
| ATP synthase γ subunit | ADC33136 | Chl | ATP synthesis | 39.75  /8.17 | 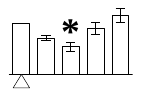 |
| ATP synthase ε subunit | YP_002000493 | ^#^Chl | ATP synthesis | 15.27  /5.19 | 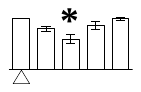 |
| ATP synthase ε subunit | YP_002364507 | ^#^Chl | ATP synthesis | 15.22  /5.44 | 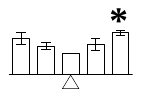 |
| ATP synthase CF0 B subunit | A1EA04 | ^#^Chl | ATP synthesis | 21.35  /9.47 | 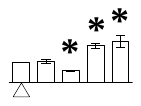 |
| **Other metabolisms (14)** | | | | | |
| Chlorophyll metabolism (8) | | | | | |
| Predicted protein, protoporphyrinogen oxidase (PPOX)* | BAK02106 | Chl | Catalyzes the formation of protoporphyrin IX | 56.20  /9.24 | 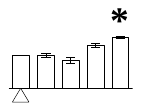 |
| Predicted protein, magnesium-protoporphyrin O-methyltransferase (MgMT)* | BAJ92664 | Chl | Catalyzes the transfer of methyl group to Mg-Pro IX | 34.74  /8.73 | 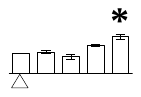 |
| Predicted protein, magnesium-protoporphyrin IX monomethyl ester [oxidative] cyclase (MgCy)* | BAJ96823 | Chl | Catalyzes the formation of protochlorophyllide | 48.16  /8.86 | 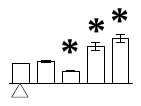 |
| Magnesium-protoporphyrin IX monomethyl ester [oxidative] cyclase (MgCy) | AAW66004 | Chl | Catalyzes the formation of protochlorophyllide | 47.91  /8.88 | 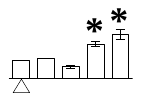 |
| Unknown protein, containing Ycf54 domain (YCF54)* | BAJ91312 | Chl | Worked as one of the components of MgCy | 23.93  /7.76 | 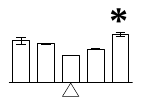 |
| Geranylgeranyl diphosphate reductase (GGDR) | AAZ67145 | Chl | Catalyze the conversion of GGPP to Phytyl-PP | 50.57  /9.26 | 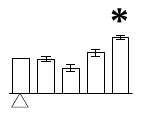 |
| Predicted protein, geranylgeranyl diphosphate reductase (GGDR)* | BAK04705 | Chl | Catalyze the conversion of GGPP to Phytyl-PP | 50.67  /9.21 | 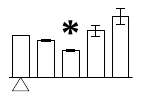 |
| Predicted protein, light harvesting-like protein 3 (LIL3)* | BAJ88054 | Chl | Stabilizing GGDR | 27.41  /6.43 | 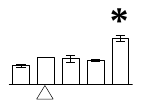 |
| Nucleotide metabolism (1) | | | | | |
| Predicted protein, nucleoside diphosphate kinase (NDPK)* | BAJ87594 | Chl, Cyt | Catalyzes the production of (d)NTPs; Oxidative stress response | 26.54  /9.12 | 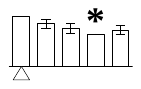 |

Table S6 *(continued from previous page.)*

| **Protein name^(^*^a^*^)^** | **Accession no.^(^*^b^*^)^** | **Loc^(c)^** | **Biological function^(^*^d^*^)^** | **MM(kD)/p*I*^(^*^e^*^)^** | **PA^(^*^f^*^)^** |
| --- | --- | --- | --- | --- | --- |
| Amino acid metabolism (3) | | | | | |
| Plastidic glutamine synthetase (GS) | BAD12059 | Chl | Catalyzes glutamate and NH_4_^+^ to form glutamine | 46.69  /6.21 | 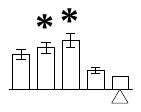 |
| Unnamed protein product, glutamine synthetase, chloroplastic (GS)* | CBN69844 | Chl | Catalyzes glutamate and NH_4_^+^ to form glutamine | 46.77  /6.21 | 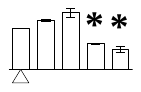 |
| Phosphoserine aminotransferase (PSAT) | AAN06833 | Chl | Involved in the formation of serine | 44.93  /8.52 | 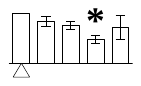 |
| Tocopherol metabolism (2) | | | | | |
| Predicted protein, containing cd05121, activator of bc1 complex kinase and similar protein domain (ABC1K)* | BAK08115 | Chl | Phosphorylated tocopherol cyclase | 78.47  /6.07 | 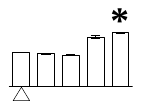 |
| Os11g0216300, containing cd05121, activator of bc1 complex kinases and similar protein domain (ABC1K)* | NP_001067509 | Chl | Phosphorylated tocopherol cyclase | 82.89  /7.6 | 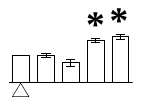 |
| **Stress and defense (8)** | | | | | |
| Thioredoxin peroxidase (TPx) | AAC78473 | Chl | ROS scavenging | 28.13  /6.33 | 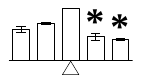 |
| Predicted protein, 2-Cys peroxiredoxin BAS1, chloroplastic-like (Prx)* | BAJ98505 | Chl | ROS scavenging | 28.23  /6.33 | 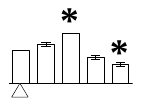 |
| Predicted protein, containing PLN02879, L-ascorbate peroxidase domain (APX)* | BAJ87776 | Chl | ROS scavenging | 37.45  /7.57 | 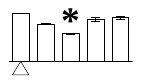 |
| Chloroplast glutathione reductase (GR) | ABQ53155 | Chl, Mit | ROS scavenging | 50.61  /6.17 | 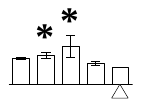 |
| Predicted protein, containing cd03041, glutathione S-transferase, N-terminal domain (GST)* | BAJ99765 | ^#^Chl, Cyt | ROS scavenging | 36.41  /8.98 | 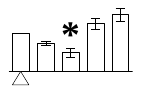 |
| Chloroplast lipocalin (CHL) | ABB02408 | Chl | Prevent lipid peroxidation | 36.85  /5.74 | 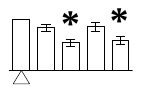 |
| Hypothetical protein SORBIDRAFT_07g015170, acclimation of photosynthesis to environment 1 (APE1)* | XP_002445398 | Chl | PSII light energy balance | 28.00  /8.91 | 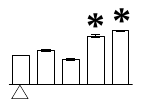 |
| Light-induced protein 1-like (LIP) | ABA61130 | Chl | Stress response | 14.15  /4.54 | 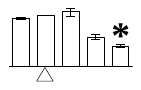 |
| **Membrane and transporting (8)** | | | | | |
| Actin | AAW78915 | ^#^Chl | Chloroplast movement | 41.73  /5.23 | 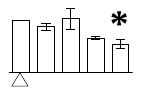 |
| Hypothetical protein OsJ_16821, actin* | EEE62039 | ^#^Chl | Chloroplast movement | 42.93  /5.24 | 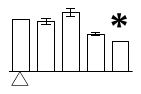 |
| Predicted protein, containing pfam04755, plastid-lipid-associated protein (PAP)* | BAJ88297 | Chl | Protect thylakoid membrane from oxidative damage | 33.22  /5.84 | 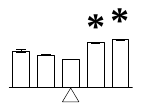 |
| Predicted protein, containing pfam04755, plastid-lipid-associated protein domain (PAP)* | BAJ86880 | Chl | Protect thylakoid membrane from oxidative damage | 32.67  /6.66 | 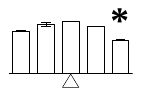 |
| Unknown protein, protein curvature thylakoid 1A (CURT1A)* | XP_002320843 | Chl | Induce thylakoid membrane curvature | 16.55  /9.23 | 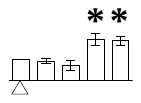 |
| Unknown protein, protein curvature thylakoid 1A (CURT1A)* | BAK02001 | Chl | Induce thylakoid membrane curvature | 16.88  /5.53 | 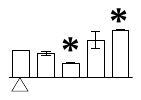 |
| Predicted protein, containing PLN03209, protein Tic 62 domain (Tic62)* | BAJ87171 | Chl | Protein translocation | 57.23  /9.18 | 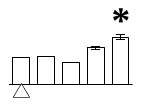 |
| Hypothetical protein SORBIDRAFT_07g007870, preprotein translocase subunit SecY (SecY)* | XP_002445291 | Chl | Protein translocation | 59.36  /9.74 | 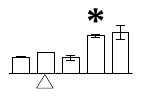 |
| **Signaling (1)** | | | | | |
| Predicted protein, calcium sensing receptor, chloroplastic (CAS)* | BAJ94250 | Chl | Signal transduction | 39.92  /9.46 | 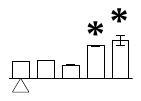 |
| **Gene expression, protein synthesis and turnover (16)** | | | | | |
| Unnamed protein product, 31 kDa ribonucleoprotein (RNP)* | CBU92445 | Chl | Transcription | 33.12  /4.6 | 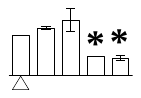 |
| Predicted protein, containing cd01586, aconitase A catalytic domain (ACO)* | BAJ85661 | Chl | Transcription | 106.92  /6.79 | 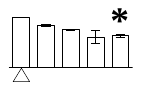 |
| Predicted protein, containing cd00552, ribosome-associated inhibitor A domain (RaiA)* | BAJ98981 | Chl | Protein synthesis | 32.87  /5.78 | 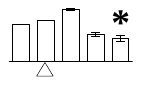 |
| Os03g0315800, containing cd05692, 30S ribosomal protein S1 domain (30S RP)* | NP_001049938 | Chl | Protein synthesis | 41.93  /4.56 | 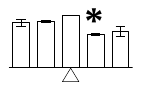 |
| Predicted protein, 50S ribosomal protein L1 (50S RP)* | BAJ90836 | Chl | Protein synthesis | 37.47  /8.25 | 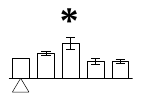 |
| Predicted protein, 50S ribosomal protein L6 (50S RP)* | BAJ92513 | Chl | Protein synthesis | 24.15  /10.13 | 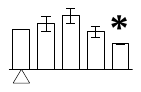 |
| Predicted protein, 50S ribosomal protein L6 (50S RP)* | BAJ93276 | Chl | Protein synthesis | 25.18  /10.08 | 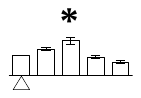 |
| 50S ribosomal protein L9 (50S RP) | Q8L803 | Chl | Protein synthesis | 21.58  /9.86 | 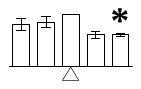 |

Table S6 *(continued from previous page.)*

| **Protein name^(^*^a^*^)^** | **Accession no.^(^*^b^*^)^** | **Loc^(c)^** | **Biological function^(^*^d^*^)^** | **MM(kD)/p*I*^(^*^e^*^)^** | **PA^(^*^f^*^)^** |
| --- | --- | --- | --- | --- | --- |
| Predicted protein, 50S ribosomal protein L9 (50S RP)* | BAK02423 | Chl | Protein synthesis | 22.03  /9.85 | 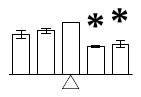 |
| Predicted protein, 50S ribosomal protein L12-2 (50S RP)* | BAJ91881 | Chl | Protein synthesis | 18.15  /5.6 | 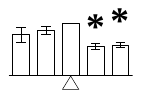 |
| Hypothetical protein SORBIDRAFT_04g024850, elongation factor Tu (EF-Tu)* | XP_002452390 | Chl | Protein synthesis | 50.72  /6.06 | 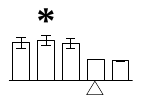 |
| Unnamed protein product, stromal 70 kDa heat shock-related protein (Hsp70)* | CBC02994 | Chl | Protein folding | 73.11  /5.18 | 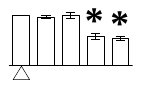 |
| Uncharacterized LOC100280354, stromal 70 kDa heat shock-related protein (Hsp70)* | NP_001146752 | Chl | Protein folding | 66.47  /4.83 | 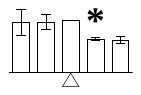 |
| Predicted protein, containing cd10747, chaperone protein DnaJ-like domain (DnaJ)* | BAJ93457 | Chl | Protein folding | 51.90  /9.68 | 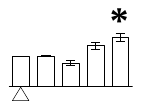 |
| Predicted protein, thylakoid lumenal 15 kDa protein (TL15)* | BAJ97105 | Chl | Protein folding | 20.92  /6.06 | 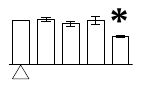 |
| Hypothetical protein SORBIDRAFT_09g000350, containing cd01926, peptidyl-prolyl cis-trans isomerase CYP20-2 domain (PPIase)* | XP_002439092 | Chl | Protein folding | 26.19  /9.25 | 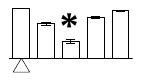 |
| **Miscellaneous and function unknown (6)** | | | | | |
| Predicted protein, containing cd00158, rhodanese homology domain (RHOD)* | BAJ98664 | Chl | Miscellaneous | 64.66  /5.15 | 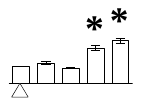 |
| Predicted protein, thylakoid rhodanese-like protein (RHOD)* | NP_001131286 | Chl | Miscellaneous | 49.43  /5.25 | 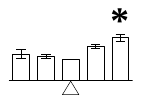 |
| Predicted protein, containing cd05265, SDR_a1 domain (SDR-a1)* | XP_002442964 | Chl | Function unknown | 42.18  /8.88 | 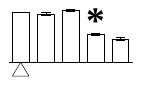 |
| Hypothetical protein OsJ_35887, containing cd05265, SDR_a1 domain (SDR-a1)* | EEE53110 | Chl | Function unknown | 41.53  /8.59 | 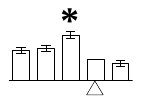 |
| Predicted protein, containing cd05243, SDR_a5 domain (SDR-a5)* | BAJ91782 | Chl | Function unknown | 42.44  /8.89 |  |
| Predicted protein, containing cd05243, SDR_a5 domain (SDR-a5)* | BAJ91568 | Chl | Function unknown | 32.93  /6.76 |  |

*Note: ^a^*The name and functional categories of the proteins identified by LC-MS/MS. Protein names marked with an asterisk (*) have been edited by us depending on searching against NCBI non-redundant protein database for functional domain. The abbreviations for the protein names are indicated in the bracket after protein names.

*^b^*Database accession numbers from NCBInr.

*^c^*Subcellular location (Loc) of the identified proteins predicted by softwares (YLoc, LocTree3, Plant-mPLoc, ngLOC, and ChloroP). Only the consistent predictions from at least two tools were accepted as a confident result. Pounds (#) indicate the subcellular localizations were predicted based on literatures listed in Supplemental Information Table S9. Chl, chloroplast; Cyt, cytoplasm; Mit, mitochondria.

*^d^*The biological function of the identified proteins.

*^e^*Theoretical molecular mass (kDa) and p*I* of identified proteins.

*^f^*The protein abundances (PA) under corresponding treatments compared with conditions marked with triangles (Δ). The asterisks (*) indicate significant differences (p < 0.05). Error bars indicate ± standard deviation. Five columns from left to right indicated the treatments of 0, 150 and 200 mM Na_2_CO_3_ for 12 and 24 h, respectively.

Abbreviations: 1,3-BPG, 1,3-bisphosphoglyceric acid; GAP, glyceraldehyde 3-phosphate; GGDR, geranylgeranyl diphosphate reductase; GGPP, geranylgeranyl pyrophosphate; LHCII, light harvesting complex II; Mg-pro IX, Mg-proporphyrin IX; MgCy, magnesium-protoporphyrin IX monomethyl ester [oxidative] cyclase; Phytyl-PP, phytyl pyrophosphate; R5P, ribose 5-phosphate; ROS, reactive oxygen species; Ru5P, ribulose-5-phosphate; RuBP, ribulose-1,5-bisphosphate; S7P, sedoheptulose 7-phosphate; Xu5P, xylulose 5-phosphate.
