## Supplemental Table 8 for "Na_2_CO_3_-Responsive Photosynthetic and ROS Scavenging Mechanisms in Chloroplasts of Alkaligrass Revealed by Phosphoproteomics"

| **Protein name^(^*^a^*^)^** | **Accession no.^(^*^b^*^)^** | **Loc^(c)^** | **Biological function^(^*^d^*^)^** | **Peptide sequence (m/z, charge, score)^(^*^e^*^)^** | **Ratio** **± S.D.^(^*^f^*^)^** | |
| --- | --- | --- | --- | --- | --- | --- |
|  |  |  |  |  | **150mM/0mM** | **200mM/0mM** |
| **Photosynthesis (25)** | | | | | | |
| **Chlorophyll a/b binding protein (11)** | | | | | | |
| Light harvesting complex I chlorophyll a-b binding protein (Lhca2) | CAA59049 | Chl | Light harvesting | APERPIWFPG**S^57^**TPPPWLDG**S^66^**LPGDFGFDPWGLGSDPESLR (1166.01, 4, 12) | 0.66±0.08 | 0.60±0.04 |
| Light harvesting complex I chlorophyll a-b binding protein (Lhca5) | XP_003618083 | Chl | Light harvesting | QSL**S^66^**YLDGSLPGDFGFDPLGLSDPEGTGGFIEPR (1195.87, 3, 16) | 1.83±0.01 | 1.22±0.28 |
| Light harvesting chlorophyll a-b binding protein of LHCII type 1-like (Lhcb1) | CDI44335 | Chl | Light harvesting, state transition | VAGGPLGEVVDPLYPGG**S^193^**LDPLGLADDPEAFAELK (746.06, 6, 13) | 1.34±0.15 | 1.65±0.20 |
| Light harvesting chlorophyll a-b binding protein of LHCII type 1-like (Lhcb1) | EMS50795 | Chl | Light harvesting, state transition | VLYLGPLSGEPP**S^68^**YLTGEFPGDYGWDTAGLSADPETFAK (1246.35, 4, 12) | 1.93±0.34 | 1.78±0.32 |
|  |  |  |  | VLYLGPLSGEPP**S^68^**YL**T^71^**GEFPGDYGWD**T^82^**AGLSADPETFAK (1246.32, 4, 14) | 0.44±0.05 | 0.42±0.04 |
| Light harvesting chlorophyll a-b binding protein of LHCII type 1-like (Lhcb1) | EMT11232 | Chl | Light harvesting, state transition | VLYLGPL**S^63^**GDPP**S^68^**YLTGEFPGDYGWD**T^82^**AGLSADPETFAK (1246.82, 4, 12) | 0.36±0.16 | 0.47±0.13 |
| Light harvesting chlorophyll a-b binding protein of LHCII type 1-like (Lhcb1) | EMT29003 | Chl | Light harvesting, state transition | VLYLGPLSGEPP**S^68^**YLNGEFPGDYGWD**T^82^**AGL**S^86^**ADPETFAK (1247.32, 4, 13) | 0.33±0.08 | 0.40±0.10 |
|  |  |  |  | VLYLGPLSGEPPSYLNGEFPGDYGWDTAGL**S^86^**ADPETFAK (1247.10, 4, 12) | 0.47±0.04 | 0.55±0.06 |
| Light harvesting chlorophyll a-b binding protein of LHCII type 1-like (Lhcb1) | CAA32109 | Chl | Light harvesting, state transition | VLYLGPLSGREPP**S^65^**YLTGEFPGDYGWDTAGLSADPETFAK (1247.61, 4, 13) | 0.45±0.01 | 0.55±0.07 |
| Light-harvesting chlorophyll a-b binding protein CP29, chloroplastic (CP29) | 1908421A | Chl | PSII disassembly, energy dissipation | LAQNLAGEIIG**T^108^**RFEDADVK (716.39, 4, 11) | 1.94±0.49 | 1.46±0.14 |
| Light-harvesting chlorophyll a-b binding protein CP29.2, chloroplastic (CP29) | CDI44415 | Chl | PSII disassembly, energy dissipation | PAEYLQYDVDSLDQNLAQNLAGEIIG**T^108^**R (1164.24, 3, 13) | 1.32±0.26 | 1.66±0.15 |
| Light-harvesting chlorophyll a-b binding protein CP29.2, chloroplastic (CP29) | XP_003562892 | Chl | PSII disassembly, energy dissipation | PAEYLQYDPD**S^95^**LDQNLAQNLAGEVIGTRFEDADIK (1537.76, 3, 10) | 17.46±10.55 | 16.77±7.96 |
| Light-harvesting chlorophyll a-b binding protein CP26, chloroplastic (CP26) | EMT03206 | Chl | Energy dissipation | TGALLLDGN**T^165^**LNYFGN**S^172^**IPINLILAVVAEVVLVGGAEYYR (1114.07, 4, 13) | 1.60±0.07 | - |
|  |  |  |  | TGALLLDGNTLNYFGN**S^172^**IPINLILAVVAEVVLVGGAEYYR (1201.63, 4, 19) | 1.15±0.36 | 1.74±0.14 |
| **Photosystem II related protein (5)** | | | | | | |
| Photosystem II subunit S (PsbS) | XP_003564708 | Chl | Energy dissipation | GIL**S^122^**QLNLETGIPIYEAEPLLLFFILFTLLGAIGALGDR (1207.64, 4, 14) | 1.30±0.19 | 1.80±0.13 |
| Photosystem II 43 kDa protein (CP43) | ABC02751 | Chl | PSII core protein | TLFNG**T^20^**FVLAGR (709.84, 2, 8) | 1.68±0.10 | 1.03±0.12 |
| Photosystem II reaction center protein H (PsbH) | P69555 | Chl | D1 maturation and incorporation | A**T^3^**QTVEDSSKPR (1044.01, 2, 14) | 2.60±1.04 | 3.28±0.50 |
|  |  |  |  | A**T^3^**Q**T^5^**VEDSSKPRPK (1308.68, 2, 14) | 3.69±1.05 | - |
| Predicted protein, oxygen-evolving enhancer protein 1, chloroplastic (PsbO)* | CCO16140 | Chl | Photosynthetic oxygen evolution, PSII D1 repair | GGSTGYDNAVALPAR**S^264^**DADDLQKENNK (954.97, 4, 13) | 0.97±0.02 | 0.62±0.01 |
| Predicted protein, containing pfam11493, thylakoid soluble phosphoprotein of 9 kDa domain (TSP9)* | BAJ97488 | Chl | State transition | VDGPAPSAGG**T^87^**ASR (834.92, 2, 14) | 1.42±0.08 | 1.74±0.05 |
| **Calvin cycle (9)** | | | | | | |
| Expressed protein, ribulose bisphosphate carboxylase/oxygenase activase (RCA)* | ABA95524 | Chl | Activate RuBisCO | GLAYDI**S^71^**DDQQDITR (1199.59, 2, 9) | 0.87±0.17 | 0.60±0.02 |
| Ribulose-1,5-bisphosphate carboxylase/oxygenase large subunit (RBL) | ABB03412 | Chl | Combines CO_2_ to produce 3-phosphoglycerate | GLDF**T^181^**KDDENVNSQPFMR (859.71, 3, 13) | 1.33±0.08 | 1.56±0.06 |
|  |  |  |  | V**T^27^**PQPGVPPEEAGAAESSTGTWTTVWTDGLTSLDR (1000.47, 4, 12) | 4.97±0.23 | 1.23±0.25 |
| Ribulose-1,5-bisphosphate carboxylase/oxygenase large subunit (RBL) | CAB85674 | Chl | Combines CO_2_ to produce 3-phosphoglycerate | AAFRVTPRPGVPPEEAGAAVAAES**S^62^**TGTWTTVWTDGLTSLDR (1203.10, 4, 12) | 1.59±0.01 | 1.10±0.38 |
| Ribulose-1,5-bisphosphate carboxylase/oxygenase large subunit (RBL) | CAA94018 | Chl | Combines CO_2_ to produce 3-phosphoglycerate | PGVPPEEAGAEVAAES**S^53^**TGTWTTVWTDGLTSLDR (1291.28, 3, 9) | 1.78±0.19 | 1.48±0.48 |
| Ribulose-1,5-bisphosphate carboxylase/oxygenase large subunit (RBL) | CAC04358 | Chl | Combines CO_2_ to produce 3-phosphoglycerate | VTPQPGVPAEEAGAAVDAE**S^51^**STGTWTTVWTDGLTSLDR (1420.37, 3, 13) | 2.00±0.66 | 1.34±0.02 |
| Ribulose-1,5-bisphosphate carboxylase/oxygenase large subunit (RBL) | AAG43944 | Chl | Combines CO_2_ to produce 3-phosphoglycerate | V**T^36^**PQPGVPPGGAGAAVAAES**S^55^**TGTWTTVWTDGLTSLDR (1392.66, 3, 15) | 1.77±0.22 | 1.26±0.21 |
|  |  |  |  | VTPQPGVPPGGAGAAVAAE**S^54^S^55^**TGTWTTVWTDGLTSLDR (1044.50, 4, 13) | 1.91±0.29 | 1.17±0.01 |
| Ribulose-1,5-bisphosphate carboxylase/oxygenase large subunit (RBL) | CAA93205 | Chl | Combines CO_2_ to produce 3-phosphoglycerate | VTPQPGVPAEEAGAAVAAE**S^54^**STGTW**T^60^**TVWTDGLTSLDR (795.56, 6, 14) | 0.51±0.14 | 0.80±0.07 |
|  |  |  |  | VTPQPGVPAEEAGAAVAAES**S^55^**TGTWTTVWTDGLTSLDR (1473.04, 3, 12) | 2.56±0.54 | 2.53±0.90 |
| Ribulose-1,5-bisphosphate carboxylase/oxygenase large subunit (RBL) | AFA27686 | Chl | Combines CO_2_ to produce 3-phosphoglycerate | VTPQPGVPPEEAGAAVAGEIG**T^69^**W**T^71^**TVWTDGLTSLDR (1040.50, 4, 16) | 2.17±0.51 | 2.74±0.18 |
|  |  |  |  | VTPQPGVPPEEAGAAVAGEIGTW**T^71^T^72^**VWTDGLTSLDR (1386.99, 3, 15) | 2.16±0.03 | 2.44±0.17 |
| Glyceraldehyde-3-phosphate dehydrogenase A, chloroplastic (GAPDH) | EMT31124 | Chl | Reversibly converts 1,3-BPG to GAP | GDS**S^92^**PLEVIAINDTGGVK (820.77, 3, 17) | 1.72±0.08 | 0.99±0.53 |
| **Carbohydrate and energy metabolism (12)** | | | | | | |
| **Energy metabolism (3)** | | | | | | |
| ATP synthase subunit alpha, chloroplastic | EMS64844 | Chl | ATP synthesis | NPLIAAA**S^9^**VIAAGLAVGLA**S^21^**IGPGVGQGTAAGQAVEGIAR (1243.97, 3, 13) | 1.18±0.37 | 1.62±0.13 |

**Table S8 Na_2_CO_3_-responsive phosphoproteins in alkaligrass leaves**

Table S8 (continued from previous page.)

| **Protein name^(^*^a^*^)^** | **Accession no.^(^*^b^*^)^** | **Loc^(c)^** | **Biological function^(^*^d^*^)^** | **Peptide sequence (m/z, charge, score)^(^*^e^*^)^** | **Ratio ± S.D.^(^*^f^*^)^** | |
| --- | --- | --- | --- | --- | --- | --- |
|  |  |  |  |  | **150mM/0mM** | **200mM/0mM** |
| ATP synthase CF1 alpha subunit | AEJ10071 | Chl | ATP synthesis | **T^43^**GLGQVMSGELVEFAEGTR (793.72, 3, 12) | 2.01±0.06 | 1.80±0.05 |
| ATP synthase beta subunit | ABR67212 | Chl | ATP synthesis | GFQLIL**S^445^**GELDALPEQAFYLVGNIDEASTK (959.00, 4, 11) | 1.06±0.03 | 1.97±0.38 |
| **Glycolysis (3)** | | | | | | |
| Glyceraldehyde-3-phosphate dehydrogenase (GAPDH) | AFS33113 | Cyt | Reversibly converts 1,3-BPG to GAP | FGIVEGLMTTVHAM**T^184^**ATQK (681.35, 4, 8) | - | 1.90±0.14 |
| Hypothetical protein, phosphoglycerate kinase, cytosolic (PGK)* | XP_006300324 | ^#^Cyt | Reversibly catalyzes 3-PG to produce 1,3-BPG | LSELLGVEVVMAND**S^98^**IGEEVQKLVAALPEGGVLLLENVR (1129.11, 4, 12) | 1.57±0.01 | 1.16±0.15 |
| 2,3-bisphosphoglycerate-independent phosphoglycerate mutase (PGAM) | XP_003564482 | Cyt | Catalyzes the interconversion of 3PGA and 2PGA | AHGTAVGLPSDDDMGN**S^81^**EVGHNALGAGR (1036.13, 3, 21) | 1.63±0.11 | 1.17±0.12 |
| **Pyruvate metabolism (1)** | | | | | | |
| Phosphoenolpyruvate carboxylase (PEPC) | BAD36412 | Cyt | Catalyzes HCO_3_^-^ and PEP to form oxaloacetate | LS**S^18^**IDAQLR (693.87, 2, 13) | 2.25±0.23 | 1.28±0.11 |
| **Starch and sucrose metabolism (4)** | | | | | | |
| Phosphoglucomutase, cytoplasmic (PGM) | EMT08717 | ^#^Cyt, Chl | Reversibly converts G1P to G6P | ATGAFILTA**S^99^**HNPGGPTEDFGIK (1103.91, 3, 13) | 1.58±0.03 | 0.85±0.06 |
| UDP-glucose 6-dehydrogenase (UGDH) | EMT06309 | Cyt | Reversibly converts UDP-glucose to UDP-glucuronate | FDWDHPVHLQPM**S^337^**PTTTK (942.47, 3, 12) | 0.56±0.02 | 0.60±0.01 |
| UDP-glucose 6-dehydrogenase (UGDH) | EMT21987 | ^#^Cyt | Reversibly converts UDP-glucose to UDP-glucuronate | FDWDHPMHLQPT**S^393^**PTAVK (699.60, 4, 12) | 0.70±0.02 | 0.60±0.02 |
| Sucrose-phosphate synthase (SPS) | EMS63629 | ^#^Cyt, Nuc | Catalyzes UDP-glucose and F6P to form UDP and S6P | SDDATEVSETD**S^638^**PGDSLR (1132.99, 2, 24) | 0.61±0.01 | 0.65±0.01 |
| **Other glycometabolism (1)** | | | | | | |
| Phosphoglycerate kinase, chloroplastic (PGK) | XP_003568189 | Chl | Reversibly catalyzes 3-PG to produce 1,3-BPG | PGVVALDEAVTVGSV**T^481^** (797.39, 2, 10) | 1.12±0.01 | 0.51±0.11 |
| **Stress and defense (2)** | | | | | | |
| Predicted protein, containing pfam12481, aluminium induced protein domain (AIP)* | XP_004960054 | Chl | Stress response | QVAHAPQELN**S^18^**PR (915.96, 2, 18) | 2.05±0.15 | 2.87±0.03 |
| Wheat cold induced 16 | BAN63108 | Cyt, Nuc | Stress response | QY**S^76^**SGGTEK (823.40, 2, 14) | 0.64±0.02 | 0.71±0.04 |
|  |  |  |  | QYS**S^77^**GGTEK (548.94, 3, 11) | 0.61±0.05 | 0.80±0.03 |
|  |  |  |  | GAA**S^130^**LSGK (689.88, 2, 11) | 1.21±0.47 | 0.39±0.03 |
| **Membrane and transporting (4)** | | | | | | |
| Hypothetical protein, fructokinase-like 2, chloroplastic (FLN)* | EMS47290 | ^#^Chl, Nuc | Chloroplast thylakoids development | VAEQL**S^113^**DDEGEDQSK (1169.55, 2, 14) | 0.58±0.01 | 0.76±0.06 |
| Zinc finger protein VAR3, chloroplastic (VAR3) | EMS67681 | Chl | Chloroplast development | SD**S^1098^**QVFLFANSK (677.69, 3, 12) | 1.69±0.18 | 1.29±0.06 |
| Predicted protein,villin-2-like* | BAJ91166 | ^#^Chl, Nuc | Actin reverse polymerization | AAAVAALSSVLTAEQSG**S^414^**SDNLR (868.11, 3, 22) | 1.43±0.32 | 1.61±0.12 |
| Hypothetical protein, containing pfam00335, tetraspanin family protein domain* | EMS54049 | PM | Intracellular trafficking | AMNKPAEYD**S^200^**DDEIIGTAR (934.12, 3, 13) | 0.86±0.07 | 1.52±0.02 |
| **Signaling (8)** | | | | | | |
| Predicted protein, containing cd05574, catalytic domain of phototropin-like* | AAC05084 | ^#^PM | Photoreceptor | DEDPLLD**S^384^**DDERPESFDDELR (964.42, 3, 13) | 0.53±0.07 | 1.06±0.18 |
| Calmodulin 3 (CaM) | AGW21711 | Cyt | Ca^2+^ sensor | TVMR**S^8^**LGQNPTEAELQAMINEVDADGNGTIDFPEFLNLMAR (1169.81, 4, 13) | 1.29±0.55 | 1.62±0.08 |
|  |  |  |  | SLGQNP**T^14^**EAELQAMINEVDADGNGTIDFPEFLNLMAR (1463.35, 3, 9) | 1.90±0.22 | 2.46±0.87 |
|  |  |  |  | SLGQNPTEAELQAMINEVDADGNG**T^32^**IDFPEFLNLMAR (1469.03, 3, 20) | 2.59±0.45 | 2.16±0.49 |
| Unknown, calmodulin (CaM)* | ABR18064 | Cyt | Ca^2+^ sensor | SLGQNPTEAELQDMI**S^54^**EVDADGNGTIDFPEFLNLMAR (1468.68, 3, 19) | 3.07±0.30 | 2.36±0.07 |
| Calmodulin (CaM) | AAR99410 | Cyt | Ca^2+^ sensor | SLGQNPTEAELQDMINEVDADGNGTIDFPEFLNL**T^73^**AR (1468.69, 3, 23) | 2.53±0.47 | 2.00±0.14 |
| Inactive receptor kinase-like | XP_004970381 | PM | Signal transduction | EEW**T^564^**AEVFDVDLLR (702.67, 3, 8) | 1.05±0.61 | 2.12±0.25 |
| Phytosulfokine receptor kinase (PSKR) | AAT01376 | PM | PSK signaling | A**S^345^**AEVLGK (731.91, 2, 14) | 0.61±0.01 | 1.28±0.32 |
| Hypothetical protein, containing cd01459, VWA copine domain* | EMT11740 | Nuc | Cellular signaling | SS**S^390^**FGQQTSGFQQSDSFKQR (1002.79, 3, 13) | 1.71±0.02 | 1.56±0.03 |
|  |  |  |  | SS**S^390^**FGQQTSGFQQSD**S^403^**FKQR (1002.79, 3, 13) | 2.03±0.28 | 1.70±0.07 |
| Auxin-repressed 12.5 kDa protein (ARP1) | ABA95233 | ^#^Cyt, Nuc | Hormone signaling | SLGANLFDRPQPN**S^108^**PTVYDWLYSDETR (1175.89, 3, 16) | 0.50±0.08 | 1.08±0.18 |
| **Gene expression, protein synthesis and turnover (11)** | | | | | | |
| Hypothetical protein, histone H2A* | XP_002465460 | Nuc | DNA modification | G**T^60^**GAPVYLAAVLEYLAAEVLELAGNAAR (1449.74, 2, 18) | - | 2.36±0.37 |

Table S8 (continued from previous page.)

| **Protein name^(^*^a^*^)^** | **Accession no.^(^*^b^*^)^** | **Loc^(c)^** | **Biological function^(^*^d^*^)^** | **Peptide sequence (m/z, charge, score)^(^*^e^*^)^** | **Ratio ± S.D.^(^*^f^*^)^** | |
| --- | --- | --- | --- | --- | --- | --- |
|  |  |  |  |  | **150mM/0mM** | **200mM/0mM** |
| Histone H2A.6-like | XP_003569004 | Nuc | DNA modification | VG**S^55^**GAPV**Y^60^**LAAVLEYLAAEVLELAGNAAR (782.62, 4, 13) | 0.85±0.00 | 0.64±0.01 |
|  |  |  |  | PV**Y^60^**LAAVLEYLAAEVLELAGNAAR (967.52, 3, 16) | 1.37±0.11 | 1.97±0.10 |
|  |  |  |  | PVYLAAVLE**Y^67^**LAAEVLELAGNAAR (991.20, 3, 13) | 1.69±0.05 | 1.27±0.06 |
| Histone H2AX-like | XP_006479831 | Nuc | DNA modification | VGAGAPVYL**S^57^**AVLEYLAAEVLELAGNAAR (829.93, 4, 13) | 1.06±0.11 | 1.52±0.02 |
| Hypothetical protein, histone H2AX-like* | ESW34361 | Nuc | DNA modification | GSGSPVYL**S^61^**AVLEYLAAEVLELAGNAAR (967.49, 3, 14) | 1.37±0.11 | 1.97±0.09 |
| 30S ribosomal protein 1, chloroplastic | XP_003559644 | Chl | Protein synthesis | EWQTAAAAAFSE**S^189^**DVDEEEDEDELVEVIGAEDEETVLTK (1243.07, 4, 9) | 0.91±0.27 | 0.63±0.02 |
| 60S ribosomal protein L13-1 | EMT10064 | Cyt | Protein synthesis | AGDS**T^156^**PEELANATQVQGDYMPIAR (973.46, 3, 19) | 1.54±0.01 | 1.41±0.06 |
| Uncharacterized protein, pfam06273 plant specific eukaryotic initiation factor 4B* | XP_003562957 | Nuc | Protein synthesis | GVDALASDLEKT**S^372^**PVGR (801.76, 3, 15) | 0.83±0.20 | 0.59±0.10 |
| Translation initiation factor 5A | ABB29987 | Cyt | Protein synthesis | SD**T^4^**DEHHFESK (1031.47, 2, 13) | 0.62±0.04 | 0.75±0.12 |
| Predicted protein, heat shock cognate 70 kDa protein* | BAJ86014 | Cyt | Protein folding | VQDLLLLDVTPL**S^406^**QGLE (1119.10, 2, 7) | - | 0.48±0.06 |
| Cytosolic heat shock protein 90 | AAP87284 | Cyt | Protein folding | EI**S^220^**DDEDEEEK (1013.45, 2, 17) | 0.90±0.03 | 1.66±0.11 |
| 26S protease regulatory subunit 6A homolog | XP_003570339 | Cyt | Protein degradation | SS**S^4^**PTPAPAAAPAAPMAVDETEDDQL**S^28^**SMSTDDIVR (1264.20, 3, 8) | 0.79±0.02 | 0.42±0.05 |
| **Unknown (1)** | | | | | | |
| Hypothetical protein | EMT31442 | Nuc | Function unknown | GPSGFPGAGGSGSD**S^44^**DEPQEYYTGGEK (1107.83, 3, 16) | 0.64±0.03 | 0.92±0.26 |

*Note: ^a^*The name and functional categories of the proteins identified by LC-MS/MS, protein names marked with an asterisk (*) have been edited by us depending on searching against NCBI non-redundant protein database for functional domain. The abbreviations for the protein names are indicated in the bracket after protein names.

*^b^*Database accession number from NCBInr.

*^c^*Protein subcellular localization (Loc) predicted by softwares (YLoc, Cello, Plant-mPLoc, ngLOC, and ChloroP). Pounds (#) indicate the subcellular localizations were predicted based on literatures. Chl, chloroplast; Cyt, cytoplasm; Mit, mitochondria; Nuc, nucleus; PM, plasma membrane.

*^d^*Biological function of the identified proteins.

*^e^*Identified phosphopeptide, the phosphorylated amino acids sites are highlighted. Precursor m/z, charge, and peptide identification scores are indicated in the bracket after each phosphopeptide.

*^f^*The average ratios of phosphopeptides between control (0mM) and treatment (150mM and 200mM for 24h) were quantified according to the isobaric tags for relative and absolute quantification (iTRAQ) from three biological replicates, error bar indicates ± standard deviation (S.D.). ‘-’ indicate that the phosphorylation site without quantitative information in three replicates.

1,3-BPG, 1,3-bisphosphoglycerate; 3-PG, 3-phosphoglycerate; 2PGA, 2-phosphoglycerate; 3PGA, 3-phosphoglycerate; F6P, fructose 6-phosphate; G1P, glucose 1-phosphate; G6P, glucose 6-phosphate; GAP, glyceraldehyde 3-phosphate; PEP, phosphoenolpyruvate; PSK, phytosulfokine; S6P, sucrose-6-phosphate; UDP-glucose, uridine diphosphate glucose.
