## Supplemental Table 14 for "Na_2_CO_3_-Responsive Photosynthetic and ROS Scavenging Mechanisms in Chloroplasts of Alkaligrass Revealed by Phosphoproteomics"

**Table S14 Primers used for overexpression of *PtFBA* in *Synechocystis* 6803**

| **Name** | **Primer sequence (5’–3’)** | **Purpose** |
| --- | --- | --- |
| Primers used for *PtFBA* gene clone | | |
| *PtFBA*-FP-1 | CAATGGCGTCTGCTACTCTCCTCA | Amplification of full length cDNA of *Ptfba* |
| *PtFBA*-RP-1 | CGTCAGTTCAGGTCGCTCCACT |  |
| Primers used to construct the P*psbAII-Ptfba* expression vector | | |
| *PtFBA*-FP-2 | GGAATTCCATATGGCGTCTGCTACTCT | Amplification of *Ptfba* encoding gene fragment |
| *PtFBA*-RP-2 | GGAATTCCATATGCTAGAGATTGGTCAGTTC |  |
| *slr0168*-FP | GAGTAGTTCCCTCAACACCAGT | Segregation analysis |
| *slr0168*-RP | TTCCAGGCCACATTGTTGTC |  |
| Primers used for RT-PCR | | |
| *PtFBA*-FP-3 | GGAATTCCATATGGCGTCTGCTACTCT | *Ptfba* transcript |
| *PtFBA*-RP-3 | GGAATTCCATATGCTAGAGATTGGTCAGTTC |  |
| *16 S rRNA*-FP | CGACTGCTAATACCCAATGTGC | *16 S rRNA* transcript |
| *16 S rRNA*-RP | GTCCCTCAGTGTCAGTTTCAGC |  |
